# A role for fibroblast and mural cell subsets in a nerve ligation model of neuropathic pain

**DOI:** 10.1101/2024.12.11.627455

**Authors:** Sara Villa-Hernandez, Julia Vlachaki Walker, Zoe Hore, Laura Fedele, Irene Zebochin, Yuening Li, Harvey Davis, Takashi Kanda, Fumitaka Shimizu, Leonie S. Taams, Franziska Denk

## Abstract

Neuropathic pain is a particularly intractable type of chronic pain that can result from physical nerve damage due to surgery or entrapment. Here, we present data which suggest that a particular subclass of fibroblast and mural cells may be implicated in the sensory neuron dysfunction that is characteristic of this pain state.

In a mouse model of traumatic painful neuropathy, we used cell sorting, nerve tissue clearing and RNA sequencing to study mesenchymal lineage cells. With cell sorting (n = 4 mouse nerves) and tissue clearing (n = 5), we show that fibroblasts and mural cells positive for the platelet-derived growth factor receptor beta (*Pdgfrb*) gene are increased in number for at least two months post-nerve damage. Moreover, single cell RNA sequencing data (n = 4) from our own lab and those of three other laboratories reveal that Pdgfrb+ cells express high levels of known and putative pro-algesic mediators. Bulk sequencing of sorted Pdgfrb+ fibroblasts (n = 10) and Pdgfrb+/Cd146+ mural cells (n = 11) further indicate that many of these mediators are upregulated in neuropathy.

We go on to demonstrate that a human nerve pericyte line releases a selection of these pro-algesic mediators at protein level. Moreover, conditioned media from stimulated human pericytes induces intra-cellular changes in human induced pluripotent stem cell derived sensory neurons (n = 5 independent differentiations); these changes (phosphorylation of the transcription factor signal transducer and activator of transcription 3, STAT3) have been previously linked to sensory neuron activation.

In summary, our data indicate that mesenchymal cell abnormalities should be considered when developing novel strategies to tackle neuropathic pain.

**Graphical abstract:** 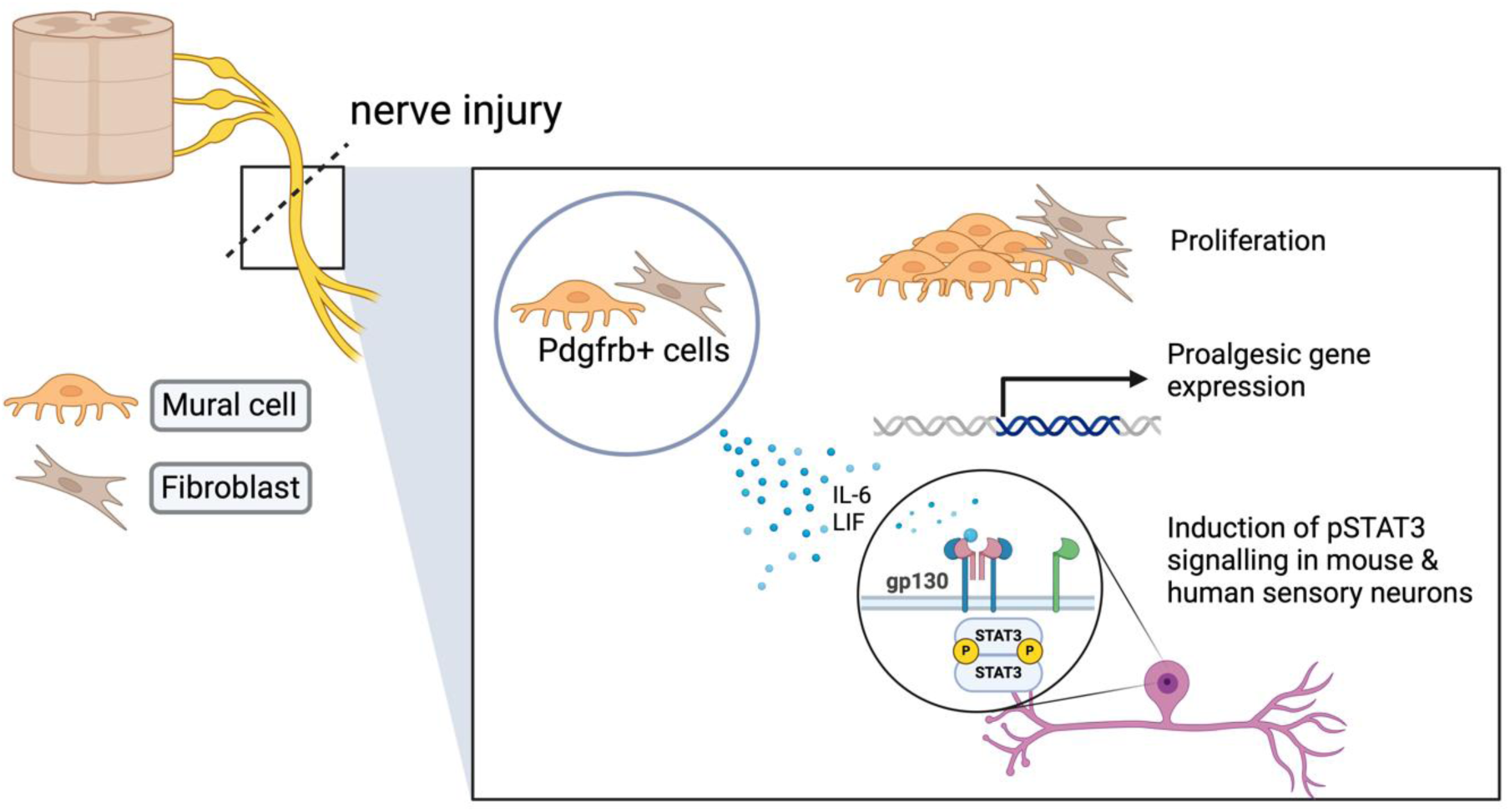

## Introduction

Neuropathic pain results from injury or disease of the nervous system, for example, when a nerve is trapped in sciatica or damaged due to diabetes. It is a particularly unpleasant kind of pain, often described with adjectives such as ‘shooting’, ‘burning’ or ‘stabbing’. Unfortunately, it is also rather common; a recent epidemiological study estimates that 9% of individuals in the UK live with the condition, with middle- to older-age women particularly affected (Baskozos et al., 2023).

Mechanistically, neuropathic pain arises as a result of peripheral and central nervous system hyperactivity (Colloca et al., 2017). In the periphery, this hyperactivity is believed to be significantly driven by the local inflammatory milieu; most work to date has focused on immune cells – suggesting involvement of e.g. macrophages (Gylfadottir et al., 2022; Liang et al., 2020; Sandy-Hindmarch et al., 2024), natural killer cells (Davies et al., 2019) and other adaptive immune cell populations (Davoli-Ferreira et al., 2020; Singh et al., 2022). However, it is becoming clearer that mesenchymal lineage cells like fibroblasts (Shinotsuka and Denk, 2022) are also likely to play significant roles.

From both animal (Singhmar et al., 2020) and human studies (Hofmann et al., 2024; Kreß et al., 2021), evidence is accumulating that certain fibroblast subtypes and the mediators they release can be pro-algesic in the context of neuropathic pain. This makes perfect sense when we consider that fibroblasts are well-known to be crucial drivers of chronic inflammatory states (Parsonage et al., 2005), with tissue-independent, universal sub-types present in pathologically inflamed tissues in conditions as wide-ranging as rheumatoid arthritis, interstitial lung disease, ulcerative colitis and atopic dermatitis (Buechler et al., 2021; Korsunsky et al., 2022).

One subtype of note is a population of NOTCH3+ (neurogenic locus notch homolog protein 3) fibroblasts that has been shown to proliferate in the sub-lining of joint synovial membrane in arthritis (Wei et al., 2020). This population is also positive for marker genes *THY1* (thymus-cell antigen 1), *CCL19* (C-C motif chemokine ligand 19) and *PDGFRB* (platelet-derived growth factor receptor beta); it has been reported to be a universal pathological subset present in inflamed skin, colon, joint and lung (Buechler et al., 2021; Korsunsky et al., 2022). Crucially, Wei and colleagues demonstrated that the identity of this cell sub-population is positionally determined, with signals from perivascular endothelial cells controlling their phenotype as well as that of NOTCH3+ mural cells. These could be either vascular smooth muscle cells, if located around arteries or arterioles, or pericytes, if wrapped around capillaries (Davis and Attwell, 2023).

Mural cells have been very poorly studied in neuropathic pain to date, although pioneering work and proposals by Heike Rittner’s group suggest they could play a role in the blood-nerve barrier breakdown that has been observed in some experimental models of neuropathic pain (Reinhold et al., 2023). They are also thought to be involved in the pathophysiology of conditions that can cause neuropathic pain, such as diabetes (Hamilton et al., 2010; Mateus Gonçalves et al., 2023).

Here, we present novel evidence for the involvement of both pathological fibroblasts and mural cells in neuropathic pain. We show that these cell types contain particularly high levels of known and putative pro-algesic mediators in a mouse model of traumatic neuropathy. We further demonstrate that the number of mural cells in nerve is increased for 2 months after model induction, with significant pain-relevant transcriptional dysregulation observable in both Pdgfrb+ fibroblasts and mural cells. Finally, we show that human nerve pericytes, when stimulated, release pro-algesic mediators at protein level and can induce intracellular activation of induced pluripotent stem cell (IPSC)-derived sensory neurons.

## Materials and Methods

### Animals

We used either C57BL/6J wild-type mice purchased from Charles River, UK, or Pdgfrb-CreERT2-tdTomato mice, with both strains obtained from the Jackson Laboratory: RRID:IMSR_JAX:029684, allele: Tg(Pdgfrb-cre/ERT2)6096Rha/J (Gerl et al., 2015) & RRID:IMSR_JAX:007908, Ai14 tdTomato allele: Gt(ROSA)26Sortm14(CAG-tdTomato)Hze/J (Madisen et al., 2010). Pdgfrb-CreERT2-tdTomato mice were bred in-house at King’s College London to be homozygous for tdTomato and hemizygous for the Pdgfrb-CreERT2 allele. All mice were housed in individually ventilated cages in a 12-hour light/dark cycle with food and water available ad libitum. Experiments were conducted in accordance with the United Kingdom Home Office Legislation (Scientific Procedures Act, 1986) and were approved by the Home Office to be conducted at King’s College London. All mice were adult at the point of nerve injury (i.e. at least 8 weeks of age), with both males and females used as indicated in the individual experimental methods below. To induce tdTomato in Pdgfrb+ cells, tamoxifen was injected once intraperitoneally at 75 mg/kg, one week before surgery. All mice tolerated the surgery well, but during the tissue clearing study, one sham-operated mouse had to be culled due to malocclusion. During the FACS experiment for bulk-sequencing, one mouse had to be excluded as it did not have any tdTomato signal. For single cell sequencing, the n numbers refer to a pool of animals; in all other instances n numbers refer to individual mice.

### Neuropathic pain model

Partial sciatic nerve ligation (PSNL) was carried out according to previously published protocols (Liang et al., 2020). Briefly, after pre-operative subcutaneous administration of 0.1 mg/kg buprenorphine (Henry Schein, Melville, NY, Cat# 988-3245), mice were anaesthetised with isoflurane. An incision was made in the skin over the left thigh of the mouse, and the underlying muscle was blunt dissected. Approximately two-thirds of the sciatic nerve was ligated with a 5.0 VICRYL absorbable suture (Ethicon, Cincinnati, OH, Cat# W9982). The wound was closed with a metal clip (Reflex, 7mm, Fine Science Tools, Foster City, CA, Cat# 12032-07), which was removed 7-10 days after the procedure. Sham surgery was performed in identical fashion, but the nerve was only visually exposed by blunt dissection without being mechanically disturbed. Animals were not randomised into sham or surgery groups, but rather age-matched littermates were chosen for one or the other procedure. Allocation was thus not concealed. The success of surgery and persistence of nerve damage was confirmed by checking that all PSNL mice continued to display the expected postural abnormalities in the operated hind-leg (Liang et al., 2020).

### Flow cytometry

At acute (5 days after PSNL) and chronic (2 months after PSNL) timepoints, mice were transcardially perfused with phosphate buffered saline, after which ipsi- and contralateral sciatic nerves were dissected into F12 solution (Gibco, New York, NY, Cat# 21765-029) and trimmed to a 0.5 cm (data in **Figure 2A**) or a 1 cm piece around the lesion site (all other experiments). In either case, the distal and proximal pieces were kept at the same length (0.25 or 0.5 cm each). The suture was removed at this point, if still present. A single cell solution was generated as described previously (Liang et al., 2020). Cells were stained with the nuclear marker DRAQ5, a live-dead marker and antibodies as outlined in **Supplementary Table 1A&B**. The data were analysed in FlowJo, according to the gating strategies outlined in **Supplementary Figures** 1 **& 2.** Analysis did not proceed blind to condition, but identical gates were used for all samples within an experiment and initially placed with the help of fluorescence-minus one controls. Cell counts were used as the outcome measure and compared between groups using a non-parametric Mann Whitney t-test; sensitivity analysis indicates that with n = 10, this t-test will have an 80% chance to detect effect sizes of d = 1.36 or larger, with alpha at 0.05.

### Single-cell RNA sequencing (scRNA-seq)

At acute (4-6 days after PSNL) and chronic (2 months after PSNL) timepoints, C57BL/6J wild-type mice were transcardially perfused with phosphate buffered saline, after which ipsilateral sciatic nerves were dissected to a 0.5 cm piece around the lesion site, as described above. A single cell solution was generated as described previously (Liang et al., 2020), after which cells were stained with the nuclear marker DRAQ5, the live-dead stain DAPI, PercCP-Cy5-5-conjugated myelin beads, BUV737-CD45 to stain immune cells and BV786-Ter119 to stain erythrocyte precursors. DRAQ5+DAPI-,Myelin-,CD45-,Ter119-singlets were sorted into 1.5 ml tubes, pre-coated with foetal calf serum (FBS), containing 3 µl of 3% BSA in PBS buffer and immediately processed by the NIHR Maudsley Biomedical Research Centre (BRC) Genomics & Biomarker Core Facility at King’s College London for library preparation (single cell 3’ version 3 on a Chromium 10X instrument) and sequencing (Illumina NextSeq system).

Experiments were conducted in 2 batches; see **Supplementary Table 2** for breakdown of individual mice and samples. In the first batch, we processed n = 4 injured and n = 4 sham nerves (50% male/female), collected them two months after surgery and pooled them into two 10X wells (injury vs. sham) in the same FACS, library preparation and sequencing run. In the second batch, we processed n = 4 sham and n = 4 injured nerves each, at two different timepoints: 5 days and 2 months after surgery. 50% of samples were male, 50% female; they were run on the same day and the same 10X chip, pooled by group across 4 wells (sham-day5, PSNL-day5, sham-month2, PSNL-month2). Library preparation and sequencing proceeded at the same time.

The resulting fastq files were aligned to mouse (Mus_musculus.GRCm38) in Google Colabs using kb-bustools (Melsted et al., 2021), with kallisto version 0.46.2 and bustools version 0.40.0. Count matrices were then imported into R and further analysed in Seurat (version 4.3). The R scripts used to generate the figures in this manuscript are available in **Supplementary R-Notebooks 1-3**. Raw data, aligned reads and the corresponding Seurat objects have been deposited on the Gene Expression Omnibus website under accession number GSE283177.

### FACS followed by bulk RNA sequencing

Nerve cells were dissociated from Pdgfrb-CreERT2-tdTomato mice as described in the ‘Single-cell RNA sequencing’ section. This time, we processed ipsilateral and contralateral nerves 5 days and 2 months after PSNL (see **Supplementary Table 2** for a list of animals & sequencing samples). Cells were stained using the panel outlined in **Supplementary Table 1C** and gated according to the strategy shown in **Supplementary Figure 3**. Pdgfrb+ and Pdgfrb+/Cd146+ populations were sorted into 4µl of Smart-seq2 lysis buffer. We sorted a maximum of 100 cells per tube, so as to not overload the solution. For day 5 samples, sequencing libraries were then prepared in batch controlled fashion using the Smart-seq2 protocol described by Picelli and colleagues (Picelli et al., 2014). Some samples failed at FACS and some showed degradation at cDNA level and had to be excluded before further processing. The remaining samples were n = 10 ipsi (4 male/ 6 female) and n = 14 contra (5 male/ 9 female) Pdgfrb+ samples and n = 11 ipsi (3 male/ 8 female) and n = 13 contra (5 male/ 8 female) Pdgfrb+/Cd146+ samples. The over-representation of female samples was due to more female animals having been generated in the litters chosen for these experiments. Sequencing was performed on these samples by Novogene Europe. Reads were aligned to mouse (Mus_musculus.GRCm38) using kallisto version 0.50.1 (Bray et al., 2016). Differential expression analysis was performed using DESeq2 (Love et al., 2014). Ligand-receptor analysis was performed using the ICELLNET package (Massenet-Regad and Soumelis, 2024), as well as in Excel using ligand-receptor pairs published by the Lewis lab (Armingol et al., 2021), specifically those derived from (Shao et al., 2020). Expression levels of receptors on mouse sensory neurons were derived from (Renthal et al., 2020), while expression level of receptors on human sensory neurons were derived from pseudobulk data generated from two separate post-mortem studies (Jung et al., 2023; Nguyen et al., 2021). Fastq files and processed data have been deposited on the Gene Expression Omnibus website under accession number GSE283281. Corresponding R scripts are available in **Supplementary R-Notebook 4**.

### Human IPSC-derived sensory neurons

Sensory neurons were derived from the Kute4 iPSC line (HPSI0714i-kute_4) as detailed in (Li et al., 2024b). Briefly, iPSCs were maintained using StemFlex media in vitronectin coated plates. They were passaged at ∼70% confluency using versene. iPSCs were moved onto GelTrex coated plates prior to being differentiated into sensory neurons using the Chambers protocol (Chambers et al., 2012). On day 11, the immature neurons were replated on GelTrex coated glass coverslips at a density of 75,000 cells per coverslip. They were maintained with twice weekly media changes using N2 media with addition of 25 ng/ml of growth factors (NT3, BDNF, GDNF, NGF) until maturity (day 50). Once a week, the medium was additionally supplemented with GelTrex to prevent cell detachment.

### Human pericyte cell lines

A vial of human placental pericytes was purchased by PromoCell (#C-12980) and a human peripheral nerve pericyte line was obtained under MTA from Profs. Takashi Kanda and Fumitaka Shimizu from Yamaguchi University. Both pericyte lines were maintained in Pericyte Medium (ScienCell, #1201), containing 1% pericyte growth supplement (ScienCell, #1252), 2% FBS (ScienCell, #0010) and 1% penicillin/streptomycin solution (ScienCell, #0503). Media was changed twice weekly, and cells were passaged using Accutase (Gibco, # A11105-01) when 70% confluency was reached.

### Pericyte cytokine treatment

Pericytes (passage 5 for the placental line, passage 26 for the nerve pericyte line) were plated onto 6-well plates at a density of 150,000 cells per well. The following day, media were swapped for serum-starved media (containing Pericyte Basal Media and penicillin/streptomycin only). After 24 hours, the media were again changed to serum starved media with the addition of human recombinant cytokines (10 ng/ml TNFα, BioLegend #570102, 10 ng/ml IL-17, Proteintech #Hz-1113, 10 ng/ml IL-4, BioLegend #574004, 100 ng/ml IFNγ, BioLegend #570202). The cells were then incubated with the cytokines for 4 hours, after which the conditioned media were collected and stored at -80°C until use for Luminex or conditioned media experiments.

### Luminex Assay

Pericyte cell supernatants, either untreated or treated with the cytokines listed above were diluted either 1:2 or 1:100 and ran using a Luminex Flexmap Assay (ThermoFisher, Lot# L152023) to probe for IL-6, NGF, LIF and CCL2 expression. Media alone was also run as a control at a 1:2 dilution.

### Conditioned media incubation and Western Blot

Once mature (Day 50), iPSC derived sensory neurons were incubated for 1 hour with media collected from cytokine-treated human nerve pericytes or control unstimulated pericytes. Following the incubation, cells were rinsed with PBS and harvested using 1x Laemmli Buffer with 10% βMercaptoethanol (ITWReagents, #A1108). The experiment was repeated with 5 independent neuronal differentiations, each treated with a different batch of pericyte conditioned media.

Samples were denatured by heating at 95°C for 5 minutes. The samples and a ladder (Thermo Scientific, #26619) were then loaded on 8-16% Tris-Glycine Mini Protein Gels (Thermofisher, XP08165BOX) and run at 100V for 90 minutes. 30 µl of each sample was loaded on the gel; given that neurons were seeded at the same density throughout trials, protein quantification was not performed. Following separation, proteins were transferred to methanol activated PVDF membranes (Fisher Scientific, #10617554) at 100V for 60 minutes. Membranes were blocked using 5% milk in TBS-Tween for 45 minutes, prior to overnight primary antibody incubation with pSTAT3 (1:1000, Cell Signalling, #9145T) and GAPDH (1:1000, Cell Signalling, #5174). The antibody solutions were made using 1% BSA and 0.1% sodium azide in TBS-Tween. The next day, membranes were washed 3x 10 minutes in TBS-Tween and incubated for 1 hour at room temperature with secondary antibodies (1:5000, Abcam, #ab205718) in TBS-Tween. They were then washed once more 3x 5 minutes in TBS-Tween. Staining was visualized by incubating the membranes with ECL Western Blotting Substrate (GE Healthcare, #RPN2232) and imaged using the iBright Imaging System (Invitrogen). To visualize STAT3 (1:1000, Cell Signalling #12640), membranes were stripped for 15 minutes using ReBlot Plus Strong Antibody Stripping Solution (Millipore, #2504), and the above-described blocking and staining process was repeated. Densitometric analysis was conducted using Fiji to determine pSTAT3/STAT3 ratios.

### Tissue clearing and immunostaining

For whole sciatic nerve staining and tissue clearing, at acute (5 days after PSNL) and chronic (2 months after PSNL) timepoints, Pdgfrb-CreERT2-tdTomato mice were transcardially perfused with 20 ml warm Heparin solution (Merck, #H3149-25KU) diluted to 20U/ml in PBS (Thermo Scientific #BR0014G), followed by 10 ml warm FITC-Albumin-Gelatin: 5% Gelatin (SLS, #G9382-500G), 0.2% FITC-Albumin, (Merck #A9771-100MG) in PBS. Mice were then placed on ice for 30 minutes to set the gelatin. Where still present, the suture was carefully removed, and ipsilateral and contralateral sciatic nerve collected and placed in 4% paraformaldehyde overnight. Nerves were washed in PBS and stored at 4°C until ready to for tissue clearing for which we followed the published CUBIC protocol (Susaki et al., 2014). Briefly, nerves were gently shaken in blocking buffer at 37°C for two days, consisting of 0.3% Triton-X-100 (Sigma, #T-9284) and 2% BSA (Sigma, #A3294-100G) in PBS, with 1 μg/ml DAPI (SLS, #D9542-1M). They were then washed in 0.2% Triton and placed in CUBIC R1 solution: 25% N,N,N’,N’-tetrakis (Sigma, #122262-1L), 15% Triton X-100, 25% Urea (Sigma, #U-6504) in distilled water. Nerves were kept in CUBIC R1 for 5 days, refreshing the solution once during incubation the period. After that, they were transferred into CUBIC R2 solution for 24hr: 50% Sucrose, 25% Urea, 10% Triethanolamine (Sigma, #90279-500ML), 0.1% Triton in distilled water. Nerves were mounted on microscope slides in home-made wells containing CUBIC R2 solution and imaged on a Zeiss LSM 710 confocal microscope using a 20x objective (20x/0.8na).

### Imaging and analysis

To image whole, cleared sciatic nerves, the 20x objective (0.8na) was used, and clearly visible blood vessels were identified based on FITC staining. A z-stack at 2-8 μm intervals was taken for each nerve region selected, capturing the whole thickness of the selected vessel. For sham ipsilateral and PSNL contralateral samples, two distinct regions per nerve were selected to image from. For PSNL ipsilateral samples, one or two images from the area of the lesion and one or two from either side of the lesion were taken. When collecting the first batch of samples for the 5-day timepoint, there were no markings to aid with identification of the orientation of the nerve; therefore, at the 5 day timepoint, we could only define images as captured at the “Site of Lesion” and “Away from Lesion”. In this one batch, we also did not perform any sham surgeries. For later experiments (including a second batch at day 5 post-PSNL), sham surgeries were preformed, and the sciatic nerve was marked using a diagonal incision during collection; this meant that regions upstream and downstream from the lesion could be conclusively identified during imaging. The two batches of the day 5 experiments are shown separately in **Supplementary Figure 4**.

Analysis of the images proceeded in FIJI (Schindelin et al., 2012) by an experimenter blinded to treatment group. In brief, vessel area was determined based on FITC signal. FITC thresholds were set based on vessel visibility. Tdtomato thresholds were set as recommended by the software for the day 5 experiments. For the 2-month imaging data, the thresholds set by the software appeared to differ between experimental conditions. We therefore chose to apply the FITC thresholds to the TdTomato images instead for this timepoint, as these did not appear to be impacted by experimental condition (**Supplementary Figure 5C & G**).

Once thresholds were set, the FITC signal was artificially broadened to identify tdtomato positive cells in close proximity to vessels (putative mural cells). DAPI staining was then used to identify nuclei using the StarDist Plugin (Schmidt et al., 2018). Nuclei which overlapped with putative mural cells (tdtomato/FITC double positive) and putative fibroblasts (tdtomato alone) were exported as regions of interest (ROIs) and further analysed in Excel. ROIs were exported for five z-planes per nerve region, each 8 μm apart, and an average of all measurements taken for each nerve region. Where two images per nerve region were captured and analysed, an average was taken for that region.

For all experiments, the number of nuclei was determined by counting all ROIs with an area of at least 10 μm^2^; the number of mural cells was determined by counting the number of nuclei overlapping with tdTomato-FITC co-expression; while the number of tdTomato positive cells was determined by counting the number of nuclei overlapping with tdTomato expression. The vessels in the analysed images were also classified as arterioles, venules, epineurial and endoneurial capillaries by a blinded experimenter, based on morphology (**Supplementary Table 3**). This was done to ensure the image selection did not bias the results of the analysis. In the chronic injury samples, extensive morphological and vascular disorganisation was observed, which means vessel classification in these conditions may be less accurate.

## Results

### Nerve fibroblasts and mural cell populations contain particularly high levels of known and putative pro-algesic mediators

We performed scRNA-seq on mesenchymal lineage cells in mouse nerves 5 days and 2 months after partial sciatic nerve ligation (PSNL). For this, we dissected nerve and performed cell sorting to select live nucleated events and remove myelin, CD45+ immune cells and Ter119+ erythrocyte precursors before proceeding to 10X Chromium sequencing. The resulting dataset underwent quality control (**Supplementary Figure 6**) and was examined for batch effects (**Supplementary Figure 7**); unsupervised clustering was then performed which indicated the presence of 11 different cell-subtypes (**Figure 1A**). We annotated these based on their marker gene expression (**Figure 1B, Supplementary Figure 8, Supplementary R-Notebook 1**) to be fibroblasts, endothelial cells (EC), Notch3+ mural cells (MC) and Schwann cells (SC), with a myelinating sub-cluster primarily present at day 5 post-PSNL (**Supplementary Figure 7B**). At that early time point, there were also a few immune cells (IC), which presumably escaped CD45+ deselection during sorting. Fibroblast clusters could be split into known universal sub-types positive for collagen type XV alpha 1 chain (*Col15a1*), peptidase inhibitor 16 (*Pi16*) and *Ccl19* (Buechler et al., 2021; Korsunsky et al., 2022). We also observed a cell population positive for claudin 1 (*Cldn1*), which expressed high levels of the glucose transporter Glut1 (*Slc2a1*), a known marker for mesenchymal cells at the blood-nerve barrier (Malong et al., 2023). Looking further into these data, we noticed that well-known and more putative pro-algesic mediators are prominently expressed in fibroblast and mural cell populations, both of which are Pdgfrb+ (**Figure 1C**). These include: interleukin-6 (*Il-6*), nerve growth factor (*Ngf*), C-C motif chemokine ligand 2 (*Ccl2*) and leukemia inhibitor factor (*Lif*). When comparing our data to previously published scRNA-seq datasets on mouse sciatic nerve (Carr et al., 2019; Kalinski et al., 2020; Wolbert et al., 2020), both in naïve condition and after nerve injury (sciatic nerve crush), we found that this was a reproducible result (**Figure 1D**): in all datasets, *Il6*, *Ngf*, *Lif* and *Ccl2* are prominently expressed in either Pdgfrb+ mural cells or fibroblasts.

**Figure 1:**
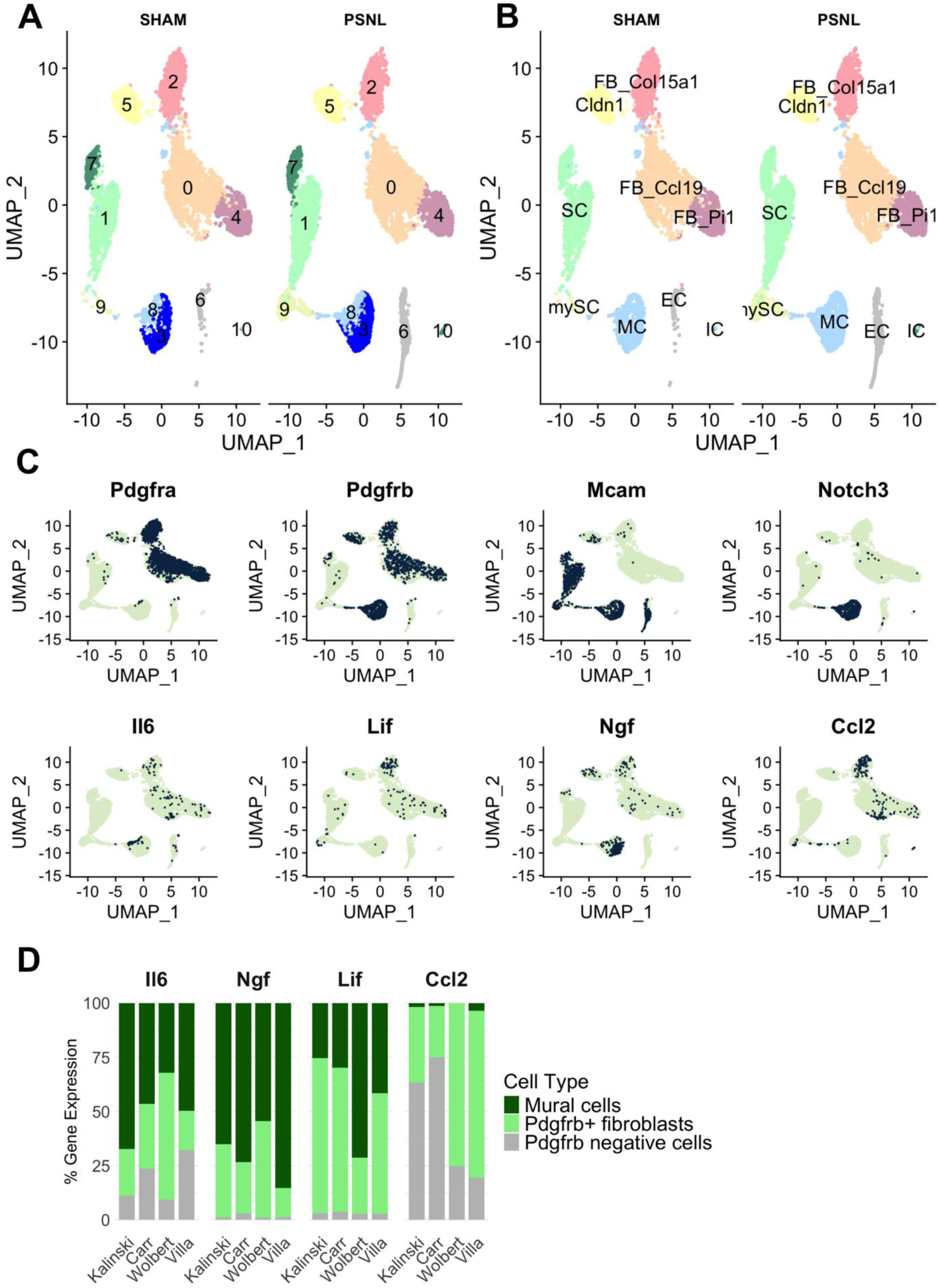
scRNA-seq of mesenchymal cells within mouse nerves reveals fibroblasts and mural cells to be rich in pro-algesic mediators. **A)** Unsupervised clustering indicated that we could distinguish 11 cell clusters which we then annotated according to their marker gene expression as shown in **(B)** (see also **Supplementary Figure 8**). n = 3 per condition (sham vs. partial sciatic nerve ligation, PSNL), across two timepoints (n = 1, day 5 post-PSNL; n = 2 month 2 post-PSNL). Cluster IDs: fibroblasts (FB) positive for Col15a1, Ccl19, Pi16; barrier cells (Cldn1), mural cells (MC); Schwann cells (SC); myelinating Schwann cells (mySC); endothelial cells (EC); immune cells (IC). **C)** Notch3 and Mcam (melanoma cell adhesion molecule = Cd146) double positive mural cells and Pdgfrb+ fibroblasts express known and putative pro-algesic mediators Il-6, Ngf, Ccl2 and Lif. Each dot is a cell. The greener the cell, the higher the expression of a particular gene. **D)** The proportion of pro-algesic mediator gene expression across mouse sciatic nerve cells as identified by our own (Villa) and three other, previously published scRNA-seq datasets (Carr et al., 2019; Kalinski et al., 2020; Wolbert et al., 2020). Notch3/Cd146/Pdgfrb+ mural cells and Pdgfrb+ cells consistently contained higher levels of these mediators than other (Pdgfrb negative) cell populations. (Wolbert et al., 2020). All previously published work did not sort mesenchymal lineage specifically, so Pdgfrb negative cells included immune, endothelial and Schwann cell populations. See **Supplementary R-Notebook 2** and **Supplementary Table 4** for more information on how Pdgfrb+ and negative clusters were selected.

### The number of Cd146+/Pdgfrb+ mural cells increases with nerve injury and remains upregulated for months after model induction

To track what happens to Notch3+ mural cells with injury, we next performed flow cytometry and/or FACS to quantify their numbers in mouse nerve after PSNL. We found that Cd146+ as well as Cd146+/Pdgfrb+ cells were increased 5 days after nerve injury. This was observed across three different experiments performed by different experimenters with distinct flow cytometry panels (**Supplementary Figures 1-3**) and with Pdgfrb as an antibody or genetically labelled using PdgfrbCre-ERT2-tdTomato mice (**Figure 2A-C**). In one of the FACS experiments, we also checked numbers 2 months after nerve injury (**Figure 2D**); once again, Cd146+/Pdgfrb+ numbers were still increased, both in terms of absolute numbers and relative to total live cells (**Supplementary Figure 9**).

**Figure 2:**
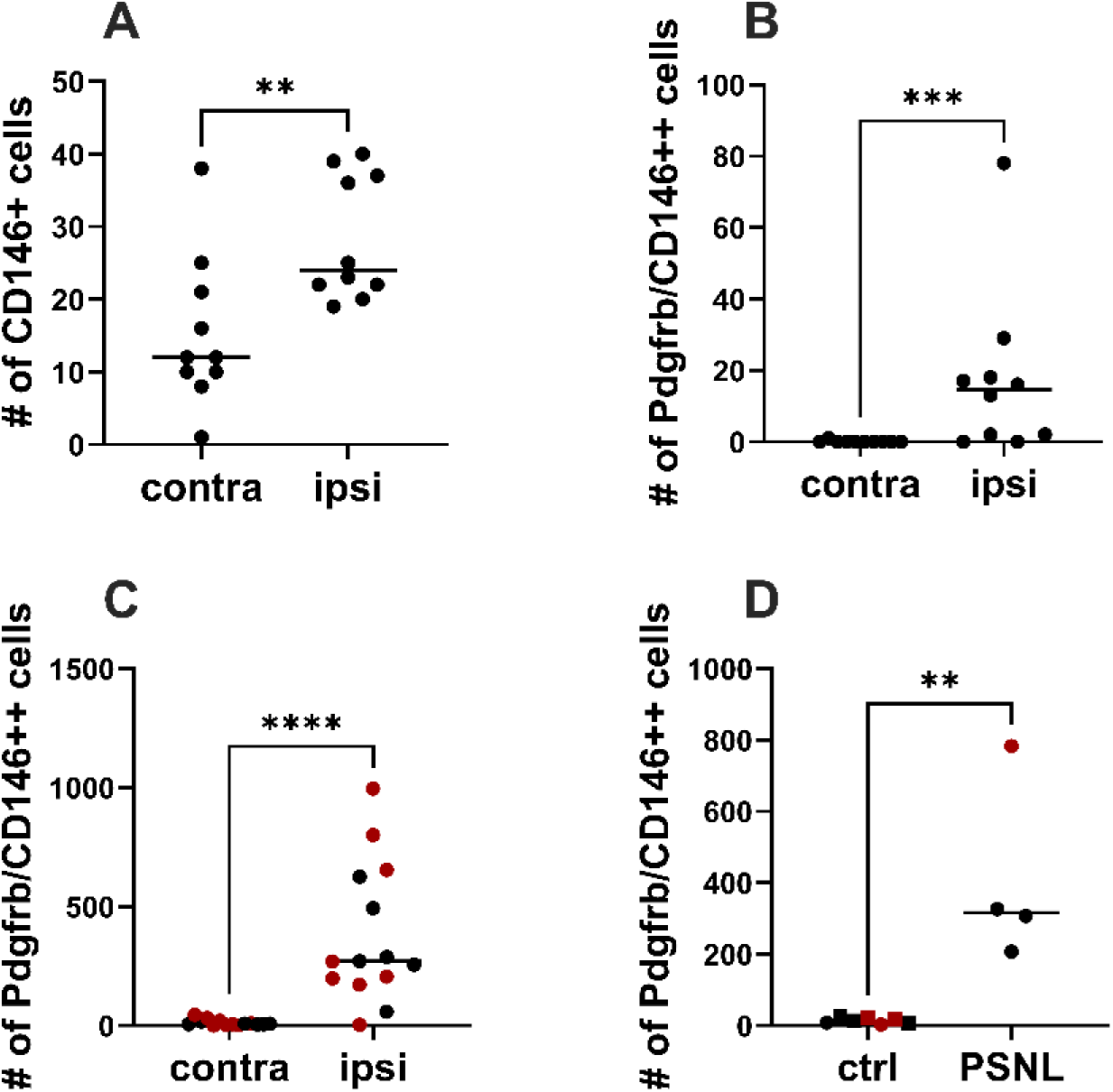
Nerve mural cells are persistently increased upon traumatic nerve injury, as measured by flow cytometry and FACS. Three independent investigators using different mouse lines, antibody panels, gating strategies and flow cytometers/sorters all detected higher numbers of Cd146+ (/PDGFRB+) mural cells in ipsilateral mouse sciatic nerves five days after partial ligation compared to their uninjured, contralateral counterparts (contra). **A)** Male C57BL/6J mice (n = 10) stained with antibody panel in **Supplementary Table 1A. B)** Male C57BL/6J mice (n = 10) stained with antibody panel in **Supplementary Table 1B**. **C)** Male and female Pdgfrb-tdTomato mice (n = 14) stained with antibody panel in **Supplementary Table 1C**. Datasets were analysed with a non-parametric Mann Whitney t-test, p = 0.0026 for A, p = 0.0007 for B, p < 0.0001 for C. See **Supplementary Figures 1-3** for corresponding gating strategies. **D)** Two months post-PSNL, the increase in Cd146+(/Pdgfrb+) double positive cells in PSNL nerves (n = 4) was still apparent compared to uninjured samples (n = 5 sham denoted by squares, n = 2 contralateral denoted by circles). Mann Whitney non-parametric t-test p = 0.0061. Black symbols = male mice, red symbols = female mice.

To further investigate the localisation of mural cells, as well as overall Pdgfrb+ populations in the injured nerve, a whole cleared tissue approach was taken. By perfusing PdgfrbCreERT2-tdTomato mice with FITC-albumin gelatin and staining the sciatic nerve with DAPI before clearing, we were able to examine the number of Pdgfrb+ tdTomato-expressing cells both adjacent to and away from the vasculature in injured mouse nerves (**Figure 3A**); the former will be mural cells (either pericytes or vascular smooth muscle cells), while the latter will consist of Pdgfrb+ endoneurial fibroblasts or Pdgfrb+ perineurial cells. Using this method, we did not observe a significant increase in the number of mural cells 5 days post-PSNL (**Figure 3B**), but we did observed a significant increase in Pdgfrb+ cells, specifically at the site of lesion (**Figure 3C**; p = 0.0157 compared to contralateral nerve, post-hoc comparison after significant Kruskal-Wallis ANOVA).

**Figure 3:**
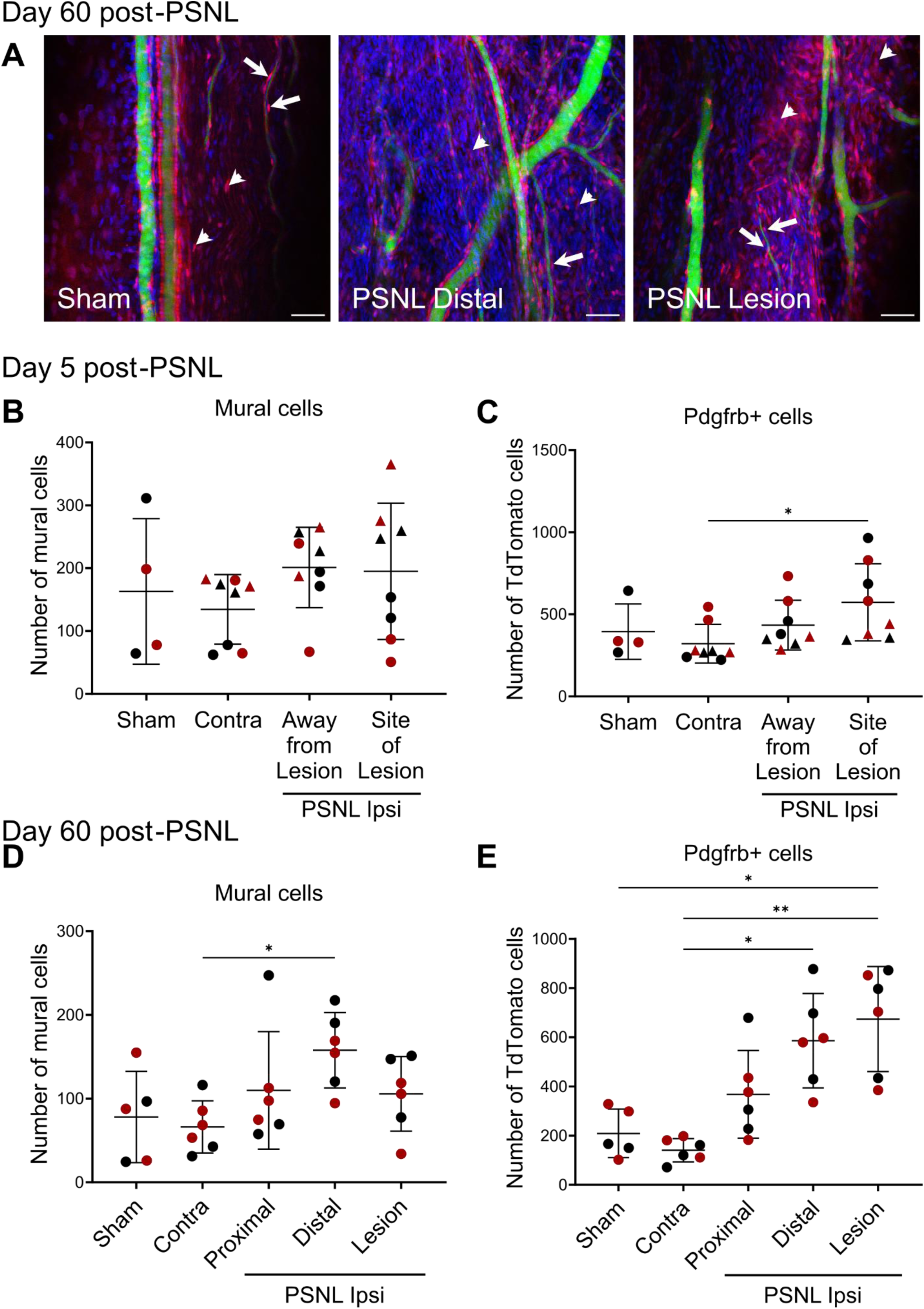
Nerve mural and stromal cells are increased upon traumatic nerve injury, as measured by confocal imaging of cleared sciatic nerve. **A)** Representative maximum intensity projection images of the sciatic nerve of PdgfrbCre x tdTomato mice perfused with FITC-albumin-gelatin 60 days after sham or PSNL surgery. Arrows point to examples of mural cells and arrowheads of Pdgfrb+ cells (scale bar = 50 μm). **B)** Number of mural cells and **C)** Pdgfrb positive (stromal) cells 5 days post PSNL or sham surgery. Experiment carried out in two batches (triangles = batch1, n = 4; circles = batch 2, n = 4). **D)** Number of mural cells and **E)** Pdgfrb positive (stromal) cells 60 days post PSNL or sham surgery (n = 5-6). Datasets were analysed with a non-parametric Kruskal-Wallis one-way ANOVA with Dunn’s multiple comparisons (black symbols = males, red symbols = females).

At day 60 post-PSNL, where we had a higher sample size in a single batch, our results were more clearly in line with our flow cytometry data; we observed a significant increase in the number of mural cells in the distal part of the nerve (**Figure 3D**; p = 0.0318 compared to contralateral nerve, post-hoc comparison after significant Kruskal-Wallis ANOVA). It is important to bear in mind that our images contained a variety of blood vessel types. However, the proportions of different vessel types imaged in each condition are unlikely to be driving the observed injury-induced increase in mural cells: our own in-house observations suggest that arterioles have the highest mural cell density in peripheral nerve followed by endoneurial capillaries; neither of these were disproportionally represented in our images of distal ipsilateral chronic PSNL nerves (**Supplementary Table 3**). At the chronic timepoint, we also observed a significant increase in the number of Pdgfrb+ cells both at the site of lesion (**Figure 3E**; p = 0.0238 compared to sham, p = 0.0019 compared to contralateral) and in the distal part of the nerve (p = 0.0101 compared to contralateral). Finally, total nuclei numbers were consistently increased: in the distal part of the nerve at the 60-day timepoint (p = 0.0025) and at the site of lesion at both timepoints: 5 days (**Supplementary Figure 5A**; p = 0.0009 compared to contralateral nerve) and 60 days (**Supplementary Figure 5E**; p = 0.0228 compared to contralateral, p = 0.0196 compared to sham). This was an expected result, given that we have previously shown that high levels of inflammation persist in nerve months after PSNL surgery (Liang et al., 2020). Our current data suggest that this chronic inflammation is not limited to changes in immune cell numbers and phenotypes, but also encompasses Pdgfrb+ mural cells and fibroblasts. From our immunostaining experiments, it appears that the lesion site and the regions downstream of the lesion are particularly affected, i.e. stromal cell dysregulation is highest in what would likely be areas of Wallerian degeneration (Coleman and Höke, 2020).

### Bulk RNA-seq of Pdgfrb+ mesenchymal cells reveals upregulation of pain-relevant mediators

To obtain well-powered differential gene expression data for Pdgfrb+ fibroblasts and Pdgfrb+/CD146+ mural cells, we performed bulk sequencing on these populations, sorted 5 days after PSNL (see gating strategy in **Supplementary Figure 3**). The resulting sequencing data confirmed at a molecular level that our FACS gating strategy was successful, with Pdgfrb+ fibroblasts expressing significantly higher levels of fibroblast marker genes (e.g. *Pdgfra* = platelet-derived growth factor receptor alpha, *Pi16*, *Pdpn* = podoplanin) and lower levels of mural cell marker genes (*Mcam*, *Notch3*, *Pdgfrb*, **Figure 4A**). Contamination was limited to the occasional Schwann cell transcripts, with no discernible pattern across conditions (**Supplementary Figure 10).**

**Figure 4:**
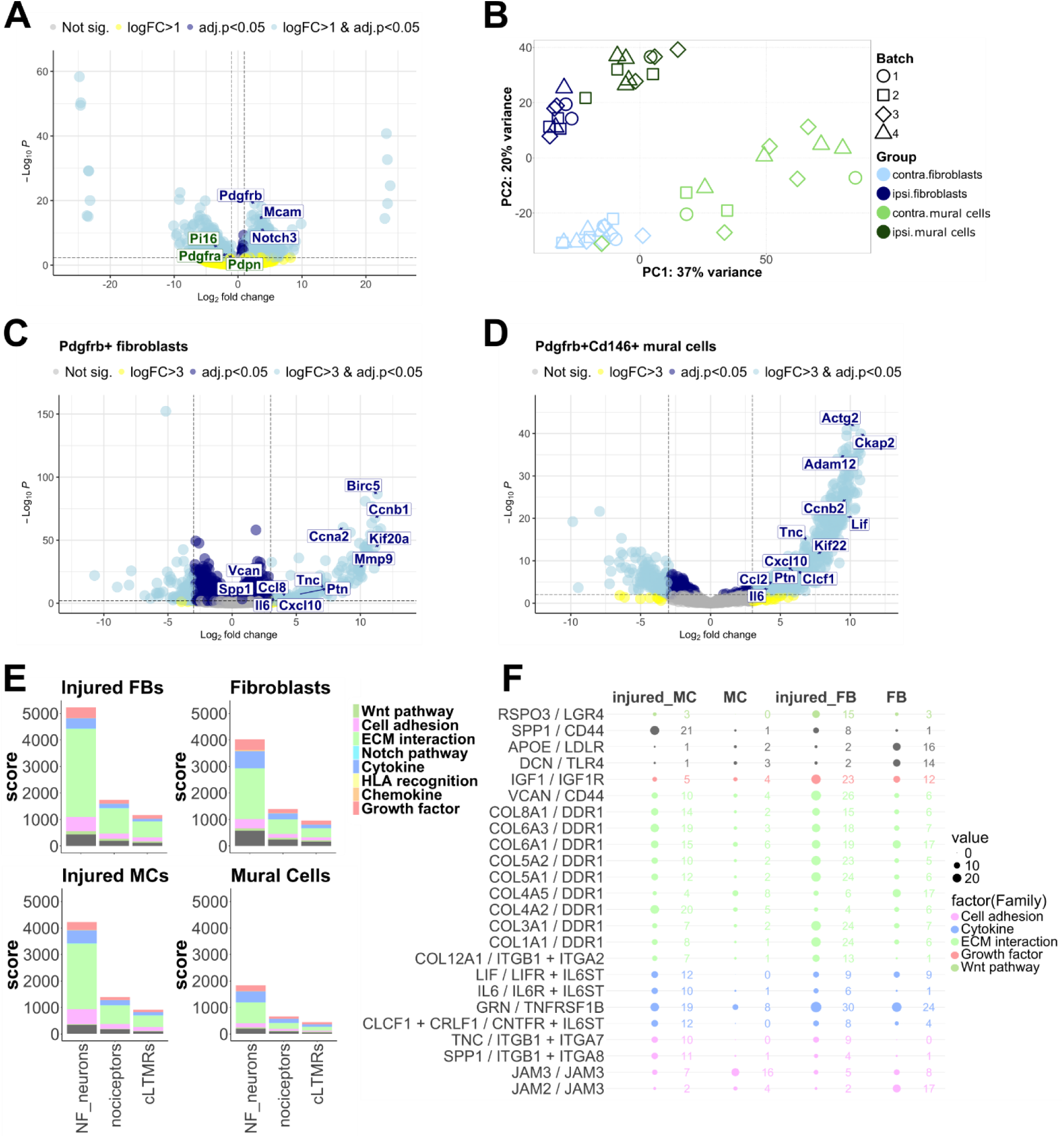
Bulk sequencing analysis of sorted nerve Pdgfrb+ fibroblasts and Pdgfrb+/CD146+ mural cells shows that both populations display significant pain-relevant transcriptional regulation that has potential to affect communication between these cell types and sensory neurons. **A)** Differential expression confirmed our sorting strategy was successful, with mural cell markers (blue labels) significantly upregulated in Pgfrb+/CD146+ double positive samples and fibroblast markers (green labels) significantly upregulated in Pdgfrb+ single positive samples. **B)** Principal component analysis of all samples analysed in the experiment. Cell type and experimental state (ipsi vs. contra = injured vs. contralateral nerve) are clearly distinguishable, even at whole transcriptome level. **C** Volcano plots of genes regulated at log_2_FC > 3 (= eightfold difference compared to control) in ipsilateral compared to contralateral nerve, 5 days after PSNL for fibroblasts and & **D)** mural cells. Each dot is a gene; occasional labels are displayed, all of them for genes that are significantly regulated at adjusted p < 0.05 (i.e. dots are above the horizontal dotted line). In light blue are genes that are significantly regulated more than 8x (to the left or right of the vertical dotted lines); in dark blue are those significantly regulated at lower fold changes. In yellow and grey are genes which are not significantly regulated. **E**) Ligand-receptor interactions between mouse sensory neurons and injured or uninjured nerve Pdgfrb+ fibroblasts (FBs) or mural cells (MCs). Communication scores are plotted for different ligand-receptor families. The higher the score, the more matching ligand-receptor pairs are expressed in a particular family in the sender (fibroblasts/mural cells after nerve injury) and receiver cells: neurofilament positive sensory neurons (NF), nociceptors and C-low threshold mechanoreceptors (cLTMRs). The grey colour indicates other interactions that do not fall within the most commonly occurring families. **F)** Plotting the top ligand receptor pairings that are differentially expressed between injured & uninjured mural cells (MCs) and fibroblasts (FBs) five days after PSNL. The colours denote the function of each ligand-receptor pair, the size of the bubbles and corresponding numbers represent the communication scores.

Nerve injury had a marked effect on gene expression in both cell types, detectable even at whole transcriptome level via principal component analysis (**Figure 4B**). Differential expression analysis using the algorithm DESeq2 (Love et al., 2014) revealed that PSNL induced 1171 upregulated and 1368 downregulated genes in fibroblasts at adjusted p < 0.05, once a stringent expression cut-off was applied (**Supplementary Table 5**). Similarly, 1073 genes were up- and 496 genes downregulated in mural cells. Regulation of the latter was more prominent, with 958 genes significantly upregulated more than eightfold, while for fibroblasts this was limited to 159 (**Figure 4C&D**). Amongst the genes more highly expressed with injury were known, and putative, pro-algesic mediators, like *Il6*, *Ccl2*, *Lif*, cardiotrophin like cytokine factor 1 (*Clcf1*), osteopontin (*Spp1*) and matrix metalloproteinase 9 (*Mmp9*). *Il6*, *Lif* and *Clcf1* are all upstream of glycoprotein 130 receptor signalling, which has been previously involved in neuropathic pain (Kalpachidou et al., 2022).

Ligand-receptor analysis was performed against a mouse dataset, containing gene expression levels of different sensory neuron subpopulations pre- and post-spared nerve injury (Renthal et al., 2020). Results suggested that both nerve fibroblasts and mural cells have the potential to communicate with sensory neuron sub-populations through a variety of pathways, with extracellular matrix interactions, cytokines, growth factors and cell adhesion molecules particularly over-represented (**Figure 4E**, **Supplementary Table 6** & **Supplementary R-Notebook 4**). Of particular interest, it appears that gene regulation after nerve injury enables significantly more ligand-receptor interactions between mural cells and sensory neurons (p = 0.0000056); this is true across sensory neuron subtypes (**Supplementary R-Notebook 4**). Known and putative pro-algesic mediators once again stood out, with *Il6*, *Lif*, and *Clcf1* enriched in mural cells upon PSNL and able to bind sensory neurons directly (**Figure 4F**). In addition to this more formal analysis, we also derived known ligand-receptor pairs for all genes that were significantly (adj.p < 0.05) up- or down-regulated in mural cells and fibroblasts (**Supplementary Table 7)**. For these, we chose our ligand-receptor pairs based on expression of corresponding receptors in published human post-mortem sensory neuron RNA sequencing datasets (Jung et al., 2023; Nguyen et al., 2021). This was to enhance the translational relevance of our findings. Once again, we identified very similar pairings that included *Il6*, *Lif*, *Clcf1* and *Ccl2*. Thus, our results were in agreement across species and across two independent ligand-receptor databases (Massenet-Regad and Soumelis, 2024; Shao et al., 2020).

Finally, we checked whether our genes of interest were enriched in particular mural cell subsets. It is known that brain pericytes and vascular smooth muscle cells are transcriptionally diverse in both mouse (Zeisel et al., 2018) and human (Yang et al., 2022). We also found this to be the case in mouse sciatic nerve, across all the datasets we examined (**Supplementary R-Notebook 3**). In our own scRNA-seq data, we detected three pericyte and two vascular smooth muscle cell (vSMC) populations (**Figure 5**). The former were positive for expected markers like vitronectin (*Vtn*) and *Abcc9* (ATP binding cassette subfamily C member 9), while the latter expressed contractile genes actin alpha 2 (*Acta2*) and transgelin (*Tagln*) at the expected higher levels. Pericytes could further be divided into those that lacked expression of *Adgfr5* (adhesion G-protein coupled receptor F5, possibly resembling the PER2 population described by Zeisel et al., 2018) and those that did not (resembling PER1 & 2 populations in Zeisel et al., 2018). There was also a very small number of *Acta2*-negative pericytes, possibly thin-strand capillary pericytes which reportedly express no or relatively little *Acta2* (Dalkara et al., 2024). Among vSMCs, a specific population was visible only at day 5 post-PSNL or sham surgery; it resembled the arteriolar SMCs described by Yang et al., 2022, expressing higher levels of *Slit3* (slit guidance ligand 3) and *Ctnna3* (catenin alpha 3).

**Figure 5:**
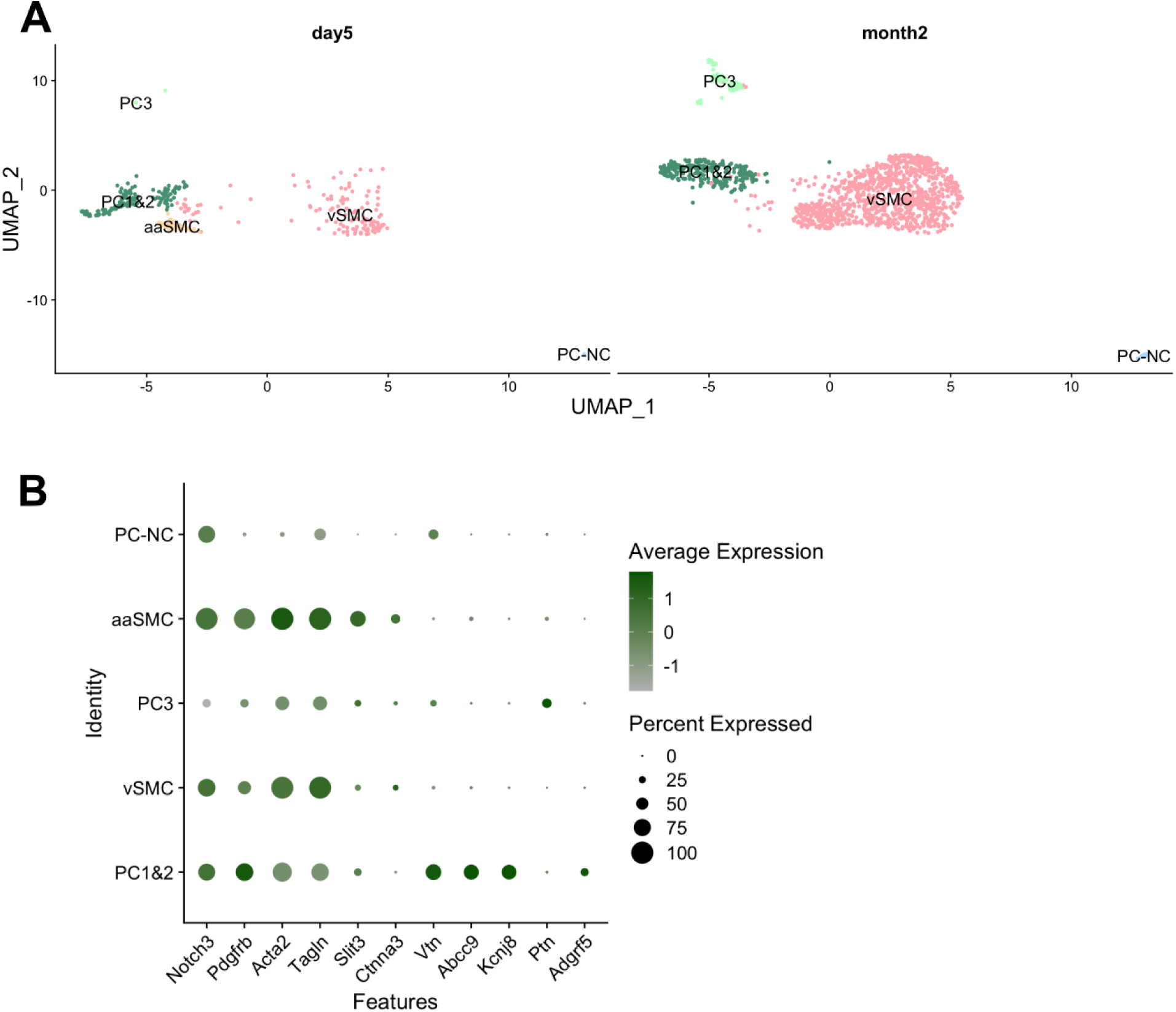
Mural cells in sciatic nerve display the expected molecular diversity. Mouse mural cells sub-cluster into five groups, resembling cell types previously described in the brain, with one (aaSMC) detected only day 5 after PSNL or sham surgery (A). Acta2 & Tagln high vascular smooth muscle cells (vSMC), arteriolar SMCs enriched for Slit3 and Ctnna3 (aaSMC), Vtn & Abcc9 & Kcnj8 positive pericytes (PC1&2), Adgrf5 negative (PC3) and Acta2 negative pericytes (PC-NC). Expression of marker genes is plotted in B. No prominent differences were detected between sham and PSNL groups at either timepoints.

At month 2, there was no consistent difference in the proportion of the various mural cell populations across injury states; at day 5, the PSNL sample did appear distinct from its sham control with increased PC1&2 and decreased vSMCs. It remains to be determined whether this is a replicable finding (**Supplementary Table 5**). Furthermore, some of the genes which we found to be upregulated in mural cells upon nerve injury appeared differentially enriched in one cell type over another, with an over-representation of *Il6*, *Lif*, *Clcf1*, *Ccl2* and tenascin-C (*Tnc*) in pericytes, while *Ngf* appeared more prominent in vascular smooth muscle cells (**Figure 6**). This was statistically significant for *Il6*, *Clcf1*, *Ccl2* and *Tnc* (**Supplementary Table 8**). *Tnc* has been linked to inflammation in painful diseases like arthritis, with its knockout reported to be protected from erosive arthritis in mice (Midwood et al., 2009).

**Figure 6:**
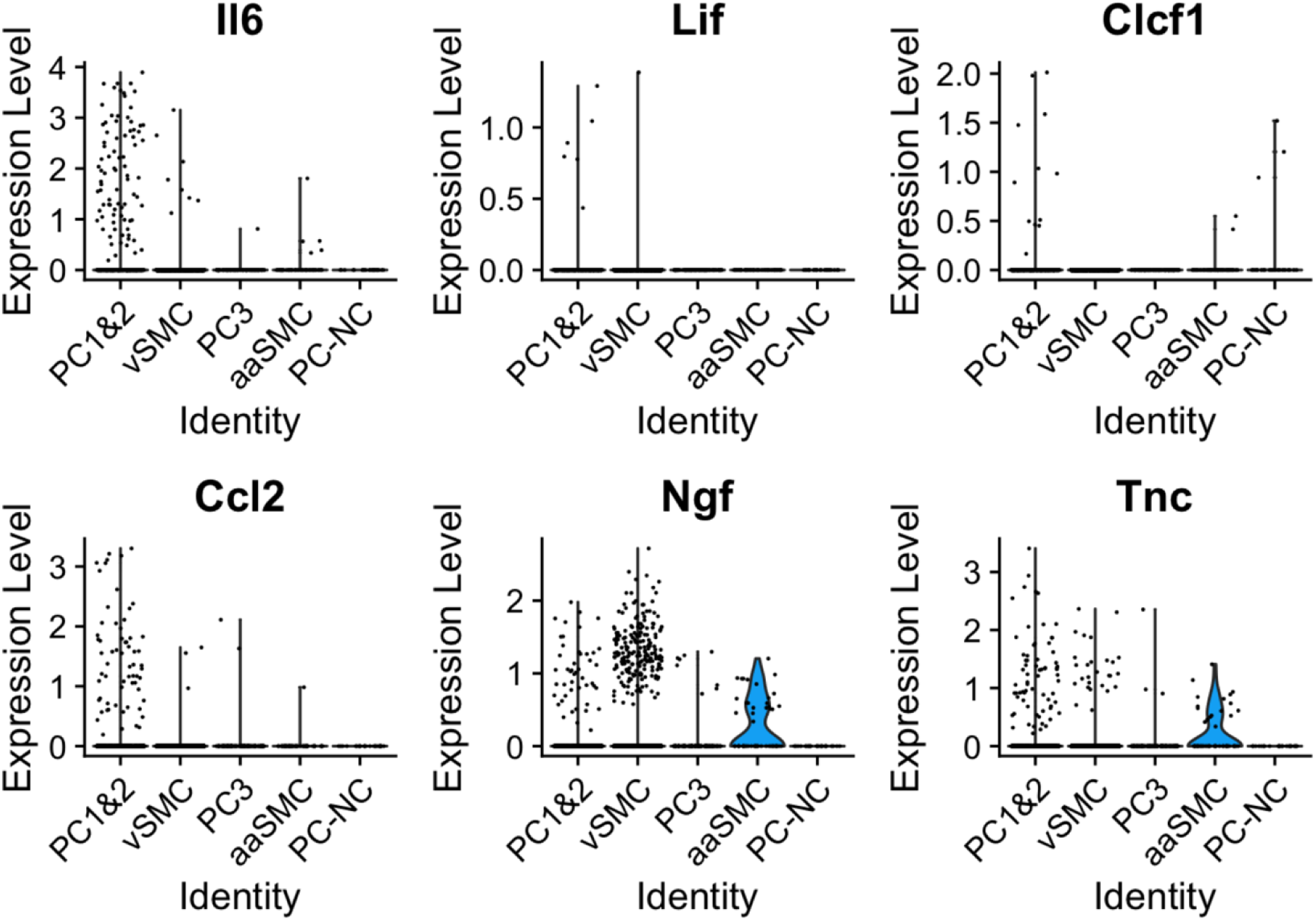
Some of the known and putative pro-algesic mediators appear differentially expressed in specific mural cell subtypes. Plotted here is a selection of mediators, the majority of which we identified as differentially expressed in pericytes in our bulk sequencing experiment (Il6, Lif, Clcf1, Ccl2 and Tnc). Their expression patterns appeared to differ across mural cell populations, with significant enrichment of Il6, Ccl2 and Tnc in PC1&2 pericytes, and Clcf1 in Acta2 negative pericytes (PC-NC).

### A human nerve-derived pericyte line expresses known pro-algesic mediators

Our findings from mouse indicate that Pdgfrb+ cells in sciatic nerve express high levels of pro-algesic mediators that are further up-regulated with nerve injury. We also present evidence to indicate that this effect is particularly striking in Pdgfrb+/Cd146+ mural cells, which are likely equivalent to the pericytes/vascular smooth muscle cells we visualised with tissue clearing; their numbers remained increased two months after PSNL.

To test whether pericytes have similar pro-algesic potential in human, we studied two immortalized pericyte cell lines. Due to ease of availability, we started with a commercially available line derived from human placenta. In an effort to increase our translational relevance, we then also used a second line derived specifically from human sciatic nerve to match the tissue we were studying in mouse (Shimizu et al., 2011).

We cultured these cells and analysed the media they released using Luminex protein analysis. IL-6, NGF, and CCL2 were detectable in both pericyte lines, while LIF was absent (**Figure 7**). NGF and IL-6 appeared constitutively expressed, while CCL2 could only be detected once pericytes were stimulated with pro-inflammatory cytokines TNF or TNF and IL-17. IL-6 expression was also highly up-regulated in these two stimulation conditions.

**Figure 7:**
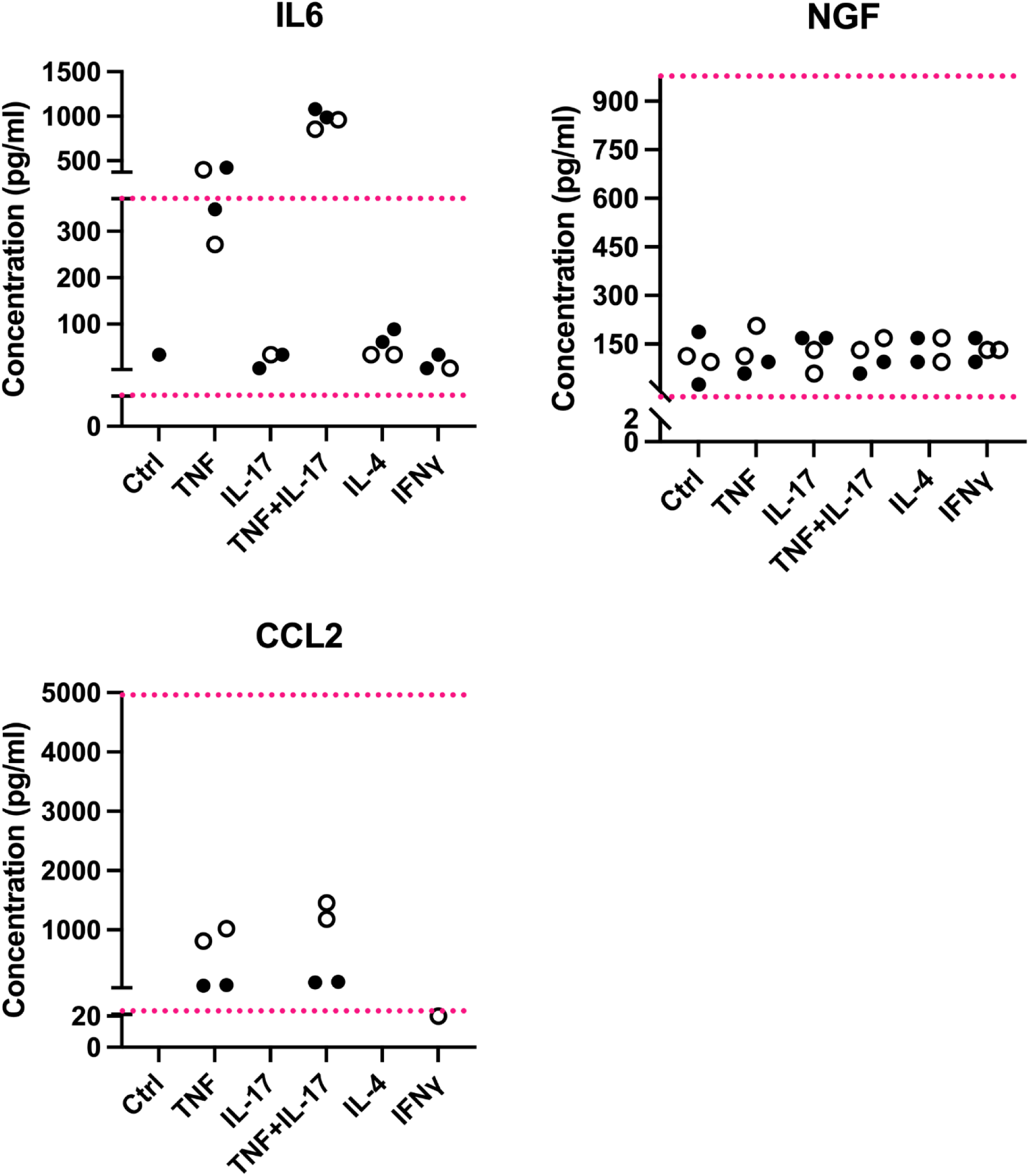
Known pro-algesic mediators IL-6, NGF and CCL2 are detectable in human pericyte lines. Each dot is a well of cells. Filled circles shows the expression levels obtained from media taken from a placental pericyte line, open circles those obtained from a nerve pericyte line. Ctrl = regular pericyte media; other columns: pericytes treated for four hours with TNF (10 ng/ml), IL-17 (10 ng/ml), TNF+IL-17 (both at 10 ng/ml), IL-4 (10 ng/ml) or IFNγ (100 ng/ml). The red dotted lines represent the upper and lower limits of the standard curves for each protein, i.e. the upper and lower detection limits. Shown here are 1:100 dilutions for IL-6 and NGF and 1:2 dilutions for CCL2.

### Conditioned medium from human sciatic nerve pericytes induces intracellular activation of human IPSC-cell derived sensory neurons

Finally, we tested whether human sciatic nerve pericytes are able to impact human sensory neurons. We once again compared naïve pericytes with those stimulated with cytokines for four hours (TNF or TNF and IL-17). After treatment, media were removed and used on human stem-cell derived sensory neurons for one hour. Western blotting revealed that conditioned medium from stimulated pericytes was able to induce phosphorylation of the transcription factor STAT3 in sensory neurons (**Figure 8**). Phosphorylation of STAT3 is a known consequence of peripheral nerve injury (Qiu et al., 2005; Schwaiger et al., 2000), and particularly noteworthy in the context of our study: several of the mediators we identified as up-regulated with nerve injury in mouse mural cells are cytokines which activate the JAK/STAT pathway (Il6, Lif and Clcf1). Our ligand-receptor analyses suggest these should be able to bind mouse and human sensory neurons directly, signalling via the g130 receptor (*Il6st*) and its various co-receptors (e.g. soluble Il-6, Lif and Cntf receptors). Our conditioned medium experiments now experimentally support the notion that substances released from stimulated pericytes, such as IL6 shown in **Figure 7**, can activate sensory neurons directly.

**Figure 8:**
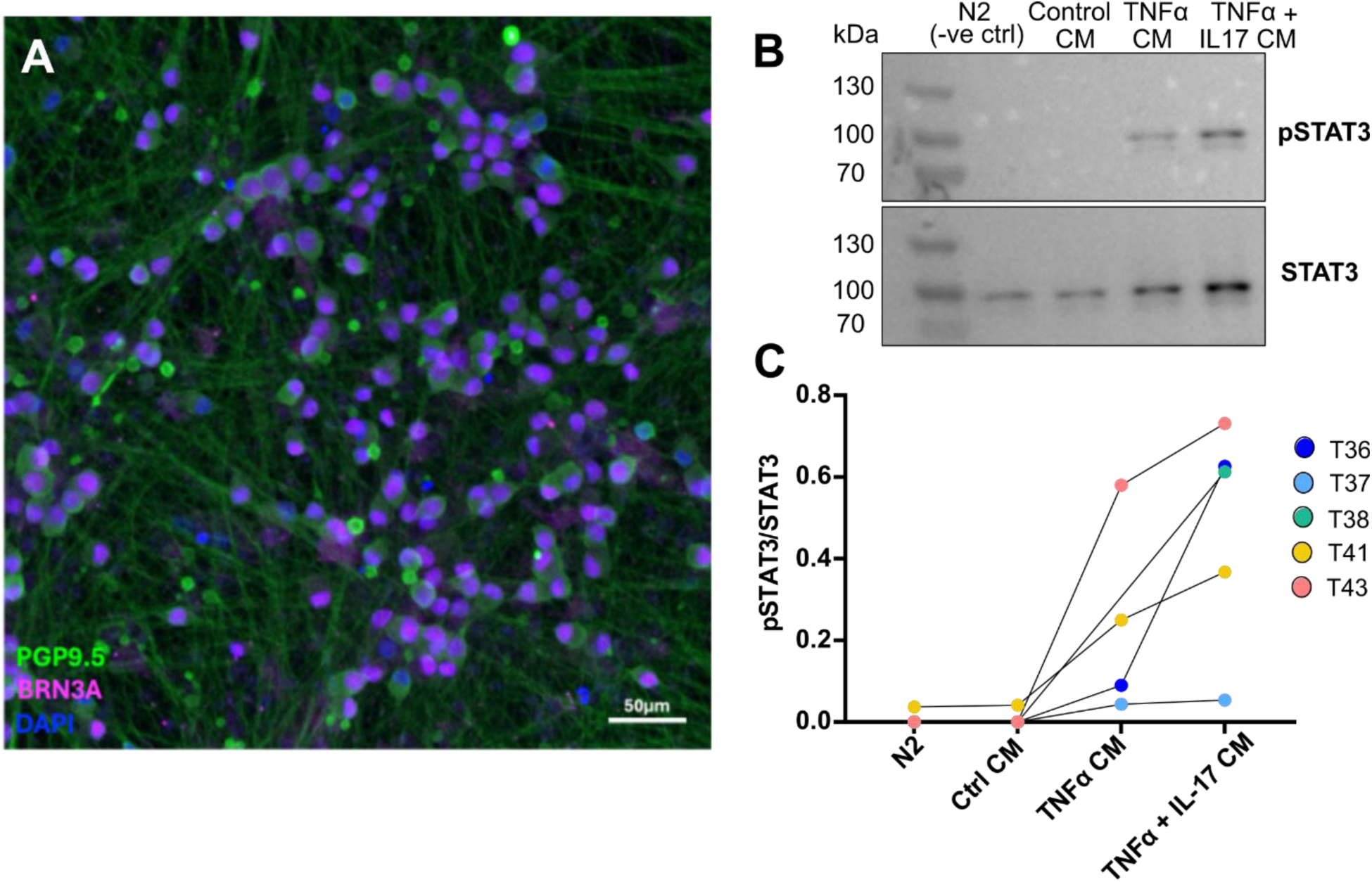
Conditioned media derived from human nerve-derived pericytes induce phospho-STAT3 in human IPSC-derived sensory neurons. (A) Representative immunocytochemistry image of stem-cell derived sensory neurons stained with pan-neuronal marker PGP9.5, sensory neuron transcription factor BRN3A and nuclear marker DAPI. (B) Western blot of neuronal pSTAT3 and STAT3 following conditioned media incubation. See Supplementary Materials for all full-length Western Blots. (C) Quantification of neuronal pSTAT3/STAT3 following incubation with cytokine-activated pericyte conditioned media (CM). Each dot represents an independent neuron differentiation (T36-43, n = 5). N2 = neuronal control medium.

## Discussion

In summary, we present a series of experiments that indicate that Pdgfrb+ fibroblasts and mural cells in sciatic nerve express prominent known and putative pro-algesic mediators that are further upregulated upon painful nerve injury. We also obtained both flow cytometry and immunostaining data to suggest that the number of Pdgfrb+ mural cells is persistently increased in damaged sciatic nerve, detectable up to 2 months after an initial insult. Finally, we provide evidence that the pro-algesic potential of mural cells translates to human, with a nerve pericyte line expressing some of the same pro-algesic mediators and inducing STAT3 phosphorylation in human stem-cell derived sensory neurons upon activation. It remains to be determined whether this translates into regulation of pain-relevant genes as well as neuronal hyperactivity, as we have previously observed following stimulation with JAK-STAT cytokines (Li et al., 2024a).

Some of our day 5 flow cytometry results at first glance appear to be contradicted by later tissue clearing data. Specifically, with flow cytometry we observed an increase in mural cells five days after nerve injury **(Figure 2A-C)**, while quantifying these cells in situ in cleared nerves only showed a significant increase in Pdgfrb+ cells in general, rather than Pdgfrb+ mural cells more specifically. We believe that this is a reflection of the challenges associated with attempting to capture changes in a small population of cells while being faced with not insignificant within- and across-batch variation; for example, the number of mural cells we determined in our day 5 post-PSNL imaging experiments ranged from 50-350, with batch 1 displaying higher counts and injury-induced changes more in line with our flow cytometry results (**Supplementary Figure 4**). At the two-month time-point, the increase in Pdgfrb+ cells replicated across both flow cytometry and tissue clearing, suggesting that, if anything, the dysregulation in Pdgfrb+ fibroblasts and mural cells worsens over time.

Fibroblasts have been implicated previously in nerve-injury induced pain (Singhmar et al., 2020), with a particular focus on a Pi16+ population, which is contained amongst our Pdgfrb+ cells (**Figure 4**). In general, it is important to note that the Pdgfrb+ fibroblasts we observed in mouse nerves are not homogeneous, but consist of several different sub-populations, including known universal sub-types positive for marker genes *Col15a1*, *Pi16*, *Ccl19* (Buechler et al., 2021; Korsunsky et al., 2022). In our single-cell sequencing data, it appeared that Col15a1+ fibroblasts were reduced in number upon nerve injury (making up 2-11% of all cells in PSNL vs. 18-33% in sham samples). However, given the small sample size (n = 3), it remains to be determined whether this is a replicable effect.

Mural cells have not been explicitly studied to date with respect to traumatic neuropathic pain states. In contrast, in the central nervous system (CNS), their altered functions in ischemic and neurodegenerative conditions are well-described (Dalkara et al., 2024). Thus, pericytes contract when blood flow is interrupted in a mouse model of ischemic stroke and remain contracted even when middle cerebral artery occlusion is lifted (Yemisci et al., 2009). There is evidence to suggest that this failure to allow re-perfusion is due to pericytes dying in a state of rigour, with their ‘corpses’ constricting the capillaries they envelop, presumably until they are removed by phagocytic cells (Dalkara et al., 2024; Hall et al., 2014).

Whether similar mechanisms will be at play in our specific nerve injury model is less clear. While peripheral nerves can develop the same re-perfusion injuries that are detectable in CNS, it is generally very difficult to render them ischemic. In fact, most rodent studies report that crush or focal nerve injuries result in hyperemia, i.e. increased rather than decreased blood flow (Zochodne, 2018). This also appears to be the case for human entrapment neuropathies, specifically carpal tunnel syndrome, where hypervascularisation appears to correlate with symptom severity (Kutlar et al., 2017). There are exceptions to this, with ischemia reported in specific models, like chronic constriction injury (Levy and Zochodne, 1998), or chronic neuromas (Xu and Zochodne, 2002) in which disruption of blood flow appeared to vary across zones, with some showing abnormal levels of angiogenesis, while others appeared to be ischemic. It is tempting to speculate that our result, of a long-lasting increase in pericyte numbers, is more in keeping with angiogenesis. In the clinic, the nature of blood flow alterations (hyperemia vs. ischemia) and the degree of inflammation will likely vary across individuals and nerve injury types. As a result, there have been calls to revise the current grading system for peripheral nerve injuries (Yeoh et al., 2022), which was first developed in the 1950s (Sunderland, 1951).

Whatever the consequence of pericyte dysfunction on blood flow, our data indicate traumatic nerve injury alters mural cell and Pdfgrb+ fibroblast transcription in a way that is likely to sensitize nociceptive neurons. Ironically, this may be an unfortunate side-effect of the system trying to remodel and repair its blood supply, with many of the pro-algesic mediators we observe upregulated after injury (*Ngf*, *Il6*, *Lif*, *Ccl2*) also having known roles in angiogenesis, vasodilation and vascular remodelling (Funamoto et al., 2000; Hilgendorf et al., 2024; Nian et al., 2004; Peplow, 2014). As expected, given the discovery of universal, cross-tissue fibroblast sub-types, our results in nerve parallel what is being reported in joint, where fibroblast dysfunction is increasingly being linked to sensory neuron hyperactivity and pain (Shinotsuka and Denk, 2022; Wijesinghe et al., 2024), most recently also in patient cohorts (Bai et al., 2024).

In conclusion, we propose that mesenchymal cell abnormalities should be considered when developing novel strategies to tackle pain as a result of physical nerve entrapment. They are also likely to be relevant to other conditions which can cause chronic pain, including arthritis and diabetic neuropathy, where alterations in pericyte and blood-nerve-barrier function have consistently been identified (Richner et al., 2018). Indeed, a recent single cell atlas of human sural nerves taken from individuals living with polyneuropathy identified vascular smooth muscle cells and perineural fibroblasts as two of the cell types that were more abundant in disease and that correlated with the extent of disability experienced by the donors (Heming et al., 2024). More work is now needed to turn these novel mechanistic insights into future drug development opportunities to finally provide relief for the millions of people suffering from neuropathic pain every day.

## Supporting information

Supplementary R Notebooks

## Acknowledgements

SV and LF were funded by an Ono Pharmaceuticals Rising Stars Award held by FD. YL by the Wellcome Trust as part of the Neuro-Immune Interactions in Health & Disease Wellcome Trust PhD Programme (218452/Z/19/Z). IZ, JVW, LT and FD were funded by a Wellcome Trust Collaborative Award 224257/Z/21/Z. HD was supported by a Sir Henry Wellcome Fellowship (224049/Z/21/Z).

We acknowledge King’s College London as the source of HPSI0714i-kute_4 human iPSC line which was generated under the Human Induced Pluripotent Stem Cell Initiative funded by a grant from the Wellcome Trust and Medical Research Council, supported by the Wellcome Trust (WT098051) and the NIHR/Wellcome Trust Clinical Research Facility, and acknowledges Life Science Technologies Corporation as the provider of Cytotune.

This research was funded in whole or in part by UK Research & Innovation and the Wellcome Trust. For the purpose of Open Access, the author has applied a CC BY public copyright licence to any Author Accepted Manuscript (AAM) version arising from this submission.

None of the authors have any conflicts of interest to declare in relation to this work.

