## Supplementary Tables & Figures for "A role for fibroblast and mural cell subsets in a nerve ligation model of neuropathic pain"

### Supplementary Figures

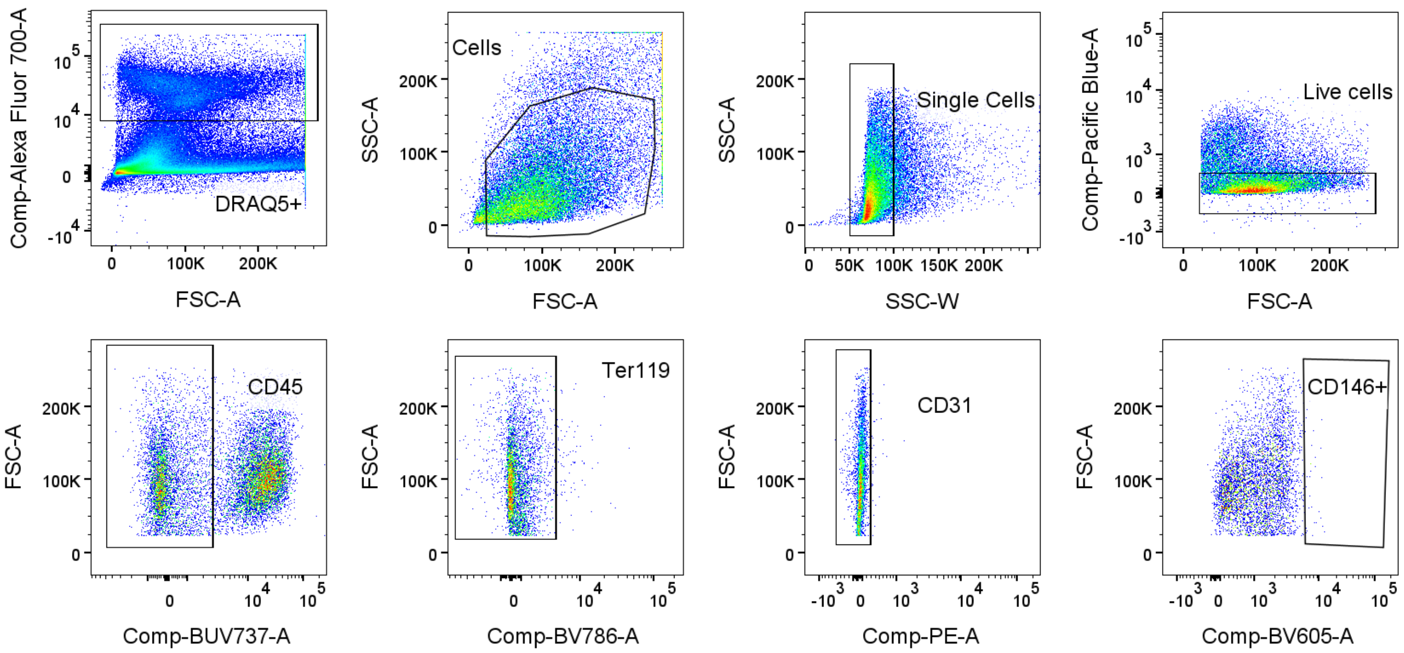


**Supplementary Figure 1: Gating strategy for flow cytometry experiment shown in Figure 2A.** Nucleated events were selected with DRAQ5; followed by selection of single, live cells and exclusion of immune cells (CD45 neg), erythrocyte precursors (Ter119 neg) and endothelial cells (CD31 neg). CD146 positive mural cells were then quantified. The sample shown here is from an ipsilateral nerve. All gates were placed using fluorescence minus one controls, with the CD146 FMO shown here.

**FMO**

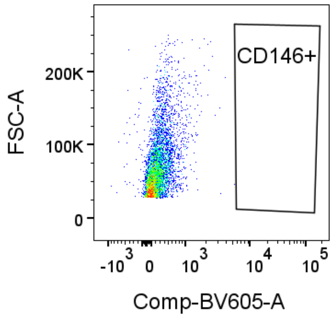

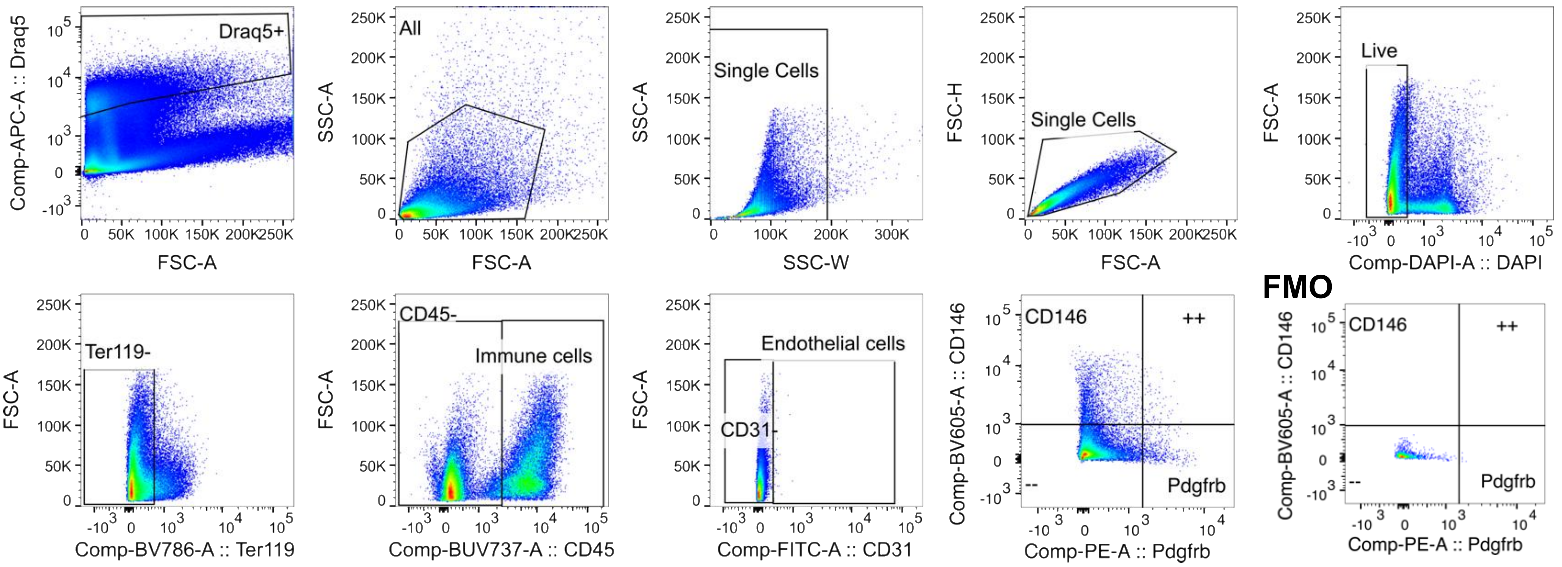


**Supplementary Figure 2: Gating strategy for flow cytometry experiment shown in Figure 2B.** Nucleated events were selected with DRAQ5; followed by selection of single, live cells and exclusion of erythrocyte precursors (Ter119 neg), immune cells (CD45 neg) and endothelial cells (CD31 neg). Pdgfrb/CD146 double positive mural cells were then quantified. The sample shown here is from an ipsilateral nerve. All gates were placed using fluorescence minus one controls, with the CD146 FMO shown here.

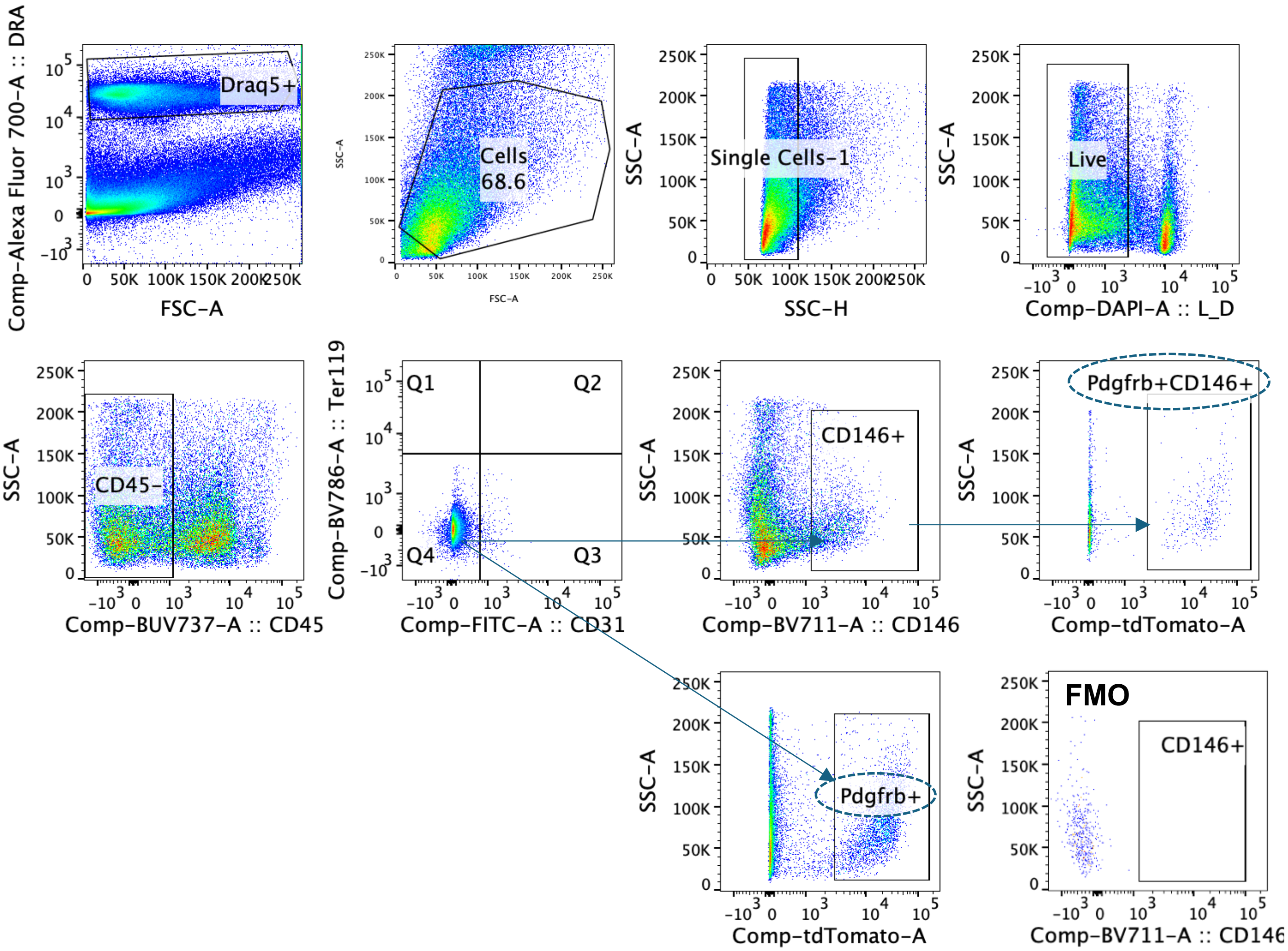


**Supplementary Figure 3: Gating strategy for FACS experiments shown in Figures 2C&D.** Nucleated events were selected with DRAQ5; followed by selection of single, live cells and exclusion of immune cells (CD45 neg), erythrocyte precursors (Ter119 neg) and endothelial cells (CD31 neg). Two populations, here indicated by dotted circles, were then quantified: CD146+ events further selected to also be Pdgfrb-tdtomato positive versus all Pdgfrb-tdtomato positive cells. The sample shown here is from an ipsilateral nerve, two months after partial sciatic nerve ligation. All gates were placed using fluorescence minus one controls, with the CD146 FMO shown here.

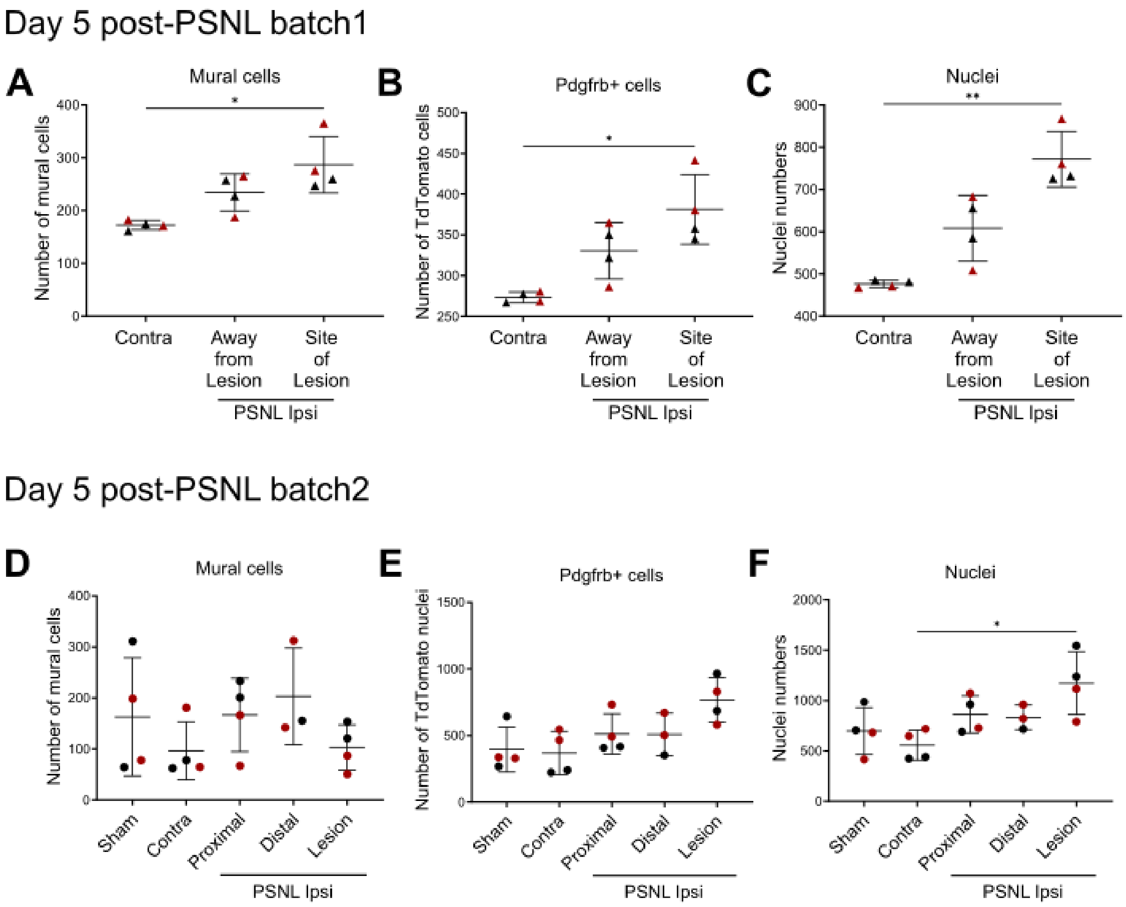


**Supplementary Figure 4. Similar results were observed across two batches of day 5 post-PSNL traumatic nerve injury, as measured by confocal imaging of cleared sciatic nerve.**

A-C) 5 days post PSNL cell numbers were significantly increased at the site of lesion in ipsilateral, injured compared to contralateral nerves: A) mural cells ($p=0.0134$); B) Pdgfrb+ cells ($p=0.0134$); C) total number of nuclei ($p=0.0051$). n = 4 nerves per group. In a second batch of samples, we observed similar trends, but these were only significant for total nuclei ($p=0.0315$), n = 4 nerves per group. Datasets were analysed with a non-parametric Kruskal-Wallis one-way ANOVA with Dunn's multiple comparisons (black symbols = males, red symbols = females).

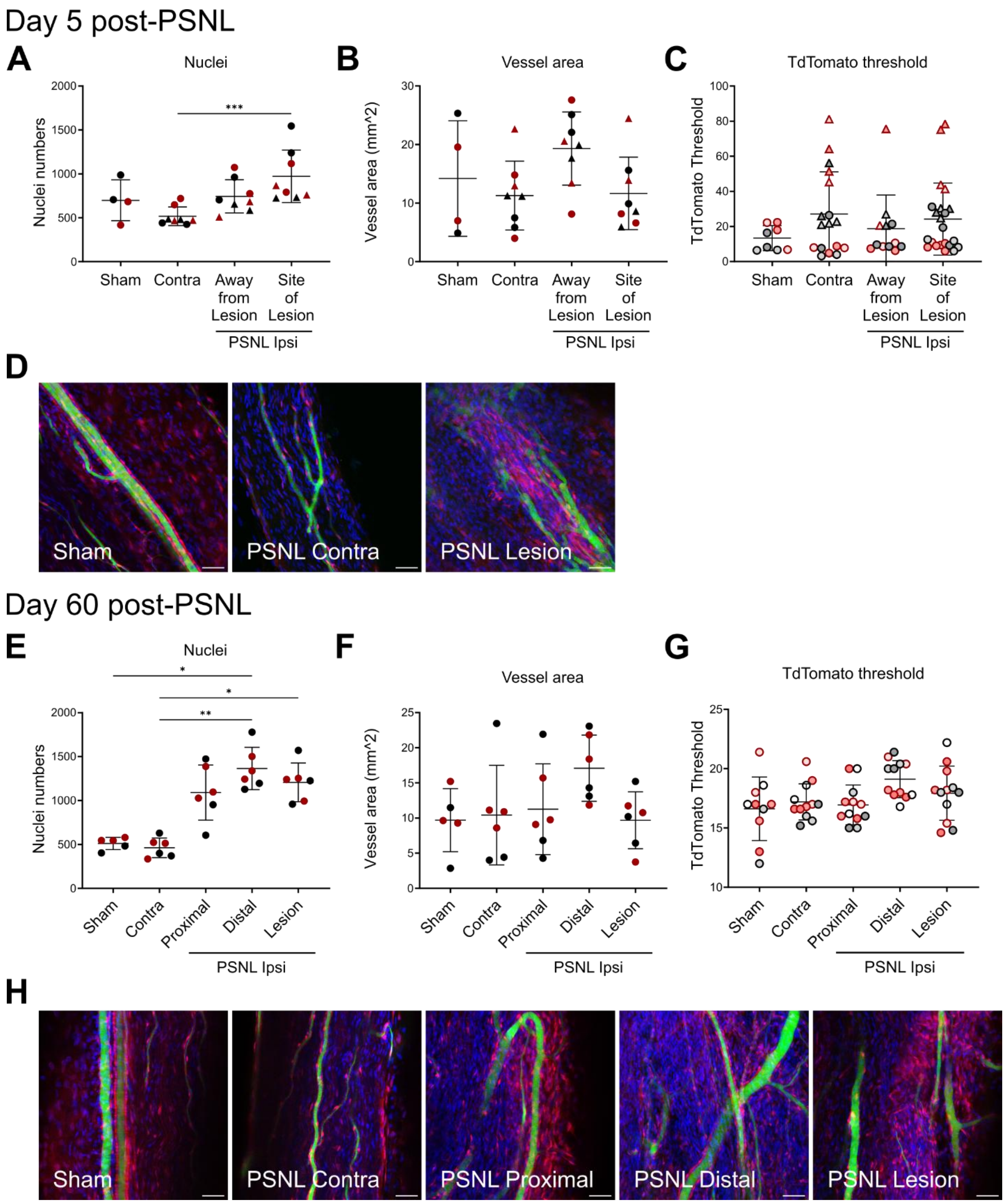


**Supplementary Figure 5. Nerve immune cells are increased upon traumatic nerve injury, as measured by confocal imaging of cleared sciatic nerve.** A) Number of nuclei and B) vessel area 5 days post PSNL or sham surgery. Experiment carried out in two batches (triangles = batch1, n=4; circles = batch 2, n=4). C) Threshold used in the TdTomato channel for the 5 day analysis. D) Representative maximum intensity projection images of the sciatic nerve of PdgfrbCre x TdTomato mice perfused with FITC-albumin-gelatin 5 days after sham or PSNL surgery. E) Number of nuclei and F) vessel area per nerve region 60 days post PSNL or sham surgery (n=5-6). G) Threshold used in the TdTomato channel for the day 60 analysis. H) Representative maximum intensity projection images of the sciatic nerve of PdgfrbCre x TdTomato mice perfused with FITC-albumin-gelatine 60 days after sham or PSNL surgery. Datasets were analysed with a non-parametric Kruskal-Wallis one-way ANOVA with Dunn's multiple comparisons (black symbols = males, red symbols = females). For A, B, E & F each symbol represents a mouse. For C & G each symbol represents an analysed z stack. Scale bars = 50μm.

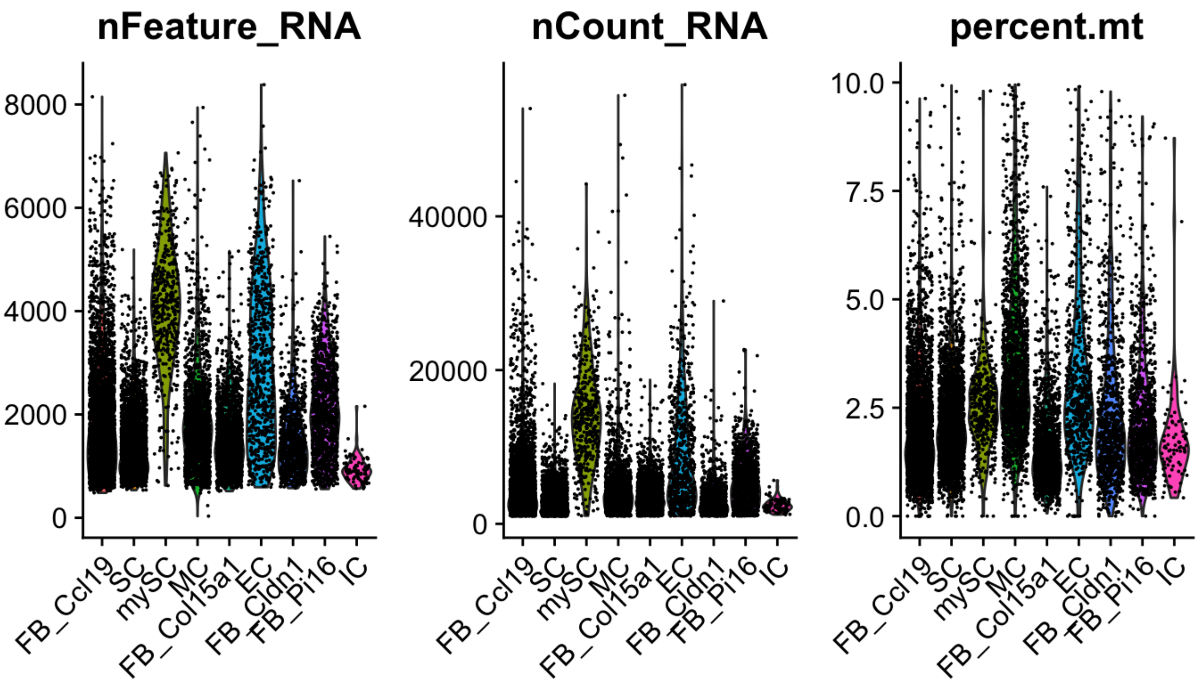


**Supplementary Figure 6: Quality control measures for the different clusters show no major differences between them** with the exception of myelinating Schwann and endothelial cells, possibly due to proliferation and larger size, respectively. Each dot is a cell; nFeature = number of genes detected in each cell; nCount = number of molecules detected in each cell; percent.mt = percentage of mitochondrial genes in each cell starting with 'mt-' in the gene name.

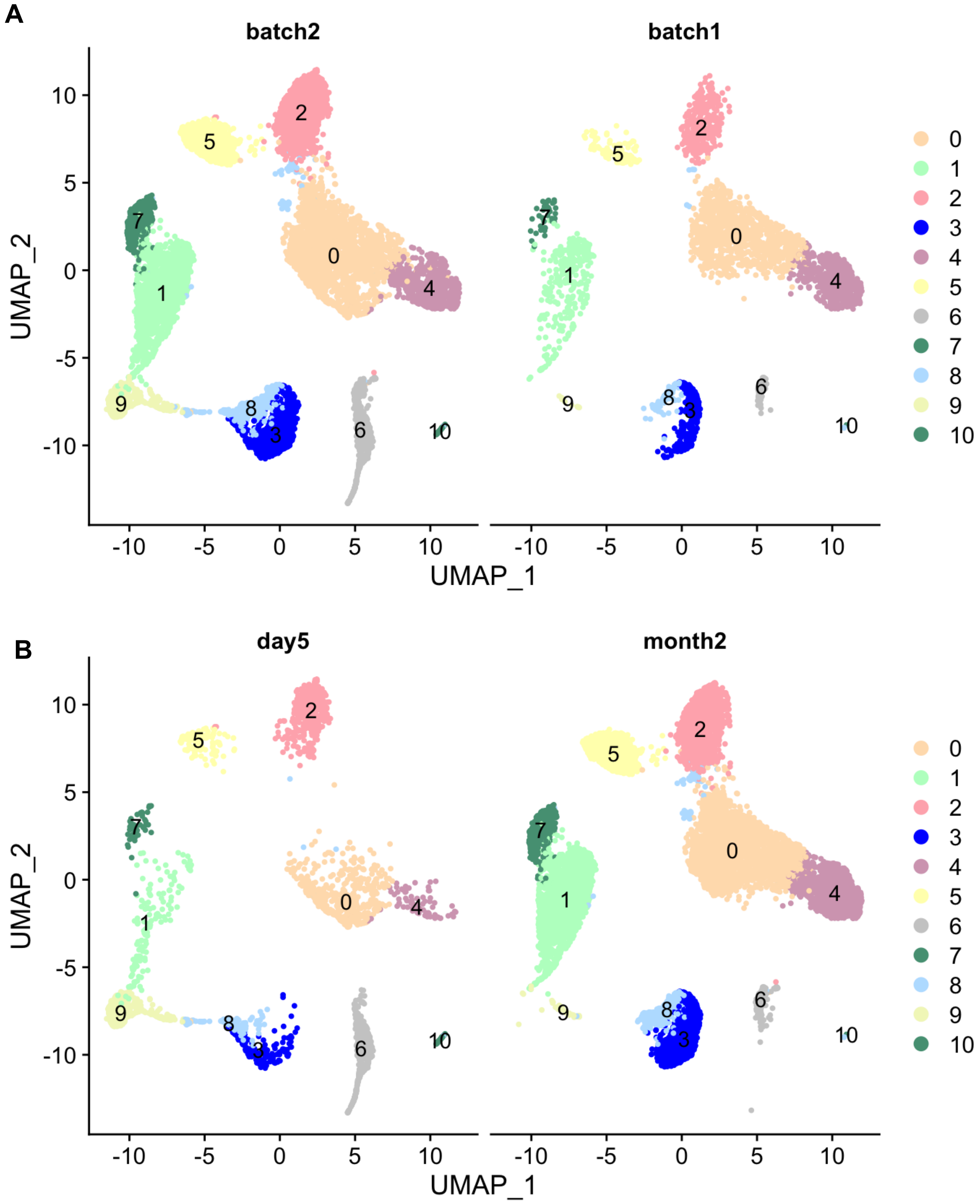


**Supplementary Figure 7: Cell-type differences appear to supersede technical batch effects, at least at a gross, reductionist level of cluster allocation in 2D.** (A) Unsupervised clustering reveals 11 different cell clusters across the two experimental batches, with batch 1 including just the chronic time point, while batch 2 includes data on chronic and acute timepoints. (B) The only striking differences are due to differences in biology, with three clusters only prominent 5 days vs 2 months after injury: cluster 9, likely myelinating Schwann cells; cluster 10: contaminating immune cells; and a preponderance of an endothelial cell cluster 6.

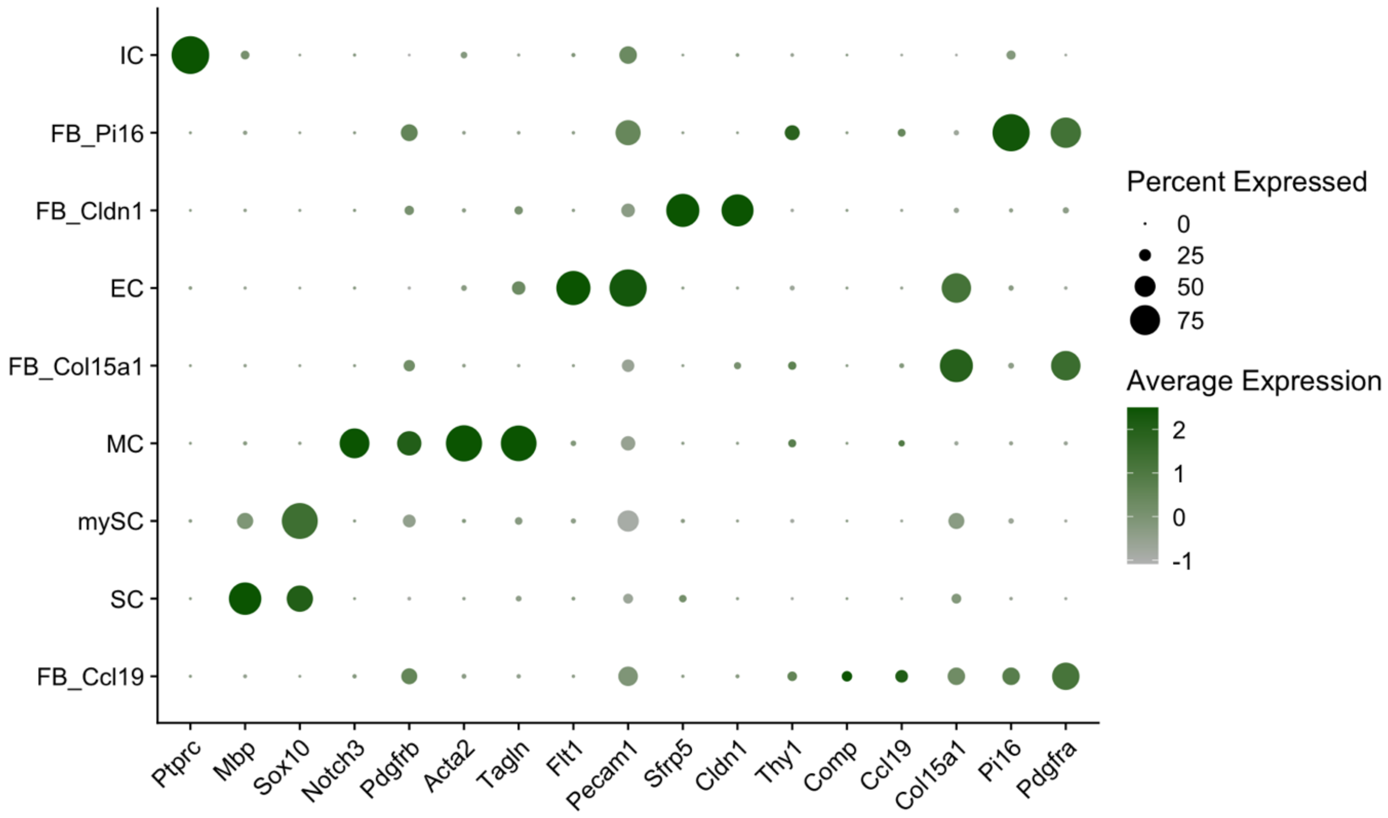


**Supplementary Figure 8: Marker gene expression by cluster.** Genes (on the x-axis) used to determine the likely cell identity of the various clusters (on the y-axis). The bigger the dot, the higher the percentage of cells expressing that particular gene in a given cluster. The 'greener' the dot, the higher the expression of the gene in the cluster.

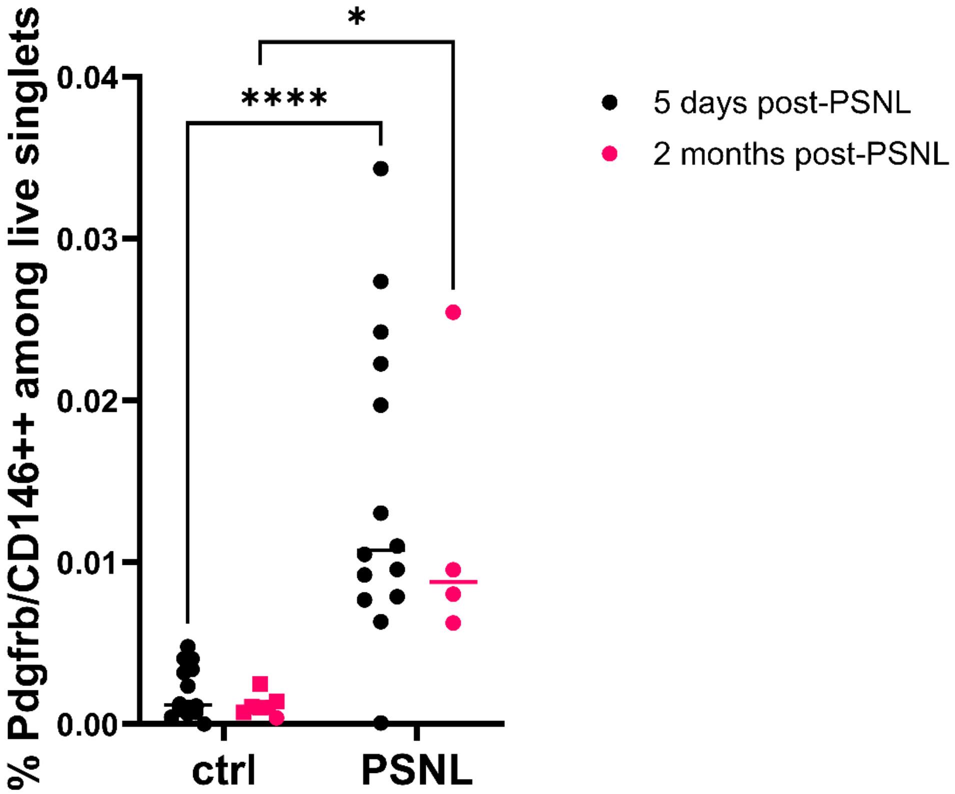


**Supplementary Figure 9. The percentage of Pdgfrb/CD146+ cells compared to the total number of live nucleated events appeared increased after PSNL at both 5 days and 2 months post-nerve injury.** Each dot is a mouse nerve; in black: n = 14, day 5 post-surgery, ctrl = contralateral, PSNL = ipsilateral, injured nerves; in pink: month 2 post-surgery, ctrl = contralateral uninjured nerves (n = 2, circles) and ipsilateral sham nerves (n = 5, squares), PSNL = ipsilateral, injured nerves (n = 4). A two-way ANOVA revealed a main effect of Injury, regardless of timepoint: $F(1, 35) = 25.11$, $p < 0.0001$, with Sidak's post-hoc tests: **** $p < 0.0001$, * $p = 0.0183$.

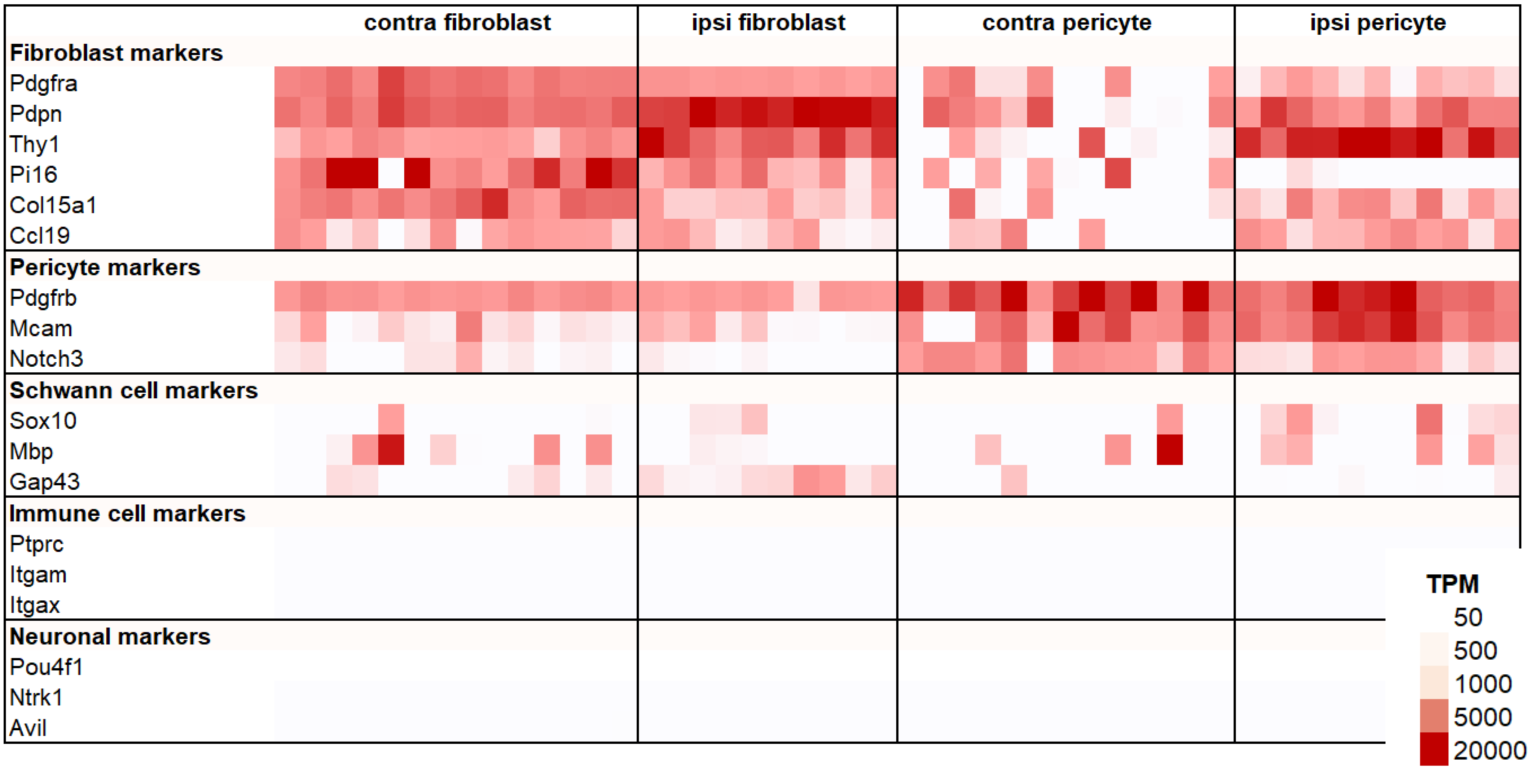


**Supplementary Figure 10. Expression level of key marker genes indicates occasional contamination with Schwann-cell related transcripts.** Gene expression levels (transcript per million, TPM) for a select range of markers indicates no immune and neuronal transcript contamination and only occasional contamination with Schwann-cell related transcripts.

### Supplementary Tables

**a) Panel for Figure 2A**

| Excitation Laser | Emission | Colour | Epitope | Catalogue # | Dilution |
|---|---|---|---|---|---|
| **UV** | Near-IR | BUV737 | CD45 | BD Bioscience 748371 | 1:300 |
| **Violet** | Far red | BV785 | Ter119 | Biolegend 116245 | 1:300 |
| | Orange/Red | BV605 | CD146 | BD Bioscience 740434 | 1:300 |
| | Violet | ~451 nm | Live/Dead | Invitrogen L34955 | 1:300 |
| **Yellow/Green** | Yellow/Orange | PE | CD31 | Biolegend 102507 | 1:300 |
| | | PE-Cy7 | CD90 | Biolegend 140309 | 1:300 |
| **Red** | Far red | DRAQ5 | Nucleated events | Biolegend 424101 | 1:5000 |
| **N/A** | N/A | N/A | FC block | Biolegend 101301 | 1:20 |

**b) Panel for Figure 2B**

| Excitation Laser | Emission | Colour | Epitope | Catalogue # | Dilution |
|---|---|---|---|---|---|
| **UV** | 450 | DAPI | Dead | SLS D9542-1M | 10ng/ml |
| | 737 | BUV737 | CD45 | BD 748371 | 1:300 |
| **Violet** | 610 | BV605 | CD146 | BD 740434 | 1:300 |
| | 780 | BV785 | Ter119 | Biolegend 116245 | 1:300 |
| **Blue** | 525 | FITC | CD31 | Biolegend 102405 | 1:300 |
| **Yellow/Green** | 575 | PE | PDGFRB | Biolegend 136005 | 1:300 |
| | 780 | PE-Cy7 | Thy1.2 | Biolegend 140309 | 1:300 |
| **Red** | 660+780 | APC | DRAQ5 | Biolegend 424101 | 1:10000 |
| **N/A** | N/A | | FC block | Biolegend 101320 | 1:20 |

**c) Panel for Figures 2C&D**

| Excitation Laser | Emission | Colour | Epitope | Catalogue # | Dilution |
|---|---|---|---|---|---|
| **UV** | 450 | DAPI | Dead | SLS D9542-1M | 10ng/ml |
| | 737 | BUV737 | CD45 | BD 748371 | 1:300 |
| **Violet** | 780 | BV785 | Ter119 | Biolegend 116245 | 1:300 |
| | 711 | BV711 | CD146 | BD 740827 | 1:300 |
| **Blue** | 525 | FITC | CD31 | Biolegend 102405 | 1:300 |
| **Yellow/Green** | 581 | tdtomato | PDGFRB | transgene | |
| | 780 | PE-Cy7 | Myelin | | |
| **Red** | 660+780 | Far-Red | DRAQ5 | Biolegend 424101 | 1:10000 |
| **N/A** | N/A | | FC block | Biolegend 101320 | 1:20 |

**Supplementary Table 1: Flow and FACS panels used to label mesenchymal lineage cells.** See **Supplementary Figures 1-3** for corresponding gating strategies.

| Day 5 post-PSNL batch 1 | | % images containing: | | | | Number of images analysed |
|---|---|---|---|---|---|---|
| | | Venules | Arterioles | Epineurial capillaries | Endoneurial vessels | |
| PSNL | Contra | 0.0% | 77.8% | 88.9% | 55.6% | 9 |
| PSNL Ipsi | Away from Lesion | 50.0% | 37.5% | 87.5% | 50.0% | 8 |
| | Site of Lesion | 50.0% | 25.0% | 100.0% | 25.0% | 4 |

| Day 5 post-PSNL batch 2 | | % images containing: | | | | Number of images analysed |
|---|---|---|---|---|---|---|
| | | Venules | Arterioles | Epineurial capillaries | Endoneurial vessels | |
| Sham | | 0.0% | 50.0% | 75.0% | 75.0% | 8 |
| PSNL | Contra | 0.0% | 12.5% | 87.5% | 100.0% | 8 |
| PSNL Ipsi | Proximal | 0.0% | 100.0% | 100.0% | 25.0% | 8 |
| | Distal | 50.0% | 50.0% | 100.0% | 16.7% | 6 |
| | Lesion | 37.5% | 37.5% | 75.0% | 25.0% | 8 |

| Day 60 post-PSNL | | % images containing: | | | | Number of images analysed |
|---|---|---|---|---|---|---|
| | | Venules | Arterioles | Epineurial capillaries | Endoneurial vessels | |
| Sham | | 20.0% | 50.0% | 80.0% | 90.0% | 10 |
| PSNL | Contra | 0.0% | 58.3% | 83.3% | 100.0% | 12 |
| PSNL Ipsi | Proximal | 16.7% | 66.7% | 83.3% | 91.7% | 12 |
| | Distal | 25.0% | 50.0% | 83.3% | 91.7% | 12 |
| | Lesion | 0.0% | 91.7% | 100.0% | 58.3% | 12 |

**Supplementary Table 3: Percentage of images containing each type of blood vessel.**

Table showing the percentage of images containing each type of blood vessel characterised in each experimental batch. The last column shows the number of z-stack images obtained per condition and nerve section.

| **Kalinski et al.** | Immune | **MC** | EC | **FB** | ? | SC |
|---|---|---|---|---|---|---|
| Il6 | 0.5 | **10.7** | 6.3 | **3.4** | 0.2 | 0.1 |
| Ngf | 0.2 | **28.4** | 0.3 | **14.7** | 0.5 | 1 |
| Ccl2 | 36.9 | **19.5** | 8.6 | **55.1** | 17.5 | 21 |
| Lif | 0.4 | **0.6** | 0.1 | **11.5** | 0.8 | 8.1 |
| Notch3 | 1 | **90.1** | 3.2 | **7.1** | 1.1 | 1.1 |
| Pdgfrb | 1.4 | **85.8** | 2.6 | **55.5** | 9.3 | 10.8 |

| **Carr et al.** | **FB2** | **FB** | **?** | CD45 | EC | SC | **PC** |
|---|---|---|---|---|---|---|---|
| Il6 | **5** | **5.7** | **5.6** | 1.2 | 11 | 0.6 | **8.4** |
| Ngf | **4.9** | **1.1** | **3.5** | 0.2 | 0.6 | 0.4 | **9.8** |
| Ccl2 | **42.7** | **15** | **29.5** | 17.6 | 9.3 | 8.6 | **13** |
| Lif | **6** | **1.9** | **3.3** | 0.6 | 0.4 | 3.8 | **0.2** |
| Notch3 | **2.6** | **3.9** | **5.1** | 0.8 | 5.5 | 0.5 | **54** |
| Pdgfrb | **17.2** | **14.3** | **12.4** | 1 | 4.1 | 5.2 | **45.2** |

| **Wolbert et al.** | **nmSC** | CD4 | MC | **mySC** | **fibro** | **vSMC/PC** | EC | CD8 | pDC | BC | EP |
|---|---|---|---|---|---|---|---|---|---|---|---|
| Il6 | **1.3** | 0.2 | 1.6 | **0.1** | **1.3** | **1.1** | 0 | 0 | 0 | 0.1 | 0 |
| Ngf | **2.5** | 0 | 0.2 | **0.2** | **0.2** | **3.3** | 0 | 0.1 | 0 | 0.1 | 0 |
| Ccl2 | **11.4** | 1.2 | 25.8 | **2.7** | **6.1** | **2.2** | 2.3 | 0.8 | 0 | 1.2 | 0 |
| Lif | **0.8** | 0.5 | 0 | **0** | **0** | **0** | 0 | 0 | 0 | 0 | 0 |
| Notch3 | **0** | 0.1 | 0 | **0.2** | **0.6** | **26.7** | 0 | 0.1 | 1.8 | 0.1 | 0 |
| Pdgfrb | **5.6** | 0 | 0 | **4.2** | **5.8** | **30** | 0.6 | 0.2 | 0 | 0.1 | **5.5** |

| **in-house data** | FB_Ccl19 | mySC | **MC** | EC | FB_Pi16 | FB_Col15a1 | FB_Cldn1 | SC | IC |
|---|---|---|---|---|---|---|---|---|---|
| Il6 | 4.3 | 0.5 | **5.7** | 11.1 | 3.1 | 2.3 | 0.2 | 0 | 0 |
| Ngf | 2.2 | 3.2 | **16.2** | 0.4 | 1.6 | 3.2 | 2.6 | 0.3 | 0 |
| Ccl2 | 6.1 | 20 | **4** | 2.2 | 4.5 | 11.6 | 0.7 | 0.8 | 3.5 |
| Lif | 2.5 | 17.4 | **0.4** | 0.3 | 2.7 | 3.8 | 0.4 | 0.5 | 0 |
| Notch3 | 4.1 | 0.9 | **74.6** | 1.2 | 0.9 | 0.7 | 1.3 | 0.2 | 1.2 |
| Pdgfrb | **36.3** | **27.5** | **59.4** | 1.3 | **38.8** | **23.6** | **18.1** | 2.8 | 0 |

**Supplementary Table 4: Percentage of cells that express a particular gene in each scRNA-seq cluster.** Data from four scRNA-seq experiments are shown: re-analysis of GSE153762 (Kalinski et al., ELife, 2020), GSE120678 (Carr et al., Cell Stem Cell, 2019) and GSE142541 (Wolbert et al., PNAS, 2020) and our own in-house scRNA-seq data presented in this paper. In yellow are the clusters which were considered to be Notch3+ mural cells; in green the ones which were considered Pdgfrb+ cells; all other cell clusters were considered Pdgfrb negative. Figure 1D plots the percentages for each gene for mural cells and the percentages for each gene averaged across Pdgfrb+ or Pdgfrb- clusters.

Column titles indicate cluster IDs: FB & fibro = fibroblasts; MC, PC, vVSMC/PC = mural cells; SC & mySC = Schwann cells; EC = endothelial cells; IC = immune cells; CD45 = immune cells; DC = dendritic cells; BC = B cells; CD4 & CD8 = T cells; “?” in Kalinski et al. includes what they annotate to be hybrid immune-vascular cells and chondrocytes; “?” in Carr et al. are cells expressing a mixture of mesenchymal markers, annotated by the authors as dividing mesenchymal cells.

See **Supplementary R-Notebook 2** for corresponding scripts.

| | SHAM_D5 | SHAM_M2.1 | SHAM_M2.2 | SNL_D5 | SNL_M2.1 | SNL_M2.2 |
|---|---|---|---|---|---|---|
| **PC1&2** | 35 (**22%**) | 6 (**18%**) | 79 (**21%**) | 113 (**67%**) | 82 (**22%**) | 94 (**15%**) |
| **PC3** | 3 (**2%**) | 1 (**3%**) | 58 (**16%**) | 0 | 9 (**2%**) | 35 (**5%**) |
| **PC-NC** | 1 (**1%**) | 0 | 0 | 0 | 2 (**1%**) | 20 (**3%**) |
| **vSMC** | 109 (**69%**) | 27 (**79%**) | 231 (**63%**) | 15 (**9%**) | 281 (**75%**) | 495 (**77%**) |
| **aaSMC** | 11 (**7%**) | 0 | 0 | 41 (**24%**) | 0 | 0 |
| **Total #** | **159** | **34** | **368** | **169** | **374** | **644** |

**Supplementary Table 5: Number of cells in each mural cell cluster, split by experimental sample.** Numbers in brackets represent the percentage of cells amongst all mural cells. Mouse mural cells clustered into five groups, resembling cell types previously described in the brain: Vtn, Abcc9, Kcnj8 positive pericytes (PC1&2), Adgrf5 negative (PC3) and Acta2 negative pericytes (PC-NC); vascular smooth muscle cells (vSMC) and arteriolar smooth muscle cells. Experimental samples: sham samples 5 days (SHAM_D5) and two months after surgery (SHAM_M2.1 & SHAM_M2.2); PSNL samples 5 days (SNL_D5) and two months after surgery (SNL_M2.1 & SNL_M2.2).

### Full length Western blots

**Full length Westerns for the blots quantified in the manuscript. Each set of blots was performed on an independent differentiation (T36-43) of Kute4 iPSC into sensory neurons. Conditions were as follows, with conditions 1, 3, 4 and 5 of blot T43 displayed in the manuscript:**

1. **N2 media only (-ve control)**
2. N2 media + IL6 + sIL6R
3. **Control pericyte conditioned media**
4. **TNFα treated pericyte conditioned media**
5. **TNFα + IL-17 treated pericyte conditioned media**
6. TNFα treated pericyte conditioned media + sIL6R
7. TNFα + IL-17 treated pericyte conditioned media + sIL6R
8. Clean pericyte media + TNFα
9. Clean pericyte media + TNFα + IL-17
10. N2 media + IL6 + sIL6R + Tocilizumab
11. TNFα treated pericyte conditioned media + Tocilizumab
12. TNFα + IL-17 treated pericyte conditioned media + Tocilizumab
13. IL-17 treated pericyte conditioned media
14. IL-4 treated pericyte conditioned media
15. IFNγ treated pericyte conditioned media
16. IL-17 treated pericyte conditioned media + sIL6R
17. IL-4 treated pericyte conditioned media + sIL6R
18. IFNγ treated pericyte conditioned media + sIL6R
19. Clean pericyte media + TNFα + IL-17 + IL-4 + IFNγ

**T36**

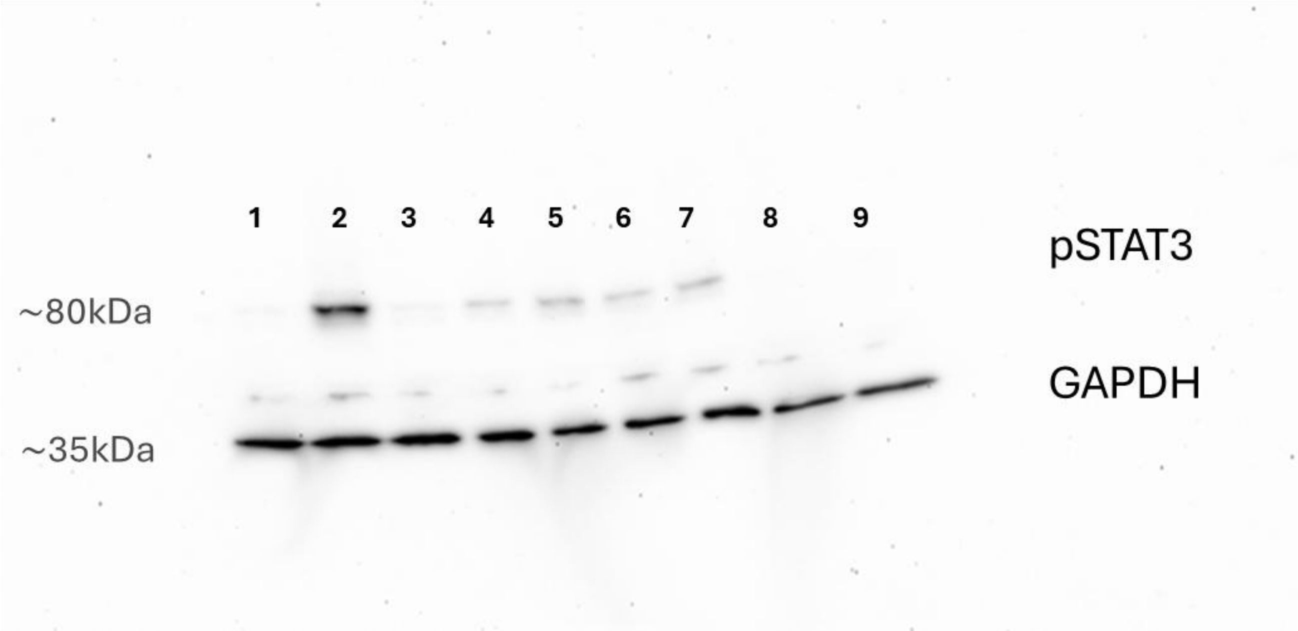


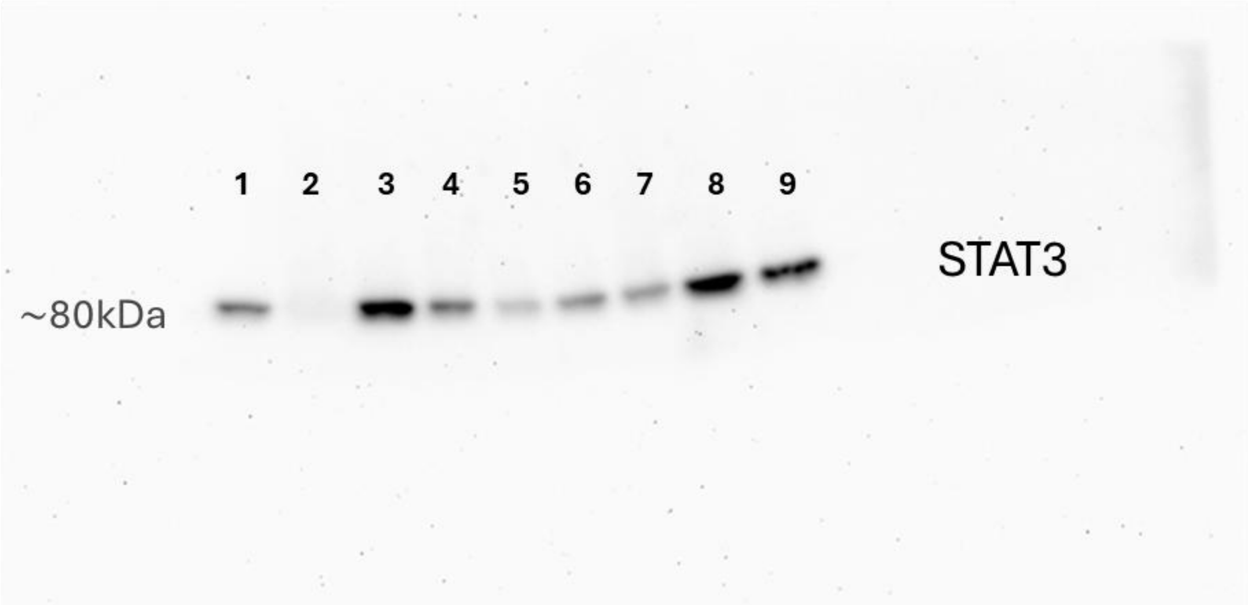

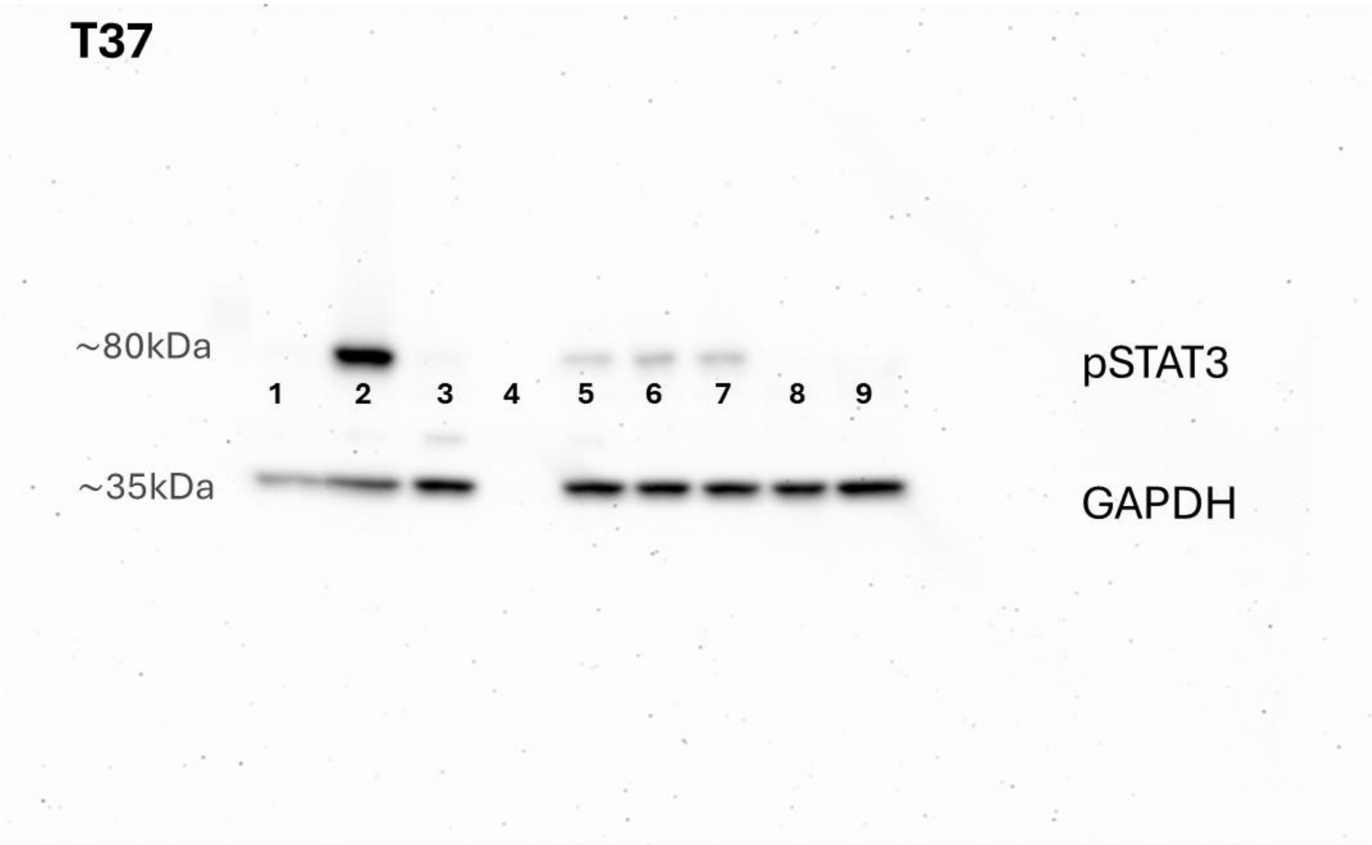
T37
~80kDa
pSTAT3
1 2 3 4 5 6 7 8 9
~35kDa
GAPDH

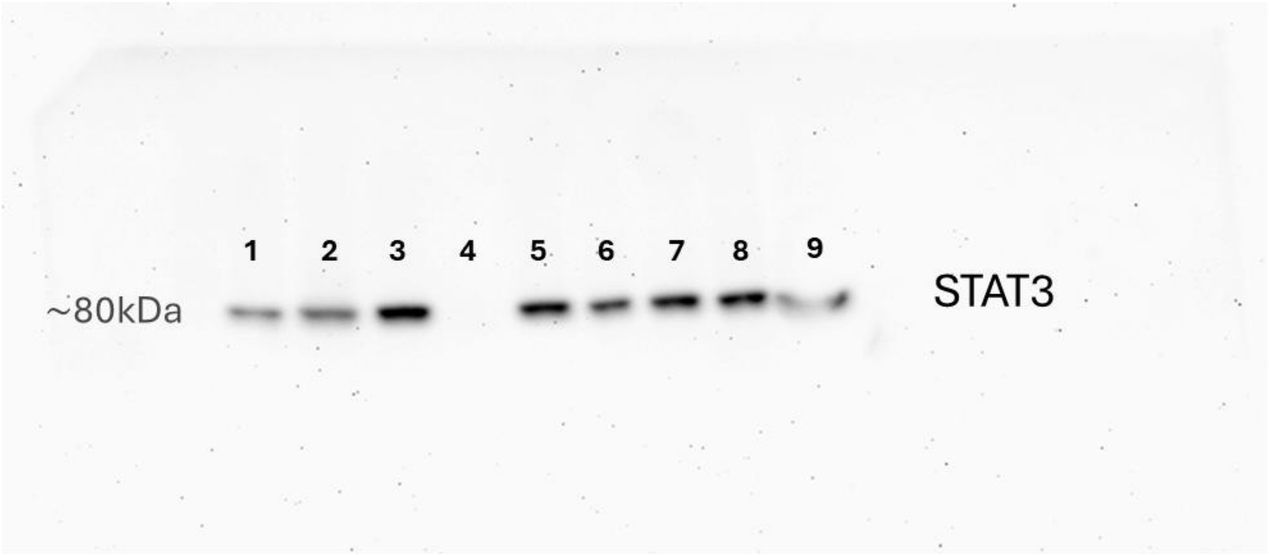
1 2 3 4 5 6 7 8 9
~80kDa
STAT3

# T38

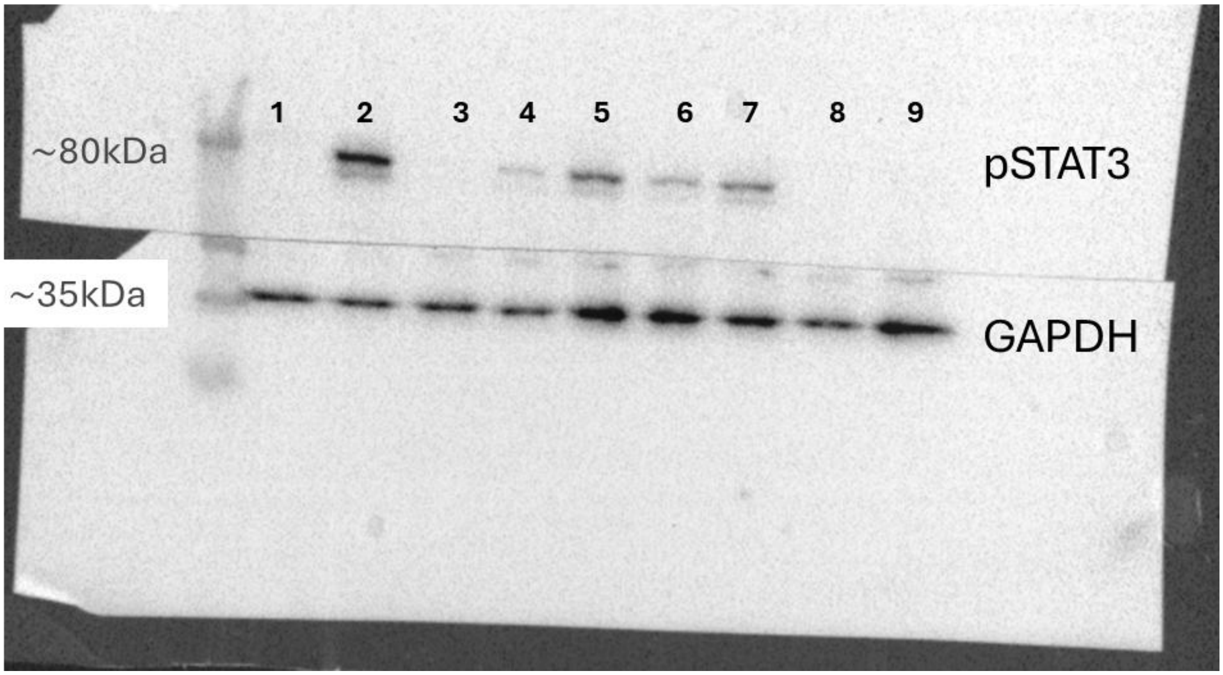


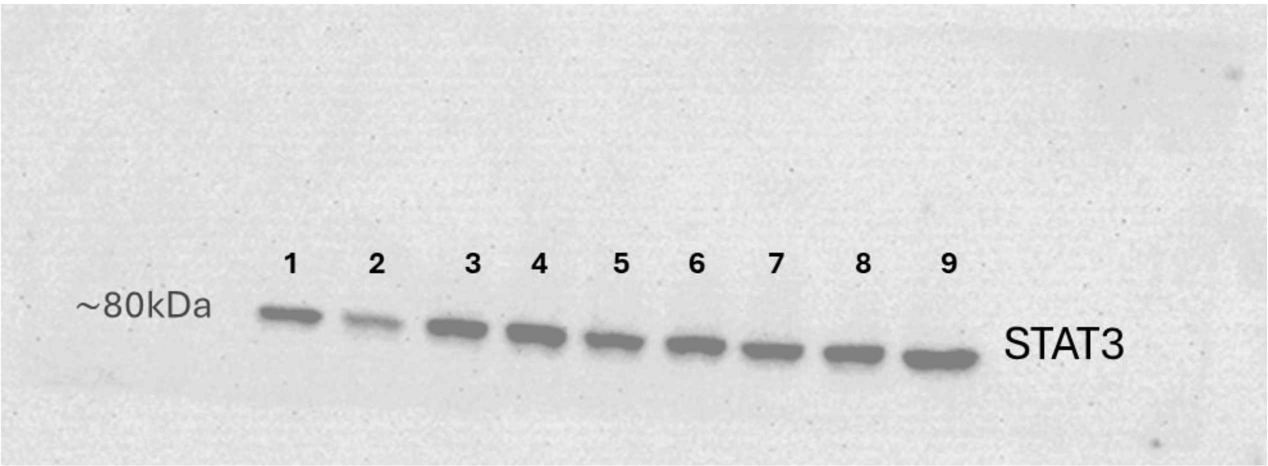

**T41**

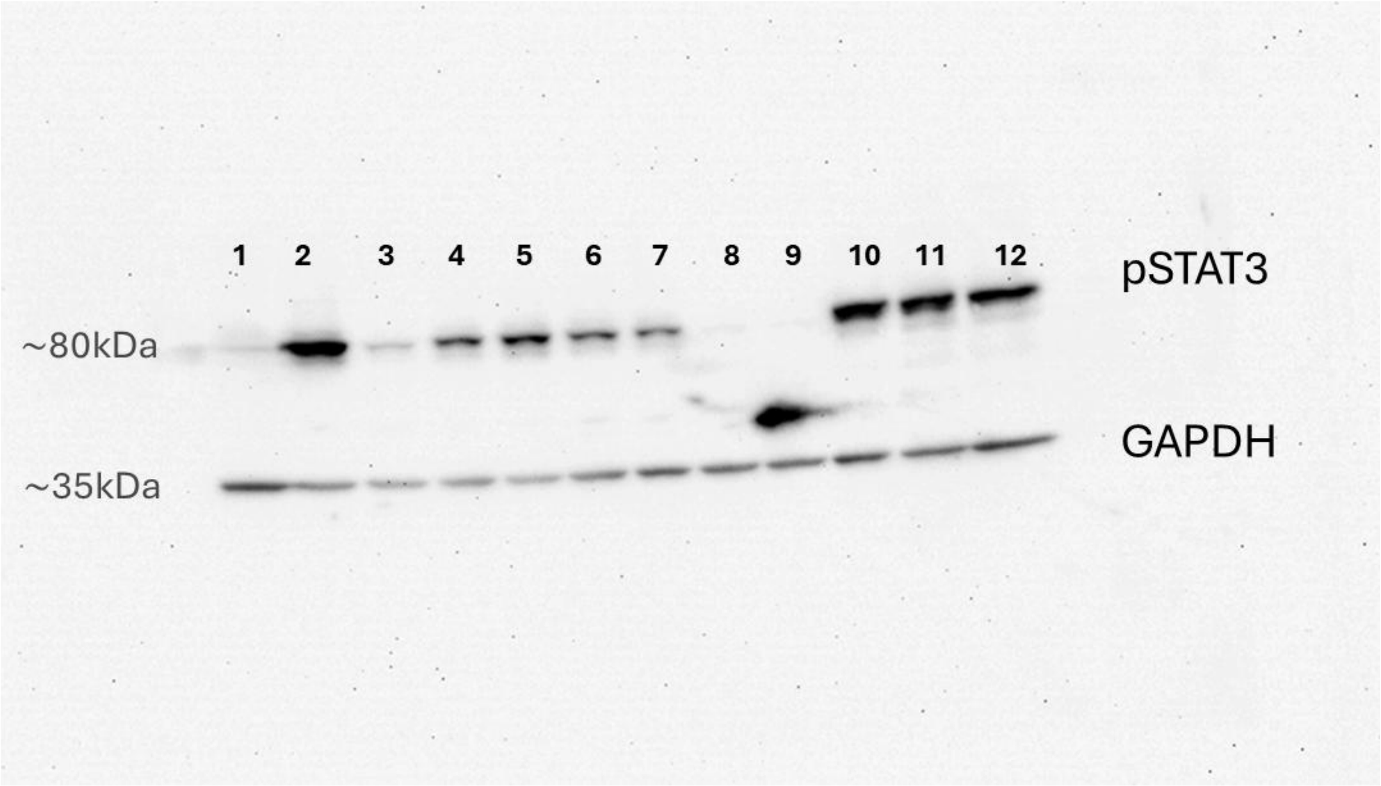
1 2 3 4 5 6 7 8 9 10 11 12
pSTAT3
~80kDa
GAPDH
~35kDa

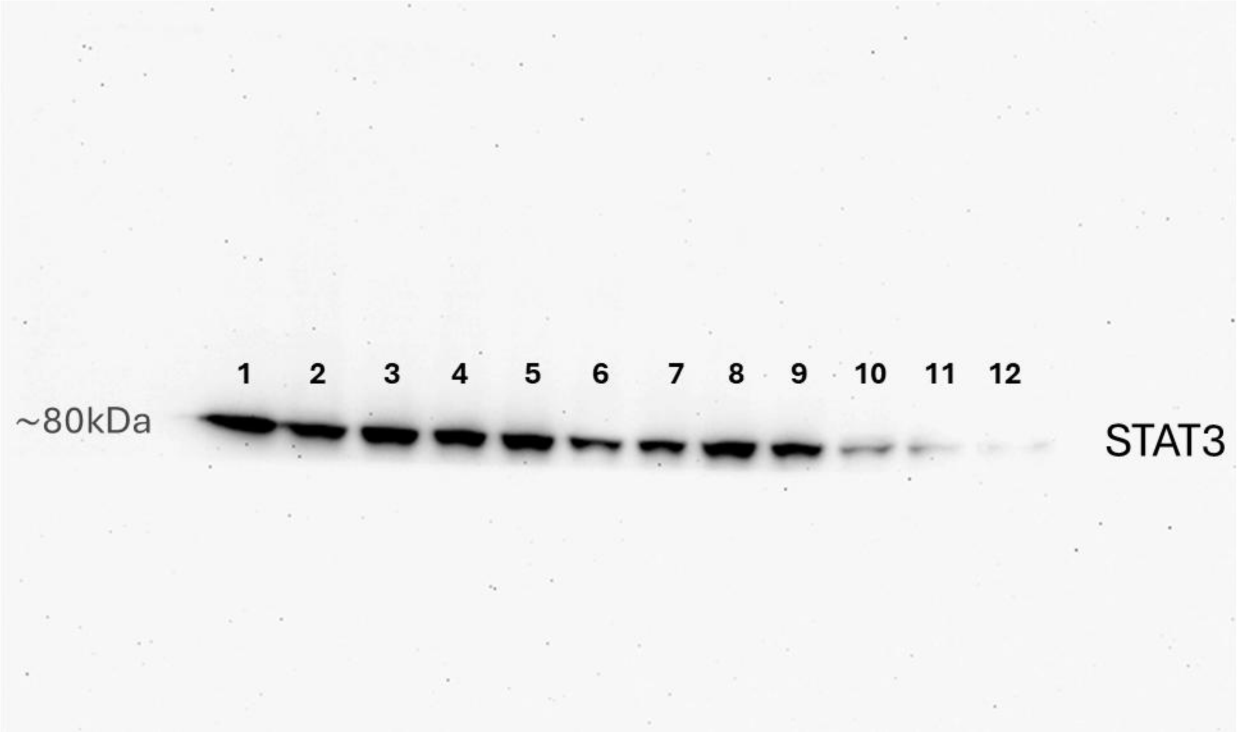
1 2 3 4 5 6 7 8 9 10 11 12
~80kDa
STAT3

**T43**

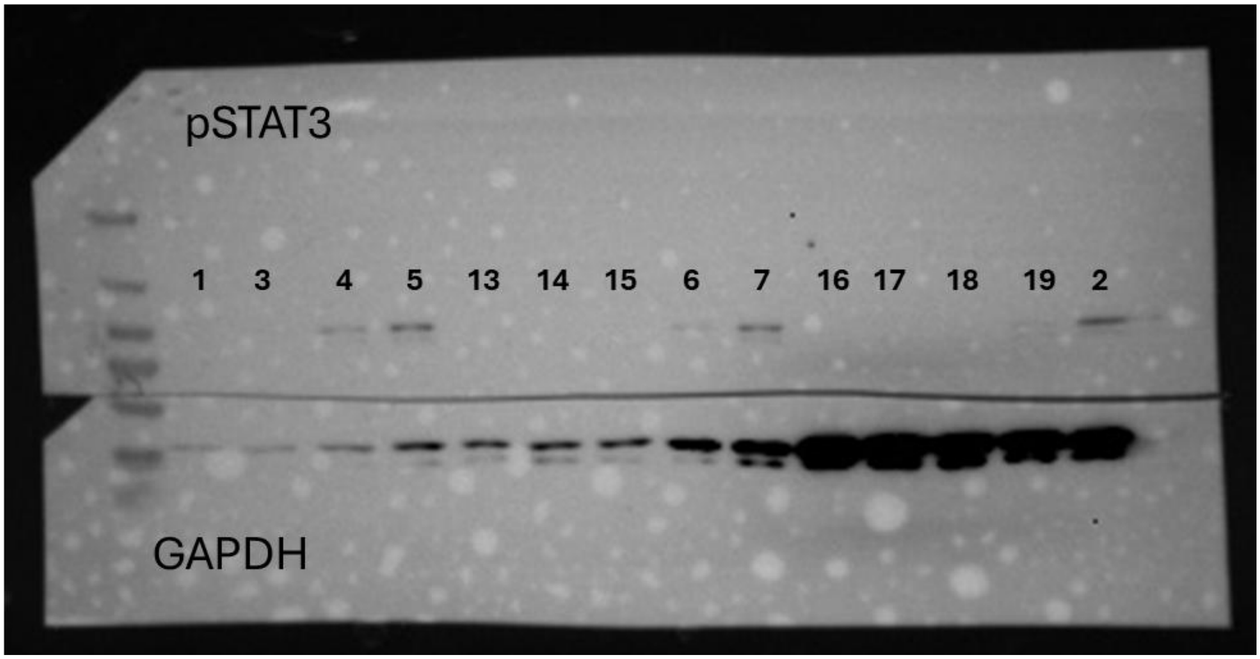

pSTAT3
1 3 4 5 13 14 15 6 7 16 17 18 19 2
GAPDH


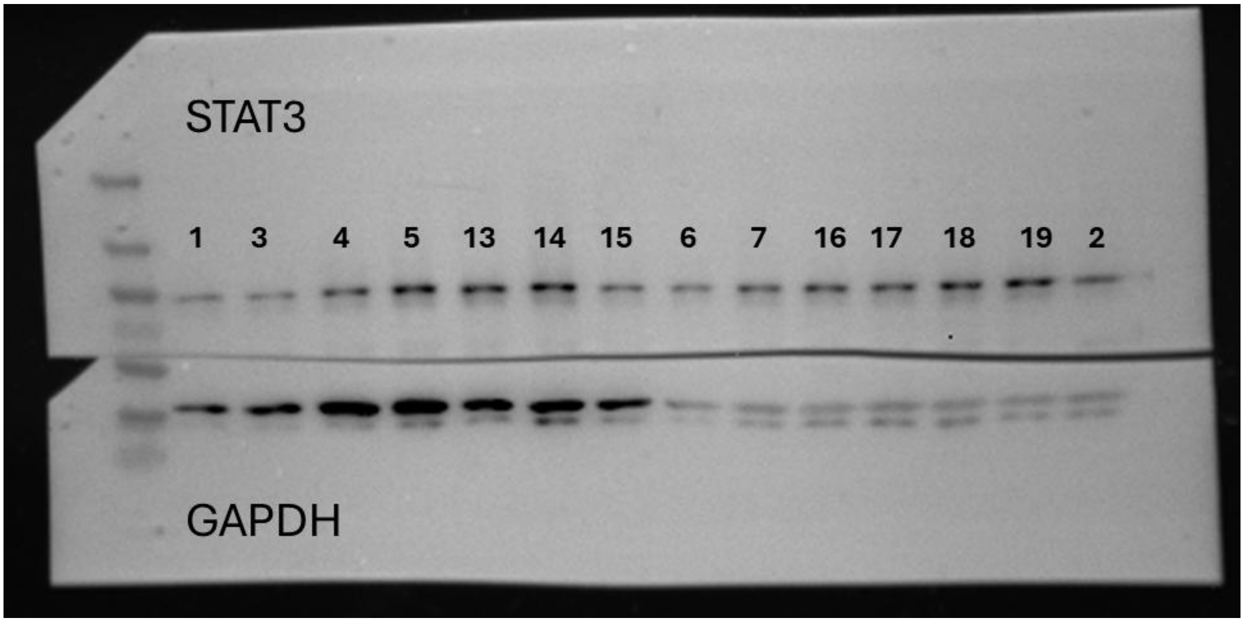

STAT3
1 3 4 5 13 14 15 6 7 16 17 18 19 2
GAPDH
