## Supplementary R Notebooks for "A role for fibroblast and mural cell subsets in a nerve ligation model of neuropathic pain": Suppl_RNotebook_1.rev.html

scRNA\_seq of nerve mesenchyme


Code 

- Show All Code
- Hide All Code
- Download Rmd

### scRNA\_seq of nerve mesenchyme

This is an R Markdown
Notebook. When you execute code within the notebook, the results appear
beneath the code.

Try executing this chunk by clicking the *Run* button within
the chunk or by placing your cursor inside it and pressing
*Ctrl+Shift+Enter*.

The kb-python output used in this notebook was generated as described
by the Pachter lab, here: https://github.com/pachterlab/kb\_python


```
#If you don't have the following packages download them. Next call them with the library() function
library(Seurat)
library(BUSpaRse)
```


```
Registered S3 methods overwritten by 'dbplyr':
  method         from
  print.tbl_lazy     
  print.tbl_sql
```


```
library(ggplot2)
library(tidyverse)
```


```
── Attaching packages ─────────────────────────────────────────────────────────────────────── tidyverse 1.3.1 ──
✔ tibble  3.2.1     ✔ dplyr   1.1.2
✔ tidyr   1.3.0     ✔ stringr 1.5.1
✔ readr   2.0.2     ✔ forcats 0.5.1
✔ purrr   1.0.1     
Warning: package 'tibble' was built under R version 4.1.2Warning: package 'tidyr' was built under R version 4.1.2Warning: package 'purrr' was built under R version 4.1.2Warning: package 'dplyr' was built under R version 4.1.2── Conflicts ────────────────────────────────────────────────────────────────────────── tidyverse_conflicts() ──
✖ dplyr::filter() masks stats::filter()
✖ dplyr::lag()    masks stats::lag()
```


```
library(DropletUtils)
```


```
Warning: package 'DropletUtils' was built under R version 4.1.2Loading required package: SingleCellExperiment
Loading required package: SummarizedExperiment
Loading required package: MatrixGenerics
Loading required package: matrixStats
Warning: package 'matrixStats' was built under R version 4.1.2
Attaching package: 'matrixStats'

The following object is masked from 'package:dplyr':

    count


Attaching package: 'MatrixGenerics'

The following objects are masked from 'package:matrixStats':

    colAlls, colAnyNAs, colAnys, colAvgsPerRowSet, colCollapse, colCounts, colCummaxs, colCummins,
    colCumprods, colCumsums, colDiffs, colIQRDiffs, colIQRs, colLogSumExps, colMadDiffs, colMads,
    colMaxs, colMeans2, colMedians, colMins, colOrderStats, colProds, colQuantiles, colRanges,
    colRanks, colSdDiffs, colSds, colSums2, colTabulates, colVarDiffs, colVars, colWeightedMads,
    colWeightedMeans, colWeightedMedians, colWeightedSds, colWeightedVars, rowAlls, rowAnyNAs,
    rowAnys, rowAvgsPerColSet, rowCollapse, rowCounts, rowCummaxs, rowCummins, rowCumprods,
    rowCumsums, rowDiffs, rowIQRDiffs, rowIQRs, rowLogSumExps, rowMadDiffs, rowMads, rowMaxs,
    rowMeans2, rowMedians, rowMins, rowOrderStats, rowProds, rowQuantiles, rowRanges, rowRanks,
    rowSdDiffs, rowSds, rowSums2, rowTabulates, rowVarDiffs, rowVars, rowWeightedMads,
    rowWeightedMeans, rowWeightedMedians, rowWeightedSds, rowWeightedVars

Loading required package: GenomicRanges
Warning: package 'GenomicRanges' was built under R version 4.1.2Loading required package: stats4
Loading required package: BiocGenerics

Attaching package: 'BiocGenerics'

The following objects are masked from 'package:dplyr':

    combine, intersect, setdiff, union

The following objects are masked from 'package:stats':

    IQR, mad, sd, var, xtabs

The following objects are masked from 'package:base':

    anyDuplicated, append, as.data.frame, basename, cbind, colnames, dirname, do.call, duplicated,
    eval, evalq, Filter, Find, get, grep, grepl, intersect, is.unsorted, lapply, Map, mapply,
    match, mget, order, paste, pmax, pmax.int, pmin, pmin.int, Position, rank, rbind, Reduce,
    rownames, sapply, setdiff, sort, table, tapply, union, unique, unsplit, which.max, which.min

Loading required package: S4Vectors
Warning: package 'S4Vectors' was built under R version 4.1.3
Attaching package: 'S4Vectors'

The following objects are masked from 'package:dplyr':

    first, rename

The following object is masked from 'package:tidyr':

    expand

The following objects are masked from 'package:base':

    expand.grid, I, unname

Loading required package: IRanges

Attaching package: 'IRanges'

The following objects are masked from 'package:dplyr':

    collapse, desc, slice

The following object is masked from 'package:purrr':

    reduce

Loading required package: GenomeInfoDb
Warning: package 'GenomeInfoDb' was built under R version 4.1.2Loading required package: Biobase
Welcome to Bioconductor

    Vignettes contain introductory material; view with 'browseVignettes()'. To cite Bioconductor,
    see 'citation("Biobase")', and for packages 'citation("pkgname")'.


Attaching package: 'Biobase'

The following object is masked from 'package:MatrixGenerics':

    rowMedians

The following objects are masked from 'package:matrixStats':

    anyMissing, rowMedians


Attaching package: 'SummarizedExperiment'

The following object is masked from 'package:SeuratObject':

    Assays

The following object is masked from 'package:Seurat':

    Assays
```


```
library(patchwork)
```


In the following code we first ask R to list the files in the folder
with the kb-python output files (modify the paths to point to our kb
python output folders). Next we create the count matrices. Finally we
have R write the dimensions of the matrix so we can see how many genes
and cells/droplets we have.


```
#lists the output files from Kb-python. Write the path for the output folder in the (). Remember to check that it is "/" and not "\". Make sure your counts_unfiltered folder contains files with the following extensions: barcodes.txt; genes.text; cells_x_genes.mtx

folders <- c("/Users/franziskadenk/Library/CloudStorage/OneDrive-SharedLibraries-King'sCollegeLondon/Denk Lab SharePoint - Mesenchymal_MS/scRNA_seq_data/counts_batch2/ShamD5/counts_unfiltered", 
             "/Users/franziskadenk/Library/CloudStorage/OneDrive-SharedLibraries-King'sCollegeLondon/Denk Lab SharePoint - Mesenchymal_MS/scRNA_seq_data/counts_batch2/ShamM2/counts_unfiltered", 
"/Users/franziskadenk/Library/CloudStorage/OneDrive-SharedLibraries-King'sCollegeLondon/Denk Lab SharePoint - Mesenchymal_MS/scRNA_seq_data/counts_batch2/SNLD5/counts_unfiltered", 
"/Users/franziskadenk/Library/CloudStorage/OneDrive-SharedLibraries-King'sCollegeLondon/Denk Lab SharePoint - Mesenchymal_MS/scRNA_seq_data/counts_batch2/SNLM2/counts_unfiltered",
"/Users/franziskadenk/Library/CloudStorage/OneDrive-SharedLibraries-King'sCollegeLondon/Denk Lab SharePoint - Mesenchymal_MS/scRNA_seq_data/counts_batch1/SHAM/counts_unfiltered",
"/Users/franziskadenk/Library/CloudStorage/OneDrive-SharedLibraries-King'sCollegeLondon/Denk Lab SharePoint - Mesenchymal_MS/scRNA_seq_data/counts_batch1/SNL/counts_unfiltered")
```


```
short_names <- c("SHAM_D5", "SHAM_M2.2", "SNL_D5", "SNL_M2.2", "SHAM_M2.1", "SNL_M2.1")
```


```
results_list <- list()
```


```
#This reads in the following files: .barcodes.txt, .genes.txt and .mtx and makes dgCMatrix files
for (i in seq_along(folders)) {
  folder <- folders[i]
  short_name <- short_names[i]
  
  result <- read_count_output(folder, name = "cells_x_genes", tcc = FALSE)
  
  results_list[[short_name]] <- result
}
```


```
as(<dgTMatrix>, "dgCMatrix") is deprecated since Matrix 1.5-0; do as(., "CsparseMatrix") instead
```


```
summary(results_list)
```


```
Warning: NAs introduced by coercion to integer range
```


```
          Length Class     Mode
SHAM_D5   NA     dgCMatrix S4  
SHAM_M2.2 NA     dgCMatrix S4  
SNL_D5    NA     dgCMatrix S4  
SNL_M2.2  NA     dgCMatrix S4  
SHAM_M2.1 NA     dgCMatrix S4  
SNL_M2.1  NA     dgCMatrix S4
```


```
for (x in results_list) {
  print(dim(x))
}
```


```
[1]  55421 249825
[1]  55421 407894
[1]  55421 438576
[1]  55421 518193
[1]  55421 261842
[1]  55421 311562
```


Now we have created the count matrices with the counts and ensembl
gene IDs. Next we want to change the ensembl gene IDs
(ENSMUSG00000028072) to gene symbols (eg, Ntrk1).


```
#you can see the first few rownames of each matrix with this code.
for (x in results_list) {
  print(head(rownames(x)))
}
```


```
[1] "ENSMUSG00000102693.1" "ENSMUSG00000064842.1" "ENSMUSG00000051951.5"
[4] "ENSMUSG00000102851.1" "ENSMUSG00000103377.1" "ENSMUSG00000104017.1"
[1] "ENSMUSG00000102693.1" "ENSMUSG00000064842.1" "ENSMUSG00000051951.5"
[4] "ENSMUSG00000102851.1" "ENSMUSG00000103377.1" "ENSMUSG00000104017.1"
[1] "ENSMUSG00000102693.1" "ENSMUSG00000064842.1" "ENSMUSG00000051951.5"
[4] "ENSMUSG00000102851.1" "ENSMUSG00000103377.1" "ENSMUSG00000104017.1"
[1] "ENSMUSG00000102693.1" "ENSMUSG00000064842.1" "ENSMUSG00000051951.5"
[4] "ENSMUSG00000102851.1" "ENSMUSG00000103377.1" "ENSMUSG00000104017.1"
[1] "ENSMUSG00000102693.1" "ENSMUSG00000064842.1" "ENSMUSG00000051951.5"
[4] "ENSMUSG00000102851.1" "ENSMUSG00000103377.1" "ENSMUSG00000104017.1"
[1] "ENSMUSG00000102693.1" "ENSMUSG00000064842.1" "ENSMUSG00000051951.5"
[4] "ENSMUSG00000102851.1" "ENSMUSG00000103377.1" "ENSMUSG00000104017.1"
```


```
#import the t2g file that was generated during kb_python during the pseudoalignment 
t2g <- read_tsv("/Users/franziskadenk/Library/CloudStorage/OneDrive-SharedLibraries-King'sCollegeLondon/Denk Lab SharePoint - Mesenchymal_MS/scRNA_seq_data/t2g.txt", col_names = FALSE, col_types = cols(.default = col_character()))
head(t2g)
```


```
#remove the coloumns we don't need
ens2sym<- t2g %>% 
  dplyr::select(X2,X3) %>% distinct()
# a dictionary like vector with names as ensemble ids and values as gene symbol
gene_map<- ens2sym %>% tibble::deframe() 
head(gene_map)
```


```
ENSMUSG00000102693.1 ENSMUSG00000064842.1 ENSMUSG00000051951.5 
     "4933401J01Rik"            "Gm26206"               "Xkr4" 
ENSMUSG00000102851.1 ENSMUSG00000103377.1 ENSMUSG00000104017.1 
           "Gm18956"            "Gm37180"            "Gm37363"
```


```
# change the ensembel id with gene symbol
rownames(results_list$SHAM_D5) <- gene_map[rownames(results_list$SHAM_D5)] %>% unname()
rownames(results_list$SHAM_M2.2) <- gene_map[rownames(results_list$SHAM_M2.2)] %>% unname()
rownames(results_list$SNL_D5) <- gene_map[rownames(results_list$SNL_D5)] %>% unname()
rownames(results_list$SNL_M2.2) <- gene_map[rownames(results_list$SNL_M2.2)] %>% unname()
rownames(results_list$SHAM_M2.1) <- gene_map[rownames(results_list$SHAM_M2.1)] %>% unname()
rownames(results_list$SNL_M2.1) <- gene_map[rownames(results_list$SNL_M2.1)] %>% unname()
```


```
#you can see the first few rownames of each matrix with this code.
for (x in results_list) {
  print(head(rownames(x)))
}
```


```
[1] "4933401J01Rik" "Gm26206"       "Xkr4"          "Gm18956"      
[5] "Gm37180"       "Gm37363"      
[1] "4933401J01Rik" "Gm26206"       "Xkr4"          "Gm18956"      
[5] "Gm37180"       "Gm37363"      
[1] "4933401J01Rik" "Gm26206"       "Xkr4"          "Gm18956"      
[5] "Gm37180"       "Gm37363"      
[1] "4933401J01Rik" "Gm26206"       "Xkr4"          "Gm18956"      
[5] "Gm37180"       "Gm37363"      
[1] "4933401J01Rik" "Gm26206"       "Xkr4"          "Gm18956"      
[5] "Gm37180"       "Gm37363"      
[1] "4933401J01Rik" "Gm26206"       "Xkr4"          "Gm18956"      
[5] "Gm37180"       "Gm37363"
```


```
#how many unique molecular identifiers (UMIs) per barcode? In other words this means how many counts (UMI) pr cell (barcode). This can be used to filter out empty droplets later:
for (x in results_list) {
  tot_counts <- Matrix::colSums(x)
  print(summary(tot_counts))
}
```


```
    Min.  1st Qu.   Median     Mean  3rd Qu.     Max. 
    0.00     1.00     1.00    31.54     1.00 58503.00 
    Min.  1st Qu.   Median     Mean  3rd Qu.     Max. 
    0.00     1.00     1.00    40.76     2.00 20126.00 
    Min.  1st Qu.   Median     Mean  3rd Qu.     Max. 
    0.00     1.00     1.00    44.59     2.00 58932.00 
    Min.  1st Qu.   Median     Mean  3rd Qu.     Max. 
    0.00     1.00     1.00    33.74     2.00 23283.00 
    Min.  1st Qu.   Median     Mean  3rd Qu.     Max. 
    0.00     0.00     1.00    15.66     5.00 19921.00 
   Min. 1st Qu.  Median    Mean 3rd Qu.    Max. 
    0.0     0.0     1.0    24.1     2.0 21623.0
```


```
# Create Seurat Objects
seurat_objects <- list()

# Iterate over the results list and create Seurat objects
for (sample_id in names(results_list)) {
  # Create a Seurat object for the current matrix
  seurat_object <- CreateSeuratObject(counts = results_list[[sample_id]], project = sample_id)
  
  # Add sample ID to meta.data
  seurat_object$sample.ID <- sample_id
  
  # Add the Seurat object to the list
  seurat_objects[[sample_id]] <- seurat_object
}
```


```
Warning: Non-unique features (rownames) present in the input matrix, making uniqueWarning: Non-unique features (rownames) present in the input matrix, making uniqueWarning: Non-unique features (rownames) present in the input matrix, making uniqueWarning: Non-unique features (rownames) present in the input matrix, making uniqueWarning: Non-unique features (rownames) present in the input matrix, making uniqueWarning: Non-unique features (rownames) present in the input matrix, making unique
```


```
seurat_objects
```


```
$SHAM_D5
An object of class Seurat 
55421 features across 249825 samples within 1 assay 
Active assay: RNA (55421 features, 0 variable features)

$SHAM_M2.2
An object of class Seurat 
55421 features across 407894 samples within 1 assay 
Active assay: RNA (55421 features, 0 variable features)

$SNL_D5
An object of class Seurat 
55421 features across 438576 samples within 1 assay 
Active assay: RNA (55421 features, 0 variable features)

$SNL_M2.2
An object of class Seurat 
55421 features across 518193 samples within 1 assay 
Active assay: RNA (55421 features, 0 variable features)

$SHAM_M2.1
An object of class Seurat 
55421 features across 261842 samples within 1 assay 
Active assay: RNA (55421 features, 0 variable features)

$SNL_M2.1
An object of class Seurat 
55421 features across 311562 samples within 1 assay 
Active assay: RNA (55421 features, 0 variable features)
```


```
for (x in seurat_objects) {
  print(x@meta.data)
}
```


```
Srt <- merge(x = seurat_objects[[1]], y = seurat_objects[-1], add.cell.ids=names(seurat_objects))
```


```
Srt@meta.data
```


```
Srt
```


```
An object of class Seurat 
55421 features across 2187892 samples within 1 assay 
Active assay: RNA (55421 features, 0 variable features)
```


```
table(Srt$sample.ID)
```


```
  SHAM_D5 SHAM_M2.1 SHAM_M2.2    SNL_D5  SNL_M2.1  SNL_M2.2 
   249825    261842    407894    438576    311562    518193
```


```
Srt <- CalculateBarcodeInflections(Srt)
BarcodeInflectionsPlot(Srt)
```


```
FeatureScatter(Srt, feature1 = "nCount_RNA", feature2 = "nFeature_RNA")
```


```
Rasterizing points since number of points exceeds 100,000.
To disable this behavior set `raster=FALSE`
```


cut-off of nCount\_RNA chosen to be relatively high, as lower cut-off
(nCount\_RNA > 100 was also tried) ended up with some clusters being
distinguished mainly by genes indicating cellular stress.


```
Srt2 <- SubsetByBarcodeInflections(Srt)
Srt2 <- subset(Srt2, subset = nCount_RNA > 1000)
```


```
#finding a subset of mitochondrial genes. "^mt-" for mouse. 
Srt2[["percent.mt"]] <- PercentageFeatureSet(Srt2, pattern = "^mt-")
```


```
plot1 <- FeatureScatter(Srt2, feature1 = "nCount_RNA", feature2 = "percent.mt")
plot2 <- FeatureScatter(Srt2, feature1 = "nCount_RNA", feature2 = "nFeature_RNA")
plot1 + plot2
```


```
#we make some plots to further clean up the data:
VlnPlot(Srt2, features = c("nFeature_RNA", "nCount_RNA", "percent.mt"), ncol = 3)
```


```
#remove cells with high mitochondrial gene content: 
Srt3 <- subset(Srt2, subset = percent.mt < 10)
```


```
#check that data look reasonable post QC:
VlnPlot(Srt3, features = c("nFeature_RNA", "nCount_RNA", "percent.mt"), ncol = 3)
```


```
#check number of cells per biological sample:
table(Srt3$orig.ident)
```


```
  SHAM_D5 SHAM_M2.1 SHAM_M2.2    SNL_D5  SNL_M2.1  SNL_M2.2 
     1095       543      3390      1481      2325      3775
```


add some more metadata:


```
Srt3 <- (RenameIdents(Srt3, `SHAM_D5` = "SHAM", `SHAM_M2.1` = "SHAM", `SHAM_M2.2` = "SHAM", `SNL_D5` = "PSNL", `SNL_M2.1` = "PSNL", `SNL_M2.2` = "PSNL"))
Srt3[["Injury"]] <- Idents(object = Srt3)
```


```
Idents(Srt3) <- Srt3$orig.ident
Srt3 <- (RenameIdents(Srt3, `SHAM_D5` = "batch2", `SHAM_M2.1` = "batch1", `SHAM_M2.2` = "batch2", `SNL_D5` = "batch2", `SNL_M2.1` = "batch1", `SNL_M2.2` = "batch2"))
Srt3[["batch"]] <- Idents(object = Srt3)
```


```
Idents(Srt3) <- Srt3$orig.ident
Srt3 <- (RenameIdents(Srt3, `SHAM_D5` = "day5", `SHAM_M2.1` = "month2", `SHAM_M2.2` = "month2", `SNL_D5` = "day5", `SNL_M2.1` = "month2", `SNL_M2.2` = "month2"))
Srt3[["Timepoint"]] <- Idents(object = Srt3)
```


```
Srt3@meta.data
```


run unsupervised clustering first without integration to see the
extent of the batch effect


```
RNA_N <- NormalizeData(Srt3)
```


```
Performing log-normalization
0%   10   20   30   40   50   60   70   80   90   100%
[----|----|----|----|----|----|----|----|----|----|
**************************************************|
```


```
RNA_V <- FindVariableFeatures(RNA_N)
```


```
Calculating gene variances
0%   10   20   30   40   50   60   70   80   90   100%
[----|----|----|----|----|----|----|----|----|----|
**************************************************|
Calculating feature variances of standardized and clipped values
0%   10   20   30   40   50   60   70   80   90   100%
[----|----|----|----|----|----|----|----|----|----|
**************************************************|
```


```
RNA_S <- ScaleData(RNA_V)
```


```
Centering and scaling data matrix

  |                                                                                                            
  |                                                                                                      |   0%
  |                                                                                                            
  |===================================================                                                   |  50%
  |                                                                                                            
  |======================================================================================================| 100%
```


```
PCA <- RunPCA(RNA_S, verbose=TRUE)
```


```
PC_ 1 
Positive:  Col1a2, Col3a1, Col1a1, Ogn, Igfbp6, Mfap5, Gpc3, Tnxb, Lum, C3 
       Col6a2, Apod, Dpt, Dpep1, Igf1, Fn1, Gas1, Htra3, Ccl11, Col6a3 
       Col5a1, Smoc2, Col5a2, Sfrp4, Myoc, Itm2a, Col14a1, Ctsk, Gpx3, Cd248 
Negative:  Egfl7, Cd93, Ctla2a, Esam, Ptprb, Plvap, Mmrn2, Sox18, Emcn, S1pr1 
       Aplnr, Adgrf5, Grrp1, Myct1, Rasip1, Cdh5, Ehd4, Flt1, Fam167b, Kdr 
       Tie1, Tmem252, Adgrl4, Lrg1, Icam2, Fli1, Fabp4, Mecom, Eng, Gimap4 
PC_ 2 
Positive:  Ly6a, Fn1, Ly6c1, Fstl1, Tnxb, C3, Dpt, Igf1, Mfap5, Col3a1 
       Fbn1, Col6a2, Igfbp6, Ly6e, Fbln2, Ogn, Cd248, Loxl1, Fndc1, Col6a3 
       Col14a1, Klf4, Aebp1, Gas1, Htra3, Lsp1, Gpx3, Gpc3, Dpep1, Prrx1 
Negative:  Cryab, Plp1, Gatm, Mpz, Cnp, Kcna1, Mal, Gpm6b, Mbp, Itgb8 
       Dbi, Sox10, Cadm1, Cadm4, Stmn1, Atp1a2, Ank3, S100a4, Ldhb, Kcna2 
       Gjc3, Lbh, Erbb3, Gas2l3, Sostdc1, Pou3f1, Pdgfa, Foxd3, Cdh19, Ednrb 
PC_ 3 
Positive:  Plp1, Gatm, Gpm6b, Cnp, Itgb8, Stmn1, Mpz, Kcna1, Arpc1b, Mal 
       Gas7, Cadm4, Cadm1, Sox10, Ank3, Mbp, Dbi, Erbb3, Gas2l3, Gjc3 
       S100a4, Sostdc1, Pmp22, Foxd3, Cdh19, Arpc1a, Fabp5, Itga6, Pou3f1, Reln 
Negative:  Acta2, Myh11, Tpm2, Myl9, Gucy1a1, Tagln, Mylk, Pcp4l1, Mustn1, Lmod1 
       Ptp4a3, Notch3, Rcan2, Des, Npy1r, Tesc, Gm13889, Mrvi1, Tinagl1, Rgs7bp 
       Nrip2, Filip1l, Ndufa4l2, Olfr558, Ppp1r14a, Dgkb, Rgs4, Map3k20, Rbpms2, Pdlim3 
PC_ 4 
Positive:  Vim, Lgals1, Hsp90b1, Crip1, Fkbp1a, Npm1, Col1a2, Tubb5, Fbln2, Ly6a 
       Calr, Aebp1, Tuba1a, Ly6c1, Fstl1, Tagln2, Pmepa1, Ece1, Pdlim1, Col5a2 
       Col4a1, Ran, Heg1, Col1a1, Col4a2, Col5a1, Dbi, Col14a1, Cald1, Dpysl3 
Negative:  Tyrobp, Il1b, S100a8, S100a9, Fcer1g, Cd52, Hdc, Clec4d, Ccr1, Ptprc 
       Csf3r, Cxcl2, Plek, Coro1a, Samsn1, Mxd1, Slpi, Il1r2, Acod1, Cd53 
       Clec4e, Sorl1, Clec7a, Mmp9, Trem1, Lilr4b, Gm44292, Spi1, Cxcr2, Cd300ld 
PC_ 5 
Positive:  Timp1, S100a11, Mki67, Cdca3, Tpx2, Tnc, Ccnb2, Birc5, Postn, Anxa1 
       Nt5dc2, Racgap1, Uchl1, Prc1, Dclk1, Cenpf, Hmmr, Nes, Adamts5, Crlf1 
       Cd44, Ckap2l, Cenpe, Cthrc1, Cdkn3, Ccdc80, Col14a1, Cks2, Ccna2, Pmepa1 
Negative:  Smoc2, Tenm2, Sfrp5, Cldn1, Klf5, Cdh5, F11r, Cavin2, Apoe, Trib2 
       Thbs4, Lypd2, Cp, Clec1a, Krt19, Ramp2, Apod, Jam2, Itgb4, Fmo2 
       C130074G19Rik, Ptprb, Slc2a1, Wnt6, Abcb1a, Ptch1, Egfl7, Fxyd6, Fzd2, Col15a1
```


```
ElbowPlot(PCA)
```


```
DimPlot(cbmc, split.by="Injury", label=TRUE,  cols = c("5" = "#ffffba", "2" = "#ffb3ba", "0" = "#ffdfba", "4" = "#d5a6bd", "1" = "#baffc9", "7" = "#559e83", "9" = "#f2f6c3", "8" = "#bae1ff", "3" = "blue","6" = "#cccccc", "10" = "#559e83"), pt.size=0.7)
```


```
DimPlot(cbmc, split.by="Timepoint", label=TRUE,  cols = c("5" = "#ffffba", "2" = "#ffb3ba", "0" = "#ffdfba", "4" = "#d5a6bd", "1" = "#baffc9", "7" = "#559e83", "9" = "#f2f6c3", "8" = "#bae1ff", "3" = "blue","6" = "#cccccc", "10" = "#559e83"), pt.size=0.7)
```


continued without integration first, as technical batch seems to be
superseded by cell type, at least at ‘macro-level’. obtain cluster
markers.


```
RNA_markers <- FindAllMarkers(cbmc, min.diff.pct = 0.2)
```


```
Calculating cluster 0
```


```
  |                                                  | 0 % ~calculating  
  |+                                                 | 1 % ~10s          
  |++                                                | 3 % ~10s          
  |++                                                | 4 % ~10s          
  |+++                                               | 5 % ~10s          
  |++++                                              | 7 % ~09s          
  |++++                                              | 8 % ~09s          
  |+++++                                             | 9 % ~09s          
  |++++++                                            | 11% ~09s          
  |++++++                                            | 12% ~08s          
  |+++++++                                           | 13% ~08s          
  |++++++++                                          | 14% ~08s          
  |++++++++                                          | 16% ~08s          
  |+++++++++                                         | 17% ~08s          
  |++++++++++                                        | 18% ~08s          
  |++++++++++                                        | 20% ~08s          
  |+++++++++++                                       | 21% ~07s          
  |++++++++++++                                      | 22% ~07s          
  |++++++++++++                                      | 24% ~07s          
  |+++++++++++++                                     | 25% ~07s          
  |++++++++++++++                                    | 26% ~07s          
  |++++++++++++++                                    | 28% ~07s          
  |+++++++++++++++                                   | 29% ~07s          
  |++++++++++++++++                                  | 30% ~06s          
  |++++++++++++++++                                  | 32% ~06s          
  |+++++++++++++++++                                 | 33% ~06s          
  |++++++++++++++++++                                | 34% ~06s          
  |++++++++++++++++++                                | 36% ~06s          
  |+++++++++++++++++++                               | 37% ~06s          
  |++++++++++++++++++++                              | 38% ~06s          
  |++++++++++++++++++++                              | 39% ~06s          
  |+++++++++++++++++++++                             | 41% ~05s          
  |++++++++++++++++++++++                            | 42% ~05s          
  |++++++++++++++++++++++                            | 43% ~05s          
  |+++++++++++++++++++++++                           | 45% ~05s          
  |++++++++++++++++++++++++                          | 46% ~05s          
  |++++++++++++++++++++++++                          | 47% ~05s          
  |+++++++++++++++++++++++++                         | 49% ~05s          
  |+++++++++++++++++++++++++                         | 50% ~04s          
  |++++++++++++++++++++++++++                        | 51% ~04s          
  |+++++++++++++++++++++++++++                       | 53% ~04s          
  |+++++++++++++++++++++++++++                       | 54% ~04s          
  |++++++++++++++++++++++++++++                      | 55% ~04s          
  |+++++++++++++++++++++++++++++                     | 57% ~04s          
  |+++++++++++++++++++++++++++++                     | 58% ~04s          
  |++++++++++++++++++++++++++++++                    | 59% ~04s          
  |+++++++++++++++++++++++++++++++                   | 61% ~04s          
  |+++++++++++++++++++++++++++++++                   | 62% ~03s          
  |++++++++++++++++++++++++++++++++                  | 63% ~03s          
  |+++++++++++++++++++++++++++++++++                 | 64% ~03s          
  |+++++++++++++++++++++++++++++++++                 | 66% ~03s          
  |++++++++++++++++++++++++++++++++++                | 67% ~03s          
  |+++++++++++++++++++++++++++++++++++               | 68% ~03s          
  |+++++++++++++++++++++++++++++++++++               | 70% ~03s          
  |++++++++++++++++++++++++++++++++++++              | 71% ~03s          
  |+++++++++++++++++++++++++++++++++++++             | 72% ~02s          
  |+++++++++++++++++++++++++++++++++++++             | 74% ~02s          
  |++++++++++++++++++++++++++++++++++++++            | 75% ~02s          
  |+++++++++++++++++++++++++++++++++++++++           | 76% ~02s          
  |+++++++++++++++++++++++++++++++++++++++           | 78% ~02s          
  |++++++++++++++++++++++++++++++++++++++++          | 79% ~02s          
  |+++++++++++++++++++++++++++++++++++++++++         | 80% ~02s          
  |+++++++++++++++++++++++++++++++++++++++++         | 82% ~02s          
  |++++++++++++++++++++++++++++++++++++++++++        | 83% ~02s          
  |+++++++++++++++++++++++++++++++++++++++++++       | 84% ~01s          
  |+++++++++++++++++++++++++++++++++++++++++++       | 86% ~01s          
  |++++++++++++++++++++++++++++++++++++++++++++      | 87% ~01s          
  |+++++++++++++++++++++++++++++++++++++++++++++     | 88% ~01s          
  |+++++++++++++++++++++++++++++++++++++++++++++     | 89% ~01s          
  |++++++++++++++++++++++++++++++++++++++++++++++    | 91% ~01s          
  |+++++++++++++++++++++++++++++++++++++++++++++++   | 92% ~01s          
  |+++++++++++++++++++++++++++++++++++++++++++++++   | 93% ~01s          
  |++++++++++++++++++++++++++++++++++++++++++++++++  | 95% ~00s          
  |+++++++++++++++++++++++++++++++++++++++++++++++++ | 96% ~00s          
  |+++++++++++++++++++++++++++++++++++++++++++++++++ | 97% ~00s          
  |++++++++++++++++++++++++++++++++++++++++++++++++++| 99% ~00s          
  |++++++++++++++++++++++++++++++++++++++++++++++++++| 100% elapsed=09s
```


```
Calculating cluster 1
```


```
  |                                                  | 0 % ~calculating  
  |+                                                 | 1 % ~18s          
  |++                                                | 2 % ~17s          
  |++                                                | 3 % ~17s          
  |+++                                               | 4 % ~17s          
  |+++                                               | 5 % ~16s          
  |++++                                              | 7 % ~16s          
  |++++                                              | 8 % ~16s          
  |+++++                                             | 9 % ~16s          
  |+++++                                             | 10% ~15s          
  |++++++                                            | 11% ~15s          
  |+++++++                                           | 12% ~15s          
  |+++++++                                           | 13% ~15s          
  |++++++++                                          | 14% ~15s          
  |++++++++                                          | 15% ~15s          
  |+++++++++                                         | 16% ~15s          
  |+++++++++                                         | 18% ~14s          
  |++++++++++                                        | 19% ~14s          
  |++++++++++                                        | 20% ~14s          
  |+++++++++++                                       | 21% ~14s          
  |+++++++++++                                       | 22% ~14s          
  |++++++++++++                                      | 23% ~13s          
  |+++++++++++++                                     | 24% ~13s          
  |+++++++++++++                                     | 25% ~13s          
  |++++++++++++++                                    | 26% ~13s          
  |++++++++++++++                                    | 27% ~12s          
  |+++++++++++++++                                   | 29% ~12s          
  |+++++++++++++++                                   | 30% ~12s          
  |++++++++++++++++                                  | 31% ~12s          
  |++++++++++++++++                                  | 32% ~12s          
  |+++++++++++++++++                                 | 33% ~12s          
  |++++++++++++++++++                                | 34% ~11s          
  |++++++++++++++++++                                | 35% ~11s          
  |+++++++++++++++++++                               | 36% ~11s          
  |+++++++++++++++++++                               | 37% ~11s          
  |++++++++++++++++++++                              | 38% ~11s          
  |++++++++++++++++++++                              | 40% ~10s          
  |+++++++++++++++++++++                             | 41% ~10s          
  |+++++++++++++++++++++                             | 42% ~10s          
  |++++++++++++++++++++++                            | 43% ~10s          
  |++++++++++++++++++++++                            | 44% ~10s          
  |+++++++++++++++++++++++                           | 45% ~10s          
  |++++++++++++++++++++++++                          | 46% ~10s          
  |++++++++++++++++++++++++                          | 47% ~10s          
  |+++++++++++++++++++++++++                         | 48% ~09s          
  |+++++++++++++++++++++++++                         | 49% ~09s          
  |++++++++++++++++++++++++++                        | 51% ~09s          
  |++++++++++++++++++++++++++                        | 52% ~09s          
  |+++++++++++++++++++++++++++                       | 53% ~09s          
  |+++++++++++++++++++++++++++                       | 54% ~08s          
  |++++++++++++++++++++++++++++                      | 55% ~08s          
  |+++++++++++++++++++++++++++++                     | 56% ~08s          
  |+++++++++++++++++++++++++++++                     | 57% ~08s          
  |++++++++++++++++++++++++++++++                    | 58% ~07s          
  |++++++++++++++++++++++++++++++                    | 59% ~07s          
  |+++++++++++++++++++++++++++++++                   | 60% ~07s          
  |+++++++++++++++++++++++++++++++                   | 62% ~07s          
  |++++++++++++++++++++++++++++++++                  | 63% ~07s          
  |++++++++++++++++++++++++++++++++                  | 64% ~06s          
  |+++++++++++++++++++++++++++++++++                 | 65% ~06s          
  |+++++++++++++++++++++++++++++++++                 | 66% ~06s          
  |++++++++++++++++++++++++++++++++++                | 67% ~06s          
  |+++++++++++++++++++++++++++++++++++               | 68% ~06s          
  |+++++++++++++++++++++++++++++++++++               | 69% ~05s          
  |++++++++++++++++++++++++++++++++++++              | 70% ~05s          
  |++++++++++++++++++++++++++++++++++++              | 71% ~05s          
  |+++++++++++++++++++++++++++++++++++++             | 73% ~05s          
  |+++++++++++++++++++++++++++++++++++++             | 74% ~05s          
  |++++++++++++++++++++++++++++++++++++++            | 75% ~04s          
  |++++++++++++++++++++++++++++++++++++++            | 76% ~04s          
  |+++++++++++++++++++++++++++++++++++++++           | 77% ~04s          
  |++++++++++++++++++++++++++++++++++++++++          | 78% ~04s          
  |++++++++++++++++++++++++++++++++++++++++          | 79% ~04s          
  |+++++++++++++++++++++++++++++++++++++++++         | 80% ~03s          
  |+++++++++++++++++++++++++++++++++++++++++         | 81% ~03s          
  |++++++++++++++++++++++++++++++++++++++++++        | 82% ~03s          
  |++++++++++++++++++++++++++++++++++++++++++        | 84% ~03s          
  |+++++++++++++++++++++++++++++++++++++++++++       | 85% ~03s          
  |+++++++++++++++++++++++++++++++++++++++++++       | 86% ~03s          
  |++++++++++++++++++++++++++++++++++++++++++++      | 87% ~02s          
  |++++++++++++++++++++++++++++++++++++++++++++      | 88% ~02s          
  |+++++++++++++++++++++++++++++++++++++++++++++     | 89% ~02s          
  |++++++++++++++++++++++++++++++++++++++++++++++    | 90% ~02s          
  |++++++++++++++++++++++++++++++++++++++++++++++    | 91% ~02s          
  |+++++++++++++++++++++++++++++++++++++++++++++++   | 92% ~01s          
  |+++++++++++++++++++++++++++++++++++++++++++++++   | 93% ~01s          
  |++++++++++++++++++++++++++++++++++++++++++++++++  | 95% ~01s          
  |++++++++++++++++++++++++++++++++++++++++++++++++  | 96% ~01s          
  |+++++++++++++++++++++++++++++++++++++++++++++++++ | 97% ~01s          
  |+++++++++++++++++++++++++++++++++++++++++++++++++ | 98% ~00s          
  |++++++++++++++++++++++++++++++++++++++++++++++++++| 99% ~00s          
  |++++++++++++++++++++++++++++++++++++++++++++++++++| 100% elapsed=17s
```


```
Calculating cluster 2
```


```
  |                                                  | 0 % ~calculating  
  |+                                                 | 1 % ~06s          
  |++                                                | 3 % ~06s          
  |++                                                | 4 % ~06s          
  |+++                                               | 5 % ~06s          
  |++++                                              | 7 % ~06s          
  |++++                                              | 8 % ~06s          
  |+++++                                             | 9 % ~06s          
  |++++++                                            | 11% ~05s          
  |++++++                                            | 12% ~05s          
  |+++++++                                           | 13% ~05s          
  |++++++++                                          | 14% ~05s          
  |++++++++                                          | 16% ~05s          
  |+++++++++                                         | 17% ~05s          
  |++++++++++                                        | 18% ~05s          
  |++++++++++                                        | 20% ~05s          
  |+++++++++++                                       | 21% ~05s          
  |++++++++++++                                      | 22% ~05s          
  |++++++++++++                                      | 24% ~05s          
  |+++++++++++++                                     | 25% ~05s          
  |++++++++++++++                                    | 26% ~05s          
  |++++++++++++++                                    | 28% ~04s          
  |+++++++++++++++                                   | 29% ~04s          
  |++++++++++++++++                                  | 30% ~04s          
  |++++++++++++++++                                  | 32% ~04s          
  |+++++++++++++++++                                 | 33% ~04s          
  |++++++++++++++++++                                | 34% ~04s          
  |++++++++++++++++++                                | 36% ~04s          
  |+++++++++++++++++++                               | 37% ~04s          
  |++++++++++++++++++++                              | 38% ~04s          
  |++++++++++++++++++++                              | 39% ~04s          
  |+++++++++++++++++++++                             | 41% ~04s          
  |++++++++++++++++++++++                            | 42% ~04s          
  |++++++++++++++++++++++                            | 43% ~04s          
  |+++++++++++++++++++++++                           | 45% ~03s          
  |++++++++++++++++++++++++                          | 46% ~03s          
  |++++++++++++++++++++++++                          | 47% ~03s          
  |+++++++++++++++++++++++++                         | 49% ~03s          
  |+++++++++++++++++++++++++                         | 50% ~03s          
  |++++++++++++++++++++++++++                        | 51% ~03s          
  |+++++++++++++++++++++++++++                       | 53% ~03s          
  |+++++++++++++++++++++++++++                       | 54% ~03s          
  |++++++++++++++++++++++++++++                      | 55% ~03s          
  |+++++++++++++++++++++++++++++                     | 57% ~03s          
  |+++++++++++++++++++++++++++++                     | 58% ~03s          
  |++++++++++++++++++++++++++++++                    | 59% ~03s          
  |+++++++++++++++++++++++++++++++                   | 61% ~02s          
  |+++++++++++++++++++++++++++++++                   | 62% ~02s          
  |++++++++++++++++++++++++++++++++                  | 63% ~02s          
  |+++++++++++++++++++++++++++++++++                 | 64% ~02s          
  |+++++++++++++++++++++++++++++++++                 | 66% ~02s          
  |++++++++++++++++++++++++++++++++++                | 67% ~02s          
  |+++++++++++++++++++++++++++++++++++               | 68% ~02s          
  |+++++++++++++++++++++++++++++++++++               | 70% ~02s          
  |++++++++++++++++++++++++++++++++++++              | 71% ~02s          
  |+++++++++++++++++++++++++++++++++++++             | 72% ~02s          
  |+++++++++++++++++++++++++++++++++++++             | 74% ~02s          
  |++++++++++++++++++++++++++++++++++++++            | 75% ~02s          
  |+++++++++++++++++++++++++++++++++++++++           | 76% ~01s          
  |+++++++++++++++++++++++++++++++++++++++           | 78% ~01s          
  |++++++++++++++++++++++++++++++++++++++++          | 79% ~01s          
  |+++++++++++++++++++++++++++++++++++++++++         | 80% ~01s          
  |+++++++++++++++++++++++++++++++++++++++++         | 82% ~01s          
  |++++++++++++++++++++++++++++++++++++++++++        | 83% ~01s          
  |+++++++++++++++++++++++++++++++++++++++++++       | 84% ~01s          
  |+++++++++++++++++++++++++++++++++++++++++++       | 86% ~01s          
  |++++++++++++++++++++++++++++++++++++++++++++      | 87% ~01s          
  |+++++++++++++++++++++++++++++++++++++++++++++     | 88% ~01s          
  |+++++++++++++++++++++++++++++++++++++++++++++     | 89% ~01s          
  |++++++++++++++++++++++++++++++++++++++++++++++    | 91% ~01s          
  |+++++++++++++++++++++++++++++++++++++++++++++++   | 92% ~00s          
  |+++++++++++++++++++++++++++++++++++++++++++++++   | 93% ~00s          
  |++++++++++++++++++++++++++++++++++++++++++++++++  | 95% ~00s          
  |+++++++++++++++++++++++++++++++++++++++++++++++++ | 96% ~00s          
  |+++++++++++++++++++++++++++++++++++++++++++++++++ | 97% ~00s          
  |++++++++++++++++++++++++++++++++++++++++++++++++++| 99% ~00s          
  |++++++++++++++++++++++++++++++++++++++++++++++++++| 100% elapsed=06s
```


```
Calculating cluster 3
```


```
  |                                                  | 0 % ~calculating  
  |+                                                 | 1 % ~21s          
  |++                                                | 2 % ~20s          
  |++                                                | 3 % ~20s          
  |+++                                               | 4 % ~20s          
  |+++                                               | 6 % ~19s          
  |++++                                              | 7 % ~19s          
  |++++                                              | 8 % ~19s          
  |+++++                                             | 9 % ~19s          
  |+++++                                             | 10% ~18s          
  |++++++                                            | 11% ~18s          
  |+++++++                                           | 12% ~18s          
  |+++++++                                           | 13% ~20s          
  |++++++++                                          | 14% ~19s          
  |++++++++                                          | 16% ~19s          
  |+++++++++                                         | 17% ~19s          
  |+++++++++                                         | 18% ~18s          
  |++++++++++                                        | 19% ~18s          
  |++++++++++                                        | 20% ~18s          
  |+++++++++++                                       | 21% ~17s          
  |++++++++++++                                      | 22% ~17s          
  |++++++++++++                                      | 23% ~17s          
  |+++++++++++++                                     | 24% ~16s          
  |+++++++++++++                                     | 26% ~16s          
  |++++++++++++++                                    | 27% ~16s          
  |++++++++++++++                                    | 28% ~16s          
  |+++++++++++++++                                   | 29% ~15s          
  |+++++++++++++++                                   | 30% ~15s          
  |++++++++++++++++                                  | 31% ~15s          
  |+++++++++++++++++                                 | 32% ~15s          
  |+++++++++++++++++                                 | 33% ~14s          
  |++++++++++++++++++                                | 34% ~14s          
  |++++++++++++++++++                                | 36% ~14s          
  |+++++++++++++++++++                               | 37% ~14s          
  |+++++++++++++++++++                               | 38% ~13s          
  |++++++++++++++++++++                              | 39% ~13s          
  |++++++++++++++++++++                              | 40% ~13s          
  |+++++++++++++++++++++                             | 41% ~13s          
  |++++++++++++++++++++++                            | 42% ~12s          
  |++++++++++++++++++++++                            | 43% ~12s          
  |+++++++++++++++++++++++                           | 44% ~12s          
  |+++++++++++++++++++++++                           | 46% ~12s          
  |++++++++++++++++++++++++                          | 47% ~11s          
  |++++++++++++++++++++++++                          | 48% ~11s          
  |+++++++++++++++++++++++++                         | 49% ~11s          
  |+++++++++++++++++++++++++                         | 50% ~11s          
  |++++++++++++++++++++++++++                        | 51% ~10s          
  |+++++++++++++++++++++++++++                       | 52% ~10s          
  |+++++++++++++++++++++++++++                       | 53% ~10s          
  |++++++++++++++++++++++++++++                      | 54% ~10s          
  |++++++++++++++++++++++++++++                      | 56% ~09s          
  |+++++++++++++++++++++++++++++                     | 57% ~09s          
  |+++++++++++++++++++++++++++++                     | 58% ~09s          
  |++++++++++++++++++++++++++++++                    | 59% ~09s          
  |++++++++++++++++++++++++++++++                    | 60% ~09s          
  |+++++++++++++++++++++++++++++++                   | 61% ~08s          
  |++++++++++++++++++++++++++++++++                  | 62% ~08s          
  |++++++++++++++++++++++++++++++++                  | 63% ~08s          
  |+++++++++++++++++++++++++++++++++                 | 64% ~08s          
  |+++++++++++++++++++++++++++++++++                 | 66% ~07s          
  |++++++++++++++++++++++++++++++++++                | 67% ~07s          
  |++++++++++++++++++++++++++++++++++                | 68% ~07s          
  |+++++++++++++++++++++++++++++++++++               | 69% ~07s          
  |+++++++++++++++++++++++++++++++++++               | 70% ~06s          
  |++++++++++++++++++++++++++++++++++++              | 71% ~06s          
  |+++++++++++++++++++++++++++++++++++++             | 72% ~06s          
  |+++++++++++++++++++++++++++++++++++++             | 73% ~06s          
  |++++++++++++++++++++++++++++++++++++++            | 74% ~05s          
  |++++++++++++++++++++++++++++++++++++++            | 76% ~05s          
  |+++++++++++++++++++++++++++++++++++++++           | 77% ~05s          
  |+++++++++++++++++++++++++++++++++++++++           | 78% ~05s          
  |++++++++++++++++++++++++++++++++++++++++          | 79% ~04s          
  |++++++++++++++++++++++++++++++++++++++++          | 80% ~04s          
  |+++++++++++++++++++++++++++++++++++++++++         | 81% ~04s          
  |++++++++++++++++++++++++++++++++++++++++++        | 82% ~04s          
  |++++++++++++++++++++++++++++++++++++++++++        | 83% ~04s          
  |+++++++++++++++++++++++++++++++++++++++++++       | 84% ~03s          
  |+++++++++++++++++++++++++++++++++++++++++++       | 86% ~03s          
  |++++++++++++++++++++++++++++++++++++++++++++      | 87% ~03s          
  |++++++++++++++++++++++++++++++++++++++++++++      | 88% ~03s          
  |+++++++++++++++++++++++++++++++++++++++++++++     | 89% ~02s          
  |+++++++++++++++++++++++++++++++++++++++++++++     | 90% ~02s          
  |++++++++++++++++++++++++++++++++++++++++++++++    | 91% ~02s          
  |+++++++++++++++++++++++++++++++++++++++++++++++   | 92% ~02s          
  |+++++++++++++++++++++++++++++++++++++++++++++++   | 93% ~01s          
  |++++++++++++++++++++++++++++++++++++++++++++++++  | 94% ~01s          
  |++++++++++++++++++++++++++++++++++++++++++++++++  | 96% ~01s          
  |+++++++++++++++++++++++++++++++++++++++++++++++++ | 97% ~01s          
  |+++++++++++++++++++++++++++++++++++++++++++++++++ | 98% ~00s          
  |++++++++++++++++++++++++++++++++++++++++++++++++++| 99% ~00s          
  |++++++++++++++++++++++++++++++++++++++++++++++++++| 100% elapsed=21s
```


```
Calculating cluster 4
```


```
  |                                                  | 0 % ~calculating  
  |+                                                 | 1 % ~13s          
  |++                                                | 2 % ~13s          
  |++                                                | 3 % ~13s          
  |+++                                               | 4 % ~12s          
  |+++                                               | 5 % ~12s          
  |++++                                              | 6 % ~12s          
  |++++                                              | 7 % ~12s          
  |+++++                                             | 8 % ~12s          
  |+++++                                             | 9 % ~12s          
  |++++++                                            | 10% ~11s          
  |++++++                                            | 11% ~11s          
  |+++++++                                           | 12% ~11s          
  |+++++++                                           | 14% ~11s          
  |++++++++                                          | 15% ~11s          
  |++++++++                                          | 16% ~11s          
  |+++++++++                                         | 17% ~11s          
  |+++++++++                                         | 18% ~11s          
  |++++++++++                                        | 19% ~11s          
  |++++++++++                                        | 20% ~10s          
  |+++++++++++                                       | 21% ~10s          
  |+++++++++++                                       | 22% ~10s          
  |++++++++++++                                      | 23% ~10s          
  |++++++++++++                                      | 24% ~10s          
  |+++++++++++++                                     | 25% ~10s          
  |++++++++++++++                                    | 26% ~10s          
  |++++++++++++++                                    | 27% ~10s          
  |+++++++++++++++                                   | 28% ~09s          
  |+++++++++++++++                                   | 29% ~09s          
  |++++++++++++++++                                  | 30% ~09s          
  |++++++++++++++++                                  | 31% ~09s          
  |+++++++++++++++++                                 | 32% ~09s          
  |+++++++++++++++++                                 | 33% ~09s          
  |++++++++++++++++++                                | 34% ~09s          
  |++++++++++++++++++                                | 35% ~08s          
  |+++++++++++++++++++                               | 36% ~08s          
  |+++++++++++++++++++                               | 38% ~08s          
  |++++++++++++++++++++                              | 39% ~08s          
  |++++++++++++++++++++                              | 40% ~08s          
  |+++++++++++++++++++++                             | 41% ~08s          
  |+++++++++++++++++++++                             | 42% ~08s          
  |++++++++++++++++++++++                            | 43% ~08s          
  |++++++++++++++++++++++                            | 44% ~07s          
  |+++++++++++++++++++++++                           | 45% ~07s          
  |+++++++++++++++++++++++                           | 46% ~07s          
  |++++++++++++++++++++++++                          | 47% ~07s          
  |++++++++++++++++++++++++                          | 48% ~07s          
  |+++++++++++++++++++++++++                         | 49% ~07s          
  |+++++++++++++++++++++++++                         | 50% ~07s          
  |++++++++++++++++++++++++++                        | 51% ~06s          
  |+++++++++++++++++++++++++++                       | 52% ~06s          
  |+++++++++++++++++++++++++++                       | 53% ~06s          
  |++++++++++++++++++++++++++++                      | 54% ~06s          
  |++++++++++++++++++++++++++++                      | 55% ~06s          
  |+++++++++++++++++++++++++++++                     | 56% ~06s          
  |+++++++++++++++++++++++++++++                     | 57% ~06s          
  |++++++++++++++++++++++++++++++                    | 58% ~06s          
  |++++++++++++++++++++++++++++++                    | 59% ~05s          
  |+++++++++++++++++++++++++++++++                   | 60% ~05s          
  |+++++++++++++++++++++++++++++++                   | 61% ~05s          
  |++++++++++++++++++++++++++++++++                  | 62% ~05s          
  |++++++++++++++++++++++++++++++++                  | 64% ~05s          
  |+++++++++++++++++++++++++++++++++                 | 65% ~05s          
  |+++++++++++++++++++++++++++++++++                 | 66% ~05s          
  |++++++++++++++++++++++++++++++++++                | 67% ~04s          
  |++++++++++++++++++++++++++++++++++                | 68% ~04s          
  |+++++++++++++++++++++++++++++++++++               | 69% ~04s          
  |+++++++++++++++++++++++++++++++++++               | 70% ~04s          
  |++++++++++++++++++++++++++++++++++++              | 71% ~04s          
  |++++++++++++++++++++++++++++++++++++              | 72% ~04s          
  |+++++++++++++++++++++++++++++++++++++             | 73% ~04s          
  |+++++++++++++++++++++++++++++++++++++             | 74% ~03s          
  |++++++++++++++++++++++++++++++++++++++            | 75% ~03s          
  |+++++++++++++++++++++++++++++++++++++++           | 76% ~03s          
  |+++++++++++++++++++++++++++++++++++++++           | 77% ~03s          
  |++++++++++++++++++++++++++++++++++++++++          | 78% ~03s          
  |++++++++++++++++++++++++++++++++++++++++          | 79% ~03s          
  |+++++++++++++++++++++++++++++++++++++++++         | 80% ~03s          
  |+++++++++++++++++++++++++++++++++++++++++         | 81% ~03s          
  |++++++++++++++++++++++++++++++++++++++++++        | 82% ~02s          
  |++++++++++++++++++++++++++++++++++++++++++        | 83% ~02s          
  |+++++++++++++++++++++++++++++++++++++++++++       | 84% ~02s          
  |+++++++++++++++++++++++++++++++++++++++++++       | 85% ~02s          
  |++++++++++++++++++++++++++++++++++++++++++++      | 86% ~02s          
  |++++++++++++++++++++++++++++++++++++++++++++      | 88% ~02s          
  |+++++++++++++++++++++++++++++++++++++++++++++     | 89% ~02s          
  |+++++++++++++++++++++++++++++++++++++++++++++     | 90% ~01s          
  |++++++++++++++++++++++++++++++++++++++++++++++    | 91% ~01s          
  |++++++++++++++++++++++++++++++++++++++++++++++    | 92% ~01s          
  |+++++++++++++++++++++++++++++++++++++++++++++++   | 93% ~01s          
  |+++++++++++++++++++++++++++++++++++++++++++++++   | 94% ~01s          
  |++++++++++++++++++++++++++++++++++++++++++++++++  | 95% ~01s          
  |++++++++++++++++++++++++++++++++++++++++++++++++  | 96% ~01s          
  |+++++++++++++++++++++++++++++++++++++++++++++++++ | 97% ~00s          
  |+++++++++++++++++++++++++++++++++++++++++++++++++ | 98% ~00s          
  |++++++++++++++++++++++++++++++++++++++++++++++++++| 99% ~00s          
  |++++++++++++++++++++++++++++++++++++++++++++++++++| 100% elapsed=13s
```


```
Calculating cluster 5
```


```
  |                                                  | 0 % ~calculating  
  |+                                                 | 1 % ~09s          
  |++                                                | 2 % ~08s          
  |++                                                | 4 % ~08s          
  |+++                                               | 5 % ~08s          
  |++++                                              | 6 % ~08s          
  |++++                                              | 8 % ~07s          
  |+++++                                             | 9 % ~07s          
  |+++++                                             | 10% ~07s          
  |++++++                                            | 11% ~07s          
  |+++++++                                           | 12% ~07s          
  |+++++++                                           | 14% ~07s          
  |++++++++                                          | 15% ~07s          
  |+++++++++                                         | 16% ~07s          
  |+++++++++                                         | 18% ~07s          
  |++++++++++                                        | 19% ~07s          
  |++++++++++                                        | 20% ~07s          
  |+++++++++++                                       | 21% ~06s          
  |++++++++++++                                      | 22% ~06s          
  |++++++++++++                                      | 24% ~06s          
  |+++++++++++++                                     | 25% ~06s          
  |++++++++++++++                                    | 26% ~06s          
  |++++++++++++++                                    | 28% ~06s          
  |+++++++++++++++                                   | 29% ~06s          
  |+++++++++++++++                                   | 30% ~06s          
  |++++++++++++++++                                  | 31% ~06s          
  |+++++++++++++++++                                 | 32% ~06s          
  |+++++++++++++++++                                 | 34% ~06s          
  |++++++++++++++++++                                | 35% ~05s          
  |+++++++++++++++++++                               | 36% ~05s          
  |+++++++++++++++++++                               | 38% ~05s          
  |++++++++++++++++++++                              | 39% ~05s          
  |++++++++++++++++++++                              | 40% ~05s          
  |+++++++++++++++++++++                             | 41% ~05s          
  |++++++++++++++++++++++                            | 42% ~05s          
  |++++++++++++++++++++++                            | 44% ~05s          
  |+++++++++++++++++++++++                           | 45% ~05s          
  |++++++++++++++++++++++++                          | 46% ~04s          
  |++++++++++++++++++++++++                          | 48% ~04s          
  |+++++++++++++++++++++++++                         | 49% ~04s          
  |+++++++++++++++++++++++++                         | 50% ~04s          
  |++++++++++++++++++++++++++                        | 51% ~04s          
  |+++++++++++++++++++++++++++                       | 52% ~04s          
  |+++++++++++++++++++++++++++                       | 54% ~04s          
  |++++++++++++++++++++++++++++                      | 55% ~04s          
  |+++++++++++++++++++++++++++++                     | 56% ~04s          
  |+++++++++++++++++++++++++++++                     | 58% ~04s          
  |++++++++++++++++++++++++++++++                    | 59% ~03s          
  |++++++++++++++++++++++++++++++                    | 60% ~03s          
  |+++++++++++++++++++++++++++++++                   | 61% ~03s          
  |++++++++++++++++++++++++++++++++                  | 62% ~03s          
  |++++++++++++++++++++++++++++++++                  | 64% ~03s          
  |+++++++++++++++++++++++++++++++++                 | 65% ~03s          
  |++++++++++++++++++++++++++++++++++                | 66% ~03s          
  |++++++++++++++++++++++++++++++++++                | 68% ~03s          
  |+++++++++++++++++++++++++++++++++++               | 69% ~03s          
  |+++++++++++++++++++++++++++++++++++               | 70% ~02s          
  |++++++++++++++++++++++++++++++++++++              | 71% ~02s          
  |+++++++++++++++++++++++++++++++++++++             | 72% ~02s          
  |+++++++++++++++++++++++++++++++++++++             | 74% ~02s          
  |++++++++++++++++++++++++++++++++++++++            | 75% ~02s          
  |+++++++++++++++++++++++++++++++++++++++           | 76% ~02s          
  |+++++++++++++++++++++++++++++++++++++++           | 78% ~02s          
  |++++++++++++++++++++++++++++++++++++++++          | 79% ~02s          
  |++++++++++++++++++++++++++++++++++++++++          | 80% ~02s          
  |+++++++++++++++++++++++++++++++++++++++++         | 81% ~02s          
  |++++++++++++++++++++++++++++++++++++++++++        | 82% ~01s          
  |++++++++++++++++++++++++++++++++++++++++++        | 84% ~01s          
  |+++++++++++++++++++++++++++++++++++++++++++       | 85% ~01s          
  |++++++++++++++++++++++++++++++++++++++++++++      | 86% ~01s          
  |++++++++++++++++++++++++++++++++++++++++++++      | 88% ~01s          
  |+++++++++++++++++++++++++++++++++++++++++++++     | 89% ~01s          
  |+++++++++++++++++++++++++++++++++++++++++++++     | 90% ~01s          
  |++++++++++++++++++++++++++++++++++++++++++++++    | 91% ~01s          
  |+++++++++++++++++++++++++++++++++++++++++++++++   | 92% ~01s          
  |+++++++++++++++++++++++++++++++++++++++++++++++   | 94% ~01s          
  |++++++++++++++++++++++++++++++++++++++++++++++++  | 95% ~00s          
  |+++++++++++++++++++++++++++++++++++++++++++++++++ | 96% ~00s          
  |+++++++++++++++++++++++++++++++++++++++++++++++++ | 98% ~00s          
  |++++++++++++++++++++++++++++++++++++++++++++++++++| 99% ~00s          
  |++++++++++++++++++++++++++++++++++++++++++++++++++| 100% elapsed=08s
```


```
Calculating cluster 6
```


```
  |                                                  | 0 % ~calculating  
  |+                                                 | 1 % ~41s          
  |+                                                 | 2 % ~41s          
  |++                                                | 3 % ~42s          
  |++                                                | 4 % ~41s          
  |+++                                               | 5 % ~41s          
  |+++                                               | 6 % ~41s          
  |++++                                              | 7 % ~40s          
  |++++                                              | 8 % ~40s          
  |+++++                                             | 9 % ~39s          
  |+++++                                             | 10% ~39s          
  |++++++                                            | 11% ~38s          
  |++++++                                            | 12% ~38s          
  |+++++++                                           | 13% ~37s          
  |+++++++                                           | 14% ~37s          
  |++++++++                                          | 15% ~36s          
  |++++++++                                          | 16% ~36s          
  |+++++++++                                         | 17% ~35s          
  |+++++++++                                         | 18% ~35s          
  |++++++++++                                        | 19% ~34s          
  |++++++++++                                        | 20% ~34s          
  |+++++++++++                                       | 21% ~34s          
  |+++++++++++                                       | 22% ~33s          
  |++++++++++++                                      | 23% ~34s          
  |++++++++++++                                      | 24% ~33s          
  |+++++++++++++                                     | 25% ~33s          
  |+++++++++++++                                     | 26% ~32s          
  |++++++++++++++                                    | 27% ~32s          
  |++++++++++++++                                    | 28% ~31s          
  |+++++++++++++++                                   | 29% ~31s          
  |+++++++++++++++                                   | 30% ~30s          
  |++++++++++++++++                                  | 31% ~30s          
  |++++++++++++++++                                  | 32% ~29s          
  |+++++++++++++++++                                 | 33% ~29s          
  |+++++++++++++++++                                 | 34% ~29s          
  |++++++++++++++++++                                | 35% ~28s          
  |++++++++++++++++++                                | 36% ~28s          
  |+++++++++++++++++++                               | 37% ~27s          
  |+++++++++++++++++++                               | 38% ~27s          
  |++++++++++++++++++++                              | 39% ~26s          
  |++++++++++++++++++++                              | 40% ~26s          
  |+++++++++++++++++++++                             | 41% ~25s          
  |+++++++++++++++++++++                             | 42% ~25s          
  |++++++++++++++++++++++                            | 43% ~25s          
  |++++++++++++++++++++++                            | 44% ~24s          
  |+++++++++++++++++++++++                           | 45% ~24s          
  |+++++++++++++++++++++++                           | 46% ~23s          
  |++++++++++++++++++++++++                          | 47% ~23s          
  |++++++++++++++++++++++++                          | 48% ~22s          
  |+++++++++++++++++++++++++                         | 49% ~22s          
  |+++++++++++++++++++++++++                         | 50% ~22s          
  |++++++++++++++++++++++++++                        | 51% ~21s          
  |++++++++++++++++++++++++++                        | 52% ~21s          
  |+++++++++++++++++++++++++++                       | 53% ~20s          
  |+++++++++++++++++++++++++++                       | 54% ~20s          
  |++++++++++++++++++++++++++++                      | 55% ~19s          
  |++++++++++++++++++++++++++++                      | 56% ~19s          
  |+++++++++++++++++++++++++++++                     | 57% ~18s          
  |+++++++++++++++++++++++++++++                     | 58% ~18s          
  |++++++++++++++++++++++++++++++                    | 59% ~18s          
  |++++++++++++++++++++++++++++++                    | 60% ~17s          
  |+++++++++++++++++++++++++++++++                   | 61% ~17s          
  |+++++++++++++++++++++++++++++++                   | 62% ~16s          
  |++++++++++++++++++++++++++++++++                  | 63% ~16s          
  |++++++++++++++++++++++++++++++++                  | 64% ~15s          
  |+++++++++++++++++++++++++++++++++                 | 65% ~15s          
  |+++++++++++++++++++++++++++++++++                 | 66% ~15s          
  |++++++++++++++++++++++++++++++++++                | 67% ~14s          
  |++++++++++++++++++++++++++++++++++                | 68% ~14s          
  |+++++++++++++++++++++++++++++++++++               | 69% ~13s          
  |+++++++++++++++++++++++++++++++++++               | 70% ~13s          
  |++++++++++++++++++++++++++++++++++++              | 71% ~12s          
  |++++++++++++++++++++++++++++++++++++              | 72% ~12s          
  |+++++++++++++++++++++++++++++++++++++             | 73% ~12s          
  |+++++++++++++++++++++++++++++++++++++             | 74% ~11s          
  |++++++++++++++++++++++++++++++++++++++            | 75% ~11s          
  |++++++++++++++++++++++++++++++++++++++            | 76% ~10s          
  |+++++++++++++++++++++++++++++++++++++++           | 77% ~10s          
  |+++++++++++++++++++++++++++++++++++++++           | 78% ~09s          
  |++++++++++++++++++++++++++++++++++++++++          | 79% ~09s          
  |++++++++++++++++++++++++++++++++++++++++          | 80% ~09s          
  |+++++++++++++++++++++++++++++++++++++++++         | 81% ~08s          
  |+++++++++++++++++++++++++++++++++++++++++         | 82% ~08s          
  |++++++++++++++++++++++++++++++++++++++++++        | 83% ~07s          
  |++++++++++++++++++++++++++++++++++++++++++        | 84% ~07s          
  |+++++++++++++++++++++++++++++++++++++++++++       | 85% ~06s          
  |+++++++++++++++++++++++++++++++++++++++++++       | 86% ~06s          
  |++++++++++++++++++++++++++++++++++++++++++++      | 87% ~06s          
  |++++++++++++++++++++++++++++++++++++++++++++      | 88% ~05s          
  |+++++++++++++++++++++++++++++++++++++++++++++     | 89% ~05s          
  |+++++++++++++++++++++++++++++++++++++++++++++     | 90% ~04s          
  |++++++++++++++++++++++++++++++++++++++++++++++    | 91% ~04s          
  |++++++++++++++++++++++++++++++++++++++++++++++    | 92% ~03s          
  |+++++++++++++++++++++++++++++++++++++++++++++++   | 93% ~03s          
  |+++++++++++++++++++++++++++++++++++++++++++++++   | 94% ~03s          
  |++++++++++++++++++++++++++++++++++++++++++++++++  | 95% ~02s          
  |++++++++++++++++++++++++++++++++++++++++++++++++  | 96% ~02s          
  |+++++++++++++++++++++++++++++++++++++++++++++++++ | 97% ~01s          
  |+++++++++++++++++++++++++++++++++++++++++++++++++ | 98% ~01s          
  |++++++++++++++++++++++++++++++++++++++++++++++++++| 99% ~00s          
  |++++++++++++++++++++++++++++++++++++++++++++++++++| 100% elapsed=43s
```


```
Calculating cluster 7
```


```
  |                                                  | 0 % ~calculating  
  |+                                                 | 1 % ~20s          
  |++                                                | 2 % ~21s          
  |++                                                | 3 % ~21s          
  |+++                                               | 4 % ~20s          
  |+++                                               | 5 % ~19s          
  |++++                                              | 6 % ~19s          
  |++++                                              | 7 % ~19s          
  |+++++                                             | 8 % ~18s          
  |+++++                                             | 9 % ~18s          
  |++++++                                            | 11% ~19s          
  |++++++                                            | 12% ~18s          
  |+++++++                                           | 13% ~18s          
  |+++++++                                           | 14% ~18s          
  |++++++++                                          | 15% ~17s          
  |++++++++                                          | 16% ~17s          
  |+++++++++                                         | 17% ~17s          
  |+++++++++                                         | 18% ~17s          
  |++++++++++                                        | 19% ~17s          
  |++++++++++                                        | 20% ~16s          
  |+++++++++++                                       | 21% ~16s          
  |++++++++++++                                      | 22% ~16s          
  |++++++++++++                                      | 23% ~16s          
  |+++++++++++++                                     | 24% ~15s          
  |+++++++++++++                                     | 25% ~15s          
  |++++++++++++++                                    | 26% ~15s          
  |++++++++++++++                                    | 27% ~15s          
  |+++++++++++++++                                   | 28% ~14s          
  |+++++++++++++++                                   | 29% ~14s          
  |++++++++++++++++                                  | 31% ~14s          
  |++++++++++++++++                                  | 32% ~14s          
  |+++++++++++++++++                                 | 33% ~13s          
  |+++++++++++++++++                                 | 34% ~13s          
  |++++++++++++++++++                                | 35% ~13s          
  |++++++++++++++++++                                | 36% ~13s          
  |+++++++++++++++++++                               | 37% ~13s          
  |+++++++++++++++++++                               | 38% ~12s          
  |++++++++++++++++++++                              | 39% ~12s          
  |++++++++++++++++++++                              | 40% ~12s          
  |+++++++++++++++++++++                             | 41% ~12s          
  |++++++++++++++++++++++                            | 42% ~12s          
  |++++++++++++++++++++++                            | 43% ~11s          
  |+++++++++++++++++++++++                           | 44% ~11s          
  |+++++++++++++++++++++++                           | 45% ~11s          
  |++++++++++++++++++++++++                          | 46% ~11s          
  |++++++++++++++++++++++++                          | 47% ~11s          
  |+++++++++++++++++++++++++                         | 48% ~10s          
  |+++++++++++++++++++++++++                         | 49% ~10s          
  |++++++++++++++++++++++++++                        | 51% ~10s          
  |++++++++++++++++++++++++++                        | 52% ~10s          
  |+++++++++++++++++++++++++++                       | 53% ~09s          
  |+++++++++++++++++++++++++++                       | 54% ~09s          
  |++++++++++++++++++++++++++++                      | 55% ~09s          
  |++++++++++++++++++++++++++++                      | 56% ~09s          
  |+++++++++++++++++++++++++++++                     | 57% ~09s          
  |+++++++++++++++++++++++++++++                     | 58% ~08s          
  |++++++++++++++++++++++++++++++                    | 59% ~08s          
  |++++++++++++++++++++++++++++++                    | 60% ~08s          
  |+++++++++++++++++++++++++++++++                   | 61% ~08s          
  |++++++++++++++++++++++++++++++++                  | 62% ~08s          
  |++++++++++++++++++++++++++++++++                  | 63% ~08s          
  |+++++++++++++++++++++++++++++++++                 | 64% ~07s          
  |+++++++++++++++++++++++++++++++++                 | 65% ~07s          
  |++++++++++++++++++++++++++++++++++                | 66% ~07s          
  |++++++++++++++++++++++++++++++++++                | 67% ~07s          
  |+++++++++++++++++++++++++++++++++++               | 68% ~07s          
  |+++++++++++++++++++++++++++++++++++               | 69% ~06s          
  |++++++++++++++++++++++++++++++++++++              | 71% ~06s          
  |++++++++++++++++++++++++++++++++++++              | 72% ~06s          
  |+++++++++++++++++++++++++++++++++++++             | 73% ~06s          
  |+++++++++++++++++++++++++++++++++++++             | 74% ~06s          
  |++++++++++++++++++++++++++++++++++++++            | 75% ~05s          
  |++++++++++++++++++++++++++++++++++++++            | 76% ~05s          
  |+++++++++++++++++++++++++++++++++++++++           | 77% ~05s          
  |+++++++++++++++++++++++++++++++++++++++           | 78% ~05s          
  |++++++++++++++++++++++++++++++++++++++++          | 79% ~04s          
  |++++++++++++++++++++++++++++++++++++++++          | 80% ~04s          
  |+++++++++++++++++++++++++++++++++++++++++         | 81% ~04s          
  |++++++++++++++++++++++++++++++++++++++++++        | 82% ~04s          
  |++++++++++++++++++++++++++++++++++++++++++        | 83% ~04s          
  |+++++++++++++++++++++++++++++++++++++++++++       | 84% ~03s          
  |+++++++++++++++++++++++++++++++++++++++++++       | 85% ~03s          
  |++++++++++++++++++++++++++++++++++++++++++++      | 86% ~03s          
  |++++++++++++++++++++++++++++++++++++++++++++      | 87% ~03s          
  |+++++++++++++++++++++++++++++++++++++++++++++     | 88% ~02s          
  |+++++++++++++++++++++++++++++++++++++++++++++     | 89% ~02s          
  |++++++++++++++++++++++++++++++++++++++++++++++    | 91% ~02s          
  |++++++++++++++++++++++++++++++++++++++++++++++    | 92% ~02s          
  |+++++++++++++++++++++++++++++++++++++++++++++++   | 93% ~02s          
  |+++++++++++++++++++++++++++++++++++++++++++++++   | 94% ~01s          
  |++++++++++++++++++++++++++++++++++++++++++++++++  | 95% ~01s          
  |++++++++++++++++++++++++++++++++++++++++++++++++  | 96% ~01s          
  |+++++++++++++++++++++++++++++++++++++++++++++++++ | 97% ~01s          
  |+++++++++++++++++++++++++++++++++++++++++++++++++ | 98% ~00s          
  |++++++++++++++++++++++++++++++++++++++++++++++++++| 99% ~00s          
  |++++++++++++++++++++++++++++++++++++++++++++++++++| 100% elapsed=22s
```


```
Calculating cluster 8
```


```
  |                                                  | 0 % ~calculating  
  |+                                                 | 1 % ~05s          
  |++                                                | 2 % ~05s          
  |++                                                | 3 % ~05s          
  |+++                                               | 4 % ~05s          
  |+++                                               | 5 % ~05s          
  |++++                                              | 6 % ~05s          
  |++++                                              | 7 % ~05s          
  |+++++                                             | 8 % ~05s          
  |+++++                                             | 9 % ~04s          
  |++++++                                            | 10% ~04s          
  |++++++                                            | 11% ~05s          
  |+++++++                                           | 12% ~04s          
  |+++++++                                           | 13% ~04s          
  |++++++++                                          | 14% ~04s          
  |++++++++                                          | 15% ~04s          
  |+++++++++                                         | 16% ~04s          
  |+++++++++                                         | 17% ~04s          
  |++++++++++                                        | 18% ~04s          
  |++++++++++                                        | 19% ~04s          
  |+++++++++++                                       | 20% ~04s          
  |+++++++++++                                       | 21% ~04s          
  |++++++++++++                                      | 22% ~04s          
  |++++++++++++                                      | 23% ~04s          
  |+++++++++++++                                     | 24% ~04s          
  |+++++++++++++                                     | 25% ~04s          
  |++++++++++++++                                    | 26% ~04s          
  |++++++++++++++                                    | 27% ~04s          
  |+++++++++++++++                                   | 28% ~04s          
  |+++++++++++++++                                   | 29% ~04s          
  |++++++++++++++++                                  | 30% ~04s          
  |++++++++++++++++                                  | 31% ~03s          
  |+++++++++++++++++                                 | 32% ~03s          
  |+++++++++++++++++                                 | 33% ~03s          
  |++++++++++++++++++                                | 34% ~03s          
  |++++++++++++++++++                                | 35% ~03s          
  |+++++++++++++++++++                               | 36% ~03s          
  |+++++++++++++++++++                               | 37% ~03s          
  |++++++++++++++++++++                              | 38% ~03s          
  |++++++++++++++++++++                              | 39% ~03s          
  |+++++++++++++++++++++                             | 40% ~03s          
  |+++++++++++++++++++++                             | 41% ~03s          
  |++++++++++++++++++++++                            | 42% ~03s          
  |++++++++++++++++++++++                            | 43% ~03s          
  |+++++++++++++++++++++++                           | 44% ~03s          
  |+++++++++++++++++++++++                           | 45% ~03s          
  |++++++++++++++++++++++++                          | 46% ~03s          
  |++++++++++++++++++++++++                          | 47% ~03s          
  |+++++++++++++++++++++++++                         | 48% ~03s          
  |+++++++++++++++++++++++++                         | 49% ~03s          
  |++++++++++++++++++++++++++                        | 51% ~03s          
  |++++++++++++++++++++++++++                        | 52% ~03s          
  |+++++++++++++++++++++++++++                       | 53% ~03s          
  |+++++++++++++++++++++++++++                       | 54% ~03s          
  |++++++++++++++++++++++++++++                      | 55% ~03s          
  |++++++++++++++++++++++++++++                      | 56% ~03s          
  |+++++++++++++++++++++++++++++                     | 57% ~03s          
  |+++++++++++++++++++++++++++++                     | 58% ~03s          
  |++++++++++++++++++++++++++++++                    | 59% ~03s          
  |++++++++++++++++++++++++++++++                    | 60% ~02s          
  |+++++++++++++++++++++++++++++++                   | 61% ~02s          
  |+++++++++++++++++++++++++++++++                   | 62% ~02s          
  |++++++++++++++++++++++++++++++++                  | 63% ~02s          
  |++++++++++++++++++++++++++++++++                  | 64% ~02s          
  |+++++++++++++++++++++++++++++++++                 | 65% ~02s          
  |+++++++++++++++++++++++++++++++++                 | 66% ~02s          
  |++++++++++++++++++++++++++++++++++                | 67% ~02s          
  |++++++++++++++++++++++++++++++++++                | 68% ~02s          
  |+++++++++++++++++++++++++++++++++++               | 69% ~02s          
  |+++++++++++++++++++++++++++++++++++               | 70% ~02s          
  |++++++++++++++++++++++++++++++++++++              | 71% ~02s          
  |++++++++++++++++++++++++++++++++++++              | 72% ~02s          
  |+++++++++++++++++++++++++++++++++++++             | 73% ~02s          
  |+++++++++++++++++++++++++++++++++++++             | 74% ~02s          
  |++++++++++++++++++++++++++++++++++++++            | 75% ~01s          
  |++++++++++++++++++++++++++++++++++++++            | 76% ~01s          
  |+++++++++++++++++++++++++++++++++++++++           | 77% ~01s          
  |+++++++++++++++++++++++++++++++++++++++           | 78% ~01s          
  |++++++++++++++++++++++++++++++++++++++++          | 79% ~01s          
  |++++++++++++++++++++++++++++++++++++++++          | 80% ~01s          
  |+++++++++++++++++++++++++++++++++++++++++         | 81% ~01s          
  |+++++++++++++++++++++++++++++++++++++++++         | 82% ~01s          
  |++++++++++++++++++++++++++++++++++++++++++        | 83% ~01s          
  |++++++++++++++++++++++++++++++++++++++++++        | 84% ~01s          
  |+++++++++++++++++++++++++++++++++++++++++++       | 85% ~01s          
  |+++++++++++++++++++++++++++++++++++++++++++       | 86% ~01s          
  |++++++++++++++++++++++++++++++++++++++++++++      | 87% ~01s          
  |++++++++++++++++++++++++++++++++++++++++++++      | 88% ~01s          
  |+++++++++++++++++++++++++++++++++++++++++++++     | 89% ~01s          
  |+++++++++++++++++++++++++++++++++++++++++++++     | 90% ~01s          
  |++++++++++++++++++++++++++++++++++++++++++++++    | 91% ~01s          
  |++++++++++++++++++++++++++++++++++++++++++++++    | 92% ~00s          
  |+++++++++++++++++++++++++++++++++++++++++++++++   | 93% ~00s          
  |+++++++++++++++++++++++++++++++++++++++++++++++   | 94% ~00s          
  |++++++++++++++++++++++++++++++++++++++++++++++++  | 95% ~00s          
  |++++++++++++++++++++++++++++++++++++++++++++++++  | 96% ~00s          
  |+++++++++++++++++++++++++++++++++++++++++++++++++ | 97% ~00s          
  |+++++++++++++++++++++++++++++++++++++++++++++++++ | 98% ~00s          
  |++++++++++++++++++++++++++++++++++++++++++++++++++| 99% ~00s          
  |++++++++++++++++++++++++++++++++++++++++++++++++++| 100% elapsed=06s
```


```
Calculating cluster 9
```


```
  |                                                  | 0 % ~calculating  
  |+                                                 | 1 % ~01m 03s      
  |++                                                | 2 % ~01m 03s      
  |++                                                | 3 % ~01m 02s      
  |+++                                               | 4 % ~01m 01s      
  |+++                                               | 5 % ~59s          
  |++++                                              | 6 % ~58s          
  |++++                                              | 7 % ~57s          
  |+++++                                             | 8 % ~56s          
  |+++++                                             | 9 % ~55s          
  |++++++                                            | 10% ~54s          
  |++++++                                            | 11% ~53s          
  |+++++++                                           | 12% ~53s          
  |+++++++                                           | 14% ~52s          
  |++++++++                                          | 15% ~51s          
  |++++++++                                          | 16% ~51s          
  |+++++++++                                         | 17% ~50s          
  |+++++++++                                         | 18% ~49s          
  |++++++++++                                        | 19% ~48s          
  |++++++++++                                        | 20% ~47s          
  |+++++++++++                                       | 21% ~47s          
  |+++++++++++                                       | 22% ~46s          
  |++++++++++++                                      | 23% ~45s          
  |++++++++++++                                      | 24% ~45s          
  |+++++++++++++                                     | 25% ~44s          
  |++++++++++++++                                    | 26% ~43s          
  |++++++++++++++                                    | 27% ~43s          
  |+++++++++++++++                                   | 28% ~42s          
  |+++++++++++++++                                   | 29% ~41s          
  |++++++++++++++++                                  | 30% ~41s          
  |++++++++++++++++                                  | 31% ~40s          
  |+++++++++++++++++                                 | 32% ~39s          
  |+++++++++++++++++                                 | 33% ~39s          
  |++++++++++++++++++                                | 34% ~38s          
  |++++++++++++++++++                                | 35% ~38s          
  |+++++++++++++++++++                               | 36% ~37s          
  |+++++++++++++++++++                               | 38% ~36s          
  |++++++++++++++++++++                              | 39% ~36s          
  |++++++++++++++++++++                              | 40% ~35s          
  |+++++++++++++++++++++                             | 41% ~35s          
  |+++++++++++++++++++++                             | 42% ~34s          
  |++++++++++++++++++++++                            | 43% ~34s          
  |++++++++++++++++++++++                            | 44% ~33s          
  |+++++++++++++++++++++++                           | 45% ~32s          
  |+++++++++++++++++++++++                           | 46% ~32s          
  |++++++++++++++++++++++++                          | 47% ~31s          
  |++++++++++++++++++++++++                          | 48% ~30s          
  |+++++++++++++++++++++++++                         | 49% ~30s          
  |+++++++++++++++++++++++++                         | 50% ~29s          
  |++++++++++++++++++++++++++                        | 51% ~29s          
  |+++++++++++++++++++++++++++                       | 52% ~28s          
  |+++++++++++++++++++++++++++                       | 53% ~27s          
  |++++++++++++++++++++++++++++                      | 54% ~27s          
  |++++++++++++++++++++++++++++                      | 55% ~26s          
  |+++++++++++++++++++++++++++++                     | 56% ~25s          
  |+++++++++++++++++++++++++++++                     | 57% ~25s          
  |++++++++++++++++++++++++++++++                    | 58% ~24s          
  |++++++++++++++++++++++++++++++                    | 59% ~24s          
  |+++++++++++++++++++++++++++++++                   | 60% ~23s          
  |+++++++++++++++++++++++++++++++                   | 61% ~22s          
  |++++++++++++++++++++++++++++++++                  | 62% ~22s          
  |++++++++++++++++++++++++++++++++                  | 64% ~21s          
  |+++++++++++++++++++++++++++++++++                 | 65% ~21s          
  |+++++++++++++++++++++++++++++++++                 | 66% ~20s          
  |++++++++++++++++++++++++++++++++++                | 67% ~19s          
  |++++++++++++++++++++++++++++++++++                | 68% ~19s          
  |+++++++++++++++++++++++++++++++++++               | 69% ~18s          
  |+++++++++++++++++++++++++++++++++++               | 70% ~18s          
  |++++++++++++++++++++++++++++++++++++              | 71% ~17s          
  |++++++++++++++++++++++++++++++++++++              | 72% ~16s          
  |+++++++++++++++++++++++++++++++++++++             | 73% ~16s          
  |+++++++++++++++++++++++++++++++++++++             | 74% ~15s          
  |++++++++++++++++++++++++++++++++++++++            | 75% ~14s          
  |+++++++++++++++++++++++++++++++++++++++           | 76% ~14s          
  |+++++++++++++++++++++++++++++++++++++++           | 77% ~13s          
  |++++++++++++++++++++++++++++++++++++++++          | 78% ~13s          
  |++++++++++++++++++++++++++++++++++++++++          | 79% ~12s          
  |+++++++++++++++++++++++++++++++++++++++++         | 80% ~11s          
  |+++++++++++++++++++++++++++++++++++++++++         | 81% ~11s          
  |++++++++++++++++++++++++++++++++++++++++++        | 82% ~10s          
  |++++++++++++++++++++++++++++++++++++++++++        | 83% ~10s          
  |+++++++++++++++++++++++++++++++++++++++++++       | 84% ~09s          
  |+++++++++++++++++++++++++++++++++++++++++++       | 85% ~08s          
  |++++++++++++++++++++++++++++++++++++++++++++      | 86% ~08s          
  |++++++++++++++++++++++++++++++++++++++++++++      | 88% ~07s          
  |+++++++++++++++++++++++++++++++++++++++++++++     | 89% ~07s          
  |+++++++++++++++++++++++++++++++++++++++++++++     | 90% ~06s          
  |++++++++++++++++++++++++++++++++++++++++++++++    | 91% ~05s          
  |++++++++++++++++++++++++++++++++++++++++++++++    | 92% ~05s          
  |+++++++++++++++++++++++++++++++++++++++++++++++   | 93% ~04s          
  |+++++++++++++++++++++++++++++++++++++++++++++++   | 94% ~04s          
  |++++++++++++++++++++++++++++++++++++++++++++++++  | 95% ~03s          
  |++++++++++++++++++++++++++++++++++++++++++++++++  | 96% ~02s          
  |+++++++++++++++++++++++++++++++++++++++++++++++++ | 97% ~02s          
  |+++++++++++++++++++++++++++++++++++++++++++++++++ | 98% ~01s          
  |++++++++++++++++++++++++++++++++++++++++++++++++++| 99% ~01s          
  |++++++++++++++++++++++++++++++++++++++++++++++++++| 100% elapsed=58s
```


```
Calculating cluster 10
```


```
  |                                                  | 0 % ~calculating  
  |+                                                 | 1 % ~01m 33s      
  |++                                                | 2 % ~01m 34s      
  |++                                                | 3 % ~01m 35s      
  |+++                                               | 4 % ~01m 33s      
  |+++                                               | 5 % ~01m 32s      
  |++++                                              | 6 % ~01m 30s      
  |++++                                              | 7 % ~01m 29s      
  |+++++                                             | 8 % ~01m 28s      
  |+++++                                             | 9 % ~01m 27s      
  |++++++                                            | 10% ~01m 26s      
  |++++++                                            | 11% ~01m 25s      
  |+++++++                                           | 12% ~01m 24s      
  |+++++++                                           | 13% ~01m 23s      
  |++++++++                                          | 14% ~01m 22s      
  |++++++++                                          | 15% ~01m 21s      
  |+++++++++                                         | 16% ~01m 20s      
  |+++++++++                                         | 18% ~01m 19s      
  |++++++++++                                        | 19% ~01m 18s      
  |++++++++++                                        | 20% ~01m 17s      
  |+++++++++++                                       | 21% ~01m 16s      
  |+++++++++++                                       | 22% ~01m 16s      
  |++++++++++++                                      | 23% ~01m 15s      
  |++++++++++++                                      | 24% ~01m 14s      
  |+++++++++++++                                     | 25% ~01m 13s      
  |+++++++++++++                                     | 26% ~01m 12s      
  |++++++++++++++                                    | 27% ~01m 11s      
  |++++++++++++++                                    | 28% ~01m 10s      
  |+++++++++++++++                                   | 29% ~01m 09s      
  |+++++++++++++++                                   | 30% ~01m 08s      
  |++++++++++++++++                                  | 31% ~01m 07s      
  |++++++++++++++++                                  | 32% ~01m 06s      
  |+++++++++++++++++                                 | 33% ~01m 05s      
  |++++++++++++++++++                                | 34% ~01m 04s      
  |++++++++++++++++++                                | 35% ~01m 03s      
  |+++++++++++++++++++                               | 36% ~01m 02s      
  |+++++++++++++++++++                               | 37% ~01m 01s      
  |++++++++++++++++++++                              | 38% ~01m 00s      
  |++++++++++++++++++++                              | 39% ~59s          
  |+++++++++++++++++++++                             | 40% ~58s          
  |+++++++++++++++++++++                             | 41% ~57s          
  |++++++++++++++++++++++                            | 42% ~56s          
  |++++++++++++++++++++++                            | 43% ~55s          
  |+++++++++++++++++++++++                           | 44% ~54s          
  |+++++++++++++++++++++++                           | 45% ~53s          
  |++++++++++++++++++++++++                          | 46% ~52s          
  |++++++++++++++++++++++++                          | 47% ~51s          
  |+++++++++++++++++++++++++                         | 48% ~50s          
  |+++++++++++++++++++++++++                         | 49% ~49s          
  |++++++++++++++++++++++++++                        | 51% ~48s          
  |++++++++++++++++++++++++++                        | 52% ~47s          
  |+++++++++++++++++++++++++++                       | 53% ~46s          
  |+++++++++++++++++++++++++++                       | 54% ~45s          
  |++++++++++++++++++++++++++++                      | 55% ~44s          
  |++++++++++++++++++++++++++++                      | 56% ~43s          
  |+++++++++++++++++++++++++++++                     | 57% ~42s          
  |+++++++++++++++++++++++++++++                     | 58% ~41s          
  |++++++++++++++++++++++++++++++                    | 59% ~40s          
  |++++++++++++++++++++++++++++++                    | 60% ~39s          
  |+++++++++++++++++++++++++++++++                   | 61% ~38s          
  |+++++++++++++++++++++++++++++++                   | 62% ~37s          
  |++++++++++++++++++++++++++++++++                  | 63% ~36s          
  |++++++++++++++++++++++++++++++++                  | 64% ~35s          
  |+++++++++++++++++++++++++++++++++                 | 65% ~34s          
  |+++++++++++++++++++++++++++++++++                 | 66% ~33s          
  |++++++++++++++++++++++++++++++++++                | 67% ~32s          
  |+++++++++++++++++++++++++++++++++++               | 68% ~31s          
  |+++++++++++++++++++++++++++++++++++               | 69% ~30s          
  |++++++++++++++++++++++++++++++++++++              | 70% ~29s          
  |++++++++++++++++++++++++++++++++++++              | 71% ~28s          
  |+++++++++++++++++++++++++++++++++++++             | 72% ~27s          
  |+++++++++++++++++++++++++++++++++++++             | 73% ~26s          
  |++++++++++++++++++++++++++++++++++++++            | 74% ~25s          
  |++++++++++++++++++++++++++++++++++++++            | 75% ~24s          
  |+++++++++++++++++++++++++++++++++++++++           | 76% ~23s          
  |+++++++++++++++++++++++++++++++++++++++           | 77% ~22s          
  |++++++++++++++++++++++++++++++++++++++++          | 78% ~21s          
  |++++++++++++++++++++++++++++++++++++++++          | 79% ~20s          
  |+++++++++++++++++++++++++++++++++++++++++         | 80% ~19s          
  |+++++++++++++++++++++++++++++++++++++++++         | 81% ~18s          
  |++++++++++++++++++++++++++++++++++++++++++        | 82% ~17s          
  |++++++++++++++++++++++++++++++++++++++++++        | 84% ~16s          
  |+++++++++++++++++++++++++++++++++++++++++++       | 85% ~15s          
  |+++++++++++++++++++++++++++++++++++++++++++       | 86% ~14s          
  |++++++++++++++++++++++++++++++++++++++++++++      | 87% ~13s          
  |++++++++++++++++++++++++++++++++++++++++++++      | 88% ~12s          
  |+++++++++++++++++++++++++++++++++++++++++++++     | 89% ~11s          
  |+++++++++++++++++++++++++++++++++++++++++++++     | 90% ~10s          
  |++++++++++++++++++++++++++++++++++++++++++++++    | 91% ~09s          
  |++++++++++++++++++++++++++++++++++++++++++++++    | 92% ~08s          
  |+++++++++++++++++++++++++++++++++++++++++++++++   | 93% ~07s          
  |+++++++++++++++++++++++++++++++++++++++++++++++   | 94% ~06s          
  |++++++++++++++++++++++++++++++++++++++++++++++++  | 95% ~05s          
  |++++++++++++++++++++++++++++++++++++++++++++++++  | 96% ~04s          
  |+++++++++++++++++++++++++++++++++++++++++++++++++ | 97% ~03s          
  |+++++++++++++++++++++++++++++++++++++++++++++++++ | 98% ~02s          
  |++++++++++++++++++++++++++++++++++++++++++++++++++| 99% ~01s          
  |++++++++++++++++++++++++++++++++++++++++++++++++++| 100% elapsed=01m 37s
```


```
write.csv(RNA_markers,"/Users/franziskadenk/Library/CloudStorage/OneDrive-SharedLibraries-King'sCollegeLondon/Denk Lab SharePoint - Mesenchymal_MS/markers.csv")
```


```
#identify mural cells
custom_colours <- c("#DEEDCF", "#99D492","#56B870","#1D9A6C","#137177", "#0A2F51")
feature_plot1 <- FeaturePlot(cbmc, features = "Notch3", min.cutoff = "q05", max.cutoff = "q95", order = TRUE, cols = custom_colours)
feature_plot2 <- FeaturePlot(cbmc, features = "Pdgfrb", min.cutoff = "q05", max.cutoff = "q95", order = TRUE, cols = custom_colours)
feature_plot3 <- FeaturePlot(cbmc, features = "Mcam", min.cutoff = "q05", max.cutoff = "q95", order = TRUE, cols = custom_colours)
Idents(cbmc) <- cbmc$seurat_clusters
DimPlot_cols <- DiscretePalette(14, palette = "stepped", shuffle = FALSE)
dim_plot <- DimPlot(cbmc, label=TRUE, cols=DimPlot_cols) + theme(legend.position = "none")
combined_plot <- (feature_plot1 | feature_plot2 | feature_plot3 | dim_plot) + plot_layout(ncol = 2, nrow = 2)
print(combined_plot)
```


```
#identify myelinating and non-myelinating Schwann cells
custom_colours <- c("#DEEDCF", "#99D492","#56B870","#1D9A6C","#137177", "#0A2F51")
feature_plot1 <- FeaturePlot(cbmc, features = "Sox10", min.cutoff = "q05", max.cutoff = "q95", order = TRUE, cols = custom_colours)
feature_plot2 <- FeaturePlot(cbmc, features = "Mbp", min.cutoff = "q05", max.cutoff = "q95", order = TRUE, cols = custom_colours)
feature_plot3 <- FeaturePlot(cbmc, features = "Col15a1", min.cutoff = "q05", max.cutoff = "q95", order = TRUE, cols = custom_colours)
Idents(cbmc) <- cbmc$seurat_clusters
DimPlot_cols <- DiscretePalette(14, palette = "stepped", shuffle = FALSE)
dim_plot <- DimPlot(cbmc, label=TRUE, cols=DimPlot_cols) + theme(legend.position = "none")
combined_plot <- (feature_plot1 | feature_plot2 | feature_plot3 | dim_plot) + plot_layout(ncol = 2, nrow = 2)
print(combined_plot)
```


```
#identify universal fibroblast populations
custom_colours <- c("#DEEDCF", "#99D492","#56B870","#1D9A6C","#137177", "#0A2F51")
feature_plot1 <- FeaturePlot(cbmc, features = "Col15a1", min.cutoff = "q05", max.cutoff = "q95", order = TRUE, cols = custom_colours)
feature_plot2 <- FeaturePlot(cbmc, features = "Pi16", min.cutoff = "q05", max.cutoff = "q95", order = TRUE, cols = custom_colours)
feature_plot3 <- FeaturePlot(cbmc, features = "Pdgfra", min.cutoff = "q05", max.cutoff = "q95", order = TRUE, cols = custom_colours)
Idents(cbmc) <- cbmc$seurat_clusters
DimPlot_cols <- DiscretePalette(14, palette = "stepped", shuffle = FALSE)
dim_plot <- DimPlot(cbmc, label=TRUE, cols=DimPlot_cols) + theme(legend.position = "none")
combined_plot <- (feature_plot1 | feature_plot2 | feature_plot3 | dim_plot) + plot_layout(ncol = 2, nrow = 2)
print(combined_plot)
```


```
#identify universal pathological fibroblast populations
custom_colours <- c("#DEEDCF", "#99D492","#56B870","#1D9A6C","#137177", "#0A2F51")
feature_plot1 <- FeaturePlot(cbmc, features = "Ccl19", min.cutoff = "q05", max.cutoff = "q95", order = TRUE, cols = custom_colours)
feature_plot2 <- FeaturePlot(cbmc, features = "Comp", min.cutoff = "q05", max.cutoff = "q95", order = TRUE, cols = custom_colours)
feature_plot3 <- FeaturePlot(cbmc, features = "Notch3", min.cutoff = "q05", max.cutoff = "q95", order = TRUE, cols = custom_colours)
Idents(cbmc) <- cbmc$seurat_clusters
DimPlot_cols <- DiscretePalette(14, palette = "stepped", shuffle = FALSE)
dim_plot <- DimPlot(cbmc, label=TRUE, cols=DimPlot_cols) + theme(legend.position = "none")
combined_plot <- (feature_plot1 | feature_plot2 | feature_plot3 | dim_plot) + plot_layout(ncol = 2, nrow = 2)
print(combined_plot)
```


```
#identify other barrier and endothelial cells populations
custom_colours <- c("#DEEDCF", "#99D492","#56B870","#1D9A6C","#137177", "#0A2F51")
feature_plot1 <- FeaturePlot(cbmc, features = "Cldn1", min.cutoff = "q05", max.cutoff = "q95", order = TRUE, cols = custom_colours)
feature_plot2 <- FeaturePlot(cbmc, features = "Pecam1", min.cutoff = "q05", max.cutoff = "q95", order = TRUE, cols = custom_colours)
feature_plot3 <- FeaturePlot(cbmc, features = "Tagln", min.cutoff = "q05", max.cutoff = "q95", order = TRUE, cols = custom_colours)
Idents(cbmc) <- cbmc$seurat_clusters
DimPlot_cols <- DiscretePalette(14, palette = "stepped", shuffle = FALSE)
dim_plot <- DimPlot(cbmc, label=TRUE, cols=DimPlot_cols) + theme(legend.position = "none")
combined_plot <- (feature_plot1 | feature_plot2 | feature_plot3 | dim_plot) + plot_layout(ncol = 2, nrow = 2)
print(combined_plot)
```


```
bmc <- (RenameIdents(cbmc, `0` = "FB_Ccl19", `1` = "SC", `9` = "mySC", `8` = "MC", `2` = "FB_Col15a1", `6` = "EC", `5` = "FB_Cldn1",`7` = "SC", `4` = "FB_Pi16", `3` = "MC", `10` = "IC"))
```


```
Warning message:
In do_once((if (is_R_CMD_check()) stop else warning)("The function xfun::isFALSE() will be deprecated in the future. Please ",  :
  The function xfun::isFALSE() will be deprecated in the future. Please consider using base::isFALSE(x) or identical(x, FALSE) instead.
```


```
bmc[["cluster_IDs"]] <- Idents(object = bmc) #to add clusters to metadata.
```


```
RNA_markers <- FindAllMarkers(bmc, min.diff.pct = 0.2)
```


```
Calculating cluster FB_Ccl19
```


```
  |                                                  | 0 % ~calculating  
  |+                                                 | 1 % ~10s          
  |++                                                | 3 % ~09s          
  |++                                                | 4 % ~09s          
  |+++                                               | 5 % ~09s          
  |++++                                              | 7 % ~08s          
  |++++                                              | 8 % ~08s          
  |+++++                                             | 9 % ~08s          
  |++++++                                            | 11% ~08s          
  |++++++                                            | 12% ~08s          
  |+++++++                                           | 13% ~08s          
  |++++++++                                          | 14% ~08s          
  |++++++++                                          | 16% ~07s          
  |+++++++++                                         | 17% ~07s          
  |++++++++++                                        | 18% ~07s          
  |++++++++++                                        | 20% ~07s          
  |+++++++++++                                       | 21% ~07s          
  |++++++++++++                                      | 22% ~07s          
  |++++++++++++                                      | 24% ~07s          
  |+++++++++++++                                     | 25% ~07s          
  |++++++++++++++                                    | 26% ~07s          
  |++++++++++++++                                    | 28% ~07s          
  |+++++++++++++++                                   | 29% ~07s          
  |++++++++++++++++                                  | 30% ~06s          
  |++++++++++++++++                                  | 32% ~06s          
  |+++++++++++++++++                                 | 33% ~06s          
  |++++++++++++++++++                                | 34% ~06s          
  |++++++++++++++++++                                | 36% ~06s          
  |+++++++++++++++++++                               | 37% ~06s          
  |++++++++++++++++++++                              | 38% ~06s          
  |++++++++++++++++++++                              | 39% ~05s          
  |+++++++++++++++++++++                             | 41% ~05s          
  |++++++++++++++++++++++                            | 42% ~05s          
  |++++++++++++++++++++++                            | 43% ~05s          
  |+++++++++++++++++++++++                           | 45% ~05s          
  |++++++++++++++++++++++++                          | 46% ~05s          
  |++++++++++++++++++++++++                          | 47% ~05s          
  |+++++++++++++++++++++++++                         | 49% ~05s          
  |+++++++++++++++++++++++++                         | 50% ~04s          
  |++++++++++++++++++++++++++                        | 51% ~04s          
  |+++++++++++++++++++++++++++                       | 53% ~04s          
  |+++++++++++++++++++++++++++                       | 54% ~04s          
  |++++++++++++++++++++++++++++                      | 55% ~04s          
  |+++++++++++++++++++++++++++++                     | 57% ~04s          
  |+++++++++++++++++++++++++++++                     | 58% ~04s          
  |++++++++++++++++++++++++++++++                    | 59% ~04s          
  |+++++++++++++++++++++++++++++++                   | 61% ~04s          
  |+++++++++++++++++++++++++++++++                   | 62% ~03s          
  |++++++++++++++++++++++++++++++++                  | 63% ~03s          
  |+++++++++++++++++++++++++++++++++                 | 64% ~03s          
  |+++++++++++++++++++++++++++++++++                 | 66% ~03s          
  |++++++++++++++++++++++++++++++++++                | 67% ~03s          
  |+++++++++++++++++++++++++++++++++++               | 68% ~03s          
  |+++++++++++++++++++++++++++++++++++               | 70% ~03s          
  |++++++++++++++++++++++++++++++++++++              | 71% ~03s          
  |+++++++++++++++++++++++++++++++++++++             | 72% ~02s          
  |+++++++++++++++++++++++++++++++++++++             | 74% ~02s          
  |++++++++++++++++++++++++++++++++++++++            | 75% ~02s          
  |+++++++++++++++++++++++++++++++++++++++           | 76% ~02s          
  |+++++++++++++++++++++++++++++++++++++++           | 78% ~02s          
  |++++++++++++++++++++++++++++++++++++++++          | 79% ~02s          
  |+++++++++++++++++++++++++++++++++++++++++         | 80% ~02s          
  |+++++++++++++++++++++++++++++++++++++++++         | 82% ~02s          
  |++++++++++++++++++++++++++++++++++++++++++        | 83% ~02s          
  |+++++++++++++++++++++++++++++++++++++++++++       | 84% ~01s          
  |+++++++++++++++++++++++++++++++++++++++++++       | 86% ~01s          
  |++++++++++++++++++++++++++++++++++++++++++++      | 87% ~01s          
  |+++++++++++++++++++++++++++++++++++++++++++++     | 88% ~01s          
  |+++++++++++++++++++++++++++++++++++++++++++++     | 89% ~01s          
  |++++++++++++++++++++++++++++++++++++++++++++++    | 91% ~01s          
  |+++++++++++++++++++++++++++++++++++++++++++++++   | 92% ~01s          
  |+++++++++++++++++++++++++++++++++++++++++++++++   | 93% ~01s          
  |++++++++++++++++++++++++++++++++++++++++++++++++  | 95% ~00s          
  |+++++++++++++++++++++++++++++++++++++++++++++++++ | 96% ~00s          
  |+++++++++++++++++++++++++++++++++++++++++++++++++ | 97% ~00s          
  |++++++++++++++++++++++++++++++++++++++++++++++++++| 99% ~00s          
  |++++++++++++++++++++++++++++++++++++++++++++++++++| 100% elapsed=09s
```


```
Calculating cluster SC
```


```
  |                                                  | 0 % ~calculating  
  |+                                                 | 1 % ~21s          
  |++                                                | 2 % ~20s          
  |++                                                | 3 % ~20s          
  |+++                                               | 4 % ~19s          
  |+++                                               | 5 % ~01m 09s      
  |++++                                              | 7 % ~60s          
  |++++                                              | 8 % ~53s          
  |+++++                                             | 9 % ~48s          
  |+++++                                             | 10% ~44s          
  |++++++                                            | 11% ~41s          
  |++++++                                            | 12% ~38s          
  |+++++++                                           | 13% ~36s          
  |++++++++                                          | 14% ~34s          
  |++++++++                                          | 15% ~33s          
  |+++++++++                                         | 16% ~31s          
  |+++++++++                                         | 17% ~30s          
  |++++++++++                                        | 18% ~29s          
  |++++++++++                                        | 20% ~27s          
  |+++++++++++                                       | 21% ~26s          
  |+++++++++++                                       | 22% ~26s          
  |++++++++++++                                      | 23% ~25s          
  |++++++++++++                                      | 24% ~24s          
  |+++++++++++++                                     | 25% ~23s          
  |++++++++++++++                                    | 26% ~22s          
  |++++++++++++++                                    | 27% ~22s          
  |+++++++++++++++                                   | 28% ~21s          
  |+++++++++++++++                                   | 29% ~21s          
  |++++++++++++++++                                  | 30% ~20s          
  |++++++++++++++++                                  | 32% ~19s          
  |+++++++++++++++++                                 | 33% ~19s          
  |+++++++++++++++++                                 | 34% ~18s          
  |++++++++++++++++++                                | 35% ~18s          
  |++++++++++++++++++                                | 36% ~18s          
  |+++++++++++++++++++                               | 37% ~17s          
  |++++++++++++++++++++                              | 38% ~17s          
  |++++++++++++++++++++                              | 39% ~16s          
  |+++++++++++++++++++++                             | 40% ~16s          
  |+++++++++++++++++++++                             | 41% ~15s          
  |++++++++++++++++++++++                            | 42% ~15s          
  |++++++++++++++++++++++                            | 43% ~15s          
  |+++++++++++++++++++++++                           | 45% ~14s          
  |+++++++++++++++++++++++                           | 46% ~14s          
  |++++++++++++++++++++++++                          | 47% ~14s          
  |++++++++++++++++++++++++                          | 48% ~13s          
  |+++++++++++++++++++++++++                         | 49% ~13s          
  |+++++++++++++++++++++++++                         | 50% ~13s          
  |++++++++++++++++++++++++++                        | 51% ~12s          
  |+++++++++++++++++++++++++++                       | 52% ~12s          
  |+++++++++++++++++++++++++++                       | 53% ~12s          
  |++++++++++++++++++++++++++++                      | 54% ~11s          
  |++++++++++++++++++++++++++++                      | 55% ~11s          
  |+++++++++++++++++++++++++++++                     | 57% ~11s          
  |+++++++++++++++++++++++++++++                     | 58% ~10s          
  |++++++++++++++++++++++++++++++                    | 59% ~10s          
  |++++++++++++++++++++++++++++++                    | 60% ~10s          
  |+++++++++++++++++++++++++++++++                   | 61% ~10s          
  |+++++++++++++++++++++++++++++++                   | 62% ~09s          
  |++++++++++++++++++++++++++++++++                  | 63% ~09s          
  |+++++++++++++++++++++++++++++++++                 | 64% ~09s          
  |+++++++++++++++++++++++++++++++++                 | 65% ~08s          
  |++++++++++++++++++++++++++++++++++                | 66% ~08s          
  |++++++++++++++++++++++++++++++++++                | 67% ~08s          
  |+++++++++++++++++++++++++++++++++++               | 68% ~08s          
  |+++++++++++++++++++++++++++++++++++               | 70% ~07s          
  |++++++++++++++++++++++++++++++++++++              | 71% ~07s          
  |++++++++++++++++++++++++++++++++++++              | 72% ~07s          
  |+++++++++++++++++++++++++++++++++++++             | 73% ~06s          
  |+++++++++++++++++++++++++++++++++++++             | 74% ~06s          
  |++++++++++++++++++++++++++++++++++++++            | 75% ~06s          
  |+++++++++++++++++++++++++++++++++++++++           | 76% ~06s          
  |+++++++++++++++++++++++++++++++++++++++           | 77% ~05s          
  |++++++++++++++++++++++++++++++++++++++++          | 78% ~05s          
  |++++++++++++++++++++++++++++++++++++++++          | 79% ~05s          
  |+++++++++++++++++++++++++++++++++++++++++         | 80% ~05s          
  |+++++++++++++++++++++++++++++++++++++++++         | 82% ~04s          
  |++++++++++++++++++++++++++++++++++++++++++        | 83% ~04s          
  |++++++++++++++++++++++++++++++++++++++++++        | 84% ~04s          
  |+++++++++++++++++++++++++++++++++++++++++++       | 85% ~04s          
  |+++++++++++++++++++++++++++++++++++++++++++       | 86% ~03s          
  |++++++++++++++++++++++++++++++++++++++++++++      | 87% ~03s          
  |+++++++++++++++++++++++++++++++++++++++++++++     | 88% ~03s          
  |+++++++++++++++++++++++++++++++++++++++++++++     | 89% ~03s          
  |++++++++++++++++++++++++++++++++++++++++++++++    | 90% ~02s          
  |++++++++++++++++++++++++++++++++++++++++++++++    | 91% ~02s          
  |+++++++++++++++++++++++++++++++++++++++++++++++   | 92% ~02s          
  |+++++++++++++++++++++++++++++++++++++++++++++++   | 93% ~01s          
  |++++++++++++++++++++++++++++++++++++++++++++++++  | 95% ~01s          
  |++++++++++++++++++++++++++++++++++++++++++++++++  | 96% ~01s          
  |+++++++++++++++++++++++++++++++++++++++++++++++++ | 97% ~01s          
  |+++++++++++++++++++++++++++++++++++++++++++++++++ | 98% ~00s          
  |++++++++++++++++++++++++++++++++++++++++++++++++++| 99% ~00s          
  |++++++++++++++++++++++++++++++++++++++++++++++++++| 100% elapsed=23s
```


```
Calculating cluster mySC
```


```
  |                                                  | 0 % ~calculating  
  |+                                                 | 1 % ~58s          
  |++                                                | 2 % ~58s          
  |++                                                | 3 % ~59s          
  |+++                                               | 4 % ~59s          
  |+++                                               | 5 % ~58s          
  |++++                                              | 6 % ~57s          
  |++++                                              | 7 % ~56s          
  |+++++                                             | 8 % ~55s          
  |+++++                                             | 9 % ~54s          
  |++++++                                            | 10% ~53s          
  |++++++                                            | 11% ~52s          
  |+++++++                                           | 12% ~52s          
  |+++++++                                           | 14% ~51s          
  |++++++++                                          | 15% ~50s          
  |++++++++                                          | 16% ~50s          
  |+++++++++                                         | 17% ~49s          
  |+++++++++                                         | 18% ~48s          
  |++++++++++                                        | 19% ~48s          
  |++++++++++                                        | 20% ~47s          
  |+++++++++++                                       | 21% ~46s          
  |+++++++++++                                       | 22% ~46s          
  |++++++++++++                                      | 23% ~45s          
  |++++++++++++                                      | 24% ~44s          
  |+++++++++++++                                     | 25% ~44s          
  |++++++++++++++                                    | 26% ~43s          
  |++++++++++++++                                    | 27% ~42s          
  |+++++++++++++++                                   | 28% ~42s          
  |+++++++++++++++                                   | 29% ~41s          
  |++++++++++++++++                                  | 30% ~40s          
  |++++++++++++++++                                  | 31% ~40s          
  |+++++++++++++++++                                 | 32% ~39s          
  |+++++++++++++++++                                 | 33% ~39s          
  |++++++++++++++++++                                | 34% ~38s          
  |++++++++++++++++++                                | 35% ~37s          
  |+++++++++++++++++++                               | 36% ~37s          
  |+++++++++++++++++++                               | 38% ~36s          
  |++++++++++++++++++++                              | 39% ~36s          
  |++++++++++++++++++++                              | 40% ~35s          
  |+++++++++++++++++++++                             | 41% ~34s          
  |+++++++++++++++++++++                             | 42% ~34s          
  |++++++++++++++++++++++                            | 43% ~33s          
  |++++++++++++++++++++++                            | 44% ~33s          
  |+++++++++++++++++++++++                           | 45% ~32s          
  |+++++++++++++++++++++++                           | 46% ~32s          
  |++++++++++++++++++++++++                          | 47% ~31s          
  |++++++++++++++++++++++++                          | 48% ~30s          
  |+++++++++++++++++++++++++                         | 49% ~30s          
  |+++++++++++++++++++++++++                         | 50% ~29s          
  |++++++++++++++++++++++++++                        | 51% ~29s          
  |+++++++++++++++++++++++++++                       | 52% ~28s          
  |+++++++++++++++++++++++++++                       | 53% ~27s          
  |++++++++++++++++++++++++++++                      | 54% ~27s          
  |++++++++++++++++++++++++++++                      | 55% ~26s          
  |+++++++++++++++++++++++++++++                     | 56% ~26s          
  |+++++++++++++++++++++++++++++                     | 57% ~25s          
  |++++++++++++++++++++++++++++++                    | 58% ~25s          
  |++++++++++++++++++++++++++++++                    | 59% ~24s          
  |+++++++++++++++++++++++++++++++                   | 60% ~24s          
  |+++++++++++++++++++++++++++++++                   | 61% ~23s          
  |++++++++++++++++++++++++++++++++                  | 62% ~22s          
  |++++++++++++++++++++++++++++++++                  | 64% ~22s          
  |+++++++++++++++++++++++++++++++++                 | 65% ~21s          
  |+++++++++++++++++++++++++++++++++                 | 66% ~20s          
  |++++++++++++++++++++++++++++++++++                | 67% ~20s          
  |++++++++++++++++++++++++++++++++++                | 68% ~19s          
  |+++++++++++++++++++++++++++++++++++               | 69% ~19s          
  |+++++++++++++++++++++++++++++++++++               | 70% ~18s          
  |++++++++++++++++++++++++++++++++++++              | 71% ~17s          
  |++++++++++++++++++++++++++++++++++++              | 72% ~17s          
  |+++++++++++++++++++++++++++++++++++++             | 73% ~16s          
  |+++++++++++++++++++++++++++++++++++++             | 74% ~15s          
  |++++++++++++++++++++++++++++++++++++++            | 75% ~15s          
  |+++++++++++++++++++++++++++++++++++++++           | 76% ~14s          
  |+++++++++++++++++++++++++++++++++++++++           | 77% ~14s          
  |++++++++++++++++++++++++++++++++++++++++          | 78% ~13s          
  |++++++++++++++++++++++++++++++++++++++++          | 79% ~12s          
  |+++++++++++++++++++++++++++++++++++++++++         | 80% ~12s          
  |+++++++++++++++++++++++++++++++++++++++++         | 81% ~11s          
  |++++++++++++++++++++++++++++++++++++++++++        | 82% ~10s          
  |++++++++++++++++++++++++++++++++++++++++++        | 83% ~10s          
  |+++++++++++++++++++++++++++++++++++++++++++       | 84% ~09s          
  |+++++++++++++++++++++++++++++++++++++++++++       | 85% ~09s          
  |++++++++++++++++++++++++++++++++++++++++++++      | 86% ~08s          
  |++++++++++++++++++++++++++++++++++++++++++++      | 88% ~07s          
  |+++++++++++++++++++++++++++++++++++++++++++++     | 89% ~07s          
  |+++++++++++++++++++++++++++++++++++++++++++++     | 90% ~06s          
  |++++++++++++++++++++++++++++++++++++++++++++++    | 91% ~05s          
  |++++++++++++++++++++++++++++++++++++++++++++++    | 92% ~05s          
  |+++++++++++++++++++++++++++++++++++++++++++++++   | 93% ~04s          
  |+++++++++++++++++++++++++++++++++++++++++++++++   | 94% ~04s          
  |++++++++++++++++++++++++++++++++++++++++++++++++  | 95% ~03s          
  |++++++++++++++++++++++++++++++++++++++++++++++++  | 96% ~02s          
  |+++++++++++++++++++++++++++++++++++++++++++++++++ | 97% ~02s          
  |+++++++++++++++++++++++++++++++++++++++++++++++++ | 98% ~01s          
  |++++++++++++++++++++++++++++++++++++++++++++++++++| 99% ~01s          
  |++++++++++++++++++++++++++++++++++++++++++++++++++| 100% elapsed=58s
```


```
Calculating cluster MC
```


```
  |                                                  | 0 % ~calculating  
  |+                                                 | 1 % ~14s          
  |++                                                | 2 % ~14s          
  |++                                                | 3 % ~14s          
  |+++                                               | 5 % ~13s          
  |+++                                               | 6 % ~13s          
  |++++                                              | 7 % ~13s          
  |++++                                              | 8 % ~13s          
  |+++++                                             | 9 % ~13s          
  |++++++                                            | 10% ~13s          
  |++++++                                            | 11% ~12s          
  |+++++++                                           | 12% ~12s          
  |+++++++                                           | 14% ~12s          
  |++++++++                                          | 15% ~12s          
  |++++++++                                          | 16% ~12s          
  |+++++++++                                         | 17% ~12s          
  |++++++++++                                        | 18% ~12s          
  |++++++++++                                        | 19% ~11s          
  |+++++++++++                                       | 20% ~11s          
  |+++++++++++                                       | 22% ~11s          
  |++++++++++++                                      | 23% ~11s          
  |++++++++++++                                      | 24% ~11s          
  |+++++++++++++                                     | 25% ~11s          
  |++++++++++++++                                    | 26% ~10s          
  |++++++++++++++                                    | 27% ~10s          
  |+++++++++++++++                                   | 28% ~10s          
  |+++++++++++++++                                   | 30% ~10s          
  |++++++++++++++++                                  | 31% ~10s          
  |++++++++++++++++                                  | 32% ~10s          
  |+++++++++++++++++                                 | 33% ~09s          
  |++++++++++++++++++                                | 34% ~09s          
  |++++++++++++++++++                                | 35% ~09s          
  |+++++++++++++++++++                               | 36% ~09s          
  |+++++++++++++++++++                               | 38% ~09s          
  |++++++++++++++++++++                              | 39% ~09s          
  |++++++++++++++++++++                              | 40% ~08s          
  |+++++++++++++++++++++                             | 41% ~08s          
  |++++++++++++++++++++++                            | 42% ~08s          
  |++++++++++++++++++++++                            | 43% ~08s          
  |+++++++++++++++++++++++                           | 44% ~08s          
  |+++++++++++++++++++++++                           | 45% ~08s          
  |++++++++++++++++++++++++                          | 47% ~07s          
  |++++++++++++++++++++++++                          | 48% ~07s          
  |+++++++++++++++++++++++++                         | 49% ~07s          
  |+++++++++++++++++++++++++                         | 50% ~07s          
  |++++++++++++++++++++++++++                        | 51% ~07s          
  |+++++++++++++++++++++++++++                       | 52% ~07s          
  |+++++++++++++++++++++++++++                       | 53% ~07s          
  |++++++++++++++++++++++++++++                      | 55% ~06s          
  |++++++++++++++++++++++++++++                      | 56% ~06s          
  |+++++++++++++++++++++++++++++                     | 57% ~06s          
  |+++++++++++++++++++++++++++++                     | 58% ~06s          
  |++++++++++++++++++++++++++++++                    | 59% ~06s          
  |+++++++++++++++++++++++++++++++                   | 60% ~06s          
  |+++++++++++++++++++++++++++++++                   | 61% ~05s          
  |++++++++++++++++++++++++++++++++                  | 62% ~05s          
  |++++++++++++++++++++++++++++++++                  | 64% ~05s          
  |+++++++++++++++++++++++++++++++++                 | 65% ~05s          
  |+++++++++++++++++++++++++++++++++                 | 66% ~05s          
  |++++++++++++++++++++++++++++++++++                | 67% ~05s          
  |+++++++++++++++++++++++++++++++++++               | 68% ~04s          
  |+++++++++++++++++++++++++++++++++++               | 69% ~04s          
  |++++++++++++++++++++++++++++++++++++              | 70% ~04s          
  |++++++++++++++++++++++++++++++++++++              | 72% ~04s          
  |+++++++++++++++++++++++++++++++++++++             | 73% ~04s          
  |+++++++++++++++++++++++++++++++++++++             | 74% ~04s          
  |++++++++++++++++++++++++++++++++++++++            | 75% ~03s          
  |+++++++++++++++++++++++++++++++++++++++           | 76% ~03s          
  |+++++++++++++++++++++++++++++++++++++++           | 77% ~03s          
  |++++++++++++++++++++++++++++++++++++++++          | 78% ~03s          
  |++++++++++++++++++++++++++++++++++++++++          | 80% ~03s          
  |+++++++++++++++++++++++++++++++++++++++++         | 81% ~03s          
  |+++++++++++++++++++++++++++++++++++++++++         | 82% ~03s          
  |++++++++++++++++++++++++++++++++++++++++++        | 83% ~02s          
  |+++++++++++++++++++++++++++++++++++++++++++       | 84% ~02s          
  |+++++++++++++++++++++++++++++++++++++++++++       | 85% ~02s          
  |++++++++++++++++++++++++++++++++++++++++++++      | 86% ~02s          
  |++++++++++++++++++++++++++++++++++++++++++++      | 88% ~02s          
  |+++++++++++++++++++++++++++++++++++++++++++++     | 89% ~02s          
  |+++++++++++++++++++++++++++++++++++++++++++++     | 90% ~01s          
  |++++++++++++++++++++++++++++++++++++++++++++++    | 91% ~01s          
  |+++++++++++++++++++++++++++++++++++++++++++++++   | 92% ~01s          
  |+++++++++++++++++++++++++++++++++++++++++++++++   | 93% ~01s          
  |++++++++++++++++++++++++++++++++++++++++++++++++  | 94% ~01s          
  |++++++++++++++++++++++++++++++++++++++++++++++++  | 95% ~01s          
  |+++++++++++++++++++++++++++++++++++++++++++++++++ | 97% ~00s          
  |+++++++++++++++++++++++++++++++++++++++++++++++++ | 98% ~00s          
  |++++++++++++++++++++++++++++++++++++++++++++++++++| 99% ~00s          
  |++++++++++++++++++++++++++++++++++++++++++++++++++| 100% elapsed=14s
```


```
Calculating cluster FB_Col15a1
```


```
  |                                                  | 0 % ~calculating  
  |+                                                 | 1 % ~06s          
  |++                                                | 3 % ~06s          
  |++                                                | 4 % ~06s          
  |+++                                               | 5 % ~05s          
  |++++                                              | 7 % ~05s          
  |++++                                              | 8 % ~05s          
  |+++++                                             | 9 % ~05s          
  |++++++                                            | 11% ~05s          
  |++++++                                            | 12% ~05s          
  |+++++++                                           | 13% ~05s          
  |++++++++                                          | 14% ~05s          
  |++++++++                                          | 16% ~05s          
  |+++++++++                                         | 17% ~05s          
  |++++++++++                                        | 18% ~05s          
  |++++++++++                                        | 20% ~05s          
  |+++++++++++                                       | 21% ~04s          
  |++++++++++++                                      | 22% ~04s          
  |++++++++++++                                      | 24% ~04s          
  |+++++++++++++                                     | 25% ~04s          
  |++++++++++++++                                    | 26% ~04s          
  |++++++++++++++                                    | 28% ~04s          
  |+++++++++++++++                                   | 29% ~04s          
  |++++++++++++++++                                  | 30% ~04s          
  |++++++++++++++++                                  | 32% ~04s          
  |+++++++++++++++++                                 | 33% ~04s          
  |++++++++++++++++++                                | 34% ~04s          
  |++++++++++++++++++                                | 36% ~04s          
  |+++++++++++++++++++                               | 37% ~04s          
  |++++++++++++++++++++                              | 38% ~04s          
  |++++++++++++++++++++                              | 39% ~03s          
  |+++++++++++++++++++++                             | 41% ~03s          
  |++++++++++++++++++++++                            | 42% ~03s          
  |++++++++++++++++++++++                            | 43% ~03s          
  |+++++++++++++++++++++++                           | 45% ~03s          
  |++++++++++++++++++++++++                          | 46% ~03s          
  |++++++++++++++++++++++++                          | 47% ~03s          
  |+++++++++++++++++++++++++                         | 49% ~03s          
  |+++++++++++++++++++++++++                         | 50% ~03s          
  |++++++++++++++++++++++++++                        | 51% ~03s          
  |+++++++++++++++++++++++++++                       | 53% ~03s          
  |+++++++++++++++++++++++++++                       | 54% ~03s          
  |++++++++++++++++++++++++++++                      | 55% ~03s          
  |+++++++++++++++++++++++++++++                     | 57% ~02s          
  |+++++++++++++++++++++++++++++                     | 58% ~02s          
  |++++++++++++++++++++++++++++++                    | 59% ~02s          
  |+++++++++++++++++++++++++++++++                   | 61% ~02s          
  |+++++++++++++++++++++++++++++++                   | 62% ~02s          
  |++++++++++++++++++++++++++++++++                  | 63% ~02s          
  |+++++++++++++++++++++++++++++++++                 | 64% ~02s          
  |+++++++++++++++++++++++++++++++++                 | 66% ~02s          
  |++++++++++++++++++++++++++++++++++                | 67% ~02s          
  |+++++++++++++++++++++++++++++++++++               | 68% ~02s          
  |+++++++++++++++++++++++++++++++++++               | 70% ~02s          
  |++++++++++++++++++++++++++++++++++++              | 71% ~02s          
  |+++++++++++++++++++++++++++++++++++++             | 72% ~02s          
  |+++++++++++++++++++++++++++++++++++++             | 74% ~02s          
  |++++++++++++++++++++++++++++++++++++++            | 75% ~01s          
  |+++++++++++++++++++++++++++++++++++++++           | 76% ~01s          
  |+++++++++++++++++++++++++++++++++++++++           | 78% ~01s          
  |++++++++++++++++++++++++++++++++++++++++          | 79% ~01s          
  |+++++++++++++++++++++++++++++++++++++++++         | 80% ~01s          
  |+++++++++++++++++++++++++++++++++++++++++         | 82% ~01s          
  |++++++++++++++++++++++++++++++++++++++++++        | 83% ~01s          
  |+++++++++++++++++++++++++++++++++++++++++++       | 84% ~01s          
  |+++++++++++++++++++++++++++++++++++++++++++       | 86% ~01s          
  |++++++++++++++++++++++++++++++++++++++++++++      | 87% ~01s          
  |+++++++++++++++++++++++++++++++++++++++++++++     | 88% ~01s          
  |+++++++++++++++++++++++++++++++++++++++++++++     | 89% ~01s          
  |++++++++++++++++++++++++++++++++++++++++++++++    | 91% ~01s          
  |+++++++++++++++++++++++++++++++++++++++++++++++   | 92% ~00s          
  |+++++++++++++++++++++++++++++++++++++++++++++++   | 93% ~00s          
  |++++++++++++++++++++++++++++++++++++++++++++++++  | 95% ~00s          
  |+++++++++++++++++++++++++++++++++++++++++++++++++ | 96% ~00s          
  |+++++++++++++++++++++++++++++++++++++++++++++++++ | 97% ~00s          
  |++++++++++++++++++++++++++++++++++++++++++++++++++| 99% ~00s          
  |++++++++++++++++++++++++++++++++++++++++++++++++++| 100% elapsed=06s
```


```
Calculating cluster EC
```


```
  |                                                  | 0 % ~calculating  
  |+                                                 | 1 % ~42s          
  |+                                                 | 2 % ~42s          
  |++                                                | 3 % ~42s          
  |++                                                | 4 % ~41s          
  |+++                                               | 5 % ~41s          
  |+++                                               | 6 % ~40s          
  |++++                                              | 7 % ~39s          
  |++++                                              | 8 % ~39s          
  |+++++                                             | 9 % ~38s          
  |+++++                                             | 10% ~38s          
  |++++++                                            | 11% ~37s          
  |++++++                                            | 12% ~37s          
  |+++++++                                           | 13% ~36s          
  |+++++++                                           | 14% ~36s          
  |++++++++                                          | 15% ~35s          
  |++++++++                                          | 16% ~35s          
  |+++++++++                                         | 17% ~34s          
  |+++++++++                                         | 18% ~34s          
  |++++++++++                                        | 19% ~34s          
  |++++++++++                                        | 20% ~33s          
  |+++++++++++                                       | 21% ~33s          
  |+++++++++++                                       | 22% ~32s          
  |++++++++++++                                      | 23% ~32s          
  |++++++++++++                                      | 24% ~31s          
  |+++++++++++++                                     | 25% ~31s          
  |+++++++++++++                                     | 26% ~31s          
  |++++++++++++++                                    | 27% ~30s          
  |++++++++++++++                                    | 28% ~30s          
  |+++++++++++++++                                   | 29% ~29s          
  |+++++++++++++++                                   | 30% ~29s          
  |++++++++++++++++                                  | 31% ~29s          
  |++++++++++++++++                                  | 32% ~28s          
  |+++++++++++++++++                                 | 33% ~28s          
  |+++++++++++++++++                                 | 34% ~27s          
  |++++++++++++++++++                                | 35% ~27s          
  |++++++++++++++++++                                | 36% ~27s          
  |+++++++++++++++++++                               | 37% ~26s          
  |+++++++++++++++++++                               | 38% ~26s          
  |++++++++++++++++++++                              | 39% ~25s          
  |++++++++++++++++++++                              | 40% ~25s          
  |+++++++++++++++++++++                             | 41% ~24s          
  |+++++++++++++++++++++                             | 42% ~24s          
  |++++++++++++++++++++++                            | 43% ~24s          
  |++++++++++++++++++++++                            | 44% ~23s          
  |+++++++++++++++++++++++                           | 45% ~23s          
  |+++++++++++++++++++++++                           | 46% ~22s          
  |++++++++++++++++++++++++                          | 47% ~22s          
  |++++++++++++++++++++++++                          | 48% ~22s          
  |+++++++++++++++++++++++++                         | 49% ~21s          
  |+++++++++++++++++++++++++                         | 50% ~21s          
  |++++++++++++++++++++++++++                        | 51% ~20s          
  |++++++++++++++++++++++++++                        | 52% ~20s          
  |+++++++++++++++++++++++++++                       | 53% ~20s          
  |+++++++++++++++++++++++++++                       | 54% ~19s          
  |++++++++++++++++++++++++++++                      | 55% ~19s          
  |++++++++++++++++++++++++++++                      | 56% ~18s          
  |+++++++++++++++++++++++++++++                     | 57% ~18s          
  |+++++++++++++++++++++++++++++                     | 58% ~18s          
  |++++++++++++++++++++++++++++++                    | 59% ~17s          
  |++++++++++++++++++++++++++++++                    | 60% ~17s          
  |+++++++++++++++++++++++++++++++                   | 61% ~16s          
  |+++++++++++++++++++++++++++++++                   | 62% ~16s          
  |++++++++++++++++++++++++++++++++                  | 63% ~15s          
  |++++++++++++++++++++++++++++++++                  | 64% ~15s          
  |+++++++++++++++++++++++++++++++++                 | 65% ~15s          
  |+++++++++++++++++++++++++++++++++                 | 66% ~14s          
  |++++++++++++++++++++++++++++++++++                | 67% ~14s          
  |++++++++++++++++++++++++++++++++++                | 68% ~14s          
  |+++++++++++++++++++++++++++++++++++               | 69% ~13s          
  |+++++++++++++++++++++++++++++++++++               | 70% ~13s          
  |++++++++++++++++++++++++++++++++++++              | 71% ~12s          
  |++++++++++++++++++++++++++++++++++++              | 72% ~12s          
  |+++++++++++++++++++++++++++++++++++++             | 73% ~11s          
  |+++++++++++++++++++++++++++++++++++++             | 74% ~11s          
  |++++++++++++++++++++++++++++++++++++++            | 75% ~11s          
  |++++++++++++++++++++++++++++++++++++++            | 76% ~10s          
  |+++++++++++++++++++++++++++++++++++++++           | 77% ~10s          
  |+++++++++++++++++++++++++++++++++++++++           | 78% ~09s          
  |++++++++++++++++++++++++++++++++++++++++          | 79% ~09s          
  |++++++++++++++++++++++++++++++++++++++++          | 80% ~08s          
  |+++++++++++++++++++++++++++++++++++++++++         | 81% ~08s          
  |+++++++++++++++++++++++++++++++++++++++++         | 82% ~08s          
  |++++++++++++++++++++++++++++++++++++++++++        | 83% ~07s          
  |++++++++++++++++++++++++++++++++++++++++++        | 84% ~07s          
  |+++++++++++++++++++++++++++++++++++++++++++       | 85% ~06s          
  |+++++++++++++++++++++++++++++++++++++++++++       | 86% ~06s          
  |++++++++++++++++++++++++++++++++++++++++++++      | 87% ~05s          
  |++++++++++++++++++++++++++++++++++++++++++++      | 88% ~05s          
  |+++++++++++++++++++++++++++++++++++++++++++++     | 89% ~05s          
  |+++++++++++++++++++++++++++++++++++++++++++++     | 90% ~04s          
  |++++++++++++++++++++++++++++++++++++++++++++++    | 91% ~04s          
  |++++++++++++++++++++++++++++++++++++++++++++++    | 92% ~03s          
  |+++++++++++++++++++++++++++++++++++++++++++++++   | 93% ~03s          
  |+++++++++++++++++++++++++++++++++++++++++++++++   | 94% ~03s          
  |++++++++++++++++++++++++++++++++++++++++++++++++  | 95% ~02s          
  |++++++++++++++++++++++++++++++++++++++++++++++++  | 96% ~02s          
  |+++++++++++++++++++++++++++++++++++++++++++++++++ | 97% ~01s          
  |+++++++++++++++++++++++++++++++++++++++++++++++++ | 98% ~01s          
  |++++++++++++++++++++++++++++++++++++++++++++++++++| 99% ~00s          
  |++++++++++++++++++++++++++++++++++++++++++++++++++| 100% elapsed=42s
```


```
Calculating cluster FB_Cldn1
```


```
  |                                                  | 0 % ~calculating  
  |+                                                 | 1 % ~08s          
  |++                                                | 2 % ~08s          
  |++                                                | 4 % ~08s          
  |+++                                               | 5 % ~08s          
  |++++                                              | 6 % ~07s          
  |++++                                              | 8 % ~07s          
  |+++++                                             | 9 % ~07s          
  |+++++                                             | 10% ~07s          
  |++++++                                            | 11% ~07s          
  |+++++++                                           | 12% ~07s          
  |+++++++                                           | 14% ~07s          
  |++++++++                                          | 15% ~07s          
  |+++++++++                                         | 16% ~07s          
  |+++++++++                                         | 18% ~07s          
  |++++++++++                                        | 19% ~06s          
  |++++++++++                                        | 20% ~06s          
  |+++++++++++                                       | 21% ~06s          
  |++++++++++++                                      | 22% ~06s          
  |++++++++++++                                      | 24% ~06s          
  |+++++++++++++                                     | 25% ~06s          
  |++++++++++++++                                    | 26% ~06s          
  |++++++++++++++                                    | 28% ~06s          
  |+++++++++++++++                                   | 29% ~06s          
  |+++++++++++++++                                   | 30% ~06s          
  |++++++++++++++++                                  | 31% ~05s          
  |+++++++++++++++++                                 | 32% ~05s          
  |+++++++++++++++++                                 | 34% ~05s          
  |++++++++++++++++++                                | 35% ~05s          
  |+++++++++++++++++++                               | 36% ~05s          
  |+++++++++++++++++++                               | 38% ~05s          
  |++++++++++++++++++++                              | 39% ~05s          
  |++++++++++++++++++++                              | 40% ~05s          
  |+++++++++++++++++++++                             | 41% ~05s          
  |++++++++++++++++++++++                            | 42% ~05s          
  |++++++++++++++++++++++                            | 44% ~04s          
  |+++++++++++++++++++++++                           | 45% ~05s          
  |++++++++++++++++++++++++                          | 46% ~04s          
  |++++++++++++++++++++++++                          | 48% ~04s          
  |+++++++++++++++++++++++++                         | 49% ~04s          
  |+++++++++++++++++++++++++                         | 50% ~04s          
  |++++++++++++++++++++++++++                        | 51% ~04s          
  |+++++++++++++++++++++++++++                       | 52% ~04s          
  |+++++++++++++++++++++++++++                       | 54% ~04s          
  |++++++++++++++++++++++++++++                      | 55% ~04s          
  |+++++++++++++++++++++++++++++                     | 56% ~04s          
  |+++++++++++++++++++++++++++++                     | 58% ~03s          
  |++++++++++++++++++++++++++++++                    | 59% ~03s          
  |++++++++++++++++++++++++++++++                    | 60% ~03s          
  |+++++++++++++++++++++++++++++++                   | 61% ~03s          
  |++++++++++++++++++++++++++++++++                  | 62% ~03s          
  |++++++++++++++++++++++++++++++++                  | 64% ~03s          
  |+++++++++++++++++++++++++++++++++                 | 65% ~03s          
  |++++++++++++++++++++++++++++++++++                | 66% ~03s          
  |++++++++++++++++++++++++++++++++++                | 68% ~03s          
  |+++++++++++++++++++++++++++++++++++               | 69% ~03s          
  |+++++++++++++++++++++++++++++++++++               | 70% ~02s          
  |++++++++++++++++++++++++++++++++++++              | 71% ~02s          
  |+++++++++++++++++++++++++++++++++++++             | 72% ~02s          
  |+++++++++++++++++++++++++++++++++++++             | 74% ~02s          
  |++++++++++++++++++++++++++++++++++++++            | 75% ~02s          
  |+++++++++++++++++++++++++++++++++++++++           | 76% ~02s          
  |+++++++++++++++++++++++++++++++++++++++           | 78% ~02s          
  |++++++++++++++++++++++++++++++++++++++++          | 79% ~02s          
  |++++++++++++++++++++++++++++++++++++++++          | 80% ~02s          
  |+++++++++++++++++++++++++++++++++++++++++         | 81% ~02s          
  |++++++++++++++++++++++++++++++++++++++++++        | 82% ~01s          
  |++++++++++++++++++++++++++++++++++++++++++        | 84% ~01s          
  |+++++++++++++++++++++++++++++++++++++++++++       | 85% ~01s          
  |++++++++++++++++++++++++++++++++++++++++++++      | 86% ~01s          
  |++++++++++++++++++++++++++++++++++++++++++++      | 88% ~01s          
  |+++++++++++++++++++++++++++++++++++++++++++++     | 89% ~01s          
  |+++++++++++++++++++++++++++++++++++++++++++++     | 90% ~01s          
  |++++++++++++++++++++++++++++++++++++++++++++++    | 91% ~01s          
  |+++++++++++++++++++++++++++++++++++++++++++++++   | 92% ~01s          
  |+++++++++++++++++++++++++++++++++++++++++++++++   | 94% ~01s          
  |++++++++++++++++++++++++++++++++++++++++++++++++  | 95% ~00s          
  |+++++++++++++++++++++++++++++++++++++++++++++++++ | 96% ~00s          
  |+++++++++++++++++++++++++++++++++++++++++++++++++ | 98% ~00s          
  |++++++++++++++++++++++++++++++++++++++++++++++++++| 99% ~00s          
  |++++++++++++++++++++++++++++++++++++++++++++++++++| 100% elapsed=08s
```


```
Calculating cluster FB_Pi16
```


```
  |                                                  | 0 % ~calculating  
  |+                                                 | 1 % ~12s          
  |++                                                | 2 % ~13s          
  |++                                                | 3 % ~12s          
  |+++                                               | 4 % ~12s          
  |+++                                               | 5 % ~12s          
  |++++                                              | 6 % ~12s          
  |++++                                              | 7 % ~12s          
  |+++++                                             | 8 % ~11s          
  |+++++                                             | 9 % ~11s          
  |++++++                                            | 10% ~11s          
  |++++++                                            | 11% ~11s          
  |+++++++                                           | 12% ~11s          
  |+++++++                                           | 14% ~11s          
  |++++++++                                          | 15% ~13s          
  |++++++++                                          | 16% ~13s          
  |+++++++++                                         | 17% ~13s          
  |+++++++++                                         | 18% ~12s          
  |++++++++++                                        | 19% ~12s          
  |++++++++++                                        | 20% ~12s          
  |+++++++++++                                       | 21% ~12s          
  |+++++++++++                                       | 22% ~11s          
  |++++++++++++                                      | 23% ~11s          
  |++++++++++++                                      | 24% ~11s          
  |+++++++++++++                                     | 25% ~11s          
  |++++++++++++++                                    | 26% ~11s          
  |++++++++++++++                                    | 27% ~10s          
  |+++++++++++++++                                   | 28% ~10s          
  |+++++++++++++++                                   | 29% ~10s          
  |++++++++++++++++                                  | 30% ~10s          
  |++++++++++++++++                                  | 31% ~10s          
  |+++++++++++++++++                                 | 32% ~09s          
  |+++++++++++++++++                                 | 33% ~09s          
  |++++++++++++++++++                                | 34% ~09s          
  |++++++++++++++++++                                | 35% ~09s          
  |+++++++++++++++++++                               | 36% ~09s          
  |+++++++++++++++++++                               | 38% ~09s          
  |++++++++++++++++++++                              | 39% ~08s          
  |++++++++++++++++++++                              | 40% ~08s          
  |+++++++++++++++++++++                             | 41% ~08s          
  |+++++++++++++++++++++                             | 42% ~08s          
  |++++++++++++++++++++++                            | 43% ~08s          
  |++++++++++++++++++++++                            | 44% ~08s          
  |+++++++++++++++++++++++                           | 45% ~07s          
  |+++++++++++++++++++++++                           | 46% ~07s          
  |++++++++++++++++++++++++                          | 47% ~07s          
  |++++++++++++++++++++++++                          | 48% ~07s          
  |+++++++++++++++++++++++++                         | 49% ~07s          
  |+++++++++++++++++++++++++                         | 50% ~07s          
  |++++++++++++++++++++++++++                        | 51% ~07s          
  |+++++++++++++++++++++++++++                       | 52% ~06s          
  |+++++++++++++++++++++++++++                       | 53% ~06s          
  |++++++++++++++++++++++++++++                      | 54% ~06s          
  |++++++++++++++++++++++++++++                      | 55% ~06s          
  |+++++++++++++++++++++++++++++                     | 56% ~06s          
  |+++++++++++++++++++++++++++++                     | 57% ~06s          
  |++++++++++++++++++++++++++++++                    | 58% ~06s          
  |++++++++++++++++++++++++++++++                    | 59% ~05s          
  |+++++++++++++++++++++++++++++++                   | 60% ~05s          
  |+++++++++++++++++++++++++++++++                   | 61% ~05s          
  |++++++++++++++++++++++++++++++++                  | 62% ~05s          
  |++++++++++++++++++++++++++++++++                  | 64% ~05s          
  |+++++++++++++++++++++++++++++++++                 | 65% ~05s          
  |+++++++++++++++++++++++++++++++++                 | 66% ~05s          
  |++++++++++++++++++++++++++++++++++                | 67% ~04s          
  |++++++++++++++++++++++++++++++++++                | 68% ~04s          
  |+++++++++++++++++++++++++++++++++++               | 69% ~04s          
  |+++++++++++++++++++++++++++++++++++               | 70% ~04s          
  |++++++++++++++++++++++++++++++++++++              | 71% ~04s          
  |++++++++++++++++++++++++++++++++++++              | 72% ~04s          
  |+++++++++++++++++++++++++++++++++++++             | 73% ~04s          
  |+++++++++++++++++++++++++++++++++++++             | 74% ~03s          
  |++++++++++++++++++++++++++++++++++++++            | 75% ~03s          
  |+++++++++++++++++++++++++++++++++++++++           | 76% ~03s          
  |+++++++++++++++++++++++++++++++++++++++           | 77% ~03s          
  |++++++++++++++++++++++++++++++++++++++++          | 78% ~03s          
  |++++++++++++++++++++++++++++++++++++++++          | 79% ~03s          
  |+++++++++++++++++++++++++++++++++++++++++         | 80% ~03s          
  |+++++++++++++++++++++++++++++++++++++++++         | 81% ~02s          
  |++++++++++++++++++++++++++++++++++++++++++        | 82% ~02s          
  |++++++++++++++++++++++++++++++++++++++++++        | 83% ~02s          
  |+++++++++++++++++++++++++++++++++++++++++++       | 84% ~02s          
  |+++++++++++++++++++++++++++++++++++++++++++       | 85% ~02s          
  |++++++++++++++++++++++++++++++++++++++++++++      | 86% ~02s          
  |++++++++++++++++++++++++++++++++++++++++++++      | 88% ~02s          
  |+++++++++++++++++++++++++++++++++++++++++++++     | 89% ~02s          
  |+++++++++++++++++++++++++++++++++++++++++++++     | 90% ~01s          
  |++++++++++++++++++++++++++++++++++++++++++++++    | 91% ~01s          
  |++++++++++++++++++++++++++++++++++++++++++++++    | 92% ~01s          
  |+++++++++++++++++++++++++++++++++++++++++++++++   | 93% ~01s          
  |+++++++++++++++++++++++++++++++++++++++++++++++   | 94% ~01s          
  |++++++++++++++++++++++++++++++++++++++++++++++++  | 95% ~01s          
  |++++++++++++++++++++++++++++++++++++++++++++++++  | 96% ~01s          
  |+++++++++++++++++++++++++++++++++++++++++++++++++ | 97% ~00s          
  |+++++++++++++++++++++++++++++++++++++++++++++++++ | 98% ~00s          
  |++++++++++++++++++++++++++++++++++++++++++++++++++| 99% ~00s          
  |++++++++++++++++++++++++++++++++++++++++++++++++++| 100% elapsed=13s
```


```
Calculating cluster IC
```


```
  |                                                  | 0 % ~calculating  
  |+                                                 | 1 % ~01m 40s      
  |++                                                | 2 % ~01m 38s      
  |++                                                | 3 % ~01m 40s      
  |+++                                               | 4 % ~01m 36s      
  |+++                                               | 5 % ~01m 34s      
  |++++                                              | 6 % ~01m 31s      
  |++++                                              | 7 % ~01m 30s      
  |+++++                                             | 8 % ~01m 29s      
  |+++++                                             | 9 % ~01m 27s      
  |++++++                                            | 10% ~01m 26s      
  |++++++                                            | 11% ~01m 25s      
  |+++++++                                           | 12% ~01m 24s      
  |+++++++                                           | 13% ~01m 23s      
  |++++++++                                          | 14% ~01m 22s      
  |++++++++                                          | 15% ~01m 21s      
  |+++++++++                                         | 16% ~01m 20s      
  |+++++++++                                         | 18% ~01m 19s      
  |++++++++++                                        | 19% ~01m 18s      
  |++++++++++                                        | 20% ~01m 17s      
  |+++++++++++                                       | 21% ~01m 16s      
  |+++++++++++                                       | 22% ~01m 15s      
  |++++++++++++                                      | 23% ~01m 14s      
  |++++++++++++                                      | 24% ~01m 13s      
  |+++++++++++++                                     | 25% ~01m 12s      
  |+++++++++++++                                     | 26% ~01m 11s      
  |++++++++++++++                                    | 27% ~01m 10s      
  |++++++++++++++                                    | 28% ~01m 09s      
  |+++++++++++++++                                   | 29% ~01m 08s      
  |+++++++++++++++                                   | 30% ~01m 07s      
  |++++++++++++++++                                  | 31% ~01m 06s      
  |++++++++++++++++                                  | 32% ~01m 05s      
  |+++++++++++++++++                                 | 33% ~01m 04s      
  |++++++++++++++++++                                | 34% ~01m 03s      
  |++++++++++++++++++                                | 35% ~01m 02s      
  |+++++++++++++++++++                               | 36% ~01m 01s      
  |+++++++++++++++++++                               | 37% ~01m 00s      
  |++++++++++++++++++++                              | 38% ~60s          
  |++++++++++++++++++++                              | 39% ~59s          
  |+++++++++++++++++++++                             | 40% ~58s          
  |+++++++++++++++++++++                             | 41% ~57s          
  |++++++++++++++++++++++                            | 42% ~56s          
  |++++++++++++++++++++++                            | 43% ~55s          
  |+++++++++++++++++++++++                           | 44% ~54s          
  |+++++++++++++++++++++++                           | 45% ~53s          
  |++++++++++++++++++++++++                          | 46% ~52s          
  |++++++++++++++++++++++++                          | 47% ~51s          
  |+++++++++++++++++++++++++                         | 48% ~50s          
  |+++++++++++++++++++++++++                         | 49% ~49s          
  |++++++++++++++++++++++++++                        | 51% ~48s          
  |++++++++++++++++++++++++++                        | 52% ~47s          
  |+++++++++++++++++++++++++++                       | 53% ~46s          
  |+++++++++++++++++++++++++++                       | 54% ~45s          
  |++++++++++++++++++++++++++++                      | 55% ~44s          
  |++++++++++++++++++++++++++++                      | 56% ~43s          
  |+++++++++++++++++++++++++++++                     | 57% ~42s          
  |+++++++++++++++++++++++++++++                     | 58% ~41s          
  |++++++++++++++++++++++++++++++                    | 59% ~40s          
  |++++++++++++++++++++++++++++++                    | 60% ~39s          
  |+++++++++++++++++++++++++++++++                   | 61% ~38s          
  |+++++++++++++++++++++++++++++++                   | 62% ~37s          
  |++++++++++++++++++++++++++++++++                  | 63% ~36s          
  |++++++++++++++++++++++++++++++++                  | 64% ~35s          
  |+++++++++++++++++++++++++++++++++                 | 65% ~34s          
  |+++++++++++++++++++++++++++++++++                 | 66% ~33s          
  |++++++++++++++++++++++++++++++++++                | 67% ~32s          
  |+++++++++++++++++++++++++++++++++++               | 68% ~31s          
  |+++++++++++++++++++++++++++++++++++               | 69% ~30s          
  |++++++++++++++++++++++++++++++++++++              | 70% ~29s          
  |++++++++++++++++++++++++++++++++++++              | 71% ~28s          
  |+++++++++++++++++++++++++++++++++++++             | 72% ~27s          
  |+++++++++++++++++++++++++++++++++++++             | 73% ~26s          
  |++++++++++++++++++++++++++++++++++++++            | 74% ~25s          
  |++++++++++++++++++++++++++++++++++++++            | 75% ~24s          
  |+++++++++++++++++++++++++++++++++++++++           | 76% ~23s          
  |+++++++++++++++++++++++++++++++++++++++           | 77% ~22s          
  |++++++++++++++++++++++++++++++++++++++++          | 78% ~21s          
  |++++++++++++++++++++++++++++++++++++++++          | 79% ~20s          
  |+++++++++++++++++++++++++++++++++++++++++         | 80% ~19s          
  |+++++++++++++++++++++++++++++++++++++++++         | 81% ~18s          
  |++++++++++++++++++++++++++++++++++++++++++        | 82% ~17s          
  |++++++++++++++++++++++++++++++++++++++++++        | 84% ~16s          
  |+++++++++++++++++++++++++++++++++++++++++++       | 85% ~15s          
  |+++++++++++++++++++++++++++++++++++++++++++       | 86% ~14s          
  |++++++++++++++++++++++++++++++++++++++++++++      | 87% ~13s          
  |++++++++++++++++++++++++++++++++++++++++++++      | 88% ~12s          
  |+++++++++++++++++++++++++++++++++++++++++++++     | 89% ~11s          
  |+++++++++++++++++++++++++++++++++++++++++++++     | 90% ~10s          
  |++++++++++++++++++++++++++++++++++++++++++++++    | 91% ~09s          
  |++++++++++++++++++++++++++++++++++++++++++++++    | 92% ~08s          
  |+++++++++++++++++++++++++++++++++++++++++++++++   | 93% ~07s          
  |+++++++++++++++++++++++++++++++++++++++++++++++   | 94% ~06s          
  |++++++++++++++++++++++++++++++++++++++++++++++++  | 95% ~05s          
  |++++++++++++++++++++++++++++++++++++++++++++++++  | 96% ~04s          
  |+++++++++++++++++++++++++++++++++++++++++++++++++ | 97% ~03s          
  |+++++++++++++++++++++++++++++++++++++++++++++++++ | 98% ~02s          
  |++++++++++++++++++++++++++++++++++++++++++++++++++| 99% ~01s          
  |++++++++++++++++++++++++++++++++++++++++++++++++++| 100% elapsed=01m 37s
```


```
write.csv(RNA_markers,"/Users/franziskadenk/Library/CloudStorage/OneDrive-SharedLibraries-King'sCollegeLondon/Denk Lab SharePoint - Mesenchymal_MS/cluster_markers.csv")
```


```
Idents(bmc) <- bmc$cluster_IDs
DimPlot(bmc, split.by="Injury", label = TRUE, cols = c("FB_Cldn1" = "#ffffba", "FB_Col15a1" = "#ffb3ba", "FB_Ccl19" = "#ffdfba", "FB_Pi16" = "#d5a6bd", "SC" = "#baffc9", "mySC" = "#f2f6c3", "MC" = "#bae1ff", "EC" = "#cccccc", "IC" = "#559e83"), pt.size=0.7)
```


```
DimPlot(cbmc, split.by="Injury", label=TRUE,  cols = c("5" = "#ffffba", "2" = "#ffb3ba", "0" = "#ffdfba", "4" = "#d5a6bd", "1" = "#baffc9", "7" = "#559e83", "9" = "#f2f6c3", "8" = "#bae1ff", "3" = "blue","6" = "#cccccc", "10" = "#559e83"), pt.size=0.7)
```


```
DimPlot(bmc, split.by="Injury", label = TRUE, cols = c("FB_Cldn1" = "#ffffba", "FB_Col15a1" = "#ffb3ba", "FB_Ccl19" = "#ffdfba", "FB_Pi16" = "#d5a6bd", "SC" = "#baffc9", "mySC" = "#f2f6c3", "MC" = "#bae1ff", "EC" = "#cccccc", "IC" = "#559e83"), pt.size=0.7)
```


```
#plot Pdgfrb, Notch3, Cd146
custom_colours <- c("#DEEDCF", "#99D492","#56B870","#1D9A6C","#137177", "#0A2F51")
feature_plot1 <- FeaturePlot(cbmc, features = "Notch3", min.cutoff = "q05", max.cutoff = "q95", order = TRUE, cols = custom_colours)
feature_plot2 <- FeaturePlot(cbmc, features = "Pdgfrb", min.cutoff = "q05", max.cutoff = "q95", order = TRUE, cols = custom_colours)
feature_plot3 <- FeaturePlot(cbmc, features = "Mcam", min.cutoff = "q05", max.cutoff = "q95", order = TRUE, cols = custom_colours)
dim_plot <- DimPlot(bmc, label=TRUE, label.size = 3, cols = c("FB_Cldn1" = "#ffffba", "FB_Col15a1" = "#ffb3ba", "FB_Ccl19" = "#ffdfba", "FB_Pi16" = "#d5a6bd", "SC" = "#baffc9", "mySC" = "#f2f6c3", "MC" = "#bae1ff", "EC" = "#cccccc", "IC" = "#559e83")) + theme(legend.position = "none")
combined_plot <- (feature_plot1 | feature_plot2 | feature_plot3 | dim_plot) + plot_layout(ncol = 2, nrow = 2)
print(combined_plot)
```


```
#plot known/putative pro-algesic mediators
custom_colours <- c("#DEEDCF", "#99D492","#56B870","#1D9A6C","#137177", "#0A2F51")
feature_plot1 <- FeaturePlot(bmc, features = "Il6", min.cutoff = "q05", max.cutoff = "q95", order = TRUE, cols = custom_colours)
feature_plot2 <- FeaturePlot(bmc, features = "Ngf", min.cutoff = "q05", max.cutoff = "q95", order = TRUE, cols = custom_colours)
feature_plot3 <- FeaturePlot(bmc, features = "Ccl2", min.cutoff = "q05", max.cutoff = "q95", order = TRUE, cols = custom_colours)
feature_plot4 <- FeaturePlot(bmc, features = "Lif", min.cutoff = "q05", max.cutoff = "q95", order = TRUE, cols = custom_colours)
combined_plot <- (feature_plot1 | feature_plot2 | feature_plot3 | feature_plot4) + plot_layout(ncol = 2, nrow = 2)
print(combined_plot)
```


```
bmc@active.ident <- factor(bmc@active.ident, 
                            levels=c("MC", "FB_Ccl19", "FB_Col15a1", "FB_Pi16", "FB_Cldn1", "SC", "mySC", "EC", "IC"))
old_identities <- c("MC", "FB_Ccl19", "FB_Col15a1", "FB_Pi16", "FB_Cldn1", "SC", "mySC", "EC", "IC")
new_identities <- c("Mural Cells", "Fibroblasts-Ccl19", "Fibroblasts-Col15a1", 
                    "Fibroblasts-Pi16", "Fibroblasts-Cldn1", "Schwann Cells", "Myelinating Schwann Cells", 
                    "Endothelial Cells", "Immune Cells")
bmc@active.ident <- factor(bmc@active.ident, levels = old_identities)
identity_map <- setNames(new_identities, old_identities)
bmc@active.ident <- plyr::mapvalues(bmc@active.ident, from = old_identities, to = new_identities)
bmc[["longcluster_IDs"]] <- Idents(object = bmc)
```


```
Idents(bmc) <- bmc$longcluster_IDs
DimPlot(bmc, split.by="Injury", label = TRUE, cols = c("Fibroblasts-Cldn1" = "#ffffba", "Fibroblasts-Col15a1" = "#ffb3ba", "Fibroblasts-Ccl19" = "#ffdfba", "Fibroblasts-Pi16" = "#d5a6bd", "Schwann Cells" = "#baffc9", "Myelinating Schwann Cells" = "#f2f6c3", "Mural Cells" = "#bae1ff", "Endothelial Cells" = "#cccccc", "Immune Cells" = "#559e83"), repel=TRUE, pt.size=0.7) + theme(legend.position = "none")
```


```
Idents(bmc) <- bmc$longcluster_IDs
DimPlot(bmc, split.by="Timepoint", label = TRUE, cols = c("Fibroblasts-Cldn1" = "#ffffba", "Fibroblasts-Col15a1" = "#ffb3ba", "Fibroblasts-Ccl19" = "#ffdfba", "Fibroblasts-Pi16" = "#d5a6bd", "Schwann Cells" = "#baffc9", "Myelinating Schwann Cells" = "#f2f6c3", "Mural Cells" = "#bae1ff", "Endothelial Cells" = "#cccccc", "Immune Cells" = "#559e83"), repel=TRUE, pt.size=0.7) + theme(legend.position = "none")
```


```
Idents(bmc) <- bmc$cluster_IDs
DimPlot(bmc, split.by="sample.ID", cols = c("FB_Cldn1" = "#ffffba", "FB_Col15a1" = "#ffb3ba", "FB_Ccl19" = "#ffdfba", "FB_Pi16" = "#d5a6bd", "SC" = "#baffc9", "mySC" = "#f2f6c3", "MC" = "#bae1ff", "EC" = "#cccccc", "IC" = "#559e83"), pt.size=0.7)
```


```
VlnPlot(bmc, features = c("nFeature_RNA", "nCount_RNA", "percent.mt"), ncol = 3)
```


```
markers.to.plot <- c("Pdgfra", "Pi16", "Col15a1", "Ccl19", "Comp", "Thy1", "Cldn1", "Sfrp5", "Pecam1", "Flt1", "Tagln", "Acta2","Pdgfrb", "Notch3", "Sox10", "Mbp", "Ptprc")
p <- DotPlot(bmc, features = rev(markers.to.plot), cols = c("grey", "darkgreen", "green", "lightgreen"), dot.scale = 8) + RotatedAxis() +
     theme(axis.text.x = element_text(size = 12, angle = 45, hjust = 1)) +
     theme(plot.margin = margin(10, 20, 10, 10))
print(p)
```


```
table(Idents(bmc), bmc$sample.ID)
```


```
             SHAM_D5 SHAM_M2.1 SHAM_M2.2 SNL_D5 SNL_M2.1 SNL_M2.2
  FB_Ccl19       275       222       888    249      879      759
  SC             111        10       748     87      315     1315
  mySC             9         0        13    393       10       11
  MC             159        34       368    169      374      644
  FB_Col15a1     358       126       607     27      171      424
  EC              38         7        59    428       68       85
  FB_Cldn1        79         2       452     30      102      275
  FB_Pi16         65       142       255     14      405      262
  IC               1         0         0     84        1        0
```


save Seurat object


```
save(bmc, file ="/Users/franziskadenk/Library/CloudStorage/OneDrive-SharedLibraries-King'sCollegeLondon/Denk Lab SharePoint - Mesenchymal_MS/scRNA_seq_data/nerve_mesenchymal_cells.Robj")
```


Integrate across batches to check results are equivalent - they
appear to be, at least at macro level.


```
Idents(bmc) <- bmc$batch
combined.list <- SplitObject(bmc, split.by = "batch")
combined.list <- lapply(X = combined.list, FUN = function(x) {
  x <- NormalizeData(x)
  x <- FindVariableFeatures(x, selection.method = "vst", nfeatures = 2000)
})
```


```
Performing log-normalization
0%   10   20   30   40   50   60   70   80   90   100%
[----|----|----|----|----|----|----|----|----|----|
**************************************************|
Calculating gene variances
0%   10   20   30   40   50   60   70   80   90   100%
[----|----|----|----|----|----|----|----|----|----|
**************************************************|
Calculating feature variances of standardized and clipped values
0%   10   20   30   40   50   60   70   80   90   100%
[----|----|----|----|----|----|----|----|----|----|
**************************************************|
Performing log-normalization
0%   10   20   30   40   50   60   70   80   90   100%
[----|----|----|----|----|----|----|----|----|----|
**************************************************|
Calculating gene variances
0%   10   20   30   40   50   60   70   80   90   100%
[----|----|----|----|----|----|----|----|----|----|
**************************************************|
Calculating feature variances of standardized and clipped values
0%   10   20   30   40   50   60   70   80   90   100%
[----|----|----|----|----|----|----|----|----|----|
**************************************************|
```


```
anchors <- FindIntegrationAnchors(object.list = combined.list, dims = 1:10)
```


```
Computing 2000 integration features
Scaling features for provided objects
```


```
  |                                                  | 0 % ~calculating  
  |+++++++++++++++++++++++++                         | 50% ~00s          
  |++++++++++++++++++++++++++++++++++++++++++++++++++| 100% elapsed=01s
```


```
Finding all pairwise anchors
```


```
  |                                                  | 0 % ~calculating
```


```
Running CCA
Merging objects
Finding neighborhoods
Finding anchors
    Found 8493 anchors
Filtering anchors
    Retained 6950 anchors
```


```
  |++++++++++++++++++++++++++++++++++++++++++++++++++| 100% elapsed=02m 17s
```


```
merged.integrated <- IntegrateData(anchorset = anchors, dims = 1:10)
```


```
Merging dataset 2 into 1
Extracting anchors for merged samples
Finding integration vectors
Finding integration vector weights
0%   10   20   30   40   50   60   70   80   90   100%
[----|----|----|----|----|----|----|----|----|----|
**************************************************|
Integrating data
```


```
#integrated analysis
DefaultAssay(merged.integrated) <- "integrated"

# Run the standard workflow for visualization and clustering
integrated <- ScaleData(merged.integrated, verbose = FALSE)
integrated <- RunPCA(integrated, npcs = 30, verbose = FALSE)

# UMAP and Clustering
integrated <- RunUMAP(integrated, reduction = "pca", dims = 1:9)
```


```
16:58:06 UMAP embedding parameters a = 0.9922 b = 1.112
16:58:06 Read 12609 rows and found 9 numeric columns
16:58:06 Using Annoy for neighbor search, n_neighbors = 30
16:58:06 Building Annoy index with metric = cosine, n_trees = 50
0%   10   20   30   40   50   60   70   80   90   100%
[----|----|----|----|----|----|----|----|----|----|
**************************************************|
16:58:07 Writing NN index file to temp file /var/folders/by/d7jw59_d5v32rqlyswwsvx4c0000gn/T//RtmpnrAWww/file12ca96685dc61
16:58:07 Searching Annoy index using 1 thread, search_k = 3000
16:58:11 Annoy recall = 100%
16:58:12 Commencing smooth kNN distance calibration using 1 thread with target n_neighbors = 30
16:58:15 Initializing from normalized Laplacian + noise (using irlba)
16:58:16 Commencing optimization for 200 epochs, with 514610 positive edges
Using method 'umap'
0%   10   20   30   40   50   60   70   80   90   100%
[----|----|----|----|----|----|----|----|----|----|
**************************************************|
16:58:27 Optimization finished
```


```
integrated <- FindNeighbors(integrated, reduction = "pca", dims = 1:9)
```


```
Computing nearest neighbor graph
Computing SNN
```


```
integrated <- FindClusters(integrated, resolution = 0.20)
```


```
Modularity Optimizer version 1.3.0 by Ludo Waltman and Nees Jan van Eck

Number of nodes: 12609
Number of edges: 421906

Running Louvain algorithm...
```


```
0%   10   20   30   40   50   60   70   80   90   100%
[----|----|----|----|----|----|----|----|----|----|
**************************************************|
```


```
Maximum modularity in 10 random starts: 0.9620
Number of communities: 11
Elapsed time: 1 seconds
```


```
DimPlot(integrated, split.by="Injury", label = TRUE, cols = c("5" = "#ffffba", "2" = "#ffb3ba", "0" = "#ffdfba", "3" = "#d5a6bd", "1" = "#baffc9", "7" = "#559e83", "8" = "#f2f6c3", "4" = "#bae1ff", "9" = "blue","6" = "#cccccc", "10" = "#559e83"), pt.size=0.7)
```


```
markers.to.plot <- c("Pi16", "Col15a1", "Ccl19", "Cldn1", "Sfrp5", "Pecam1", "Flt1", "Tagln", "Acta2","Notch3", "Sox10", "Mbp", "Ptprc")
DotPlot(integrated, features = rev(markers.to.plot), cols = c("grey", "darkgreen", "green", "lightgreen"), dot.scale = 8) + RotatedAxis()
```


```
bmc_integrated <- (RenameIdents(integrated, `0` = "FB_Ccl19", `1` = "SC", `9` = "MC", `8` = "mySC", `2` = "FB_Col15a1", `6` = "EC", `5` = "FB_Cldn1",`7` = "SC", `4` = "MC", `3` = "FB_Pi16", `10` = "IC"))
bmc_integrated[["integrated_cluster_IDs"]] <- Idents(object = bmc_integrated)
```


```
DimPlot(bmc_integrated, split.by="Injury", label = TRUE, cols = c("FB_Cldn1" = "#ffffba", "FB_Col15a1" = "#ffb3ba", "FB_Ccl19" = "#ffdfba", "FB_Pi16" = "#d5a6bd", "SC" = "#baffc9", "mySC" = "#f2f6c3", "MC" = "#bae1ff", "EC" = "#cccccc", "IC" = "#559e83"), pt.size=0.7)
```


```
#plot known/putative pro-algesic mediators
custom_colours <- c("#DEEDCF", "#99D492","#56B870","#1D9A6C","#137177", "#0A2F51")
feature_plot1 <- FeaturePlot(bmc_integrated, features = "Il6", min.cutoff = "q05", max.cutoff = "q95", order = TRUE, cols = custom_colours)
feature_plot2 <- FeaturePlot(bmc_integrated, features = "Ngf", min.cutoff = "q05", max.cutoff = "q95", order = TRUE, cols = custom_colours)
feature_plot3 <- FeaturePlot(bmc_integrated, features = "Ccl2", min.cutoff = "q05", max.cutoff = "q95", order = TRUE, cols = custom_colours)
feature_plot4 <- FeaturePlot(bmc_integrated, features = "Pdgfrb", min.cutoff = "q05", max.cutoff = "q95", order = TRUE, cols = custom_colours)
combined_plot <- (feature_plot1 | feature_plot2 | feature_plot3 | feature_plot4) + plot_layout(ncol = 2, nrow = 2)
print(combined_plot)
```


```
#plot known/putative pro-algesic mediators
custom_colours <- c("#DEEDCF", "#99D492","#56B870","#1D9A6C","#137177", "#0A2F51")
feature_plot1 <- FeaturePlot(bmc_integrated, features = "Il6", min.cutoff = "q05", max.cutoff = "q95", order = TRUE, cols = custom_colours)
feature_plot2 <- FeaturePlot(bmc_integrated, features = "Ngf", min.cutoff = "q05", max.cutoff = "q95", order = TRUE, cols = custom_colours)
```


```
Warning: Could not find Ngf in the default search locations, found in RNA assay instead
```


```
feature_plot3 <- FeaturePlot(bmc_integrated, features = "Ccl2", min.cutoff = "q05", max.cutoff = "q95", order = TRUE, cols = custom_colours)
feature_plot4 <- FeaturePlot(bmc_integrated, features = "Pdgfrb", min.cutoff = "q05", max.cutoff = "q95", order = TRUE, cols = custom_colours)
```


```
Warning: Could not find Pdgfrb in the default search locations, found in RNA assay instead
```


```
combined_plot <- (feature_plot1 | feature_plot2 | feature_plot3 | feature_plot4) + plot_layout(ncol = 2, nrow = 2)
print(combined_plot)
```
