## Supplementary R Notebooks for "A role for fibroblast and mural cell subsets in a nerve ligation model of neuropathic pain": Suppl_RNotebook_2.rev.html

scRNA\_seq of nerve mesenchyme


Code 

- Show All Code
- Hide All Code
- Download Rmd

### scRNA\_seq of nerve mesenchyme

This is an R Markdown Notebook. When you execute code within the notebook, the results appear beneath the code.

Try executing this chunk by clicking the *Run* button within the chunk or by placing your cursor inside it and pressing *Ctrl+Shift+Enter*.


```
#If you don't have the following packages download them. Next call them with the library() function
library(Seurat)
library(ggplot2)
library(tidyverse)
library(patchwork)
```


We compared our data to the following other mouse nerve injury datasets: GSE153762, Kalinski et al. ELife 2020 (=Giger); 3 days post sciatic nerve crush GSE120678, Carr et al. Cell Stem Cell 2019 (=Kaplan); 9 days post sciatic nerve crush vs. uninjured GSE142541, Wolbert et al. PNAS 2020; naive sciatic nerve


```
load("D:/King's College London/Denk Lab SharePoint - Mesenchymal_MS/scRNA_seq_data/Giger.Robj")
load("D:/King's College London/Denk Lab SharePoint - Mesenchymal_MS/scRNA_seq_data/Kaplan.Robj")
load("D:/King's College London/Denk Lab SharePoint - Mesenchymal_MS/scRNA_seq_data/Wolbert.Robj")
load("D:/King's College London/Denk Lab SharePoint - Mesenchymal_MS/scRNA_seq_data/nerve_mesenchymal_cells.Robj")
```


alternative loading code for Mac:


```
load("/Users/franziskadenk/Library/CloudStorage/OneDrive-SharedLibraries-King'sCollegeLondon/Denk Lab SharePoint - Mesenchymal_MS/scRNA_seq_data/Giger.Robj")
load("/Users/franziskadenk/Library/CloudStorage/OneDrive-SharedLibraries-King'sCollegeLondon/Denk Lab SharePoint - Mesenchymal_MS/scRNA_seq_data/Kaplan.Robj")
load("/Users/franziskadenk/Library/CloudStorage/OneDrive-SharedLibraries-King'sCollegeLondon/Denk Lab SharePoint - Mesenchymal_MS/scRNA_seq_data/Wolbert.Robj")
load("/Users/franziskadenk/Library/CloudStorage/OneDrive-SharedLibraries-King'sCollegeLondon/Denk Lab SharePoint - Mesenchymal_MS/scRNA_seq_data/nerve_mesenchymal_cells.Robj")
```


```
Giger@meta.data
Kaplan@meta.data
Wolbert@meta.data
```


```
Idents(Wolbert) <- Wolbert$label_new
```


```
There were 18 warnings (use warnings() to see them)
```


```
Giger <- (RenameIdents(Giger, `Mac3` = "Immune", `PC1` = "MC", `EC1` = "EC", `dMes` = "FB", `Hyb` = "?", `Mac4` = "Immune", `PC2` = "MC",`Mac1` = "Immune", `pMes` = "FB", `eMes` = "FB", `SC1` = "SC", `SC2` = "SC",`SC3` = "SC", `EC2` = "EC", `GC` = "Immune", `MoDC` = "Immune", `Mac5` = "Immune",`cDC` = "Immune", `Mo` = "Immune", `Fb` = "?", `T/NK` = "Immune", `EC3` = "EC", `CL` = "?", `Mac2` = "Immune"))
Giger[["FD_clusterID"]] <- Idents(object = Giger)
```


```
VlnPlot(Giger, features = c("nFeature_RNA", "nCount_RNA", "percent.mt"), ncol = 3)
```


```
VlnPlot(Kaplan, features = c("nFeature_RNA", "nCount_RNA", "percent.mt"), ncol = 3)
```


```
VlnPlot(Wolbert, features = c("nFeature_RNA", "nCount_RNA", "percent.mt"), ncol = 3)
```


```
#plot Pdgfrb, Notch3, Cd146
custom_colours <- c("#DEEDCF", "#99D492","#56B870","#1D9A6C","#137177", "#0A2F51")
feature_plot1 <- FeaturePlot(Giger, features = "Notch3", min.cutoff = "q05", max.cutoff = "q95", order = TRUE, cols = custom_colours)
feature_plot2 <- FeaturePlot(Giger, features = "Pdgfrb", min.cutoff = "q05", max.cutoff = "q95", order = TRUE, cols = custom_colours)
feature_plot3 <- FeaturePlot(Giger, features = "Mcam", min.cutoff = "q05", max.cutoff = "q95", order = TRUE, cols = custom_colours)
dim_plot <- DimPlot(Giger, label=TRUE, label.size = 3) + theme(legend.position = "none")
combined_plot <- (feature_plot1 | feature_plot2 | feature_plot3 | dim_plot) + plot_layout(ncol = 2, nrow = 2)
print(combined_plot)
```


```
#plot known/putative pro-algesic mediators
custom_colours <- c("#DEEDCF", "#99D492","#56B870","#1D9A6C","#137177", "#0A2F51")
feature_plot1 <- FeaturePlot(Giger, features = "Il6", min.cutoff = "q05", max.cutoff = "q95", order = TRUE, cols = custom_colours)
feature_plot2 <- FeaturePlot(Giger, features = "Ngf", min.cutoff = "q05", max.cutoff = "q95", order = TRUE, cols = custom_colours)
feature_plot3 <- FeaturePlot(Giger, features = "Ccl2", min.cutoff = "q05", max.cutoff = "q95", order = TRUE, cols = custom_colours)
feature_plot4 <- FeaturePlot(Giger, features = "Lif", min.cutoff = "q05", max.cutoff = "q95", order = TRUE, cols = custom_colours)
combined_plot <- (feature_plot1 | feature_plot2 | feature_plot3 | feature_plot4) + plot_layout(ncol = 2, nrow = 2)
print(combined_plot)
```


```
#plot Pdgfrb, Notch3, Cd146
custom_colours <- c("#DEEDCF", "#99D492","#56B870","#1D9A6C","#137177", "#0A2F51")
feature_plot1 <- FeaturePlot(Kaplan, features = "Notch3", min.cutoff = "q05", max.cutoff = "q95", order = TRUE, cols = custom_colours)
feature_plot2 <- FeaturePlot(Kaplan, features = "Pdgfrb", min.cutoff = "q05", max.cutoff = "q95", order = TRUE, cols = custom_colours)
```


```
Could not find Pdgfrb in the default search locations, found in RNA assay instead
```


```
feature_plot3 <- FeaturePlot(Kaplan, features = "Mcam", min.cutoff = "q05", max.cutoff = "q95", order = TRUE, cols = custom_colours)
dim_plot <- DimPlot(Kaplan, label=TRUE, label.size = 3) + theme(legend.position = "none")
combined_plot <- (feature_plot1 | feature_plot2 | feature_plot3 | dim_plot) + plot_layout(ncol = 2, nrow = 2)
print(combined_plot)
```


```
#plot known/putative pro-algesic mediators
custom_colours <- c("#DEEDCF", "#99D492","#56B870","#1D9A6C","#137177", "#0A2F51")
feature_plot1 <- FeaturePlot(Kaplan, features = "Il6", min.cutoff = "q05", max.cutoff = "q95", order = TRUE, cols = custom_colours)
feature_plot2 <- FeaturePlot(Kaplan, features = "Ngf", min.cutoff = "q05", max.cutoff = "q95", order = TRUE, cols = custom_colours)
```


```
Could not find Ngf in the default search locations, found in RNA assay instead
```


```
feature_plot3 <- FeaturePlot(Kaplan, features = "Ccl2", min.cutoff = "q05", max.cutoff = "q95", order = TRUE, cols = custom_colours)
feature_plot4 <- FeaturePlot(Kaplan, features = "Lif", min.cutoff = "q05", max.cutoff = "q95", order = TRUE, cols = custom_colours)
combined_plot <- (feature_plot1 | feature_plot2 | feature_plot3 | feature_plot4) + plot_layout(ncol = 2, nrow = 2)
print(combined_plot)
```


```
#plot Pdgfrb, Notch3, Cd146
custom_colours <- c("#DEEDCF", "#99D492","#56B870","#1D9A6C","#137177", "#0A2F51")
feature_plot1 <- FeaturePlot(Wolbert, features = "Notch3", order = TRUE, cols = custom_colours, reduction = "umap")
feature_plot2 <- FeaturePlot(Wolbert, features = "Pdgfrb", order = TRUE, cols = custom_colours, reduction = "umap")
feature_plot3 <- FeaturePlot(Wolbert, features = "Mcam", order = TRUE, cols = custom_colours, reduction = "umap")
dim_plot <- DimPlot(Wolbert, label=TRUE, label.size = 3, reduction = "umap") + theme(legend.position = "none")
combined_plot <- (feature_plot1 | feature_plot2 | feature_plot3 | dim_plot) + plot_layout(ncol = 2, nrow = 2)
print(combined_plot)
```


```
#plot known/putative pro-algesic mediators
custom_colours <- c("#DEEDCF", "#99D492","#56B870","#1D9A6C","#137177", "#0A2F51")
feature_plot1 <- FeaturePlot(Wolbert, features = "Il6",order = TRUE, cols = custom_colours, reduction = "umap")
feature_plot2 <- FeaturePlot(Wolbert, features = "Ngf",order = TRUE, cols = custom_colours, reduction = "umap")
feature_plot3 <- FeaturePlot(Wolbert, features = "Ccl2",order = TRUE, cols = custom_colours, reduction = "umap")
feature_plot4 <- FeaturePlot(Wolbert, features = "Lif",order = TRUE, cols = custom_colours, reduction = "umap")
combined_plot <- (feature_plot1 | feature_plot2 | feature_plot3 | feature_plot4) + plot_layout(ncol = 2, nrow = 2)
print(combined_plot)
```


```
#get the percentage of cells within the Pdgfrb and the Pdgfrb/Notch3++ clusters that express the pro-algesic candidates:
PrctCellExpringGene <- function(object, genes, group.by = "all", assay = "RNA", datatype = "counts", threshold = 0){
    if(group.by == "all"){
        prct = unlist(lapply(genes,calc_helper, object=object, assay = assay, datatype=datatype, threshold=threshold))
        result = data.frame(Markers = genes, Cell_proportion = prct)
        return(result)
    }

    else{        
        list = SplitObject(object, group.by)
        factors = names(list)
        results = lapply(list, PrctCellExpringGene, genes=genes, assay = assay, datatype=datatype, threshold=threshold)
        results %>% reduce(full_join, by="Markers") %>% select(any_of("Markers")) -> genelist
        results %>% reduce(full_join, by="Markers") %>% select(!any_of("Markers")) %>% "colnames<-"(names(results)) -> percentages
        combined <- cbind(genelist,percentages)
        return(combined)
    }
}

calc_helper <- function(object,genes,assay,datatype,threshold){
    counts = slot(object[[assay]],datatype)
    ncells = ncol(counts)
    if(genes %in% row.names(counts)){
    round((sum(counts[genes,]>threshold)/ncells)*100,1)
    }else{return(NA)}
}
```


```
PrctCellExpringGene(Kaplan, genes = c("Il6","Ngf","Ccl2","Lif"), 
                    group.by = "FD_clusters", assay = "RNA", datatype = "counts", threshold = 0)
```


```
PrctCellExpringGene(Wolbert, genes = c("Il6","Ngf","Ccl2","Lif"), 
                    group.by = "label_new", assay = "RNA", datatype = "counts", threshold = 0)
```


```
PrctCellExpringGene(bmc, genes = c("Il6","Ngf","Ccl2","Lif"), 
                    group.by = "cluster_IDs", assay = "RNA", datatype = "counts", threshold = 0)
```


```
#export as data.frame
Giger_d <- PrctCellExpringGene(Giger, genes = c("Il6","Ngf","Ccl2","Lif"), 
                    group.by = "FD_clusterID", assay = "RNA", datatype = "counts", threshold = 0)
Kaplan_d <- PrctCellExpringGene(Kaplan, genes = c("Il6","Ngf","Ccl2","Lif"), 
                    group.by = "FD_clusters", assay = "RNA", datatype = "counts", threshold = 0)
Wolbert_d <- PrctCellExpringGene(Wolbert, genes = c("Il6","Ngf","Ccl2","Lif"), 
                    group.by = "label_new", assay = "RNA", datatype = "counts", threshold = 0)
Sara_d <- PrctCellExpringGene(bmc, genes = c("Il6","Ngf","Ccl2","Lif"), 
                    group.by = "cluster_IDs", assay = "RNA", datatype = "counts", threshold = 0)
write.csv(Giger_d, file = "D:/King's College London/Denk Lab SharePoint - Mesenchymal_MS/scRNA_seq_data/Giger.csv")
write.csv(Kaplan_d,file = "D:/King's College London/Denk Lab SharePoint - Mesenchymal_MS/scRNA_seq_data/Kaplan.csv")
write.csv(Wolbert_d,file = "D:/King's College London/Denk Lab SharePoint - Mesenchymal_MS/scRNA_seq_data/Wolbert.csv")
write.csv(Sara_d,file = "D:/King's College London/Denk Lab SharePoint - Mesenchymal_MS/scRNA_seq_data/Sara.csv")
```
