## Supplementary R Notebooks for "A role for fibroblast and mural cell subsets in a nerve ligation model of neuropathic pain": Suppl_RNotebook_3.rev.html

scRNA\_seq of nerve mesenchyme


### scRNA\_seq of nerve mesenchyme

This is an R Markdown
Notebook. When you execute code within the notebook, the results appear
beneath the code.

Try executing this chunk by clicking the *Run* button within
the chunk or by placing your cursor inside it and pressing
*Ctrl+Shift+Enter*.

```
#If you don't have the following packages download them. Next call them with the library() function
library(Seurat)
```

```
## Warning: package 'Seurat' was built under R version 4.1.2
```

```
## Attaching SeuratObject
```

```
library(ggplot2)
library(tidyverse)
```

```
## ── Attaching packages ─────────────────────────────────────── tidyverse 1.3.1 ──
```

```
## ✔ tibble  3.2.1     ✔ dplyr   1.1.2
## ✔ tidyr   1.3.0     ✔ stringr 1.5.1
## ✔ readr   2.0.2     ✔ forcats 0.5.1
## ✔ purrr   1.0.1
```

```
## Warning: package 'tibble' was built under R version 4.1.2
```

```
## Warning: package 'tidyr' was built under R version 4.1.2
```

```
## Warning: package 'purrr' was built under R version 4.1.2
```

```
## Warning: package 'dplyr' was built under R version 4.1.2
```

```
## ── Conflicts ────────────────────────────────────────── tidyverse_conflicts() ──
## ✖ dplyr::filter() masks stats::filter()
## ✖ dplyr::lag()    masks stats::lag()
```

```
library(patchwork)
```

We compared our data to the following other mouse nerve injury
datasets: GSE153762, Kalinski et al. ELife 2020 (=Giger); 3 days post
sciatic nerve crush GSE120678, Carr et al. Cell Stem Cell 2019
(=Kaplan); 9 days post sciatic nerve crush vs. uninjured GSE142541,
Wolbert et al. PNAS 2020; naive sciatic nerve

```
load("/Users/franziskadenk/Library/CloudStorage/OneDrive-SharedLibraries-King'sCollegeLondon/Denk Lab SharePoint - Mesenchymal_MS/scRNA_seq_data/Giger.Robj")
load("/Users/franziskadenk/Library/CloudStorage/OneDrive-SharedLibraries-King'sCollegeLondon/Denk Lab SharePoint - Mesenchymal_MS/scRNA_seq_data/Kaplan.Robj")
load("/Users/franziskadenk/Library/CloudStorage/OneDrive-SharedLibraries-King'sCollegeLondon/Denk Lab SharePoint - Mesenchymal_MS/scRNA_seq_data/Wolbert.Robj")
load("/Users/franziskadenk/Library/CloudStorage/OneDrive-SharedLibraries-King'sCollegeLondon/Denk Lab SharePoint - Mesenchymal_MS/scRNA_seq_data/nerve_mesenchymal_cells.Robj")
```

```
Giger@meta.data
```

```
Kaplan@meta.data
```

```
Wolbert@meta.data
```

```
Idents(Wolbert) <- Wolbert$label_new
```

```
Giger <- (RenameIdents(Giger, `Mac3` = "Immune", `PC1` = "MC", `EC1` = "EC", `dMes` = "FB", `Hyb` = "?", `Mac4` = "Immune", `PC2` = "MC",`Mac1` = "Immune", `pMes` = "FB", `eMes` = "FB", `SC1` = "SC", `SC2` = "SC",`SC3` = "SC", `EC2` = "EC", `GC` = "Immune", `MoDC` = "Immune", `Mac5` = "Immune",`cDC` = "Immune", `Mo` = "Immune", `Fb` = "?", `T/NK` = "Immune", `EC3` = "EC", `CL` = "?", `Mac2` = "Immune"))
Giger[["FD_clusterID"]] <- Idents(object = Giger)
```

```
VlnPlot(Giger, features = c("Acta2", "Kcnj8", "Abcc9", "Vtn", "Ggt1", "Ifitm1"), ncol = 3)
```

```
VlnPlot(Kaplan, features = c("Acta2", "Kcnj8", "Abcc9", "Vtn", "Ggt1", "Ifitm1"), ncol = 3)
```

```
## Warning: Could not find Ggt1 in the default search locations, found in RNA assay
## instead
```

```
VlnPlot(Wolbert, features = c("Acta2", "Kcnj8", "Abcc9", "Vtn", "Ggt1", "Ifitm1"), ncol = 3)
```

```
VlnPlot(bmc, features = c("Acta2", "Kcnj8", "Abcc9", "Vtn", "Ggt1", "Ifitm1"), ncol = 3)
```

```
#plot Acta2, Kcnj8, Abcc9
custom_colours <- c("#DEEDCF", "#99D492","#56B870","#1D9A6C","#137177", "#0A2F51")
feature_plot1 <- FeaturePlot(bmc, features = "Acta2", min.cutoff = "q05", max.cutoff = "q95", order = TRUE, cols = custom_colours)
feature_plot2 <- FeaturePlot(bmc, features = "Kcnj8", min.cutoff = "q05", max.cutoff = "q95", order = TRUE, cols = custom_colours)
feature_plot3 <- FeaturePlot(bmc, features = "Abcc9", min.cutoff = "q05", max.cutoff = "q95", order = TRUE, cols = custom_colours)
dim_plot <- DimPlot(bmc, label=TRUE, label.size = 3) + theme(legend.position = "none")
combined_plot <- (feature_plot1 | feature_plot2 | feature_plot3 | dim_plot) + plot_layout(ncol = 2, nrow = 2)
print(combined_plot)
```

```
#plot pro-algesic mediators
custom_colours <- c("#DEEDCF", "#99D492","#56B870","#1D9A6C","#137177", "#0A2F51")
feature_plot1 <- FeaturePlot(bmc, features = "Il6", min.cutoff = "q05", max.cutoff = "q95", order = TRUE, cols = custom_colours)
feature_plot2 <- FeaturePlot(bmc, features = "Lif", min.cutoff = "q05", max.cutoff = "q95", order = TRUE, cols = custom_colours)
feature_plot3 <- FeaturePlot(bmc, features = "Clcf1", min.cutoff = "q05", max.cutoff = "q95", order = TRUE, cols = custom_colours)
feature_plot4 <- FeaturePlot(bmc, features = "Ngf", min.cutoff = "q05", max.cutoff = "q95", order = TRUE, cols = custom_colours)
combined_plot <- (feature_plot1 | feature_plot2 | feature_plot3 | feature_plot4) + plot_layout(ncol = 2, nrow = 2)
print(combined_plot)
```

```
custom_colours <- c("#DEEDCF", "#99D492","#56B870","#1D9A6C","#137177", "#0A2F51")
feature_plot1 <- FeaturePlot(bmc, features = "Ccl2", min.cutoff = "q05", max.cutoff = "q95", order = TRUE, cols = custom_colours)
feature_plot2 <- FeaturePlot(bmc, features = "Cxcl10", min.cutoff = "q05", max.cutoff = "q95", order = TRUE, cols = custom_colours)
feature_plot3 <- FeaturePlot(bmc, features = "Spp1", min.cutoff = "q05", max.cutoff = "q95", order = TRUE, cols = custom_colours)
feature_plot4 <- FeaturePlot(bmc, features = "Mmp9", min.cutoff = "q05", max.cutoff = "q95", order = TRUE, cols = custom_colours)
combined_plot <- (feature_plot1 | feature_plot2 | feature_plot3 | feature_plot4) + plot_layout(ncol = 2, nrow = 2)
print(combined_plot)
```

```
#plot Acta2, Kcnj8, Abcc9
custom_colours <- c("#DEEDCF", "#99D492","#56B870","#1D9A6C","#137177", "#0A2F51")
feature_plot1 <- FeaturePlot(Giger, features = "Acta2", min.cutoff = "q05", max.cutoff = "q95", order = TRUE, cols = custom_colours)
feature_plot2 <- FeaturePlot(Giger, features = "Kcnj8", min.cutoff = "q05", max.cutoff = "q95", order = TRUE, cols = custom_colours)
feature_plot3 <- FeaturePlot(Giger, features = "Abcc9", min.cutoff = "q05", max.cutoff = "q95", order = TRUE, cols = custom_colours)
dim_plot <- DimPlot(Giger, label=TRUE, label.size = 3) + theme(legend.position = "none")
combined_plot <- (feature_plot1 | feature_plot2 | feature_plot3 | dim_plot) + plot_layout(ncol = 2, nrow = 2)
print(combined_plot)
```

```
#plot #plot Acta2, Kcnj8, Abcc9
custom_colours <- c("#DEEDCF", "#99D492","#56B870","#1D9A6C","#137177", "#0A2F51")
feature_plot1 <- FeaturePlot(Kaplan, features = "Acta2", min.cutoff = "q05", max.cutoff = "q95", order = TRUE, cols = custom_colours)
feature_plot2 <- FeaturePlot(Kaplan, features = "Kcnj8", min.cutoff = "q05", max.cutoff = "q95", order = TRUE, cols = custom_colours)
feature_plot3 <- FeaturePlot(Kaplan, features = "Abcc9", min.cutoff = "q05", max.cutoff = "q95", order = TRUE, cols = custom_colours)
dim_plot <- DimPlot(Kaplan, label=TRUE, label.size = 3) + theme(legend.position = "none")
combined_plot <- (feature_plot1 | feature_plot2 | feature_plot3 | dim_plot) + plot_layout(ncol = 2, nrow = 2)
print(combined_plot)
```

```
#plot Acta2, Kcnj8, Abcc9
custom_colours <- c("#DEEDCF", "#99D492","#56B870","#1D9A6C","#137177", "#0A2F51")
feature_plot1 <- FeaturePlot(Wolbert, features = "Acta2", order = TRUE, cols = custom_colours, reduction = "umap")
feature_plot2 <- FeaturePlot(Wolbert, features = "Kcnj8", order = TRUE, cols = custom_colours, reduction = "umap")
feature_plot3 <- FeaturePlot(Wolbert, features = "Abcc9", order = TRUE, cols = custom_colours, reduction = "umap")
dim_plot <- DimPlot(Wolbert, label=TRUE, label.size = 3, reduction = "umap") + theme(legend.position = "none")
combined_plot <- (feature_plot1 | feature_plot2 | feature_plot3 | dim_plot) + plot_layout(ncol = 2, nrow = 2)
print(combined_plot)
```

```
bmc@meta.data
```

subset just on mural cells

```
Idents(bmc) <- bmc$cluster_IDs
MCs <- subset(bmc, idents="MC")
table(Idents(bmc))
```

```
## 
##   FB_Ccl19         SC       mySC         MC FB_Col15a1         EC   FB_Cldn1 
##       3272       2586        436       1748       1713        685        940 
##    FB_Pi16         IC 
##       1143         86
```

```
MCs
```

```
## An object of class Seurat 
## 55421 features across 1748 samples within 1 assay 
## Active assay: RNA (55421 features, 2000 variable features)
##  2 dimensional reductions calculated: pca, umap
```

recluster based on mural cells

```
RNA_N <- NormalizeData(MCs)
RNA_V <- FindVariableFeatures(RNA_N)
RNA_S <- ScaleData(RNA_V)
```

```
## Centering and scaling data matrix
```

```
PCA <- RunPCA(RNA_S, verbose=TRUE)
```

```
## PC_ 1 
## Positive:  Net1, Fbxl22, Gm37800, Gm37357, Ppp1r14a, Rasd1, Pnpla2, Timp4, Rasl11a, Sh3bgr 
##     Btg2, Nrgn, Plin4, Cnn1, Gm22513, Ppp1r15a, Chchd10, Scn3a, Gadd45g, Klf2 
##     Inhba, Cebpb, Rrad, Gm43123, Actn1, AI593442, Adamts5, Trp53i11, Gja4, Id3 
## Negative:  Col3a1, Col5a1, Timp1, Vcan, Col1a1, Col5a2, Fbn1, Col1a2, Loxl2, Ccnb2 
##     Mki67, Fbln2, Cdca3, Col5a3, Cks2, Fn1, Col6a1, Cthrc1, Emp1, Postn 
##     Marcks, Cenpe, Hmmr, Birc5, Col6a2, Fstl1, Emilin2, Cdc20, Ly6a, Anln 
## PC_ 2 
## Positive:  Vcan, Fbln2, Cthrc1, Lsp1, Dpysl3, Hmmr, Emilin2, Mki67, Ccnb2, Pdpn 
##     Cdc20, Gja1, Cdca3, Anln, Birc5, Pdgfra, Ly6c1, Aspm, Ckap2, Diaph3 
##     Lox, Dcn, Cenpf, Ckap2l, Tacc3, Loxl1, Serpinf1, Adamts5, Cenpe, Gm37357 
## Negative:  Ifitm1, Rgs5, Steap4, Tmem176a, Abcc9, Kcnj8, Tmem176b, Sept4, Colec11, Rgs16 
##     Vtn, Fst, Adra2a, Tmsb4x, Cyp4b1, Sdc1, Meg3, Ccl11, Procr, Adamts12 
##     Phlda1, Cfh, Ifitm3, Cavin2, Rap2a, Id4, Arhgdib, Ccl19, Prrx1, Fhl2 
## PC_ 3 
## Positive:  Mpz, Dcn, Kcna1, Igfbp6, Gatm, Clec3b, Penk, Apod, Serpinf1, Plp1 
##     Mal, Mbp, Apoe, Inmt, Celf2, Igf1, Sfrp4, Fxyd6, Cnp, Lum 
##     Gsn, Ahnak2, Pou3f1, Abca8a, C3, Ptn, Dpep1, Dpt, Fbln1, Entpd2 
## Negative:  Cald1, Actn1, Col18a1, Ppp1r14a, Mcam, Col12a1, Col4a1, Cfl1, Itga1, Col4a2 
##     Tnc, S100a11, Cspg4, Thy1, Calm1, Lrrc32, Tpm4, Nes, Zfp469, Gja4 
##     Ckap4, Fblim1, Pdlim1, AI506816, Ripk3, Hif1a, Cnn1, Pkm, Loxl2, Timp1 
## PC_ 4 
## Positive:  Egr1, Jun, Zfp36, Socs3, Fos, Atf3, Cxcl1, Fst, Vcam1, Aspm 
##     Gm17334, Cxcl12, Mt2, Emilin2, Hmmr, Btg2, Irf1, Ckap2, Gm48942, Ptx3 
##     Fosb, Cenpf, Rgs16, Vcan, Hspa1a, Tpx2, Cdc20, Bub1, Anln, Chodl 
## Negative:  Col18a1, Col4a1, Mbp, Mpz, Col4a2, Mcam, Mal, Gatm, Kcna1, Plp1 
##     Cspg4, Cnp, Vim, Csrp2, Ncmap, Actg2, Cald1, Actn1, Pou3f1, Itga6 
##     Cnn1, Pllp, Itga4, Nes, Cd93, Calm1, Itga1, Emid1, Ccnd1, Itgb8 
## PC_ 5 
## Positive:  Ly6a, Sfrp4, Ly6c1, Clec3b, Mfap5, S100a6, Ccnb2, Igfbp6, Cdkn3, Ifi27l2a 
##     Cdca3, Fbln1, Lgals1, Pcolce, Ifitm3, Serpinf1, Dcn, Cenpa, Hmmr, Spon2 
##     Nbl1, Cks2, C3, Knstrn, Cxcl14, Cdc20, Pi16, Lum, Mmp2, Ifitm1 
## Negative:  Chodl, Myf5, Crlf1, Peg3, Pax7, Sema6a, Fgfr4, Bmp4, Pdlim4, Ncam1 
##     Jsrp1, Cdh15, Tln2, Clcn5, Zim1, Mrln, Clmn, Edn3, Tanc2, Drp2 
##     Vcam1, Pde10a, Prox1, Gm25630, St3gal5, Iqsec3, Egr1, Mt2, Chd7, Lars2
```

```
ElbowPlot(PCA)
```

```
cbmc <- FindNeighbors(PCA, dims = 1:15)
```

```
## Computing nearest neighbor graph
```

```
## Computing SNN
```

```
cbmc <- FindClusters(cbmc, resolution = 0.2)
```

```
## Modularity Optimizer version 1.3.0 by Ludo Waltman and Nees Jan van Eck
## 
## Number of nodes: 1748
## Number of edges: 58480
## 
## Running Louvain algorithm...
## Maximum modularity in 10 random starts: 0.8974
## Number of communities: 6
## Elapsed time: 0 seconds
```

```
cbmc <- RunUMAP(cbmc, dims = 1:15, reduction = "pca")
```

```
## Warning: The default method for RunUMAP has changed from calling Python UMAP via reticulate to the R-native UWOT using the cosine metric
## To use Python UMAP via reticulate, set umap.method to 'umap-learn' and metric to 'correlation'
## This message will be shown once per session
```

```
## 16:43:20 UMAP embedding parameters a = 0.9922 b = 1.112
```

```
## 16:43:20 Read 1748 rows and found 15 numeric columns
```

```
## 16:43:20 Using Annoy for neighbor search, n_neighbors = 30
```

```
## 16:43:20 Building Annoy index with metric = cosine, n_trees = 50
```

```
## 0%   10   20   30   40   50   60   70   80   90   100%
```

```
## [----|----|----|----|----|----|----|----|----|----|
```

```
## **************************************************|
## 16:43:20 Writing NN index file to temp file /var/folders/by/d7jw59_d5v32rqlyswwsvx4c0000gn/T//RtmprjkKO0/file3dde2dfd8f7b
## 16:43:20 Searching Annoy index using 1 thread, search_k = 3000
## 16:43:20 Annoy recall = 100%
## 16:43:21 Commencing smooth kNN distance calibration using 1 thread with target n_neighbors = 30
## 16:43:21 Initializing from normalized Laplacian + noise (using irlba)
## 16:43:21 Commencing optimization for 500 epochs, with 69104 positive edges
## 16:43:24 Optimization finished
```

```
DimPlot(cbmc, label=TRUE, cols = c("4" = "#559e83", "2" = "#ffb3ba", "0" = "#ffdfba", "1" = "#baffc9", "3" = "#bae1ff", "5" = "grey"), pt.size=0.7)
```

```
markers.to.plot <- c("Lama4","Slc1a3","Cldn5", "Adgrf5", "Emcn", "Apln", "Cd82", "Chst1","Cspg4", "Abcc9", "Vtn", "Ptn", "Ctnna3","Slit3","Tagln", "Acta2", "Pdgfrb", "Notch3")
p <- DotPlot(cbmc, features = rev(markers.to.plot), cols = c("grey", "darkgreen", "green", "lightgreen"), dot.scale = 6) +  RotatedAxis() +
     theme(axis.text.x = element_text(size = 12, angle = 45, hjust = 1)) +
     theme(plot.margin = margin(10, 20, 10, 10))
print(p)
```

```
markers.to.plot <- c("Slc12a2", "Slc1a5", "Slc1a3", "Col4a2", "Col4a1", "Lama4","Ptn", "Abcc9", "Vtn", "Ctnna3","Slit3","Tagln", "Acta2", "Pdgfrb", "Notch3")
p <- DotPlot(cbmc, features = rev(markers.to.plot), cols = c("grey", "darkgreen", "green", "lightgreen"), dot.scale = 6) +  RotatedAxis() +
     theme(axis.text.x = element_text(size = 12, angle = 45, hjust = 1)) +
     theme(plot.margin = margin(10, 20, 10, 10))
print(p)
```

```
Idents(cbmc) <- cbmc$seurat_clusters
cbmc <- (RenameIdents(cbmc, `1` = "PC1&2", `2` = "vSMC", `0` = "vSMC", `3` = "PC3", `4` = "aaSMC", `5` = "PC-NC"))
cbmc[["FD_clusters"]] <- Idents(object = cbmc) #to add clusters to metadata.
```

```
MC_markers <- FindAllMarkers(cbmc, min.pct = 0.1, return.thresh = 0.0000012, logfc.threshold = 2)
```

```
## Calculating cluster PC1&2
```

```
## Calculating cluster vSMC
```

```
## Calculating cluster PC3
```

```
## Calculating cluster aaSMC
```

```
## Calculating cluster PC-NC
```

```
write.csv(MC_markers,"/Users/franziskadenk/Library/CloudStorage/OneDrive-SharedLibraries-King'sCollegeLondon/Denk Lab SharePoint - Mesenchymal_MS/scRNA_seq_data/MCs_logFC2.csv")
MC_markers <- FindAllMarkers(cbmc, min.pct = 0.1, return.thresh = 0.0000012)
```

```
## Calculating cluster PC1&2
```

```
## Calculating cluster vSMC
```

```
## Calculating cluster PC3
```

```
## Calculating cluster aaSMC
```

```
## Calculating cluster PC-NC
```

```
write.csv(MC_markers,"/Users/franziskadenk/Library/CloudStorage/OneDrive-SharedLibraries-King'sCollegeLondon/Denk Lab SharePoint - Mesenchymal_MS/scRNA_seq_data/MCs_noFCcut0ff.csv")
```

```
markers.to.plot <- c("Adgrf5","Ptn", "Kcnj8", "Abcc9", "Vtn", "Ctnna3","Slit3","Tagln", "Acta2", "Pdgfrb", "Notch3")
p <- DotPlot(cbmc, features = rev(markers.to.plot), cols = c("grey", "darkgreen", "green", "lightgreen"), dot.scale = 6) +  RotatedAxis() +
     theme(axis.text.x = element_text(size = 12, angle = 45, hjust = 1)) +
     theme(plot.margin = margin(10, 20, 10, 10))
print(p)
```

```
Idents(cbmc) <- cbmc$FD_clusters
custom_colours <- c("#DEEDCF", "#99D492","#56B870","#1D9A6C","#137177", "#0A2F51")
feature_plot1 <- FeaturePlot(cbmc, features = "Notch3", min.cutoff = "q05", max.cutoff = "q95", order = TRUE, cols = custom_colours)
feature_plot2 <- FeaturePlot(cbmc, features = "Tagln", min.cutoff = "q05", max.cutoff = "q95", order = TRUE, cols = custom_colours)
feature_plot3 <- FeaturePlot(cbmc, features = "Acta2", min.cutoff = "q05", max.cutoff = "q95", order = TRUE, cols = custom_colours)
dim_plot <- DimPlot(cbmc, label=TRUE, label.size = 3, cols = c("PC1&2" = "#559e83", "vSMC" = "#ffb3ba", "aaSMC" = "#ffdfba", "PC3" = "#baffc9", "PC-NC" = "#bae1ff")) + theme(legend.position = "none")
combined_plot <- (feature_plot1 | feature_plot2 | feature_plot3 | dim_plot) + plot_layout(ncol = 2, nrow = 2)
print(combined_plot)
```

```
custom_colours <- c("#DEEDCF", "#99D492","#56B870","#1D9A6C","#137177", "#0A2F51")
feature_plot1 <- FeaturePlot(cbmc, features = "Vtn", min.cutoff = "q05", max.cutoff = "q95", order = TRUE, cols = custom_colours)
feature_plot2 <- FeaturePlot(cbmc, features = "Abcc9", min.cutoff = "q05", max.cutoff = "q95", order = TRUE, cols = custom_colours)
feature_plot3 <- FeaturePlot(cbmc, features = "Ptn", min.cutoff = "q05", max.cutoff = "q95", order = TRUE, cols = custom_colours)
feature_plot4 <- FeaturePlot(cbmc, features = "Adgrf5", min.cutoff = "q05", max.cutoff = "q95", order = TRUE, cols = custom_colours)
combined_plot <- (feature_plot1 | feature_plot2 | feature_plot3 | feature_plot4) + plot_layout(ncol = 2, nrow = 2)
print(combined_plot)
```

```
DimPlot(cbmc, label=TRUE, label.size = 5, cols = c("PC1&2" = "#559e83", "vSMC" = "#ffb3ba", "aaSMC" = "#ffdfba", "PC3" = "#baffc9", "PC-NC" = "#bae1ff"),split.by = "Injury") + theme(legend.position = "none")
```

```
DimPlot(cbmc, label=TRUE, label.size = 5, cols = c("PC1&2" = "#559e83", "vSMC" = "#ffb3ba", "aaSMC" = "#ffdfba", "PC3" = "#baffc9", "PC-NC" = "#bae1ff"),split.by = "Timepoint") + theme(legend.position = "none", text = element_text(size=18))
```

```
Idents(cbmc) <- cbmc$FD_clusters
table(Idents(cbmc), cbmc$sample.ID)
```

```
##        
##         SHAM_D5 SHAM_M2.1 SHAM_M2.2 SNL_D5 SNL_M2.1 SNL_M2.2
##   PC1&2      35         6        79    113       82       94
##   vSMC      109        27       231     15      281      495
##   PC3         3         1        58      0        9       35
##   aaSMC      11         0         0     41        0        0
##   PC-NC       1         0         0      0        2       20
```

```
VlnPlot(cbmc, features = c("Il6", "Lif", "Clcf1", "Ccl2", "Ngf", "Tnc"), ncol = 3)
```

```
MC_markers$ratio <- MC_markers$pct.1 / MC_markers$pct.2
MC_markers %>%
    group_by(cluster) %>%
    top_n(n = 5, wt = ratio) -> top5_ratio
MC_markers %>%
    group_by(cluster) %>%
    top_n(n = 5, wt = avg_log2FC) -> top5_FC
```

```
DoHeatmap(cbmc, features = top5_ratio$gene, angle = 45, size = 5) + scale_fill_gradientn(colors = c("purple", "white", "yellow"))+ theme(axis.text.y = element_text(size = 10), legend.text = element_text(size=10), legend.title = element_text(size=10))
```

```
## Warning in DoHeatmap(cbmc, features = top5_ratio$gene, angle = 45, size = 5):
## The following features were omitted as they were not found in the scale.data
## slot for the RNA assay: Gm49975, Lbx1, Dbx1, Vgll2, Traf3ip3, Cd109, Klhl30,
## Gm19705, Gm29686, Kcna6, Tcap, Slc16a11, Gucy2g, Kcnt1
```

```
## Scale for fill is already present.
## Adding another scale for fill, which will replace the existing scale.
```

```
DoHeatmap(cbmc, features = top5_FC$gene, angle = 45, size = 5) + scale_fill_gradientn(colors = c("purple", "white", "yellow"))+ theme(axis.text.y = element_text(size = 10), legend.text = element_text(size=10), legend.title = element_text(size=10))
```

```
## Warning in DoHeatmap(cbmc, features = top5_FC$gene, angle = 45, size = 5): The
## following features were omitted as they were not found in the scale.data slot
## for the RNA assay: Tesc, Myh11, Rcan2, Pcp4l1
```

```
## Scale for fill is already present.
## Adding another scale for fill, which will replace the existing scale.
```
