## Supplementary R Notebooks for "A role for fibroblast and mural cell subsets in a nerve ligation model of neuropathic pain": Suppl_RNotebook_4.rev.html

Deseq2


### Deseq2

```
library(Seurat)
```

```
## Loading required package: SeuratObject
```

```
## Loading required package: sp
```

```
## 'SeuratObject' was built under R 4.4.0 but the current version is
## 4.4.2; it is recomended that you reinstall 'SeuratObject' as the ABI
## for R may have changed
```

```
## 'SeuratObject' was built with package 'Matrix' 1.7.0 but the current
## version is 1.7.1; it is recomended that you reinstall 'SeuratObject' as
## the ABI for 'Matrix' may have changed
```

```
## 
## Attaching package: 'SeuratObject'
```

```
## The following objects are masked from 'package:base':
## 
##     intersect, t
```

```
library(Matrix)
library(tidyverse)
```

```
## ── Attaching core tidyverse packages ──────────────────────── tidyverse 2.0.0 ──
## ✔ dplyr     1.1.4     ✔ readr     2.1.5
## ✔ forcats   1.0.0     ✔ stringr   1.5.1
## ✔ ggplot2   3.5.1     ✔ tibble    3.2.1
## ✔ lubridate 1.9.4     ✔ tidyr     1.3.1
## ✔ purrr     1.0.2
```

```
## ── Conflicts ────────────────────────────────────────── tidyverse_conflicts() ──
## ✖ tidyr::expand() masks Matrix::expand()
## ✖ dplyr::filter() masks stats::filter()
## ✖ dplyr::lag()    masks stats::lag()
## ✖ tidyr::pack()   masks Matrix::pack()
## ✖ tidyr::unpack() masks Matrix::unpack()
## ℹ Use the conflicted package (<http://conflicted.r-lib.org/>) to force all conflicts to become errors
```

```
library(cowplot)
```

```
## 
## Attaching package: 'cowplot'
## 
## The following object is masked from 'package:lubridate':
## 
##     stamp
```

```
library(ggplot2)
library(patchwork)
```

```
## 
## Attaching package: 'patchwork'
## 
## The following object is masked from 'package:cowplot':
## 
##     align_plots
```

```
library(limma)
library(DESeq2)
```

```
## Loading required package: S4Vectors
## Loading required package: stats4
## Loading required package: BiocGenerics
## 
## Attaching package: 'BiocGenerics'
## 
## The following object is masked from 'package:limma':
## 
##     plotMA
## 
## The following objects are masked from 'package:lubridate':
## 
##     intersect, setdiff, union
## 
## The following objects are masked from 'package:dplyr':
## 
##     combine, intersect, setdiff, union
## 
## The following object is masked from 'package:SeuratObject':
## 
##     intersect
## 
## The following objects are masked from 'package:stats':
## 
##     IQR, mad, sd, var, xtabs
## 
## The following objects are masked from 'package:base':
## 
##     anyDuplicated, aperm, append, as.data.frame, basename, cbind,
##     colnames, dirname, do.call, duplicated, eval, evalq, Filter, Find,
##     get, grep, grepl, intersect, is.unsorted, lapply, Map, mapply,
##     match, mget, order, paste, pmax, pmax.int, pmin, pmin.int,
##     Position, rank, rbind, Reduce, rownames, sapply, saveRDS, setdiff,
##     table, tapply, union, unique, unsplit, which.max, which.min
## 
## 
## Attaching package: 'S4Vectors'
## 
## The following objects are masked from 'package:lubridate':
## 
##     second, second<-
## 
## The following objects are masked from 'package:dplyr':
## 
##     first, rename
## 
## The following object is masked from 'package:tidyr':
## 
##     expand
## 
## The following objects are masked from 'package:Matrix':
## 
##     expand, unname
## 
## The following object is masked from 'package:utils':
## 
##     findMatches
## 
## The following objects are masked from 'package:base':
## 
##     expand.grid, I, unname
## 
## Loading required package: IRanges
## 
## Attaching package: 'IRanges'
## 
## The following object is masked from 'package:lubridate':
## 
##     %within%
## 
## The following objects are masked from 'package:dplyr':
## 
##     collapse, desc, slice
## 
## The following object is masked from 'package:purrr':
## 
##     reduce
## 
## The following object is masked from 'package:sp':
## 
##     %over%
## 
## Loading required package: GenomicRanges
## Loading required package: GenomeInfoDb
## Loading required package: SummarizedExperiment
## Loading required package: MatrixGenerics
## Loading required package: matrixStats
## 
## Attaching package: 'matrixStats'
## 
## The following object is masked from 'package:dplyr':
## 
##     count
## 
## 
## Attaching package: 'MatrixGenerics'
## 
## The following objects are masked from 'package:matrixStats':
## 
##     colAlls, colAnyNAs, colAnys, colAvgsPerRowSet, colCollapse,
##     colCounts, colCummaxs, colCummins, colCumprods, colCumsums,
##     colDiffs, colIQRDiffs, colIQRs, colLogSumExps, colMadDiffs,
##     colMads, colMaxs, colMeans2, colMedians, colMins, colOrderStats,
##     colProds, colQuantiles, colRanges, colRanks, colSdDiffs, colSds,
##     colSums2, colTabulates, colVarDiffs, colVars, colWeightedMads,
##     colWeightedMeans, colWeightedMedians, colWeightedSds,
##     colWeightedVars, rowAlls, rowAnyNAs, rowAnys, rowAvgsPerColSet,
##     rowCollapse, rowCounts, rowCummaxs, rowCummins, rowCumprods,
##     rowCumsums, rowDiffs, rowIQRDiffs, rowIQRs, rowLogSumExps,
##     rowMadDiffs, rowMads, rowMaxs, rowMeans2, rowMedians, rowMins,
##     rowOrderStats, rowProds, rowQuantiles, rowRanges, rowRanks,
##     rowSdDiffs, rowSds, rowSums2, rowTabulates, rowVarDiffs, rowVars,
##     rowWeightedMads, rowWeightedMeans, rowWeightedMedians,
##     rowWeightedSds, rowWeightedVars
## 
## Loading required package: Biobase
## Welcome to Bioconductor
## 
##     Vignettes contain introductory material; view with
##     'browseVignettes()'. To cite Bioconductor, see
##     'citation("Biobase")', and for packages 'citation("pkgname")'.
## 
## 
## Attaching package: 'Biobase'
## 
## The following object is masked from 'package:MatrixGenerics':
## 
##     rowMedians
## 
## The following objects are masked from 'package:matrixStats':
## 
##     anyMissing, rowMedians
## 
## 
## Attaching package: 'SummarizedExperiment'
## 
## The following object is masked from 'package:Seurat':
## 
##     Assays
## 
## The following object is masked from 'package:SeuratObject':
## 
##     Assays
```

```
library(tximport)
library(readr)
library(biomaRt)
library(EnhancedVolcano)
```

```
## Loading required package: ggrepel
```

get sample IDs as list

```
base_dir <- "/Users/franziskadenk/Library/CloudStorage//My Drive/Sara_pericytes/output"
folders <- list.dirs(base_dir, recursive = TRUE, full.names = TRUE)
folders <- folders[grepl("CONTRA|IPSI", folders)]
count_subdirectories <- function(path) {
  parts <- unlist(strsplit(path, "/"))
  return(length(parts) - 1)  
}
subdir_counts <- sapply(folders, count_subdirectories)
max_subdirs <- max(subdir_counts)
longer_paths <- folders[subdir_counts == max_subdirs]
files <- file.path(longer_paths, "abundance.tsv")

extract_last_subfolder <- function(path) {
  parts <- unlist(strsplit(path, "/"))
  return(parts[length(parts) - 1])
}
last_subfolders <- sapply(files, extract_last_subfolder)
names(files) <- last_subfolders

all(file.exists(files))
```

```
## [1] TRUE
```

choose one of the abundance.tsv files to use as a reference to obtain
gene symbols.

```
sample <- read.table( "/Users/franziskadenk/Library/CloudStorage//My Drive/Sara_pericytes/output/IPSI_7_F/IPSI_7_F/abundance.tsv" , header=T)
```

```
mart <- useMart(biomart = "ENSEMBL_MART_ENSEMBL", dataset = "mmusculus_gene_ensembl", host = "https://www.ensembl.org/")

ensembl_ids <- sample$target_id
gene_info <- getBM(attributes = c("ensembl_transcript_id_version", "external_gene_name"),
                   filters = "ensembl_transcript_id_version",
                   values = ensembl_ids,
                   mart = mart)
head(gene_info)
```

```
##   ensembl_transcript_id_version external_gene_name
## 1          ENSMUST00000186289.2            Gm29155
## 2          ENSMUST00000185910.2            Gm29155
## 3          ENSMUST00000188753.2            Gm29157
## 4          ENSMUST00000187764.2            Gm29156
## 5         ENSMUST00000061280.17             Pcmtd1
## 6          ENSMUST00000182114.8             Pcmtd1
```

```
txi <- tximport(files, type = "kallisto", tx2gene = gene_info)
```

```
## Note: importing `abundance.h5` is typically faster than `abundance.tsv`
```

```
## reading in files with read_tsv
```

```
## 1 2 3 4 5 6 7 8 9 10 11 12 13 14 15 16 17 18 19 20 21 22 23 24 25 26 27 28 29 30 31 32 33 34 35 36 37 38 39 40 41 42 43 44 45 46 47 48 
## transcripts missing from tx2gene: 12004
## summarizing abundance
## summarizing counts
## summarizing length
```

```
names(txi)
```

```
## [1] "abundance"           "counts"              "length"             
## [4] "countsFromAbundance"
```

```
head(txi$counts)
```

```
##               CONTRA_1_F CONTRA_1_P CONTRA_10_F CONTRA_10_P CONTRA_11_F
##                1191.5619          2   1261.7233    1612.223    1180.629
## 0610005C13Rik     0.0000          0      0.0000       0.000       0.000
## 0610009E02Rik     0.0000          0      0.0000       0.000       0.000
## 0610010K14Rik   500.8661          0    661.4566       0.000     673.679
## 0610012D04Rik     0.0000          0      0.0000       0.000       0.000
## 0610025J13Rik     0.0000          0      0.0000       0.000       0.000
##               CONTRA_11_P CONTRA_12_F CONTRA_12_P CONTRA_13_F CONTRA_13_P
##                   303.802   1558.7900     38.0487   1350.6317    788.7629
## 0610005C13Rik       0.000     52.0526      0.0000      0.0000      0.0000
## 0610009E02Rik       0.000      0.0000      0.0000     17.4659      0.0000
## 0610010K14Rik    2071.002    666.1937      0.0000    697.4358   1342.8240
## 0610012D04Rik       0.000      0.0000      0.0000      0.0000     10.0000
## 0610025J13Rik       0.000      0.0000      0.0000      0.0000      0.0000
##               CONTRA_14_F CONTRA_2_F CONTRA_2_P CONTRA_3_F CONTRA_3_P
##                  1323.367  1233.7048   5.686808   1292.754  1537.7482
## 0610005C13Rik       0.000   124.1380   0.000000      1.000     0.0000
## 0610009E02Rik       0.000    31.5174   0.000000      0.000     0.0000
## 0610010K14Rik    1023.529  1311.3800   0.000000    608.394   319.4802
## 0610012D04Rik       0.000     0.0000   0.000000      0.000     0.0000
## 0610025J13Rik       0.000     0.0000   0.000000      0.000     0.0000
##               CONTRA_4_F CONTRA_4_P CONTRA_5_F CONTRA_5_P CONTRA_6_F CONTRA_6_P
##                1407.5602  867.43054  1729.1254   782.1827   636.6340   112.9856
## 0610005C13Rik     0.0000    0.00000     0.0000     0.0000     0.0000     0.0000
## 0610009E02Rik     0.0000    7.19584    41.2250     0.0000     0.0000     0.0000
## 0610010K14Rik   670.5471  570.25767   912.3052   381.1037   996.6831  1408.8533
## 0610012D04Rik     0.0000    0.00000     0.0000     0.0000     0.0000     0.0000
## 0610025J13Rik     0.0000    0.00000     0.0000     0.0000     0.0000     0.0000
##               CONTRA_7_F CONTRA_7_P CONTRA_8_F CONTRA_8_P CONTRA_9_F CONTRA_9_P
##                1462.1747   12.00603  1575.2218  5402.8420  1186.2733   2067.956
## 0610005C13Rik     0.0000    0.00000     0.0000     0.0000     1.0000      0.000
## 0610009E02Rik     0.0000    0.00000     0.0000     0.0000     0.0000      0.000
## 0610010K14Rik   668.3956  286.51657   825.7082   909.9995   469.3103   1892.377
## 0610012D04Rik     0.0000    0.00000     0.0000     0.0000     0.0000      0.000
## 0610025J13Rik     0.0000    0.00000     0.0000     0.0000     0.0000      0.000
##                IPSI_1_F  IPSI_10_F IPSI_10_P  IPSI_11_F IPSI_11_P  IPSI_12_P
##               1131.6983  825.43814 840.01386 2495.63920  1246.995 1199.64940
## 0610005C13Rik    0.0000    2.00162   2.00112    0.00000     0.000    0.00000
## 0610009E02Rik    0.0000    0.00000   0.00000    9.99906     0.000    7.37671
## 0610010K14Rik  915.2632 1108.55758 905.80373 1172.96161  1385.169 1732.41438
## 0610012D04Rik    9.0000    5.00000  10.00000    5.00000     3.000    6.00000
## 0610025J13Rik    0.0000    0.00000   0.00000    0.00000     0.000    0.00000
##               IPSI_2_F  IPSI_2_P  IPSI_3_F IPSI_3_P  IPSI_4_F IPSI_4_P
##               705.6651  835.6828 2177.6119 1375.877 1790.2635 939.1923
## 0610005C13Rik   0.0000    0.0000    0.0000    2.000    8.0000   0.0000
## 0610009E02Rik   0.0000    0.0000    0.0000    0.000    0.0000  36.6983
## 0610010K14Rik 885.3753 1095.5867  827.9023 1122.985  649.2586 722.9598
## 0610012D04Rik  34.0000   24.0000    0.0000    0.000    0.0000   3.0000
## 0610025J13Rik   0.0000    0.0000    0.0000    0.000    0.0000   0.0000
##                IPSI_5_F  IPSI_5_P IPSI_6_F  IPSI_6_P   IPSI_7_F  IPSI_7_P
##               1268.7518 1110.8727 854.6819  548.9494 1049.05362 1225.3462
## 0610005C13Rik    0.0000    0.0000   0.0000    0.0000    3.00018    1.0000
## 0610009E02Rik    0.0000    0.0000  13.8570    0.0000    0.00000    0.0000
## 0610010K14Rik  737.8657  925.7371 672.6875 1026.0671 1035.25005  655.7962
## 0610012D04Rik   15.0000   13.0000   0.0000    3.0000    0.00000    5.0000
## 0610025J13Rik    0.0000    0.0000   0.0000    0.0000    0.00000    0.0000
##                IPSI_8_F  IPSI_8_P IPSI_9_P
##               1788.4021 1509.0708 1115.998
## 0610005C13Rik    0.0000    0.0000    8.000
## 0610009E02Rik   16.2522    0.0000    0.000
## 0610010K14Rik  733.6623  837.8453  836.137
## 0610012D04Rik    4.0000    0.0000    0.000
## 0610025J13Rik    0.0000    0.0000    0.000
```

```
head(txi$abundance)
```

```
##               CONTRA_1_F CONTRA_1_P CONTRA_10_F CONTRA_10_P CONTRA_11_F
##                 34.48520   0.107728    34.64265    46.92411    28.65205
## 0610005C13Rik    0.00000   0.000000     0.00000     0.00000     0.00000
## 0610009E02Rik    0.00000   0.000000     0.00000     0.00000     0.00000
## 0610010K14Rik   51.84418   0.000000    87.91969     0.00000    71.70154
## 0610012D04Rik    0.00000   0.000000     0.00000     0.00000     0.00000
## 0610025J13Rik    0.00000   0.000000     0.00000     0.00000     0.00000
##               CONTRA_11_P CONTRA_12_F CONTRA_12_P CONTRA_13_F CONTRA_13_P
##                  8.835536    44.58365     4.31106   41.155937    13.24173
## 0610005C13Rik    0.000000     1.39853     0.00000    0.000000     0.00000
## 0610009E02Rik    0.000000     0.00000     0.00000    0.788111     0.00000
## 0610010K14Rik  246.991153    76.44888     0.00000   83.793690   147.61952
## 0610012D04Rik    0.000000     0.00000     0.00000    0.000000     1.87782
## 0610025J13Rik    0.000000     0.00000     0.00000    0.000000     0.00000
##               CONTRA_14_F CONTRA_2_F CONTRA_2_P CONTRA_3_F CONTRA_3_P
##                  24.42459   38.95637  0.3836103 46.7240040   38.12209
## 0610005C13Rik     0.00000    3.24979  0.0000000  0.0335681    0.00000
## 0610009E02Rik     0.00000    1.32145  0.0000000  0.0000000    0.00000
## 0610010K14Rik    89.59890  157.11419  0.0000000 82.4286800   91.74265
## 0610012D04Rik     0.00000    0.00000  0.0000000  0.0000000    0.00000
## 0610025J13Rik     0.00000    0.00000  0.0000000  0.0000000    0.00000
##               CONTRA_4_F CONTRA_4_P CONTRA_5_F CONTRA_5_P CONTRA_6_F CONTRA_6_P
##                 45.92872  39.681986   47.09173   68.45539   46.17392   4.773783
## 0610005C13Rik    0.00000   0.000000    0.00000    0.00000    0.00000   0.000000
## 0610009E02Rik    0.00000   0.406483    1.76790    0.00000    0.00000   0.000000
## 0610010K14Rik   73.26055  86.764156   95.24936   62.33721  121.90331 225.114511
## 0610012D04Rik    0.00000   0.000000    0.00000    0.00000    0.00000   0.000000
## 0610025J13Rik    0.00000   0.000000    0.00000    0.00000    0.00000   0.000000
##               CONTRA_7_F CONTRA_7_P CONTRA_8_F CONTRA_8_P CONTRA_9_F CONTRA_9_P
##                 45.88569   1.120147   69.51964   176.5550 29.7178585    53.2646
## 0610005C13Rik    0.00000   0.000000    0.00000     0.0000  0.0227664     0.0000
## 0610009E02Rik    0.00000   0.000000    0.00000     0.0000  0.0000000     0.0000
## 0610010K14Rik   70.17393  45.634558  129.73161   184.1787 46.7964900   217.0514
## 0610012D04Rik    0.00000   0.000000    0.00000     0.0000  0.0000000     0.0000
## 0610025J13Rik    0.00000   0.000000    0.00000     0.0000  0.0000000     0.0000
##               IPSI_1_F   IPSI_10_F  IPSI_10_P  IPSI_11_F  IPSI_11_P  IPSI_12_P
##               31.57102  18.9409972 23.4975952  56.284747  34.764650  21.947393
## 0610005C13Rik  0.00000   0.0478986  0.0471591   0.000000   0.000000   0.000000
## 0610009E02Rik  0.00000   0.0000000  0.0000000   0.364446   0.000000   0.210845
## 0610010K14Rik 98.17642 119.8464000 92.6918110 118.716690 138.107159 139.982323
## 0610012D04Rik  1.63611   0.8970100  1.7749900   0.848681   0.504354   0.781909
## 0610025J13Rik  0.00000   0.0000000  0.0000000   0.000000   0.000000   0.000000
##                IPSI_2_F  IPSI_2_P IPSI_3_F    IPSI_3_P  IPSI_4_F   IPSI_4_P
##                19.66485  36.37913 66.29663  34.6886263 71.659809  34.762712
## 0610005C13Rik   0.00000   0.00000  0.00000   0.0465815  0.248228   0.000000
## 0610009E02Rik   0.00000   0.00000  0.00000   0.0000000  0.000000   1.931410
## 0610010K14Rik 100.27936 107.51392 97.23347 115.3792930 91.776510 101.874699
## 0610012D04Rik   6.54116   3.83623  0.00000   0.0000000  0.000000   0.736012
## 0610025J13Rik   0.00000   0.00000  0.00000   0.0000000  0.000000   0.000000
##                IPSI_5_F  IPSI_5_P  IPSI_6_F   IPSI_6_P    IPSI_7_F   IPSI_7_P
##                39.48799  34.29478 23.849041  14.684740  41.5697264 49.0410157
## 0610005C13Rik   0.00000   0.00000  0.000000   0.000000   0.0698459  0.0319955
## 0610009E02Rik   0.00000   0.00000  0.619898   0.000000   0.0000000  0.0000000
## 0610010K14Rik 105.24550 113.80877 83.377791 113.007113 104.2406890 95.8648200
## 0610012D04Rik   3.64618   2.60707  0.000000   0.549484   0.0000000  1.1744800
## 0610025J13Rik   0.00000   0.00000  0.000000   0.000000   0.0000000  0.0000000
##                 IPSI_8_F  IPSI_8_P   IPSI_9_P
##                55.401240  42.93683  44.550233
## 0610005C13Rik   0.000000   0.00000   0.244177
## 0610009E02Rik   0.837033   0.00000   0.000000
## 0610010K14Rik 102.745856 107.13457 101.944405
## 0610012D04Rik   0.950310   0.00000   0.000000
## 0610025J13Rik   0.000000   0.00000   0.000000
```

```
header <- read.table("/Users/franziskadenk/Library/CloudStorage/Dropbox/2024/SaraV_pericyte_sequencing/header.txt", header=TRUE)
row.names(header)<- header$Geneid
header$Geneid <- NULL
header
```

```
##              Population Injury Batch
## CONTRA_1_F  fibroblasts contra     1
## CONTRA_1_P    pericytes contra     1
## CONTRA_10_F fibroblasts contra     3
## CONTRA_10_P   pericytes contra     4
## CONTRA_11_F fibroblasts contra     4
## CONTRA_11_P   pericytes contra     4
## CONTRA_12_F fibroblasts contra     4
## CONTRA_12_P   pericytes contra     4
## CONTRA_13_F fibroblasts contra     4
## CONTRA_13_P   pericytes contra     4
## CONTRA_14_F fibroblasts contra     4
## CONTRA_2_F  fibroblasts contra     1
## CONTRA_2_P    pericytes contra     1
## CONTRA_3_F  fibroblasts contra     2
## CONTRA_3_P    pericytes contra     2
## CONTRA_4_F  fibroblasts contra     2
## CONTRA_4_P    pericytes contra     2
## CONTRA_5_F  fibroblasts contra     2
## CONTRA_5_P    pericytes contra     3
## CONTRA_6_F  fibroblasts contra     3
## CONTRA_6_P    pericytes contra     3
## CONTRA_7_F  fibroblasts contra     3
## CONTRA_7_P    pericytes contra     3
## CONTRA_8_F  fibroblasts contra     3
## CONTRA_8_P    pericytes contra     3
## CONTRA_9_F  fibroblasts contra     3
## CONTRA_9_P    pericytes contra     3
## IPSI_1_F    fibroblasts   ipsi     1
## IPSI_10_F   fibroblasts   ipsi     4
## IPSI_10_P     pericytes   ipsi     4
## IPSI_11_F   fibroblasts   ipsi     4
## IPSI_11_P     pericytes   ipsi     4
## IPSI_12_P     pericytes   ipsi     4
## IPSI_2_F    fibroblasts   ipsi     1
## IPSI_2_P      pericytes   ipsi     1
## IPSI_3_F    fibroblasts   ipsi     2
## IPSI_3_P      pericytes   ipsi     2
## IPSI_4_F    fibroblasts   ipsi     2
## IPSI_4_P      pericytes   ipsi     2
## IPSI_5_F    fibroblasts   ipsi     2
## IPSI_5_P      pericytes   ipsi     2
## IPSI_6_F    fibroblasts   ipsi     3
## IPSI_6_P      pericytes   ipsi     3
## IPSI_7_F    fibroblasts   ipsi     3
## IPSI_7_P      pericytes   ipsi     3
## IPSI_8_F    fibroblasts   ipsi     3
## IPSI_8_P      pericytes   ipsi     3
## IPSI_9_P      pericytes   ipsi     4
```

```
dds <- DESeqDataSetFromTximport(txi, header, ~Injury + Population + Injury:Population)
```

```
## Warning in DESeqDataSet(se, design = design, ignoreRank): some variables in
## design formula are characters, converting to factors
```

```
## using counts and average transcript lengths from tximport
```

```
dds
```

```
## class: DESeqDataSet 
## dim: 29732 48 
## metadata(1): version
## assays(2): counts avgTxLength
## rownames(29732): '' 0610005C13Rik ... Zzef1 Zzz3
## rowData names(0):
## colnames(48): CONTRA_1_F CONTRA_1_P ... IPSI_8_P IPSI_9_P
## colData names(3): Population Injury Batch
```

```
rld <- vst(dds)
```

```
## using 'avgTxLength' from assays(dds), correcting for library size
```

```
plotPCA(rld, intgroup=c("Injury","Population","Batch"))
```

```
## using ntop=500 top features by variance
```

```
pcaData <- plotPCA(rld, intgroup = c("Injury", "Population", "Batch"), returnData = TRUE)
```

```
## using ntop=500 top features by variance
```

```
percentVar <- round(100 * attr(pcaData, "percentVar"))
pcaData$Batch <- as.factor(pcaData$Batch)
pcaData$Group <- interaction(pcaData$Injury, pcaData$Population)

custom_colors <- c("contra.fibroblasts" = "#bae1ff", 
                   "ipsi.fibroblasts" = "#000080", 
                   "contra.pericytes" = "#98dd7a", 
                   "ipsi.pericytes" = "#23560c")

#stroke_colors <- c("contra.fibroblasts" = "darkgrey", 
#                   "ipsi.fibroblasts" = "lightgrey", 
#                 "contra.pericytes" = "darkgrey", 
#                  "ipsi.pericytes" = "darkgrey")

# Create the PCA plot using ggplot2
ggplot(pcaData, aes(x = PC1, y = PC2, shape = Batch, colour = Group)) +
  geom_point(size = 15, stroke = 3) +  
  scale_shape_manual(values = c(21, 22, 23, 24)) +  
  scale_color_manual(values = custom_colors) +  
  #scale_fill_manual(values = custom_colors) +  
  xlab(paste0("PC1: ", percentVar[1], "% variance")) +
  ylab(paste0("PC2: ", percentVar[2], "% variance")) +
  theme_bw() +  
  theme(
    axis.title = element_text(size = 40, face = "bold", family = "Arial"),  
    axis.text = element_text(size = 35, family = "Arial"),  
    legend.title = element_text(size = 40, family = "Arial", face = "bold"),  
    legend.text = element_text(size = 35, family = "Arial"),   
    legend.key.size = unit(1.5, "lines"),
    legend.key.height = unit(2, "cm"), 
    panel.grid.major = element_line(color = "grey", linetype = "dotted"),  
    panel.grid.minor = element_blank(),  
  ) +
  labs(fill = "Group", shape = "Batch")
```

keep only genes with raw counts of at least 50 in at least 10 samples in
any of the four main experimental groups.

```
dds$Group <- factor(paste(dds$Injury, dds$Population, sep = "_"))
groups <- levels(dds$Group)

filter_counts <- function(counts, threshold, min_samples) {
  rowSums(counts >= threshold) >= min_samples
}

keep_genes_matrix <- matrix(FALSE, nrow = nrow(dds), ncol = length(groups))

for (i in seq_along(groups)) {
  group <- groups[i]
  # Get the counts for the current group
  group_samples <- which(dds$Group == group)
  group_counts <- counts(dds)[, group_samples]
    keep_genes_matrix[, i] <- filter_counts(group_counts, threshold = 50, min_samples = 10)
}

keep_genes <- rowSums(keep_genes_matrix) > 0

dds_filtered <- dds[keep_genes, ]
dim(dds_filtered)
```

```
## [1] 10713    48
```

```
design(dds) <- ~Population
celltype <- DESeq(dds)
```

```
## estimating size factors
```

```
## using 'avgTxLength' from assays(dds), correcting for library size
```

```
## estimating dispersions
```

```
## gene-wise dispersion estimates
```

```
## mean-dispersion relationship
```

```
## final dispersion estimates
```

```
## fitting model and testing
```

```
## -- replacing outliers and refitting for 6593 genes
## -- DESeq argument 'minReplicatesForReplace' = 7 
## -- original counts are preserved in counts(dds)
```

```
## estimating dispersions
```

```
## fitting model and testing
```

```
resultsNames(celltype)
```

```
## [1] "Intercept"                           "Population_pericytes_vs_fibroblasts"
```

```
res_celltype <- results(celltype, name="Population_pericytes_vs_fibroblasts")
summary(res_celltype)
```

```
## 
## out of 21705 with nonzero total read count
## adjusted p-value < 0.1
## LFC > 0 (up)       : 953, 4.4%
## LFC < 0 (down)     : 1007, 4.6%
## outliers [1]       : 0, 0%
## low counts [2]     : 5952, 27%
## (mean count < 1)
## [1] see 'cooksCutoff' argument of ?results
## [2] see 'independentFiltering' argument of ?results
```

```
resOrdered <- res_celltype[order(res_celltype$padj),]
write.csv(resOrdered,file="/Users/franziskadenk/Library/CloudStorage/OneDrive-SharedLibraries-King'sCollegeLondon/Denk Lab SharePoint - Mesenchymal_MS/Bulk seq PGOA TdTom/res_celltype.csv")
```

```
colCustom = c("grey", "yellow", "darkblue", "lightblue")
EnhancedVolcano(res_celltype,
    lab = rownames(res_celltype),
    x = 'log2FoldChange',
    y = 'pvalue',
    legendLabels=c('Not sig.','logFC>1','adj.p<0.05',
      'logFC>1 & adj.p<0.05'),
    selectLab = c('Notch3',"Pdgfrb", "Mcam", "Pdgfra", "Pdpn", "Pi16"),
    pCutoff = 0.05,
    pCutoffCol = 'padj',
    FCcutoff = 1,
    col = colCustom,
    pointSize = 10.0,
    labSize = 10, 
    labCol = c('darkblue','darkblue','darkgreen','darkblue', 'darkgreen', 'darkgreen'),
    labFace = 'bold',
    boxedLabels = TRUE,
    drawConnectors = TRUE,
    widthConnectors = 0.8,
    colConnectors = 'darkblue',
    legendLabSize = 30, 
    axisLabSize = 22)
```

subset into different populations:

```
fib_dds <- dds_filtered[, (dds$Population == "fibroblasts")]
per_dds <- dds_filtered[, (dds$Population == "pericytes")]
design(fib_dds) <- ~Injury
design(per_dds) <- ~Injury
```

```
main <- DESeq(fib_dds)
```

```
## estimating size factors
```

```
## using 'avgTxLength' from assays(dds), correcting for library size
```

```
## estimating dispersions
```

```
## gene-wise dispersion estimates
```

```
## mean-dispersion relationship
```

```
## final dispersion estimates
```

```
## fitting model and testing
```

```
## -- replacing outliers and refitting for 45 genes
## -- DESeq argument 'minReplicatesForReplace' = 7 
## -- original counts are preserved in counts(dds)
```

```
## estimating dispersions
```

```
## fitting model and testing
```

```
resultsNames(main)
```

```
## [1] "Intercept"             "Injury_ipsi_vs_contra"
```

```
res <- results(main, name="Injury_ipsi_vs_contra")
summary(res)
```

```
## 
## out of 10713 with nonzero total read count
## adjusted p-value < 0.1
## LFC > 0 (up)       : 1518, 14%
## LFC < 0 (down)     : 1714, 16%
## outliers [1]       : 0, 0%
## low counts [2]     : 0, 0%
## (mean count < 0)
## [1] see 'cooksCutoff' argument of ?results
## [2] see 'independentFiltering' argument of ?results
```

```
resOrdered <- res[order(res$padj),]
write.csv(resOrdered,file="/Users/franziskadenk/Library/CloudStorage/OneDrive-SharedLibraries-King'sCollegeLondon/Denk Lab SharePoint - Mesenchymal_MS/Bulk seq PGOA TdTom/fibroblasts.csv")
```

```
filtered_res <- res[res$log2FoldChange > 3 & res$padj < 0.05,]
dim(filtered_res)
```

```
## [1] 200   6
```

```
colCustom = c("grey", "yellow", "darkblue", "lightblue")
EnhancedVolcano(res,
    lab = rownames(res),
    x = 'log2FoldChange',
    y = 'pvalue',
    title = "", subtitle = 'Pdgfrb+ fibroblasts', subtitleLabSize = 30,
    legendLabels=c('Not sig.','logFC>3','adj.p<0.05',
      'logFC>3 & adj.p<0.05'),
    selectLab = c('Birc5',"Kif20a", "Ccnb1", "Il6", "Tnc", "Mmp9", "Spp1", "Ptn", "Cxcl10", "Ccl8", "Vcan", "Calm", "Ccna2"),
    pCutoff = 0.05,
    pCutoffCol = 'padj',
    FCcutoff = 3,
    col = colCustom,
    pointSize = 10.0,
    labSize = 10.0, 
    labCol = 'darkblue',
    labFace = 'bold',
    boxedLabels = TRUE,
    drawConnectors = TRUE,
    widthConnectors = 0.8,
    colConnectors = 'darkblue', legendLabSize = 30,
    axisLabSize = 22) + theme(plot.subtitle = element_text(face = "bold"))
```

```
main_p <- DESeq(per_dds)
```

```
## estimating size factors
```

```
## using 'avgTxLength' from assays(dds), correcting for library size
```

```
## estimating dispersions
```

```
## gene-wise dispersion estimates
```

```
## mean-dispersion relationship
```

```
## final dispersion estimates
```

```
## fitting model and testing
```

```
## -- replacing outliers and refitting for 4031 genes
## -- DESeq argument 'minReplicatesForReplace' = 7 
## -- original counts are preserved in counts(dds)
```

```
## estimating dispersions
```

```
## fitting model and testing
```

```
resultsNames(main_p)
```

```
## [1] "Intercept"             "Injury_ipsi_vs_contra"
```

```
res_p <- results(main_p, name="Injury_ipsi_vs_contra")
summary(res_p)
```

```
## 
## out of 10712 with nonzero total read count
## adjusted p-value < 0.1
## LFC > 0 (up)       : 1582, 15%
## LFC < 0 (down)     : 700, 6.5%
## outliers [1]       : 0, 0%
## low counts [2]     : 1, 0.0093%
## (mean count < 0)
## [1] see 'cooksCutoff' argument of ?results
## [2] see 'independentFiltering' argument of ?results
```

```
resOrdered <- res_p[order(res$padj),]
write.csv(resOrdered,file="/Users/franziskadenk/Library/CloudStorage/OneDrive-SharedLibraries-King'sCollegeLondon/Denk Lab SharePoint - Mesenchymal_MS/Bulk seq PGOA TdTom/pericytes.csv")
```

```
filtered_res <- res_p[res_p$log2FoldChange > 3 & res_p$padj < 0.05,]
dim(filtered_res)
```

```
## [1] 1184    6
```

```
colCustom = c("grey", "yellow", "darkblue", "lightblue")
EnhancedVolcano(res_p,
    lab = rownames(res_p),
    x = 'log2FoldChange',
    y = 'pvalue',
    title = "", subtitle = 'Pdgfrb+Cd146+ mural cells', subtitleLabSize = 30,
    legendLabels=c('Not sig.','logFC>3','adj.p<0.05',
      'logFC>3 & adj.p<0.05'),
    selectLab = c('Ccnb2',"Kif22", "Ckap2", "Il6", "Ccl2", "Lif", "Clcf1", "Tnc", "Cxcl10", "Actg2", "Adam12", "Ptn"),
    pCutoff = 0.05,
    pCutoffCol = 'padj',
    FCcutoff = 3,
    col = colCustom,
    pointSize = 10.0,
    labSize = 10.0, 
    labCol = 'darkblue',
    labFace = 'bold',
    boxedLabels = TRUE,
    drawConnectors = TRUE,
    widthConnectors = 0.8,
    colConnectors = 'darkblue', legendLabSize = 30, 
    axisLabSize = 22) + theme(plot.subtitle = element_text(face = "bold"))
```

```
all <- as.data.frame(counts(dds_filtered))
pericytes <- as.data.frame(counts(per_dds))
fibroblasts <- as.data.frame(counts(fib_dds))
head(pericytes)
```

```
##               CONTRA_1_P CONTRA_10_P CONTRA_11_P CONTRA_12_P CONTRA_13_P
##                        2        1612         304          38         789
## 0610010K14Rik          0           0        2071           0        1343
## 0610030E20Rik          1           0           0           0           0
## 1110004F10Rik          1        2476        1650           0        7231
## 1110038F14Rik          1           0        3174        1534        3033
## 1110059E24Rik          0        1588           3           0        2747
##               CONTRA_2_P CONTRA_3_P CONTRA_4_P CONTRA_5_P CONTRA_6_P CONTRA_7_P
##                        6       1538        867        782        113         12
## 0610010K14Rik          0        319        570        381       1409        287
## 0610030E20Rik          0          3        481        947         96          0
## 1110004F10Rik       4706       3468       2865       5426       6821         35
## 1110038F14Rik       2895       1121        889       1629       1263          7
## 1110059E24Rik       3555          3       1476       1613       2000       1945
##               CONTRA_8_P CONTRA_9_P IPSI_10_P IPSI_11_P IPSI_12_P IPSI_2_P
##                     5403       2068       840      1247      1200      836
## 0610010K14Rik        910       1892       906      1385      1732     1096
## 0610030E20Rik          0          0       332       148       290      200
## 1110004F10Rik       3553       3096      5717      4367      7404     6278
## 1110038F14Rik        318       1091       871      1197      1522     1206
## 1110059E24Rik          0          1      2627      2243      2368     2915
##               IPSI_3_P IPSI_4_P IPSI_5_P IPSI_6_P IPSI_7_P IPSI_8_P IPSI_9_P
##                   1376      939     1111      549     1225     1509     1116
## 0610010K14Rik     1123      723      926     1026      656      838      836
## 0610030E20Rik      371      479      680      500      342      502      602
## 1110004F10Rik     5100     2945     3594     4466     2503     3760     3112
## 1110038F14Rik      950      966      885     1120      779      743      950
## 1110059E24Rik     1688      668     1660     1834      825     1857      424
```

```
head(fibroblasts)
```

```
##               CONTRA_1_F CONTRA_10_F CONTRA_11_F CONTRA_12_F CONTRA_13_F
##                     1192        1262        1181        1559        1351
## 0610010K14Rik        501         661         674         666         697
## 0610030E20Rik        419         400         373         572         461
## 1110004F10Rik       4863        3983        4163        4264        3317
## 1110038F14Rik        879         822        1774        1472         790
## 1110059E24Rik       1561        1042        1484         865        1002
##               CONTRA_14_F CONTRA_2_F CONTRA_3_F CONTRA_4_F CONTRA_5_F
##                      1323       1234       1293       1408       1729
## 0610010K14Rik        1024       1311        608        671        912
## 0610030E20Rik         495        274        475        595        627
## 1110004F10Rik        6702       3538       3580       3811       3089
## 1110038F14Rik         815        660        623        622       1530
## 1110059E24Rik        1819       2305        883        678        503
##               CONTRA_6_F CONTRA_7_F CONTRA_8_F CONTRA_9_F IPSI_1_F IPSI_10_F
##                      637       1462       1575       1186     1132       825
## 0610010K14Rik        997        668        826        469      915      1109
## 0610030E20Rik        355        411        467        554      358       220
## 1110004F10Rik       3323       4700       3885       2906     3412      4980
## 1110038F14Rik       1300        970        634       1698      986       971
## 1110059E24Rik       1262       1715       1471       1515     1056      1471
##               IPSI_11_F IPSI_2_F IPSI_3_F IPSI_4_F IPSI_5_F IPSI_6_F IPSI_7_F
##                    2496      706     2178     1790     1269      855     1049
## 0610010K14Rik      1173      885      828      649      738      673     1035
## 0610030E20Rik       551      133      203      340      341      266      283
## 1110004F10Rik      3982     4984     4073     3135     2935     3294     6417
## 1110038F14Rik      1022      977     1019      581      535      791     1052
## 1110059E24Rik      1547     1880     1253      855      975     1766     2091
##               IPSI_8_F
##                   1788
## 0610010K14Rik      734
## 0610030E20Rik      655
## 1110004F10Rik     2641
## 1110038F14Rik      752
## 1110059E24Rik      991
```

```
library(icellnet)
```

```
##
```

```
##
```

Expression profile of mouse DRG neurons pre and post nerve injury
(SNI) derived from this paper: https://pubmed.ncbi.nlm.nih.gov/32810432/

```
partner=read.table("/Users/franziskadenk/Library/CloudStorage/Dropbox/2021/publications/Paolo&Francesca/Izzi_et_al_to_share/Renthal_gene_sums.txt", header=T)
head(partner)
```

```
##               NF_uninjured_3 NP_uninjured_3 PEP_uninjured_3 cLTMR_uninjured_3
## 0610005C13Rik     564.330107     2384.93990      2484.45273        1563.13599
## 0610006L08Rik       8.606382       39.35442        15.05586          14.32267
## 0610007P14Rik   10488.823627     7779.80402      8476.93727        2875.34192
## 0610009B22Rik    8775.650566     3662.11237      3842.09295        1813.56128
## 0610009E02Rik     389.134830      566.66555       477.43232         340.68369
## 0610009L18Rik     320.478902      118.97148       227.91330          61.90728
##               SST_uninjured_3 NF_injured_3 NP_injured_3 PEP_injured_3
## 0610005C13Rik      727.617002     192.8315   490.142926    652.691926
## 0610006L08Rik        5.492652       6.4675     1.543927      6.425388
## 0610007P14Rik     1810.823621    3574.0508  2984.052244   1705.842108
## 0610009B22Rik      921.780785    1734.5875   808.832457    490.937240
## 0610009E02Rik      167.252261     147.6948   140.569797     89.753720
## 0610009L18Rik       93.724699     139.3107   114.845178     53.605901
##               cLTMR_injured_3 SST_injured_3
## 0610005C13Rik       434.37349   114.9303042
## 0610006L08Rik         0.00000     0.8562377
## 0610007P14Rik      1482.84522   254.1778330
## 0610009B22Rik       321.03245    73.5717841
## 0610009E02Rik        96.64204    28.7685600
## 0610009L18Rik        40.30028    23.6580951
```

```
data <- merge(all, partner, by = "row.names", all = FALSE)
rownames(data) <- data$Row.names
data <- data[, -1]
head(data)
```

```
##               CONTRA_1_F CONTRA_1_P CONTRA_10_F CONTRA_10_P CONTRA_11_F
## 0610010K14Rik        501          0         661           0         674
## 0610030E20Rik        419          1         400           0         373
## 1110004F10Rik       4863          1        3983        2476        4163
## 1110038F14Rik        879          1         822           0        1774
## 1110059E24Rik       1561          0        1042        1588        1484
## 1110059G10Rik       1013          2         585         931         532
##               CONTRA_11_P CONTRA_12_F CONTRA_12_P CONTRA_13_F CONTRA_13_P
## 0610010K14Rik        2071         666           0         697        1343
## 0610030E20Rik           0         572           0         461           0
## 1110004F10Rik        1650        4264           0        3317        7231
## 1110038F14Rik        3174        1472        1534         790        3033
## 1110059E24Rik           3         865           0        1002        2747
## 1110059G10Rik           2         634           0         668         891
##               CONTRA_14_F CONTRA_2_F CONTRA_2_P CONTRA_3_F CONTRA_3_P
## 0610010K14Rik        1024       1311          0        608        319
## 0610030E20Rik         495        274          0        475          3
## 1110004F10Rik        6702       3538       4706       3580       3468
## 1110038F14Rik         815        660       2895        623       1121
## 1110059E24Rik        1819       2305       3555        883          3
## 1110059G10Rik        1144        646          0        544       1582
##               CONTRA_4_F CONTRA_4_P CONTRA_5_F CONTRA_5_P CONTRA_6_F CONTRA_6_P
## 0610010K14Rik        671        570        912        381        997       1409
## 0610030E20Rik        595        481        627        947        355         96
## 1110004F10Rik       3811       2865       3089       5426       3323       6821
## 1110038F14Rik        622        889       1530       1629       1300       1263
## 1110059E24Rik        678       1476        503       1613       1262       2000
## 1110059G10Rik        894        812        647       1187        825          1
##               CONTRA_7_F CONTRA_7_P CONTRA_8_F CONTRA_8_P CONTRA_9_F CONTRA_9_P
## 0610010K14Rik        668        287        826        910        469       1892
## 0610030E20Rik        411          0        467          0        554          0
## 1110004F10Rik       4700         35       3885       3553       2906       3096
## 1110038F14Rik        970          7        634        318       1698       1091
## 1110059E24Rik       1715       1945       1471          0       1515          1
## 1110059G10Rik       1068          2        449        499        624          0
##               IPSI_1_F IPSI_10_F IPSI_10_P IPSI_11_F IPSI_11_P IPSI_12_P
## 0610010K14Rik      915      1109       906      1173      1385      1732
## 0610030E20Rik      358       220       332       551       148       290
## 1110004F10Rik     3412      4980      5717      3982      4367      7404
## 1110038F14Rik      986       971       871      1022      1197      1522
## 1110059E24Rik     1056      1471      2627      1547      2243      2368
## 1110059G10Rik      465       549       673       805       880      1268
##               IPSI_2_F IPSI_2_P IPSI_3_F IPSI_3_P IPSI_4_F IPSI_4_P IPSI_5_F
## 0610010K14Rik      885     1096      828     1123      649      723      738
## 0610030E20Rik      133      200      203      371      340      479      341
## 1110004F10Rik     4984     6278     4073     5100     3135     2945     2935
## 1110038F14Rik      977     1206     1019      950      581      966      535
## 1110059E24Rik     1880     2915     1253     1688      855      668      975
## 1110059G10Rik      694      802      821      754      501      513      605
##               IPSI_5_P IPSI_6_F IPSI_6_P IPSI_7_F IPSI_7_P IPSI_8_F IPSI_8_P
## 0610010K14Rik      926      673     1026     1035      656      734      838
## 0610030E20Rik      680      266      500      283      342      655      502
## 1110004F10Rik     3594     3294     4466     6417     2503     2641     3760
## 1110038F14Rik      885      791     1120     1052      779      752      743
## 1110059E24Rik     1660     1766     1834     2091      825      991     1857
## 1110059G10Rik      732      612      686      390      696      407      702
##               IPSI_9_P NF_uninjured_3 NP_uninjured_3 PEP_uninjured_3
## 0610010K14Rik      836       177.8387        148.961        103.5998
## 0610030E20Rik      602      1366.1238       1981.831       2338.3739
## 1110004F10Rik     3112     48650.0071      18609.040      21980.2573
## 1110038F14Rik      950      3237.9870       1611.138       1819.1247
## 1110059E24Rik      424      6324.0156       2529.927       1974.9727
## 1110059G10Rik      460      6142.6852       3014.668       3535.7564
##               cLTMR_uninjured_3 SST_uninjured_3 NF_injured_3 NP_injured_3
## 0610010K14Rik          60.05918        9.170503     63.94313     71.39749
## 0610030E20Rik        1190.01021      656.564118    432.91717    612.03850
## 1110004F10Rik        8651.60130     5810.045548  20561.45268  10291.53379
## 1110038F14Rik         824.34358      442.010220   1050.64258    793.84248
## 1110059E24Rik        1210.24574      738.002726   2279.96667    714.00843
## 1110059G10Rik        1239.08118      902.162791   1701.48475    977.64289
##               PEP_injured_3 cLTMR_injured_3 SST_injured_3
## 0610010K14Rik      30.98715        41.75452      26.24068
## 0610030E20Rik     644.88474       531.64448      54.82659
## 1110004F10Rik    5995.50663      4157.84016    1100.74119
## 1110038F14Rik     485.35180       316.78778      85.36186
## 1110059E24Rik     383.10842       297.55592      67.03274
## 1110059G10Rik     769.47448       422.03457     112.35739
```

```
meta=read.table("/Users/franziskadenk/Library/CloudStorage/OneDrive-SharedLibraries-King'sCollegeLondon/Denk Lab SharePoint - Mesenchymal_MS/Bulk seq PGOA TdTom/meta_data.txt", header=T)
head(meta)
```

```
##            ID   Cell_type  Class
## 1  CONTRA_1_F fibroblasts contra
## 2  CONTRA_1_P   pericytes contra
## 3 CONTRA_10_F fibroblasts contra
## 4 CONTRA_10_P   pericytes contra
## 5 CONTRA_11_F fibroblasts contra
## 6 CONTRA_11_P   pericytes contra
```

```
db=as.data.frame(read.csv(curl::curl(url="https://raw.githubusercontent.com/soumelis-lab/ICELLNET/master/data/ICELLNETdb.tsv"), sep="\t",header = T, check.names=FALSE, stringsAsFactors = FALSE, na.strings = ""))
db.name.couple=name.lr.couple(db, type="Family")
head(db.name.couple)
```

```
##      Pair                     Family         
## [1,] "TIMP1 + MMP9 / LRP1"    NA             
## [2,] "LTB + LTA / LTBR"       "Cytokine"     
## [3,] "ITGAV + ITGB5 / ADGRB1" "Cell adhesion"
## [4,] "ITGAV + ITGB3 / ADGRA2" "Cell adhesion"
## [5,] "ITGAV + ITGB3 / ADGRE5" "Cell adhesion"
## [6,] "ITGA5 + ITGB1 / ADGRE5" "Cell adhesion"
```

iCellnet is designed to work with human data only. We are therefore
converting the mouse IDs we have obtained to human orthologues.

```
mouse <- useMart("ensembl", dataset = "mmusculus_gene_ensembl")
human <- useMart("ensembl", dataset = "hsapiens_gene_ensembl")
```

getLDS kept failing to connect to the server, so had to use this
round-about fix…

```
mouse_genes <- rownames(data)

mouse_ids <- getBM(attributes = c("mgi_symbol", "ensembl_gene_id"),
                   filters = "mgi_symbol",
                   values = mouse_genes,
                   mart = mouse)

mouse_to_human <- getBM(attributes = c("ensembl_gene_id", "hsapiens_homolog_ensembl_gene"),
                        filters = "ensembl_gene_id",
                        values = mouse_ids$ensembl_gene_id,
                        mart = mouse)

human_genes <- getBM(attributes = c("ensembl_gene_id", "hgnc_symbol"),
                     filters = "ensembl_gene_id",
                     values = mouse_to_human$hsapiens_homolog_ensembl_gene,
                     mart = human)

orthologues <- merge(mouse_to_human, human_genes, 
                     by.x = "hsapiens_homolog_ensembl_gene", 
                     by.y = "ensembl_gene_id", 
                     all.x = TRUE)

orthologues <- merge(orthologues, mouse_ids, 
                       by = "ensembl_gene_id", 
                       all.x = TRUE)

colnames(orthologues) <- c("Mouse_Ensembl_ID", "Human_Ensembl_ID", "HGNC.symbol", "MGI.symbol")

print(orthologues)
```

```
data2 <- data %>%
  rownames_to_column(var = "MGI.symbol")

data_with_human <- data2 %>%
  left_join(orthologues, by = "MGI.symbol")

data_with_human <- data_with_human %>%
  distinct(HGNC.symbol, .keep_all = TRUE)

data_with_human <- data_with_human[!is.na(data_with_human$HGNC.symbol), ]

rownames(data_with_human) <- data_with_human$HGNC.symbol

data_with_human$MGI.symbol <- NULL
data_with_human$Mouse_Ensembl_ID <- NULL
data_with_human$Human_Ensembl_ID <- NULL
data_with_human$HGNC.symbol <- NULL

head(data_with_human)
```

```
##          CONTRA_1_F CONTRA_1_P CONTRA_10_F CONTRA_10_P CONTRA_11_F CONTRA_11_P
## C17orf49        501          0         661           0         674        2071
## C2orf68         419          1         400           0         373           0
## C11orf58       4863          1        3983        2476        4163        1650
## C8orf33         879          1         822           0        1774        3174
## C9orf85        1561          0        1042        1588        1484           3
## KIAA1143       1013          2         585         931         532           2
##          CONTRA_12_F CONTRA_12_P CONTRA_13_F CONTRA_13_P CONTRA_14_F CONTRA_2_F
## C17orf49         666           0         697        1343        1024       1311
## C2orf68          572           0         461           0         495        274
## C11orf58        4264           0        3317        7231        6702       3538
## C8orf33         1472        1534         790        3033         815        660
## C9orf85          865           0        1002        2747        1819       2305
## KIAA1143         634           0         668         891        1144        646
##          CONTRA_2_P CONTRA_3_F CONTRA_3_P CONTRA_4_F CONTRA_4_P CONTRA_5_F
## C17orf49          0        608        319        671        570        912
## C2orf68           0        475          3        595        481        627
## C11orf58       4706       3580       3468       3811       2865       3089
## C8orf33        2895        623       1121        622        889       1530
## C9orf85        3555        883          3        678       1476        503
## KIAA1143          0        544       1582        894        812        647
##          CONTRA_5_P CONTRA_6_F CONTRA_6_P CONTRA_7_F CONTRA_7_P CONTRA_8_F
## C17orf49        381        997       1409        668        287        826
## C2orf68         947        355         96        411          0        467
## C11orf58       5426       3323       6821       4700         35       3885
## C8orf33        1629       1300       1263        970          7        634
## C9orf85        1613       1262       2000       1715       1945       1471
## KIAA1143       1187        825          1       1068          2        449
##          CONTRA_8_P CONTRA_9_F CONTRA_9_P IPSI_1_F IPSI_10_F IPSI_10_P
## C17orf49        910        469       1892      915      1109       906
## C2orf68           0        554          0      358       220       332
## C11orf58       3553       2906       3096     3412      4980      5717
## C8orf33         318       1698       1091      986       971       871
## C9orf85           0       1515          1     1056      1471      2627
## KIAA1143        499        624          0      465       549       673
##          IPSI_11_F IPSI_11_P IPSI_12_P IPSI_2_F IPSI_2_P IPSI_3_F IPSI_3_P
## C17orf49      1173      1385      1732      885     1096      828     1123
## C2orf68        551       148       290      133      200      203      371
## C11orf58      3982      4367      7404     4984     6278     4073     5100
## C8orf33       1022      1197      1522      977     1206     1019      950
## C9orf85       1547      2243      2368     1880     2915     1253     1688
## KIAA1143       805       880      1268      694      802      821      754
##          IPSI_4_F IPSI_4_P IPSI_5_F IPSI_5_P IPSI_6_F IPSI_6_P IPSI_7_F
## C17orf49      649      723      738      926      673     1026     1035
## C2orf68       340      479      341      680      266      500      283
## C11orf58     3135     2945     2935     3594     3294     4466     6417
## C8orf33       581      966      535      885      791     1120     1052
## C9orf85       855      668      975     1660     1766     1834     2091
## KIAA1143      501      513      605      732      612      686      390
##          IPSI_7_P IPSI_8_F IPSI_8_P IPSI_9_P NF_uninjured_3 NP_uninjured_3
## C17orf49      656      734      838      836       177.8387        148.961
## C2orf68       342      655      502      602      1366.1238       1981.831
## C11orf58     2503     2641     3760     3112     48650.0071      18609.040
## C8orf33       779      752      743      950      3237.9870       1611.138
## C9orf85       825      991     1857      424      6324.0156       2529.927
## KIAA1143      696      407      702      460      6142.6852       3014.668
##          PEP_uninjured_3 cLTMR_uninjured_3 SST_uninjured_3 NF_injured_3
## C17orf49        103.5998          60.05918        9.170503     63.94313
## C2orf68        2338.3739        1190.01021      656.564118    432.91717
## C11orf58      21980.2573        8651.60130     5810.045548  20561.45268
## C8orf33        1819.1247         824.34358      442.010220   1050.64258
## C9orf85        1974.9727        1210.24574      738.002726   2279.96667
## KIAA1143       3535.7564        1239.08118      902.162791   1701.48475
##          NP_injured_3 PEP_injured_3 cLTMR_injured_3 SST_injured_3
## C17orf49     71.39749      30.98715        41.75452      26.24068
## C2orf68     612.03850     644.88474       531.64448      54.82659
## C11orf58  10291.53379    5995.50663      4157.84016    1100.74119
## C8orf33     793.84248     485.35180       316.78778      85.36186
## C9orf85     714.00843     383.10842       297.55592      67.03274
## KIAA1143    977.64289     769.47448       422.03457     112.35739
```

```
data.scaled=gene.scaling(data = data_with_human, n=1, db = db)
head(data.scaled)
```

```
##        CONTRA_1_F   CONTRA_1_P CONTRA_10_F  CONTRA_10_P  CONTRA_11_F
## ABCA1  0.15647290 0.0001590172  0.16824017 0.0001590172  0.442862822
## ACKR1  0.00000000 0.0000000000  0.02452309 0.0000000000  0.028585652
## ACKR2  0.00000000 0.0000000000  0.65835606 0.0000000000  1.663420335
## ACKR3  1.73038229 0.0022356360  2.95774648 0.2073552426 10.000000000
## ACVR1  0.61332365 0.0007215572  0.97771005 0.0000000000  0.052673678
## ACVR1B 0.03830664 0.0000000000  0.14447077 0.0000000000  0.006566853
##         CONTRA_11_P CONTRA_12_F CONTRA_12_P CONTRA_13_F CONTRA_13_P CONTRA_14_F
## ABCA1  0.0001590172 0.139299042  0.00000000   0.3539722  0.52618782  0.38195925
## ACKR1  0.0000000000 0.008494445  0.00000000   0.0573929  0.03368232  0.02910271
## ACKR2  0.0000000000 0.895987534  0.00000000   4.6747176  0.00000000  1.16088820
## ACKR3  0.0078247261 2.281466577  0.00558909   5.2218869  5.18947015  8.16789627
## ACVR1  0.0000000000 0.354284600  0.00000000   0.4264403  0.93874596  0.74897641
## ACVR1B 0.0000000000 0.056912728  0.00000000   0.1844191  0.00000000  0.14228182
##         CONTRA_2_F CONTRA_2_P CONTRA_3_F   CONTRA_3_P CONTRA_4_F   CONTRA_4_P
## ABCA1  0.205291168  0.4188512 0.12880391  0.002385258 0.18541402 0.1542466562
## ACKR1  0.021716232  0.0000000 0.02400604  0.000000000 0.09661507 0.0000000000
## ACKR2  2.006232957  0.0000000 3.54499416 10.000000000 0.78691079 0.0000000000
## ACKR3  4.546724793  8.7178627 5.88978314  0.013413816 8.11256427 1.7577688352
## ACVR1  0.734545261  1.0036861 0.29872469  0.001443114 0.47478466 0.7619644353
## ACVR1B 0.004377902  0.0000000 0.02955084  0.000000000 0.07716053 0.0005472378
##          CONTRA_5_F  CONTRA_5_P CONTRA_6_F CONTRA_6_P CONTRA_7_F  CONTRA_7_P
## ABCA1  0.0003180343 0.080144654 0.21324203 0.70444607 0.42441683 0.002067223
## ACKR1  0.0000000000 0.000000000 0.01455135 0.01676729 0.02814246 0.000000000
## ACKR2  0.0000000000 0.000000000 0.75964160 0.00000000 0.41293339 0.003895598
## ACKR3  1.4391906998 1.443103063 3.95931142 6.19047619 2.60507489 0.026268723
## ACVR1  0.6450721640 0.888958508 0.25543126 1.06934782 0.69485961 0.002164672
## ACVR1B 0.0000000000 0.005472378 0.12367574 0.00000000 0.08810528 0.001094476
##        CONTRA_8_F CONTRA_8_P CONTRA_9_F   CONTRA_9_P   IPSI_1_F   IPSI_10_F
## ABCA1  0.11083497  0.0000000 0.13373344 0.0001590172 0.29227356 0.147408918
## ACKR1  0.04033015  0.0000000 0.01366498 0.0028807247 0.02260261 0.005539855
## ACKR2  2.35683677  0.0000000 0.00000000 0.0000000000 0.21036229 0.112972341
## ACKR3  4.32707355  0.3828527 2.62016544 5.5639391907 3.14050973 5.724904985
## ACVR1  0.42355409  0.0000000 0.25615282 0.4993176034 0.75474886 0.663832652
## ACVR1B 0.05527101  0.0000000 0.07880224 0.0000000000 0.26759927 0.085369092
##         IPSI_10_P  IPSI_11_F    IPSI_11_P   IPSI_12_P   IPSI_2_F     IPSI_2_P
## ABCA1  0.13309737 0.57166673 0.4741892049 0.362559151 0.39118224 1.302351e-01
## ACKR1  0.00000000 0.02577879 0.0005170531 0.005096667 0.02341512 7.386474e-05
## ACKR2  0.00000000 0.48694975 0.0000000000 0.003895598 0.52590573 0.000000e+00
## ACKR3  2.74536106 4.99049855 1.3419405321 1.957299352 2.28482003 9.501453e-01
## ACVR1  0.68980871 1.03543463 0.7727877938 1.023168153 1.02461127 6.356919e-01
## ACVR1B 0.05691273 0.16581304 0.0333815039 0.063479581 0.06949920 6.402682e-02
##          IPSI_3_F    IPSI_3_P    IPSI_4_F    IPSI_4_P   IPSI_5_F     IPSI_5_P
## ABCA1  0.74769874 0.499313918 0.641157235 0.912917581 0.98527039 7.163724e-01
## ACKR1  0.01071039 0.001698889 0.002954589 0.001698889 0.01514227 7.386474e-05
## ACKR2  0.00000000 0.000000000 0.003895598 0.000000000 0.87650954 0.000000e+00
## ACKR3  4.53331098 2.173038229 2.071875699 0.552761011 4.39805500 2.397720e-01
## ACVR1  0.70279674 0.472619986 0.855766875 0.640742821 0.54116792 8.377279e-01
## ACVR1B 0.13626220 0.039401119 0.050345875 0.062385106 0.05034587 0.000000e+00
##           IPSI_6_F   IPSI_6_P    IPSI_7_F    IPSI_7_P   IPSI_8_F    IPSI_8_P
## ABCA1  0.342363970 0.09413817 0.165377858 0.142479386 0.14899909 0.173964785
## ACKR1  0.009306957 0.00546599 0.017579807 0.002437536 0.01964802 0.004431884
## ACKR2  0.003895598 0.00000000 0.003895598 0.000000000 1.14530580 0.000000000
## ACKR3  2.452492734 0.86183769 7.411133467 0.476190476 6.68902303 0.025150905
## ACVR1  0.696302727 0.59817094 0.256874374 0.428604995 0.63713503 0.422832537
## ACVR1B 0.059101679 0.07825500 0.060196155 0.080443952 0.12422297 0.005472378
##           IPSI_9_P NF_uninjured_3 NP_uninjured_3 PEP_uninjured_3
## ABCA1  1.678585260      5.4649470      3.6269697       1.8796782
## ACKR1  0.007755797     10.0000000      1.4902853       1.3615097
## ACKR2  0.000000000      0.7909393      0.3364550       0.3880482
## ACKR3  0.113458529      0.2621640      0.1311876       0.2149630
## ACVR1  1.130680180     10.0000000      3.6645883       4.8662476
## ACVR1B 0.039948357     10.0000000      2.8203224       4.4893623
##        cLTMR_uninjured_3 SST_uninjured_3 NF_injured_3 NP_injured_3
## ABCA1         1.64655062      1.09387868  10.00000000   5.60969464
## ACKR1         0.87547461      0.15217781   3.15371686   1.09024335
## ACKR2         0.08178383      0.09312792   0.12567455   0.13922279
## ACKR3         0.10765131      0.07528543   0.07396997   0.06558743
## ACVR1         2.18457076      0.89943918   9.44139056   3.06553143
## ACVR1B        1.76739284      0.76904511   2.37914375   0.92353999
##        PEP_injured_3 cLTMR_injured_3 SST_injured_3 Symbol
## ABCA1     1.69723720      2.20737333   0.324721885  ABCA1
## ACKR1     0.47542029      0.52385271   0.054413044  ACKR1
## ACKR2     0.07083190      0.03924640   0.003203090  ACKR2
## ACKR3     0.03038932      0.02172749   0.008444671  ACKR3
## ACVR1     2.81692352      1.35081511   0.264850280  ACVR1
## ACVR1B    0.66701029      0.45682796   0.075840478 ACVR1B
```

```
CC.data.selection.P=meta$ID[which(meta$Cell_type=="pericytes")]
CC.data.selection.F=meta$ID[which(meta$Cell_type=="fibroblasts")] 
CC.data.selection.Pi=meta$ID[which(meta$Cell_type=="pericytes"& meta$Class=="ipsi")]
CC.data.selection.Fi=meta$ID[which(meta$Cell_type=="fibroblasts"&meta$Class=="ipsi")]
CC.data.selection.PC=meta$ID[which(meta$Cell_type=="pericytes"& meta$Class=="contra")]
CC.data.selection.Fc=meta$ID[which(meta$Cell_type=="fibroblasts"&meta$Class=="contra")]
PC.data.selection=meta$ID[which(meta$Cell_type%in% c("NF_neurons", "nociceptors", "cLTMRs"))]
                                   
CC_Pi=data.scaled[, CC.data.selection.Pi]
CC_Pc=data.scaled[, CC.data.selection.PC]
CC_Fi=data.scaled[, CC.data.selection.Fi]
CC_Fc=data.scaled[, CC.data.selection.Fc]
PC = data.scaled[, PC.data.selection]
```

```
score.Pi= icellnet.score(direction="out", PC.data=PC, CC.data= CC_Pi, PC = c("NF_neurons", "nociceptors", "cLTMRs"), PC.target = meta, CC.type = "RNAseq",  PC.type = "RNAseq",  db = db) 
score_Pi=as.data.frame(score.Pi[[1]])
lr1_Pi=score.Pi[[2]]
```

```
head(score_Pi)
```

```
##                   out
## NF_neurons  4251.5054
## nociceptors 1401.7023
## cLTMRs       922.6978
```

```
head(lr1_Pi)
```

```
##                        NF_neurons nociceptors   cLTMRs
## TIMP1 + MMP9 / LRP1      3.297784    4.562423 3.595718
## LTB + LTA / LTBR               NA          NA       NA
## ITGAV + ITGB5 / ADGRB1         NA          NA       NA
## ITGAV + ITGB3 / ADGRA2  10.946496    6.404581 3.952509
## ITGAV + ITGB3 / ADGRE5   3.587557    2.141840 7.821530
## ITGA5 + ITGB1 / ADGRE5   2.611858    1.559329 5.694327
```

```
score.Pc= icellnet.score(direction="out", PC.data=PC, CC.data= CC_Pc, PC = c("NF_neurons", "nociceptors", "cLTMRs"), PC.target = meta, CC.type = "RNAseq",  PC.type = "RNAseq",  db = db) 
score_Pc=as.data.frame(score.Pc[[1]])
lr1_Pc=score.Pc[[2]]
```

```
head(score_Pc)
```

```
##                   out
## NF_neurons  1837.5694
## nociceptors  663.5530
## cLTMRs       448.2429
```

```
head(lr1_Pc)
```

```
##                        NF_neurons nociceptors   cLTMRs
## TIMP1 + MMP9 / LRP1     0.1046239   0.1447452 0.114076
## LTB + LTA / LTBR               NA          NA       NA
## ITGAV + ITGB5 / ADGRB1         NA          NA       NA
## ITGAV + ITGB3 / ADGRA2  3.0476834   1.7831400 1.100443
## ITGAV + ITGB3 / ADGRE5  0.9988344   0.5963232 2.177642
## ITGA5 + ITGB1 / ADGRE5  0.7685659   0.4588485 1.675614
```

```
score.Fi= icellnet.score(direction="out", PC.data=PC, CC.data= CC_Fi, PC = c("NF_neurons", "nociceptors", "cLTMRs"), PC.target = meta, CC.type = "RNAseq",  PC.type = "RNAseq",  db = db) 
score_Fi=as.data.frame(score.Fi[[1]])
lr1_Fi=score.Fi[[2]]
```

```
head(score_Fi)
```

```
##                  out
## NF_neurons  5259.001
## nociceptors 1748.089
## cLTMRs      1163.388
```

```
head(lr1_Fi)
```

```
##                        NF_neurons nociceptors    cLTMRs
## TIMP1 + MMP9 / LRP1      1.662100    2.299485  1.812261
## LTB + LTA / LTBR               NA          NA        NA
## ITGAV + ITGB5 / ADGRB1         NA          NA        NA
## ITGAV + ITGB3 / ADGRA2  14.220657    8.320229  5.134728
## ITGAV + ITGB3 / ADGRE5   4.660616    2.782477 10.160996
## ITGA5 + ITGB1 / ADGRE5   2.121505    1.266579  4.625268
```

```
score.Fc= icellnet.score(direction="out", PC.data=PC, CC.data= CC_Fc, PC = c("NF_neurons", "nociceptors", "cLTMRs"), PC.target = meta, CC.type = "RNAseq",  PC.type = "RNAseq",  db = db) 
score_Fc=as.data.frame(score.Fc[[1]])
lr1_Fc=score.Fc[[2]]
```

```
head(score_Fc)
```

```
##                   out
## NF_neurons  4031.3936
## nociceptors 1398.8393
## cLTMRs       956.3662
```

```
head(lr1_Fc)
```

```
##                         NF_neurons nociceptors      cLTMRs
## TIMP1 + MMP9 / LRP1    0.006327962 0.008754619 0.006899654
## LTB + LTA / LTBR                NA          NA          NA
## ITGAV + ITGB5 / ADGRB1          NA          NA          NA
## ITGAV + ITGB3 / ADGRA2 9.282385340 5.430942365 3.351639772
## ITGAV + ITGB3 / ADGRE5 3.042168413 1.816232508 6.632483767
## ITGA5 + ITGB1 / ADGRE5 1.118405700 0.667709513 2.438329061
```

```
Scores=cbind(score_Pi,score_Pc, score_Fi, score_Fc)
colnames(Scores)=c("injured_PC","PC", "injured_FB", "FB")
Scores
```

```
##             injured_PC        PC injured_FB        FB
## NF_neurons   4251.5054 1837.5694   5259.001 4031.3936
## nociceptors  1401.7023  663.5530   1748.089 1398.8393
## cLTMRs        922.6978  448.2429   1163.388  956.3662
```

```
ymax=max(Scores)+1 #for later visualisations

# table of contribution of each family of molecule to the scores
LR.family.score(lr=lr1_Pi, db.couple=db.name.couple, plot=NULL)
```

```
##                  NF_neurons nociceptors      cLTMRs
## Cytokine         494.772422 196.4235639 114.2374901
## Cell adhesion    568.560158 181.8635375 143.4675541
## Innate immune     12.727475  10.5100872   3.6250161
## ECM interaction 2468.059490 695.9082382 437.9389849
## Growth factor    293.521859 115.6374851 100.7610953
## Wnt pathway       34.650214  22.9589328  16.6926015
## HLA recognition    1.075393   0.3366021   0.2442952
## Chemokine         42.540837   6.5402540   4.8062954
## Checkpoint         0.000000   0.0000000   0.0000000
## Notch pathway     10.689629   9.7920788   6.9037708
## other            324.907915 161.7315174  94.0207392
```

```
#display heatmap
LR.family.score(lr=lr1_Pi, db.couple=db.name.couple, plot="heatmap")
```

```
#display barplot
colors=grDevices::colorRampPalette(RColorBrewer::brewer.pal(8,"Set2"))(length(unique(db$Family)))
LR.family.score(lr=lr1_Pi, db.couple=db.name.couple, plot="barplot", family.col=colors)
```

```
ymax=max(Scores)+1 #for later visualisations

# table of contribution of each family of molecule to the scores
LR.family.score(lr=lr1_Pc, db.couple=db.name.couple, plot=NULL)
```

```
##                  NF_neurons nociceptors      cLTMRs
## Cytokine        411.4954164 156.9301474  92.4266239
## Cell adhesion   163.2221173  69.4550909  52.3211707
## Innate immune     9.4023476   7.0961268   3.0294520
## ECM interaction 783.2727265 220.6998522 137.9510724
## Growth factor   215.8749336  86.2454954  81.1007960
## Wnt pathway      46.7912505  29.4341313  23.8576812
## HLA recognition   0.8269545   0.2588398   0.1878578
## Chemokine        11.2347908   1.7976584   1.3418421
## Checkpoint        0.0000000   0.0000000   0.0000000
## Notch pathway     6.8049492   6.3682989   4.5094358
## other           188.6438973  85.2673631  51.5169322
```

```
#display heatmap
LR.family.score(lr=lr1_Pc, db.couple=db.name.couple, plot="heatmap")
```

```
#display barplot
colors=grDevices::colorRampPalette(RColorBrewer::brewer.pal(8,"Set2"))(length(unique(db$Family)))
LR.family.score(lr=lr1_Pc, db.couple=db.name.couple, plot="barplot", family.col=colors)
```

```
ymax=max(Scores)+1 #for later visualisations

# table of contribution of each family of molecule to the scores
LR.family.score(lr=lr1_Fi, db.couple=db.name.couple, plot=NULL)
```

```
##                   NF_neurons  nociceptors       cLTMRs
## Cytokine        3.857585e+02 159.31292615 9.225268e+01
## Cell adhesion   5.340920e+02 195.92975613 1.501207e+02
## Innate immune   6.091589e+01  34.59606622 2.428834e+01
## ECM interaction 3.324902e+03 952.59370032 5.965870e+02
## Growth factor   3.993168e+02 150.09178557 1.335133e+02
## Wnt pathway     1.247531e+02  73.55559396 5.530540e+01
## HLA recognition 4.077695e-02   0.01276334 9.263228e-03
## Chemokine       4.222554e+01   6.99918866 4.926320e+00
## Checkpoint      0.000000e+00   0.00000000 0.000000e+00
## Notch pathway   1.444591e+01  13.37514103 9.450595e+00
## other           3.725509e+02 161.62176797 9.693452e+01
```

```
#display heatmap
LR.family.score(lr=lr1_Fi, db.couple=db.name.couple, plot="heatmap")
```

```
#display barplot
colors=grDevices::colorRampPalette(RColorBrewer::brewer.pal(8,"Set2"))(length(unique(db$Family)))
LR.family.score(lr=lr1_Fi, db.couple=db.name.couple, plot="barplot", family.col=colors)
```

```
ymax=max(Scores)+1 #for later visualisations

# table of contribution of each family of molecule to the scores
LR.family.score(lr=lr1_Fc, db.couple=db.name.couple, plot=NULL)
```

```
##                   NF_neurons  nociceptors       cLTMRs
## Cytokine        6.494386e+02 233.13127190 133.30056445
## Cell adhesion   3.580371e+02 151.26376633 115.68674311
## Innate immune   1.062380e+02  49.22698550  46.97200477
## ECM interaction 1.904715e+03 538.61311359 339.86524559
## Growth factor   4.040924e+02 158.14919794 146.14479703
## Wnt pathway     8.743287e+01  62.76768670  46.18334357
## HLA recognition 4.540852e-02   0.01421304   0.01031537
## Chemokine       5.113780e+01   7.65627538   5.90488766
## Checkpoint      0.000000e+00   0.00000000   0.00000000
## Notch pathway   5.396455e+00   4.88768875   3.43792004
## other           4.648601e+02 193.12905230 118.86036041
```

```
#display heatmap
LR.family.score(lr=lr1_Fc, db.couple=db.name.couple, plot="heatmap")
```

```
#display barplot
colors=grDevices::colorRampPalette(RColorBrewer::brewer.pal(8,"Set2"))(length(unique(db$Family)))
LR.family.score(lr=lr1_Fc, db.couple=db.name.couple, plot="barplot", family.col=colors)
```

```
## Label and range definition
my.family=c("Growth factor","Chemokine","HLA recognition","Cytokine","Notch pathway","ECM interaction", "Innate Immune", "Cell adhesion", "Wnt pathway")
family.col = c("Growth factor" = "#ffadad","Chemokine" = "#ffd6a5","HLA recognition"= "#fdffb6","Cytokine" = 
"#a0c4ff","Notch pathway" = "#9bf6ff","ECM interaction" = "#caffbf", "Innate Immune" = "#bdb2ff", "Cell adhesion" = "#ffc6ff", "Wnt pathway" = "#C7E7AB")

ymax=round(max(Scores))+1 #to define the y axis range of the barplot

# to get family contribution as dataframe
LR.family.score(lr=lr1_Pi, family=my.family, db.couple=db.name.couple, plot= NULL)
```

```
##                  NF_neurons nociceptors      cLTMRs
## Growth factor    293.521859 115.6374851 100.7610953
## Chemokine         42.540837   6.5402540   4.8062954
## HLA recognition    1.075393   0.3366021   0.2442952
## Cytokine         494.772422 196.4235639 114.2374901
## Notch pathway     10.689629   9.7920788   6.9037708
## ECM interaction 2468.059490 695.9082382 437.9389849
## Innate Immune      0.000000   0.0000000   0.0000000
## Cell adhesion    568.560158 181.8635375 143.4675541
## Wnt pathway       34.650214  22.9589328  16.6926015
## other            337.635390 172.2416046  97.6457552
```

```
# barplot
contrib.family1= LR.family.score(lr=lr1_Pi, family=my.family, db.couple=db.name.couple, plot="barplot", ymax=ymax, family.col=family.col)
contrib.family2= LR.family.score(lr=lr1_Pc, family=my.family, db.couple=db.name.couple, plot="barplot", ymax=ymax, family.col=family.col)

gridExtra::grid.arrange(contrib.family1, contrib.family2, ncol=2, nrow=1)
```

```
## Label and range definition
my.family=c("Growth factor","Chemokine","HLA recognition","Cytokine","Notch pathway","ECM interaction", "Innate Immune", "Cell adhesion", "Wnt pathway")
family.col = c("Growth factor" = "#ffadad","Chemokine" = "#ffd6a5","HLA recognition"= "#fdffb6","Cytokine" = 
"#a0c4ff","Notch pathway" = "#9bf6ff","ECM interaction" = "#caffbf", "Innate Immune" = "#bdb2ff", "Cell adhesion" = "#ffc6ff", "Wnt pathway" = "#C7E7AB")

ymax=round(max(Scores))+1 #to define the y axis range of the barplot

# to get family contribution as dataframe
LR.family.score(lr=lr1_Fi, family=my.family, db.couple=db.name.couple, plot= NULL)
```

```
##                   NF_neurons  nociceptors       cLTMRs
## Growth factor   3.993168e+02 150.09178557 1.335133e+02
## Chemokine       4.222554e+01   6.99918866 4.926320e+00
## HLA recognition 4.077695e-02   0.01276334 9.263228e-03
## Cytokine        3.857585e+02 159.31292615 9.225268e+01
## Notch pathway   1.444591e+01  13.37514103 9.450595e+00
## ECM interaction 3.324902e+03 952.59370032 5.965870e+02
## Innate Immune   0.000000e+00   0.00000000 0.000000e+00
## Cell adhesion   5.340920e+02 195.92975613 1.501207e+02
## Wnt pathway     1.247531e+02  73.55559396 5.530540e+01
## other           4.334668e+02 196.21783418 1.212229e+02
```

```
# barplot
contrib.family1= LR.family.score(lr=lr1_Fi, family=my.family, db.couple=db.name.couple, plot="barplot", ymax=ymax, family.col=family.col)
contrib.family2= LR.family.score(lr=lr1_Fc, family=my.family, db.couple=db.name.couple, plot="barplot", ymax=ymax, family.col=family.col)

gridExtra::grid.arrange(contrib.family1, contrib.family2, ncol=2, nrow=1)
```

plot for revisions.

```
library(grid)
library(gridExtra)
```

```
## 
## Attaching package: 'gridExtra'
```

```
## The following object is masked from 'package:Biobase':
## 
##     combine
```

```
## The following object is masked from 'package:BiocGenerics':
## 
##     combine
```

```
## The following object is masked from 'package:dplyr':
## 
##     combine
```

```
contrib.family1= LR.family.score(lr=lr1_Fi, family=my.family, db.couple=db.name.couple, plot="barplot", ymax=ymax, family.col=family.col, title = "Injured FBs") + theme(legend.position = "none", axis.title = element_text(size = 40, face = "bold", family = "Arial"),  
    axis.text = element_text(size = 35, family = "Arial"), legend.text = element_text(size = 40, family = "Arial", face = "bold"), plot.title = element_text(size = 40, face = "bold", family = "Arial"))
contrib.family2= LR.family.score(lr=lr1_Fc, family=my.family, db.couple=db.name.couple, plot="barplot", ymax=ymax, family.col=family.col, title = "Fibroblasts") + theme(axis.title = element_text(size = 40, face = "bold", family = "Arial"), plot.title = element_text(size = 40, face = "bold", family = "Arial"), axis.text = element_text(size = 35, family = "Arial"), legend.text = element_text(size = 40, family = "Arial", face = "bold"))

legend <- gtable::gtable_filter(ggplotGrob(contrib.family2), "guide-box")

contrib.family2 <- contrib.family2 + theme(legend.position = "none")

grid.arrange(contrib.family1, contrib.family2, legend, ncol = 3, widths = c(0.8, 0.8, 0.8))
```

```
library(grid)
library(gridExtra)
contrib.family1= LR.family.score(lr=lr1_Pi, family=my.family, db.couple=db.name.couple, plot="barplot", ymax=ymax, family.col=family.col, title = "Injured MCs") + theme(legend.position = "none", axis.title = element_text(size = 40, face = "bold", family = "Arial"),  
    axis.text = element_text(size = 35, family = "Arial"), legend.text = element_text(size = 40, family = "Arial", face = "bold"), plot.title = element_text(size = 40, face = "bold", family = "Arial"))
contrib.family2= LR.family.score(lr=lr1_Pc, family=my.family, db.couple=db.name.couple, plot="barplot", ymax=ymax, family.col=family.col, title = "Mural Cells") + theme(axis.title = element_text(size = 40, face = "bold", family = "Arial"), plot.title = element_text(size = 40, face = "bold", family = "Arial"), axis.text = element_text(size = 35, family = "Arial"), legend.text = element_text(size = 40, family = "Arial", face = "bold"))

legend <- gtable::gtable_filter(ggplotGrob(contrib.family2), "guide-box")

contrib.family2 <- contrib.family2 + theme(legend.position = "none")

grid.arrange(contrib.family1, contrib.family2, legend, ncol = 3, widths = c(0.8, 0.8, 0.8))
```

```
# Custom version of LR.balloon.plot
LR.balloon.plot.custom <- function(lr = lr, thresh = 0, topn = NULL, sort.by = "sum", 
    db.name.couple = NULL, title = title, family.col = family.col) {
    if (is.null(db.name.couple)) {
        db.name.couple = name.lr.couple(db, "Family")
    }
    interactions = rownames(lr)
    lr = as.data.frame(lr)
    lr$Pair = interactions
    lr = as.data.frame(lr[stats::complete.cases(lr), ])
    lr = lr %>% dplyr::filter_if(is.numeric, dplyr::any_vars(. > 
        0)) %>% dplyr::filter_if(is.numeric, dplyr::any_vars(. >= 
        thresh))
    if (!is.null(topn)) {
        if (topn > dim(lr)[1]) {
            note(paste0("lr contains only ", dim(lr)[1], " after filtering interaction highest than theshold"))
            lr = dplyr::left_join(lr, as.data.frame(db.name.couple), 
                by = "Pair")
        }
        else {
            if (sort.by == "sum") {
                lr = lr %>% dplyr::mutate(sum = rowSums(dplyr::across(where(is.numeric)))) %>% 
                  dplyr::arrange(dplyr::desc(sum)) %>% dplyr::top_n(topn, 
                  sum)
                lr = dplyr::left_join(lr, as.data.frame(db.name.couple), 
                  by = "Pair") %>% dplyr::select(-sum)
            }
            else if (sort.by == "var") {
                lr$variance = apply(dplyr::select_if(lr, is.numeric), 
                  1, var, na.rm = TRUE)
                lr = lr %>% dplyr::arrange(dplyr::desc(variance)) %>% 
                  dplyr::top_n(topn, variance)
                lr = dplyr::left_join(lr, as.data.frame(db.name.couple), 
                  by = "Pair") %>% dplyr::select(-variance)
            }
            else stop("sort.by argument should be fixed on var or sum")
        }
    }
    else {
        lr = dplyr::left_join(lr, as.data.frame(db.name.couple), 
            by = "Pair")
    }
    if (!(is.null(lr$Subfamily))) {
        lr = lr %>% dplyr::rename(Family = Subfamily)
    }
    melted <- reshape2::melt(lr, id.vars = c("Pair", "Family"))
    melted$Family[is.na(melted$Family)] <- "NA"
    melted = melted %>% dplyr::arrange(Family, variable)
    melted <- melted %>% dplyr::mutate(row = dplyr::group_indices(melted, 
        .dots = c("Family", "Pair")))
    melted <- melted %>% dplyr::mutate(col = dplyr::group_indices(melted, 
        .dots = c("variable")))
    vars_x_axis <- c(melted %>% dplyr::arrange(col) %>% dplyr::select(variable) %>% 
        dplyr::distinct())$variable
    names_y_axis <- c(melted %>% dplyr::arrange(row) %>% dplyr::group_by(Pair) %>% 
        dplyr::distinct(Pair) %>% dplyr::select(Pair))$Pair
    plot <- ggplot2::ggplot(melted, ggplot2::aes(x = factor(col), 
        y = factor(row), color = factor(Family), size = value)) + 
        ggplot2::geom_point() +  
      ggplot2::geom_text(ggplot2::aes(label = round(value), 
        x = col + 0.4), alpha = 1, size = 10) +
        ggplot2::scale_size_area(max_size = 12) + 
        ggplot2::scale_x_discrete(breaks = 1:length(vars_x_axis), 
            labels = vars_x_axis, position = "top") + ggplot2::scale_y_discrete(breaks = 1:length(names_y_axis), 
        labels = names_y_axis) + ggplot2::scale_color_manual(values = family.col) + 
        ggplot2::theme_bw() + ggplot2::labs(title = title) + 
        ggplot2::theme(axis.line = ggplot2::element_blank(), 
            axis.title = ggplot2::element_blank(), panel.border = ggplot2::element_blank(), 
            panel.grid.major.x = ggplot2::element_blank(), panel.grid.minor.x = ggplot2::element_blank(), 
            axis.text.x = ggplot2::element_text(angle = 0), axis.ticks.x = ggplot2::element_blank(), 
            axis.ticks.y = ggplot2::element_blank())
    return(plot)
}
```

```
lr_ind=cbind(lr1_Pi[,"nociceptors"],lr1_Pc[,"nociceptors"])
colnames(lr_ind)=c("injured_MC", "MC")
balloon_custom = LR.balloon.plot.custom(lr = lr_ind, sort.by="var", thresh = 0, topn=25, db.name.couple=db.name.couple, title="Top 25 differential L-R pairs", family.col=family.col) &  theme(axis.text.x= element_text(size=40, face = "bold"), axis.text.y= element_text(size=40), legend.text = element_text(size=35), legend.title = element_text(size=40))
```

```
## Warning: There were 2 warnings in `dplyr::mutate()`.
## The first warning was:
## ℹ In argument: `row = dplyr::group_indices(melted, .dots = c("Family",
##   "Pair"))`.
## Caused by warning:
## ! The `...` argument of `group_indices()` is deprecated as of dplyr 1.0.0.
## ℹ Please `group_by()` first
## ℹ Run `dplyr::last_dplyr_warnings()` to see the 1 remaining warning.
```

```
## Warning: There was 1 warning in `dplyr::mutate()`.
## ℹ In argument: `col = dplyr::group_indices(melted, .dots = c("variable"))`.
## Caused by warning:
## ! The `...` argument of `group_indices()` is deprecated as of dplyr 1.0.0.
## ℹ Please `group_by()` first
```

```
print(balloon_custom)
```

```
lr_ind=cbind(lr1_Fi[,"nociceptors"], lr1_Fc[,"nociceptors"])
colnames(lr_ind)=c("injured_FB", "FB")
balloon_custom = LR.balloon.plot.custom(lr = lr_ind, sort.by="var", thresh = 0, topn=25, db.name.couple=db.name.couple, title="Top 25 differential L-R pairs", family.col=family.col) &  theme(axis.text.x= element_text(size=40, face = "bold"), axis.text.y= element_text(size=40), legend.text = element_text(size=35), legend.title = element_text(size=40))
```

```
## Warning: There was 1 warning in `dplyr::mutate()`.
## ℹ In argument: `row = dplyr::group_indices(melted, .dots = c("Family",
##   "Pair"))`.
## Caused by warning:
## ! The `...` argument of `group_indices()` is deprecated as of dplyr 1.0.0.
## ℹ Please `group_by()` first
```

```
## Warning: There was 1 warning in `dplyr::mutate()`.
## ℹ In argument: `col = dplyr::group_indices(melted, .dots = c("variable"))`.
## Caused by warning:
## ! The `...` argument of `group_indices()` is deprecated as of dplyr 1.0.0.
## ℹ Please `group_by()` first
```

```
print(balloon_custom)
```

```
selected_pairs <- c(
  "SPP1 / CD44", "LIF / LIFR + IL6ST", "IL6 / IL6R + IL6ST", "GRN / TNFRSF1B",
  "CLCF1 + CRLF1 / CNTFR + IL6ST", "TNC / ITGB1 + ITGA7", "JAM3 / JAM3",
  "COL6A3 / DDR1", "COL8A1 / DDR1", "COL6A1 / DDR1", "COL5A1 / DDR1",
  "COL5A2 / DDR1", "COL4A2 / DDR1", "RSPO3 / LGR4", "APOE / LDLR",
  "DCN / TLR4", "IGF1 / IGF1R", "VCAN / CD44", "COL4A5 / DDR1",
  "COL3A1 / DDR1", "COL1A1 / DDR1", "COL12A1 / ITGB1 + ITGA2",
  "JAM2 / JAM3", "SPP1 / ITGB1 + ITGA8"
)
```

```
LR.balloon.plot.custom <- function(lr = lr, thresh = 0, selected_pairs = NULL, sort.by = "sum", 
    db.name.couple = NULL, title = title, family.col = family.col) {

    if (is.null(db.name.couple)) {
        db.name.couple = name.lr.couple(db, "Family")
    }
  
    # Use rownames as the Pair column
    interactions = rownames(lr)
    lr = as.data.frame(lr)
    lr$Pair = interactions  # Add 'Pair' column with the rownames
    
    # Filter rows with complete cases
    lr = lr[stats::complete.cases(lr), ]
    lr = lr %>% dplyr::filter_if(is.numeric, dplyr::any_vars(. > 0)) %>% 
        dplyr::filter_if(is.numeric, dplyr::any_vars(. >= thresh))
    
    # Filter based on selected pairs (using rownames now)
    if (!is.null(selected_pairs)) {
        lr <- lr %>% dplyr::filter(Pair %in% selected_pairs)  # Filter based on the Pair column
    }

    # Join with family data
    lr = dplyr::left_join(lr, as.data.frame(db.name.couple), by = "Pair")

    if (!(is.null(lr$Subfamily))) {
        lr = lr %>% dplyr::rename(Family = Subfamily)
    }

    # Melt the data for plotting
    melted <- reshape2::melt(lr, id.vars = c("Pair", "Family"))
    melted$Family[is.na(melted$Family)] <- "NA"
    melted = melted %>% dplyr::arrange(Family, variable)
    melted <- melted %>% dplyr::mutate(row = dplyr::group_indices(melted, .dots = c("Family", "Pair")))
    melted <- melted %>% dplyr::mutate(col = dplyr::group_indices(melted, .dots = c("variable")))

    # Variables for axis labels
    vars_x_axis <- c(melted %>% dplyr::arrange(col) %>% dplyr::select(variable) %>% dplyr::distinct())$variable
    names_y_axis <- c(melted %>% dplyr::arrange(row) %>% dplyr::group_by(Pair) %>% dplyr::distinct(Pair) %>% dplyr::select(Pair))$Pair

    # Create plot
    plot <- ggplot2::ggplot(melted, ggplot2::aes(x = factor(col), 
        y = factor(row), color = factor(Family), size = value)) + 
        ggplot2::geom_point() +  
        ggplot2::geom_text(ggplot2::aes(label = round(value), x = col + 0.4), alpha = 1, size = 10) +
        ggplot2::scale_size_area(max_size = 12) + 
        ggplot2::scale_x_discrete(breaks = 1:length(vars_x_axis), labels = vars_x_axis, position = "top") + 
        ggplot2::scale_y_discrete(breaks = 1:length(names_y_axis), labels = names_y_axis) + 
        ggplot2::scale_color_manual(values = family.col) + 
        ggplot2::theme_bw() + ggplot2::labs(title = title) + 
        ggplot2::theme(axis.line = ggplot2::element_blank(), 
            axis.title = ggplot2::element_blank(), panel.border = ggplot2::element_blank(), 
            panel.grid.major.x = ggplot2::element_blank(), panel.grid.minor.x = ggplot2::element_blank(), 
            axis.text.x = ggplot2::element_text(angle = 0), axis.ticks.x = ggplot2::element_blank(), 
            axis.ticks.y = ggplot2::element_blank())

    return(plot)
}
```

```
lr_ind=cbind(lr1_Pi[,"nociceptors"],lr1_Pc[,"nociceptors"], lr1_Fi[,"nociceptors"], lr1_Fc[,"nociceptors"])
colnames(lr_ind)=c("injured_MC", "MC", "injured_FB", "FB")
balloon_custom = LR.balloon.plot.custom(lr = lr_ind, sort.by="var", thresh = 0, selected_pairs = selected_pairs, db.name.couple=db.name.couple, title = "", family.col=family.col) &  theme(axis.text.x= element_text(size=40, face = "bold"), axis.text.y= element_text(size=40), legend.text = element_text(size=35), legend.title = element_text(size=40))
```

```
## Warning: There was 1 warning in `dplyr::mutate()`.
## ℹ In argument: `row = dplyr::group_indices(melted, .dots = c("Family",
##   "Pair"))`.
## Caused by warning:
## ! The `...` argument of `group_indices()` is deprecated as of dplyr 1.0.0.
## ℹ Please `group_by()` first
```

```
## Warning: There was 1 warning in `dplyr::mutate()`.
## ℹ In argument: `col = dplyr::group_indices(melted, .dots = c("variable"))`.
## Caused by warning:
## ! The `...` argument of `group_indices()` is deprecated as of dplyr 1.0.0.
## ℹ Please `group_by()` first
```

```
print(balloon_custom)
```

```
matching_rows <- apply(db.name.couple, 1, function(row) any(grepl("IL6ST", row)))
result <- db.name.couple[matching_rows, , drop = FALSE]
print(result)
```

```
##       Pair                             Family    
##  [1,] "EBI3 + IL27 / IL27RA + IL6ST"   "Cytokine"
##  [2,] "EBI3 + IL12A / IL6ST + IL12RB2" "Cytokine"
##  [3,] "CLCF1 + CRLF1 / CNTFR + IL6ST"  "Cytokine"
##  [4,] "CNTF / CNTFR + LIFR + IL6ST"    "Cytokine"
##  [5,] "IL11 / IL11RA + IL6ST"          "Cytokine"
##  [6,] "IL6 / IL6R + IL6ST"             "Cytokine"
##  [7,] "LIF / LIFR + IL6ST"             "Cytokine"
##  [8,] "OSM / LIFR + IL6ST"             "Cytokine"
##  [9,] "CTF1 / LIFR + IL6ST"            "Cytokine"
## [10,] "OSM / OSMR + IL6ST"             "Cytokine"
## [11,] "CD1B / IL6ST"                   "Cytokine"
```

```
pvalue_PC=icellnet.score.pvalue(direction="out", PC.data=PC, CC.data= CC_Pi, CC.data2   = CC_Pc, PC = c("NF_neurons", "nociceptors", "cLTMRs"), PC.target = meta, CC.type = "RNAseq",  PC.type = "RNAseq",  db = db,  between="conditions", method="BH")
pvalue.plot1=pvalue.plot(pvalue_PC[[1]], PC = c("NF_neurons", "nociceptors", "cLTMRs"))
pvalue.plot1
```

```
pvalue_FB=icellnet.score.pvalue(direction="out", PC.data=PC, CC.data= CC_Fi, CC.data2   = CC_Fc, PC = c("NF_neurons", "nociceptors", "cLTMRs"), PC.target = meta, CC.type = "RNAseq",  PC.type = "RNAseq",  db = db,  between="conditions", method="BH")
pvalue.plot1=pvalue.plot(pvalue_FB[[1]], PC = c("NF_neurons", "nociceptors", "cLTMRs"))
pvalue.plot1
```

```
pvalue1=icellnet.score.pvalue(direction="out", PC.data=PC, CC.data= CC_Pi, PC = c("NF_neurons", "nociceptors", "cLTMRs"), PC.target = meta, CC.type = "RNAseq",  PC.type = "RNAseq",  db = db,  between="cells", method="BH")
pvalue.plot1=pvalue.plot(pvalue1[[1]], PC = c("NF_neurons", "nociceptors", "cLTMRs"))
pvalue.plot1
```

```
pvalue2=icellnet.score.pvalue(direction="out", PC.data=PC, CC.data= CC_Pc, PC = c("NF_neurons", "nociceptors", "cLTMRs"), PC.target = meta, CC.type = "RNAseq",  PC.type = "RNAseq",  db = db,  between="cells", method="BH")
pvalue3=icellnet.score.pvalue(direction="out", PC.data=PC, CC.data= CC_Fi, PC = c("NF_neurons", "nociceptors", "cLTMRs"), PC.target = meta, CC.type = "RNAseq",  PC.type = "RNAseq",  db = db,  between="cells", method="BH")
pvalue4=icellnet.score.pvalue(direction="out", PC.data=PC, CC.data= CC_Fc, PC = c("NF_neurons", "nociceptors", "cLTMRs"), PC.target = meta, CC.type = "RNAseq",  PC.type = "RNAseq",  db = db,  between="cells", method="BH")
```

```
pvalue_PC[[1]]
```

```
##                   pvalue
## NF_neurons  5.608651e-06
## nociceptors 5.608651e-06
## cLTMRs      5.608651e-06
```

```
pvalue_FB[[1]]
```

```
##                  pvalue
## NF_neurons  0.002009936
## nociceptors 0.004601133
## cLTMRs      0.018545259
```

```
pvalue1[[1]]
```

```
##             NF_neurons nociceptors cLTMRs
## NF_neurons          NA          NA     NA
## nociceptors  0.2142857          NA     NA
## cLTMRs       0.5000000   0.8571429     NA
```

```
pvalue2[[1]]
```

```
##             NF_neurons nociceptors cLTMRs
## NF_neurons          NA          NA     NA
## nociceptors        0.5          NA     NA
## cLTMRs             0.5   0.6428571     NA
```

```
pvalue3[[1]]
```

```
##             NF_neurons nociceptors cLTMRs
## NF_neurons          NA          NA     NA
## nociceptors  0.2142857          NA     NA
## cLTMRs       0.5000000   0.8571429     NA
```

```
pvalue4[[1]]
```

```
##             NF_neurons nociceptors cLTMRs
## NF_neurons          NA          NA     NA
## nociceptors  0.4285714          NA     NA
## cLTMRs       0.5000000   0.8571429     NA
```

```
write.csv(lr1_Pi, "/Users/franziskadenk/Library/CloudStorage/OneDrive-SharedLibraries-King'sCollegeLondon/Denk Lab SharePoint - Mesenchymal_MS/Files_to_submit/Revisions/LR_pericytes_ipsi.csv")
write.csv(lr1_Pc, "/Users/franziskadenk/Library/CloudStorage/OneDrive-SharedLibraries-King'sCollegeLondon/Denk Lab SharePoint - Mesenchymal_MS/Files_to_submit/Revisions/LR_pericytes_contra.csv")
write.csv(lr1_Fi, "/Users/franziskadenk/Library/CloudStorage/OneDrive-SharedLibraries-King'sCollegeLondon/Denk Lab SharePoint - Mesenchymal_MS/Files_to_submit/Revisions/LR_FB_ipsi.csv")
write.csv(lr1_Fc, "/Users/franziskadenk/Library/CloudStorage/OneDrive-SharedLibraries-King'sCollegeLondon/Denk Lab SharePoint - Mesenchymal_MS/Files_to_submit/Revisions/LR_FB_contra.csv")
write.csv(db, "/Users/franziskadenk/Library/CloudStorage/OneDrive-SharedLibraries-King'sCollegeLondon/Denk Lab SharePoint - Mesenchymal_MS/Files_to_submit/Revisions/LR_database_mouse.csv")
```
